# Landscape of retron diversity across the SPIRE prokaryotic metagenome resource reveals candidate novel type XI-like lineages

**DOI:** 10.64898/2026.05.14.725207

**Authors:** Nicolás Toro

## Abstract

Retrons are prokaryotic genetic elements encoding a specialized reverse transcriptase (RT) that synthesizes multicopy single-stranded DNA (msDNA) and are increasingly recognized as components of bacterial anti-phage defense systems. However, their diversity and ecological distribution across large-scale genomic resources remain poorly characterized. Here, we surveyed retron RTs across the SPIRE representative metagenome collection, a non-redundant, species-level dataset spanning diverse microbial habitats. Using type-specific hidden Markov models cross-calibrated against all known retron groups, we identified retrons representing all canonical types together with additional divergent lineages. Retron distribution was highly structured across both taxonomy and ecology, with some lineages restricted to specific bacterial phyla and others broadly distributed across habitats. Host-associated environments accounted for the largest fraction of retron candidates, although several retron types showed clear specialization for aquatic or anthropogenic niches. Systematic novelty assessment identified two candidate type XI-like lineages, TXI_C2like and TXI_noncan_h, characterized by protease-independent architectures and lineage-specific accessory modules. De novo computational analyses further identified candidate msr/msd-like non-coding RNA elements upstream of the RT genes in both lineages, indicating that these highly divergent systems retain the defining RT-ncRNA architecture of canonical retrons. Together, these findings expand the known diversity of retron systems and reveal a strong interplay between evolutionary history and ecological specialization.

## Introduction

Retrons are widespread prokaryotic genetic elements that encode a specialized reverse transcriptase (RT) capable of producing multicopy single-stranded DNA (msDNA), a distinctive RNA–DNA hybrid molecule synthesized from a dedicated non-coding RNA (ncRNA) containing the msr and msd elements (Inouye and Inouye, 1991; Lampson et al., 1989). Originally discovered as producers of unusual DNA in myxobacteria and enteric bacteria, retrons were long regarded as molecular curiosities with unknown biological functions. This view changed dramatically with the discovery that many retrons function as components of bacterial anti-phage defense systems, acting as sensors that trigger abortive infection or toxin activation upon phage challenge (Gao et al., 2020; Millman et al., 2020; Bobonis et al., 2022). More recently, retrons have been co-opted as programmable tools for genome editing and biosensing applications, further underscoring their biological and biotechnological relevance (Schubert et al., 2021; Zhao et al., 2022).

The systematic classification of retron diversity was significantly advanced by Mestre et al. (2020), who defined 13 canonical retron types (types I–XIII) based on RT phylogeny together with the architecture of associated ncRNA elements and accessory proteins, using a curated dataset of 1,928 reference sequences spanning diverse prokaryotic lineages. This framework established that distinct retron types are characterized by specific msr/msd structures, recurrent accessory effector domains, and non-random taxonomic distributions, indicating that retron diversity has been shaped by both vertical inheritance and horizontal gene transfer across the prokaryotic tree of life. Subsequent studies have expanded the catalogue of retron-associated defense mechanisms, revealing unexpectedly rich functional diversity within this element family (Gao et al., 2020; Bernheim and Sorek, 2020; Payne et al., 2022; Bobonis et al., 2022; García-Rodríguez et al., 2025).

Despite these advances, our understanding of retron diversity remains heavily biased toward cultivated isolates and well-characterized organisms. Much of the prokaryotic diversity present in environmental and host-associated microbiomes remains underrepresented in reference databases, limiting the discovery of divergent retron lineages and obscuring the full ecological breadth of these elements. Metagenomic approaches offer a powerful complement to culture-dependent surveys; however, the systematic application of retron detection pipelines to large-scale genome-resolved metagenomic resources has not yet been reported. Key questions therefore remain unresolved: how widespread are retron systems across the prokaryotic biosphere? How are canonical retron types distributed across taxonomic groups and environments? To what extent does unexplored microbial diversity harbor previously unrecognized retron lineages?

The Searchable planetary-scale microbiome REsource (SPIRE) database provides an unprecedented resource for addressing these questions. SPIRE comprises 107,078 non-redundant representative genomes, including both metagenome-assembled genomes (MAGs) and isolate representatives, spanning aquatic, terrestrial, host-associated, and anthropogenic environments (Schmidt et al., 2024). By integrating genome-resolved assemblies with standardized taxonomic and ecological annotations, SPIRE enables systematic exploration of genetic element diversity across the global microbiome.

Here, we present a systematic survey of retron RT diversity across the SPIRE resource using a panel of type-specific hidden Markov model (HMM) profiles calibrated against all canonical retron groups defined by Mestre et al. (2020). We identified 1,286 retron candidates spanning all 13 canonical types, characterized their taxonomic and ecological distributions, and implemented a novelty prioritization framework to identify lineages with divergent architectures. This analysis revealed two candidate type XI-like lineages, TXI_C2like and TXI_noncan_h, within retron RT clade 8, distinguished by protease-independent architectures and distinct lineage-specific accessory modules. The *de novo* computational identification of candidate msr/msd-like ncRNA elements in both lineages further indicates that these highly divergent systems retain the fundamental organizational features of canonical retrons.

## Materials and Methods

### Genome dataset and protein prediction

A total of 107,078 representative prokaryotic genomes from the SPIRE v01 dataset (Schmidt et al., 2024) were analysed, comprising one genome per species-level cluster defined at approximately 95% average nucleotide identity. Of these, 92,134 were metagenome-assembled genomes (MAGs), whereas the remaining 14,944 were isolate representatives derived from linked ProGenomes clusters. Protein-coding sequences were predicted *de novo* from all assemblies using Prodigal v2.6.3 (Hyatt et al., 2010) in metagenomic mode (-p meta), yielding approximately 272 million predicted proteins.

Taxonomic lineages (phylum to species) were assigned by mapping genome identifiers to their corresponding SPIRE species-level clusters, including both ProGenomes-linked specI_v4_* clusters and *de novo* 95% ANI clusters (spire_v1_095_*), using the SPIRE genome metadata file (spire_v1_genome_metadata.tsv). Full lineage strings were then reconstructed using taxonkit v0.14 (Shen and Ren, 2021) against the NCBI taxonomy database.

Ecological metadata were assigned by linking each candidate contig to its originating sample through the derived_from_sample field and querying the SPIRE microntology (spire_v1_microntology.tsv), a controlled vocabulary that provides structured, multi-tag environmental annotations for each sequenced sample. When multiple annotations were available for a given sample, the most specific descriptor was retained. Sample-level descriptors were subsequently collapsed into seven high-level environmental categories derived from the SPIRE microntology: host-associated, aquatic, terrestrial, anthropogenic, temperature-defined, salinity-defined, and pH-defined environments.

### HMM construction, candidate detection and classification

A panel of 28 type-specific hidden Markov model (HMM) profiles was constructed to cover all 13 known retron types (types I–XIII and their recognized subgroups), as well as the clade 11 and clade2_Ec107like lineages. For each group, reference sequences from Mestre et al. (2020) were aligned using MAFFT v7 (Katoh and Standley, 2013), and profile HMMs were generated using hmmbuild from HMMER v3.3 (Eddy, 2011).

Profiles were empirically cross-calibrated to define group-specific classification thresholds. For each HMM, two score distributions were calculated: (i) self-hit scores, obtained by searching the profile against its own reference sequences and recording the minimum score (pos_min), and (ii) cross-hit scores, obtained by searching the profile against all reference sequences from the remaining groups and recording the maximum score (neg_max). The classification threshold for each group was defined as the midpoint between these values, calculated as (pos_min + neg_max) / 2. Thresholds ranged from 177.2 bits for type I-B2 to 317.5 bits for clade11-A (**Supplementary Table S1**).

The approximately 272 million predicted proteins were first screened using hmmsearch (HMMER v3.3) against a general retron RT HMM constructed from the 1,928 reference RT sequences reported by Mestre et al. (2020). Searches were performed with an E-value cutoff of 1 × 10⁻¹⁰, yielding 31,280 initial hits. Sequences with bit scores ≥160 bits were retained as candidate retron RTs (n = 1,319).

The 160-bit cutoff was selected empirically based on the score distribution of the 1,928 reference RTs against the general HMM. The lowest-scoring reference sequence yielded 114.5 bits, corresponding to a highly divergent type II RT. At the 160-bit threshold, 195 reference sequences (10.1%) fell below the cutoff, predominantly from type VI (n = 91), type I-B2 (n = 36), type I-B1 (n = 34), and type IV (n = 20). This threshold therefore represents a conservative trade-off between sensitivity and specificity and likely results in underrepresentation of the most divergent retron lineages.

Among the 1,319 candidates, 33 sequences carried contig identifiers in k* format that lacked assembly sequences, gene annotations, and associated metadata were excluded from downstream biological analyses but were retained for phylogenetic placement to preserve overall tree context.

The remaining 1,286 SPIRE-native candidates were classified by scoring each sequence against the 28 type-specific HMMs using hmmsearch. Each candidate was assigned to the group producing the highest bit score. The overall computational workflow used for retron identification, classification, and novelty assessment is summarized in **Supplementary Figure S1**.

Confidence tiers were assigned based on three independent criteria: (i) HMM detection strength, assessed relative to the group-specific classification threshold; (ii) RT domain completeness, based on conservation of the canonical RT1–RT7 motif core; and (iii) consistency of phylogenetic placement with established retron clades using EPA-ng (Barbera et al., 2019). High-confidence candidates exceeded the group-specific threshold, displayed near-complete RT1–RT7 architecture, and showed coherent phylogenetic placement within or immediately adjacent to established retron clades. Low-confidence candidates showed detectable but weaker support (bit score ≥160 bits but below the group-specific threshold) and/or partial RT domain coverage, with broadly compatible but less well-supported phylogenetic placement. Borderline candidates had scores near the overlap zone between self-hit and cross-hit distributions. These were retained for completeness and included in genomic neighbourhood analyses but excluded from high-confidence biological interpretations.

### Phylogenetic reference tree and placement of SPIRE candidates

A maximum-likelihood reference tree was inferred from the 1,928 retron RT sequences compiled by Mestre et al. (2020). Reference sequences were aligned using MAFFT v7 (Katoh and Standley, 2013), and the resulting alignment was trimmed to the 203 positions corresponding to the conserved RT1–RT7 motif core by removing columns containing more than 50% gaps. The trimmed alignment was used to infer a maximum-likelihood tree with IQ-TREE 2 v2.3.2 (Minh et al., 2020). The best-fitting substitution model was selected using ModelFinder and corresponded to Q.pfam+F+I+G4. Branch support was assessed with 1,000 ultrafast bootstrap replicates.

SPIRE candidate RTs were incorporated into the reference alignment using MAFFT with the --add option (Katoh and Frith, 2012), preserving the original alignment coordinates. Phylogenetic placement was performed using EPA-ng (Barbera et al., 2019). For each candidate, the placement with the highest likelihood weight ratio (LWR) was retained. Placements were then grafted onto the reference topology using Gappa (Czech et al., 2020), and the resulting combined tree was visualized and annotated in Interactive Tree Of Life (iTOL) v6 (Letunic and Bork, 2024).

### Covariance model searches for msr/msd detection

The presence of msr/msd non-coding RNA elements in the genomic neighbourhood of each RT candidate was assessed using cmsearch from Infernal v1.1 (Nawrocki and Eddy, 2013) with a panel of 21 covariance models derived from experimentally validated msr/msd transcripts representing retron types for which structural information is available (Mestre et al., 2020). Searches were performed against genomic regions extending 10 kb upstream and downstream of each RT locus extracted from the SPIRE genome assemblies.

Covariance model searches were applied to all high-confidence candidates (n = 934), of which 540 (57.8%) yielded a confident msr/msd assignment. The most frequently detected models corresponded to Ec107_like, clade11-A, TypeIC1_IC2, TypeIA_IIAI, and clade11-B. Retron types lacking experimentally validated msr/msd structures, including types VI, VII, VIII, X, XI, and XII were not represented in the covariance model panel and therefore could not be annotated by this analysis. Accordingly, the absence of a covariance model hit in these groups should not be interpreted as evidence for the absence of an associated ncRNA. For the clade 11 lineage, two distinct msr/msd structural families, designated A and B by Mestre et al. (2020), were used to subdivide candidates into clade11-A (model A detected; n = 71), clade11-B (model B detected; n = 39), and clade11 (no confident hit to either model; n = 9).

### Genomic neighborhood analysis

Coding sequences neighbouring each RT candidate were identified from the Prodigal gene annotations (GFF format) associated with each SPIRE genome. For each RT locus, genes located on the same contig were ordered according to genomic coordinates, and neighbouring coding sequences were extracted relative to the RT position using custom Python scripts.

A broad ±5-gene window was initially used to capture the local genomic context and provided complete coverage for all 1,286 SPIRE-native candidates. For accessory protein analyses, a more stringent ±2-gene window, excluding the RT itself, was used to define tight genomic neighbourhoods. This approach retained complete neighbourhood information for 1,284 of 1,286 loci while minimizing the inclusion of unrelated genes in gene-dense regions.

Protein sequences from these tight neighbourhoods were clustered by sequence similarity using MMseqs2 (Steinegger and Söding, 2017) with thresholds of at least 30% amino acid identity and 80% alignment coverage. Cluster representatives were functionally annotated using hmmscan from HMMER v3.3 against Pfam-A v35.0 (Mistry et al., 2021) with an E-value cutoff of 1 × 10⁻⁵.

In parallel, cluster representatives were screened against a curated collection of HMMs representing known retron-associated effector families (Mestre et al., 2020) using the same significance threshold. This complementary analysis enabled the detection of functionally relevant accessory modules that were not always fully captured by general Pfam annotations.

For clusters lacking confident Pfam or effector assignments, structural homology searches were performed using HHsearch against the PDB70 database. Candidate matches were evaluated based on HHsearch probability scores and alignment quality.

### Taxonomic annotation, ecological classification, and statistical analyses

Taxonomic lineages (phylum to species) were assigned by mapping genome identifiers to their corresponding SPIRE species-level clusters using the SPIRE genome metadata file. Full lineage strings were reconstructed using taxonkit v0.14 (Shen and Ren, 2021) against the NCBI taxonomy database. Ecological metadata were assigned by linking each candidate contig to its originating sample through the SPIRE microntology (spire_v1_microntology.tsv), collapsing sample-level descriptors into seven high-level environmental categories: host-associated, aquatic, terrestrial, anthropogenic, temperature-defined, salinity-defined, and pH-defined environments.

Non-random associations between retron groups and bacterial phyla, and between retron groups and ecological biomes, were assessed using Pearson’s chi-squared test of independence applied to contingency tables of group × phylum and group × biome, respectively, retaining only groups with n ≥ 10 and phyla or biomes with n ≥ 5. Chi-squared statistics and p-values were computed using chi2_contingency (SciPy v1.11; Virtanen et al., 2020). Pearson standardized residuals were calculated as (O − E) / √E; residuals exceeding |2| were considered indicative of notable enrichment or depletion. Taxonomic breadth of each retron group was quantified using the Gini–Simpson diversity index (1 − D), where D = Σpᵢ² and pᵢ is the proportion of candidates from phylum i; values of 0 and 1 indicate complete restriction to a single phylum and maximum evenness across phyla, respectively.

Statistical associations between accessory protein clusters and retron groups were quantified using two-sided Fisher’s exact test, comparing cluster frequency within each target group against all remaining groups. Odds ratios were computed as (a × d) / (b × c); clusters exclusively present in one group received OR = ∞. Multiple testing correction used the Benjamini–Hochberg false discovery rate procedure (Benjamini and Hochberg, 1995), implemented via SciPy v1.11 (Virtanen et al., 2020). Clusters with FDR < 0.05 and OR > 2 were considered significantly enriched. All statistical analyses were implemented in Python v3.11 using NumPy v1.24, SciPy v1.11, and pandas v2.0.

### De novo identification of candidates ncRNA structures

For *de novo* discovery of candidate retron ncRNAs, the 600 bp region immediately upstream of the RT start codon was extracted from all TXI_noncan_h (n = 28) and TXI_C2like (n = 120) loci using custom Python scripts. Conserved RNA secondary structures were initially identified using CMfinder 0.4 (Yao et al., 2006) with the parameters -f 0.5 -s1 2 -s2 3 -minspan1 80 -maxspan1 300 -minspan2 80 -maxspan2 300 -combine.

Covariance models were constructed using cmbuild and calibrated with cmcalibrate from Infernal v1.1.5 (Nawrocki and Eddy, 2013). Positional validation was performed by searching the resulting models against genomic regions extending 10 kb upstream and downstream of each RT locus using cmsearch with an E-value threshold of 0.01.

Structure-guided multiple sequence alignments were generated from the 300 bp immediately upstream of the RT start codon using mLocARNA 2.0.1 (Will et al., 2007) with the parameters --plfold-span 400 --min-prob 0.03 --indel -4 --indel-open -400 --max-diff 100 --max-diff-am -60 --alifold-cons.

Covariation significance above phylogenetic expectation was assessed using R-scape 2.0.4 (Rivas et al., 2017) with an E-value threshold of 0.05.

## Results

### Global survey of retron diversity across the SPIRE genome resource

To systematically identify retron reverse transcriptases (RTs) across the SPIRE genome resource, we screened the predicted proteomes of 107,078 representative genomes using a general retron RT HMM and classified all hits with a panel of 28 cross-calibrated type-specific HMM profiles. This approach yielded 31,280 initial RT-like hits, of which 1,319 exceeded the general score threshold. After excluding 33 candidates lacking associated genomic context, a total of 1,286 SPIRE-native candidates were retained for downstream analyses. These candidates were distributed across 1,187 unique genomes, indicating that retrons are detectable in approximately 1.1% of the species-level representatives included in SPIRE (**Supplementary Table S2**).

Candidates were assigned to confidence tiers based on HMM support, RT domain completeness, and phylogenetic consistency. Of the 1,286 candidates, 934 were classified as high-confidence (72.6%), 25 as borderline (1.9%), and 327 as low-confidence (25.4%). Phylogenetic placement using EPA-ng (Barbera et al., 2019) within a maximum-likelihood reference tree built from 1,928 curated retron RTs confirmed that all candidates fell within established retron phylogenetic diversity (**Figure 1**).

**Figure 1.**
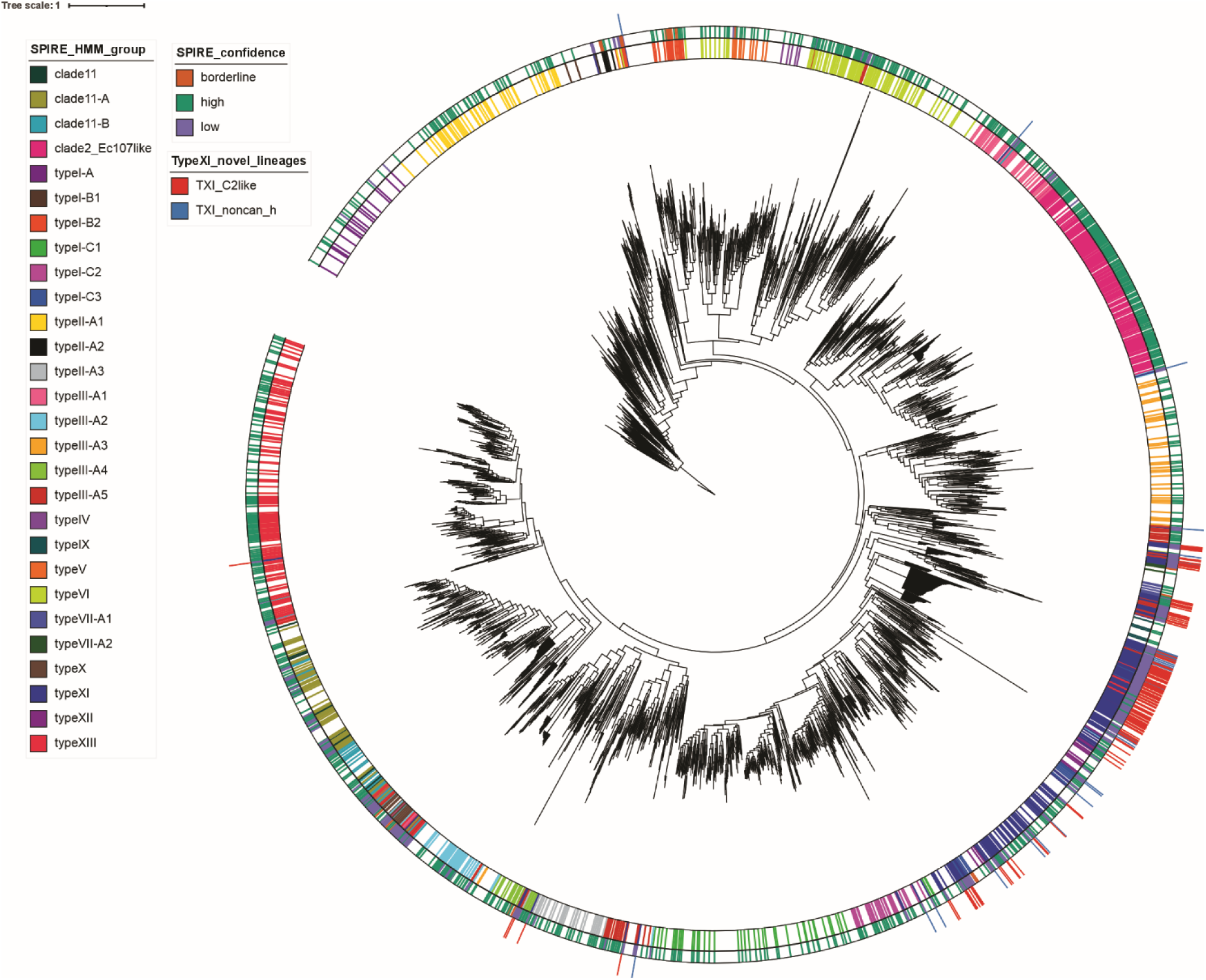
Phylogenetic placement of SPIRE-derived retron RTs in the global retron landscape. Circular maximum-likelihood phylogeny of retron reverse transcriptases (RTs) inferred from a curated reference dataset of 1,928 sequences compiled by Mestre et al. (2020). RT candidates identified in the SPIRE genome resource were incorporated into this framework by phylogenetic placement using EPA-ng (Barbera et al., 2019). The tree is shown using the original rooting of the reference phylogeny. Three concentric annotation rings summarize the main attributes of SPIRE-derived candidates. The innermost ring (SPIRE_HMM_group) indicates the HMM-based classification of each sequence into established retron groups and subgroups. The middle ring (SPIRE_confidence) denotes the confidence tier assigned to each candidate (high, borderline, or low) based on HMM support, RT motif conservation, and phylogenetic consistency. The outermost ring (TypeXI_novel_lineages) highlights sequences belonging to the two noncanonical Type XI-like lineages identified in this study, designated TXI_C2like and TXI_noncan_h, which are analyzed in detail in Figures 4 and **5**. SPIRE-derived RTs are distributed across the full breadth of the reference retron phylogeny, substantially expanding previously defined clades. Most candidates assigned to the novel Type XI-like lineages map within the reference Type XI region, although a small number of outlying placements are observed, likely reflecting uncertainty in the placement of highly divergent RT core sequences. Overall, the distribution of SPIRE candidates reveals extensive hidden retron diversity across known groups and highlights two previously unrecognized Type XI-associated radiations.

All 13 canonical retron types described by Mestre et al. (2020) were detected, together with the clade11 and clade2_Ec107like lineages, yielding 28 classification groups (**Figure 2A**, **Table 1**, and **Supplementary Table S1**). The most abundant group was type XI-like (n = 232, 18.0%), although 72% of these candidates were assigned to the low-confidence tier, well above the genome-wide average of 25.4%, indicating substantial sequence divergence within this lineage. The second most abundant group was clade2_Ec107like (n = 192, 14.6%), of which 98% were high-confidence, enabling clear separation from its sister lineage, type III-A1. Type XIII (n = 143, 11.1%) and type VI (n = 93, 7.2%) were the third and fourth most abundant groups, respectively, and were predominantly composed of high-confidence candidates. Each of the remaining 24 groups accounted for less than 6% of the total candidate set.

**Figure 2.**
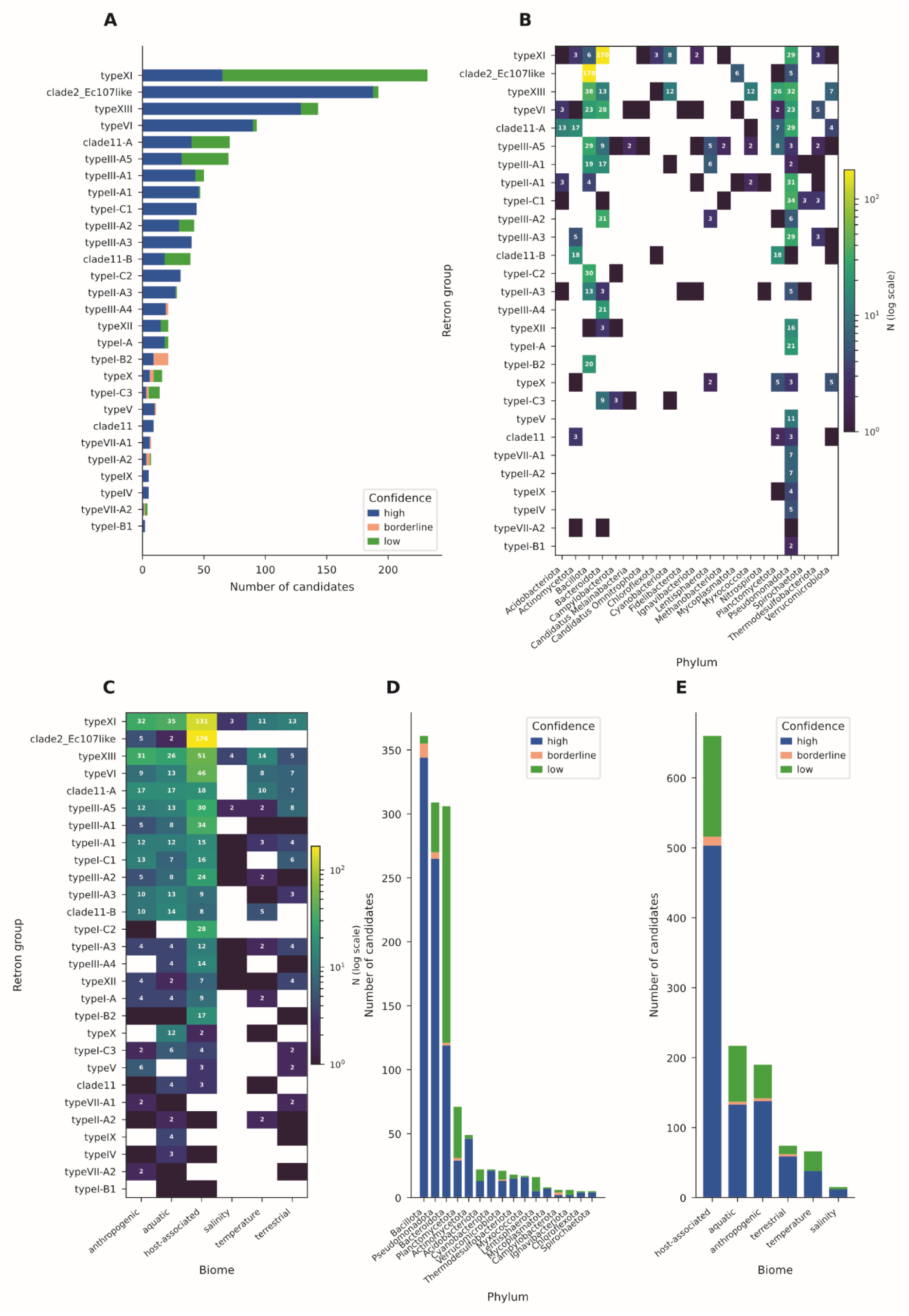
Global distribution of SPIRE-derived retron candidates across retron groups, bacterial phyla, and ecological biomes. **(A)** Distribution of retron candidates by classification group and hidden Markov model (HMM) confidence tier. Type XI-like retrons (n = 232) were the most abundant group but were dominated by low-confidence candidates (n = 167, 72%), in contrast to clade2_Ec107like (n = 192), which was almost entirely composed of high-confidence candidates (n = 188, 98%). Type XIII (n = 143), type VI (n = 93), and clade11-A (n = 71) were the next most abundant groups. Groups with notable borderline fractions included type I-B2 and type III-A4. **(B)** Distribution of retron groups across bacterial phyla (color scale indicates the log-transformed number of candidates). Type XI-like retrons were concentrated in Bacteroidota (170 of 232 candidates), whereas clade2_Ec107like was predominantly associated with Bacillota (178 of 192 members). Type XIII showed the broadest phylogenetic distribution, spanning Bacillota (n = 38), Pseudomonadota (n = 32), Planctomycetota (n = 26), and Bacteroidota (n = 13). Type VI was distributed across Bacteroidota (n = 28), Bacillota (n = 23), and Pseudomonadota (n = 23). Clade11-A was enriched in Pseudomonadota (n = 29), with additional representation in Actinomycetota (n = 17) and Acidobacteriota (n = 13). **(C)** Distribution of retron groups across high-level environmental categories (color scale indicates the log-transformed number of candidates). clade2_Ec107like showed the strongest association with host-associated environments (176 of 192 candidates, 91%). Type XI-like retrons were also predominantly assigned to host-associated environments (131 of 232 candidates, 56%) but were additionally detected in aquatic (n = 35) and anthropogenic (n = 32) categories. Type XIII displayed a broad environmental distribution, whereas type X was predominantly associated with aquatic environments (12 of 16 candidates). **(D)** Global phylum distribution stratified by HMM confidence tier. Bacillota and Pseudomonadota were dominated by high-confidence candidates (95% and 86%, respectively), consistent with the prevalence of well-characterized retron groups in these phyla. In contrast, Bacteroidota showed the lowest proportion of high-confidence candidates (39%), driven by the large contribution of low-confidence type XI-like retrons. Planctomycetota also exhibited an elevated low-confidence fraction (56%). **(E)** Global distribution of retron candidates across high-level environmental categories stratified by HMM confidence tier. Host-associated environments contained the largest number of retron candidates (n = 660, 51.3%), of which 76% were classified as high confidence. Aquatic and temperature-defined categories showed elevated low-confidence fractions (39% and 42%, respectively), consistent with the increased representation of type XI-like retrons in these categories.

**Table 1.**
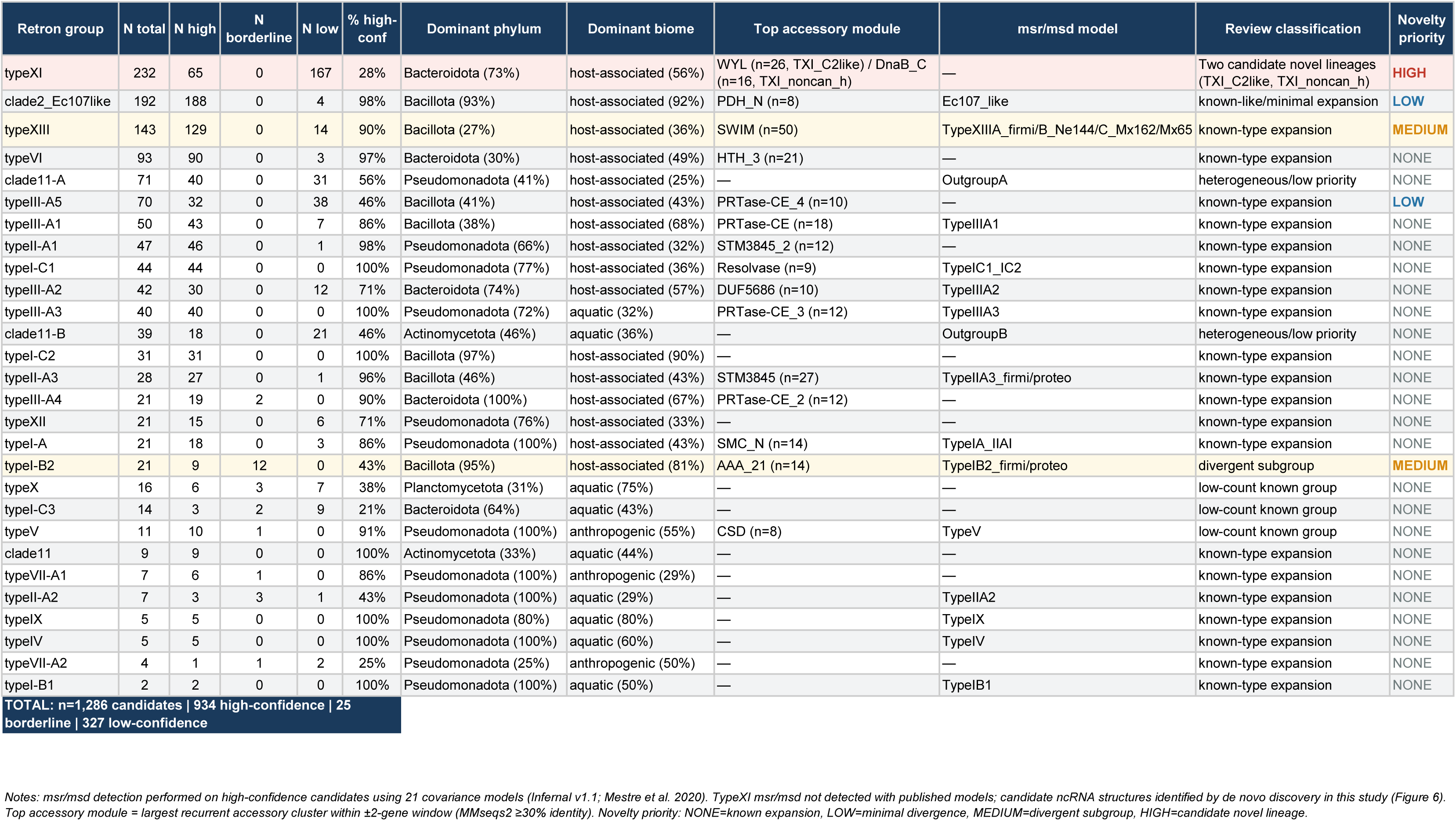
Summary of retron candidates identified across the SPIRE prokaryotic resource.

Within clade11, covariance model searches using the two known msr/msd structural families defined by Mestre et al. (2020) resolved three subgroups: clade11-A (n = 71), clade11-B (n = 39), and clade11 (n = 9), in which neither covariance model yielded a confident hit.

Retrons were detected across 13 bacterial phyla (**Figure 2B**). Retron type–phylum associations were highly significant (*χ*² = 2,764.8, df = 375, p < 10⁻³⁰⁰), and Pearson standardized residuals highlighted strong lineage-specific enrichments and depletions across phyla (**Supplementary Figure S2A**). The most represented phyla were Bacillota (n = 361), Pseudomonadota (n = 309), and Bacteroidota (n = 306), which together accounted for 75.9% of all candidates. However, confidence distributions differed markedly among these phyla. Bacillota and Pseudomonadota were dominated by high-confidence candidates (95% and 86%, respectively), reflecting the prevalence of well-characterized retron groups. By contrast, Bacteroidota showed a substantially lower high-confidence fraction (39%), driven by the predominance of type XI-like candidates. Indeed, 170 of the 232 type XI-like retrons were identified in Bacteroidota, where they represented the dominant retron lineage.

Quantitative analysis of taxonomic specificity using the Gini-Simpson diversity index (1 − D) revealed a continuum ranging from highly restricted to broadly distributed lineages (**Supplementary Figure S2B**). The most taxonomically restricted groups were clade2_Ec107like (1 − D = 0.139, 93% Bacillota), type I-C2 (1 − D = 0.062, 97% Bacillota), and type III-A4 (1 − D = 0, exclusively Bacteroidota). In contrast, type XIII (1 − D = 0.821), type VI (1 − D = 0.782), and type III-A5 (1 – D = 0.788) exhibited the broadest taxonomic distributions.

Retron-containing genomes were most frequently assigned to the host-associated category (n = 660, 51.3%), followed by aquatic (n = 217, 16.9%) and anthropogenic (n = 190, 14.8%) environments (**Figure 2C** and **2E**). This pattern was driven primarily by clade2_Ec107like, of which 91% of candidates were assigned to the host-associated category, and by type XI-like retrons, for which 55% of candidates received the same annotation. Together, these two groups accounted for nearly half of all host-associated candidates. In contrast, type X was most frequently assigned to aquatic environments (75%), whereas type V was most commonly associated with anthropogenic environments (55%). Retron type–biome associations were also highly significant (χ² = 453.3, df = 125, p = 9.2 × 10⁻³⁹; **Figure 2C**), demonstrating that ecological specialization is a pervasive feature of retron diversity.

Together, these results establish that retrons are broadly distributed across the bacterial domain but exhibit pronounced lineage-specific patterns of taxonomic and ecological specialization. In particular, the concentration of type XI-like retrons in Bacteroidota and the strong host association of clade2_Ec107like suggest that both evolutionary history and environmental context have played major roles in shaping retron diversification.

### Systematic evaluation indicates that most SPIRE retrons represent expansions of known lineages

To distinguish expanded diversity from genuine biological novelty, we systematically evaluated all 28 retron classification groups using five complementary criteria. RT phylogenetic coherence assessed whether candidates assigned to a given group formed a discrete and well-supported cluster in both the EPA-ng placement tree (**Figure 1**) and an independent maximum-likelihood reconstruction. Recurrence and specificity of accessory protein architecture evaluated whether one or more non-RT modules were consistently associated with the group and statistically enriched relative to other retron groups. Confidence class distribution reflected the proportion of high-, borderline-, and low-confidence candidates, providing a measure of how closely each group conformed to the calibrated HMM profiles. Taxonomic coherence assessed the extent to which a group was restricted to specific bacterial lineages. Ecological coherence assessed whether candidates showed non-random enrichment in particular biomes beyond that expected from host taxonomy alone. Each criterion was scored as absent (−), weak (+), moderate (++), or strong (+++), and the combined evidence was used to assign an overall novelty priority score of none, low, medium, or high (**Supplementary Table S3**).

The results are summarized in a curated evidence matrix highlighting representative lineages that span the full spectrum of observed patterns, from canonical expansions and divergent subgroups to candidate novel lineages (**Figure 3A** and **3B**). The analysis was performed at the level of curated biological lineages rather than the original HMM classification groups. Groups displaying uniformly canonical characteristics across all five criteria were omitted from the matrix because they provided limited discriminatory information.

**Figure 3.**
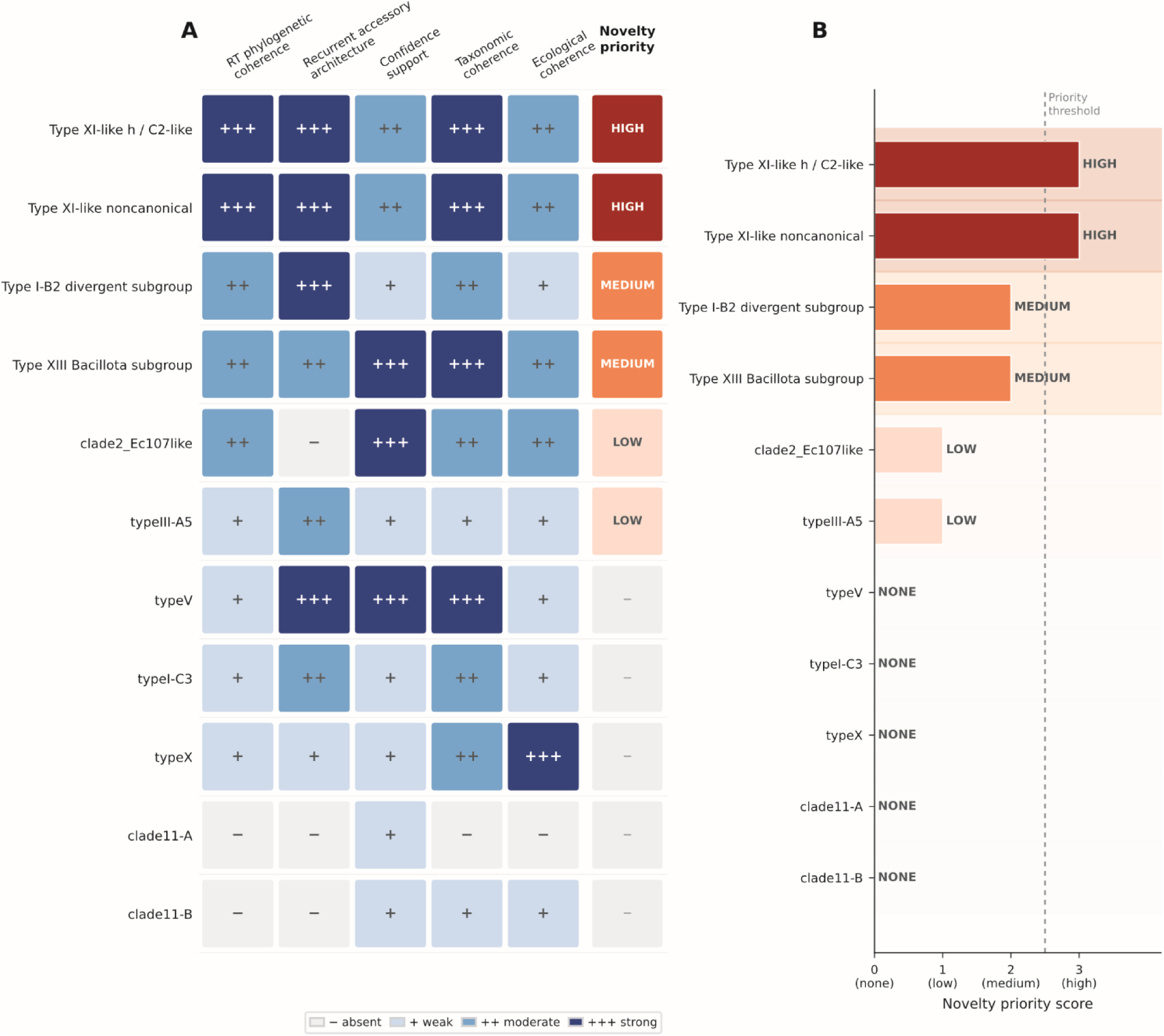
Systematic evaluation of SPIRE retron groups and novelty prioritization. **(A)** Curated evidence matrix summarizing the five criteria used to evaluate representative SPIRE retron lineages. Columns correspond to RT phylogenetic coherence, recurrence and specificity of accessory protein architecture, confidence class distribution, taxonomic coherence, and ecological coherence. Each criterion was scored on a four-point scale: absent (−), weak (+), moderate (++), or strong (+++). The composite novelty priority score (none/low/medium/high) is shown in the rightmost column. TXI_C2like and TXI_noncan_h were the only lineages assigned strong (+++) scores for RT phylogenetic coherence, recurrent accessory architecture, and taxonomic coherence simultaneously, and were prioritized as the strongest candidates for previously unrecognized retron lineages. In contrast, clade2_Ec107like showed strong (+++) confidence support and moderate (++) taxonomic coherence but was scored absent (−) for recurrent accessory architecture, reflecting the lack of non-RT accessory modules in this lineage. **(B)** Composite novelty priority scores assigned to evaluated lineages after integration of all five criteria. TXI_C2like and TXI_noncan_h were the only lineages assigned high novelty priority. The type XIII *Bacillota* subgroup and type I-B2 divergent subgroup reached medium priority. All remaining lineages received low or no novelty priority.

Overall, 21 of the 28 retron classification groups were interpreted as expansions of previously described retron lineages. Three additional groups (Types X, I-C3, and V) were classified as low-count representatives of known retron types. Clade2_Ec107like was considered a minimally divergent expansion of a previously described lineage, whereas clade11-A and clade11-B were analyzed separately but assigned low novelty priority because they lacked consistent distinguishing features. Only the type XI-like classification group, which belongs to RT Clade 8, yielded lineages with strong evidence of genuine biological novelty. Detailed analysis of its internal phylogenetic and architectural structure identified two candidate novel sublineages, TXI_C2like and TXI_noncan_h (**Figure 3A** and **3B**).

Several contrasting examples illustrate the framework’s discriminatory power. Type V showed strong accessory architecture (+++) and high confidence support (+++), yet was assigned no novelty priority because its organization fully conformed to the canonical cold-shock DNA-binding domain (CSD)-associated type V architecture, demonstrating that strong individual signals do not necessarily indicate biological novelty. Type I-B2 and the Bacillota-specific type XIII subgroup were assigned a medium priority as divergent subgroups within the established retron types. Type I-B2 combined moderate phylogenetic divergence with a distinctive ATPase–TOPRIM accessory module, whereas the Bacillota type XIII subgroup exhibited restricted taxonomic distribution and recurrent SWIM_2/DUF6849 remodelling consistent with clade-level diversification.

In contrast, clade11-A, clade11-B, type III-A5, type X, and type I-C3 displayed weak or inconsistent evidence and were assigned low or no novelty priority. Notably, clade2_Ec107like received low novelty priority despite its abundance (n = 192), consistent with its canonical phylogenetic placement and the absence of recurrent non-RT accessory modules.

Collectively, this systematic evaluation identified TXI_C2like and TXI_noncan_h as the strongest candidates for previously unrecognized retron biology, motivating their detailed characterization in the following sections.

### Type XI non-canonical lineages encode two distinct stand-alone accessory architectures

Canonical type XI retrons encode a single open reading frame in which the reverse transcriptase (RT) domain is fused to a C-terminal trypsin-like serine protease domain. Analysis of the 232 type XI-like candidates identified in SPIRE revealed that 207 loci (89.2%) lacked detectable C-terminal protease domains. Of these, 184 (88.9%) retained the conserved Region X (NAxxH) and Region Y (VTG) motifs characteristic of retron RTs and were distributed across 114 taxa. Systematic analysis of their genomic architectures and phylogenetic placement identified two recurrent non-canonical lineages, TXI_C2like (n = 126) and TXI_noncan_h (n = 29), in which the canonical protease fusion was replaced by distinct stand-alone accessory proteins encoded as separate open reading frames within the ±2-gene neighborhood of the RT (**Figure 4A**).

**Figure 4.**
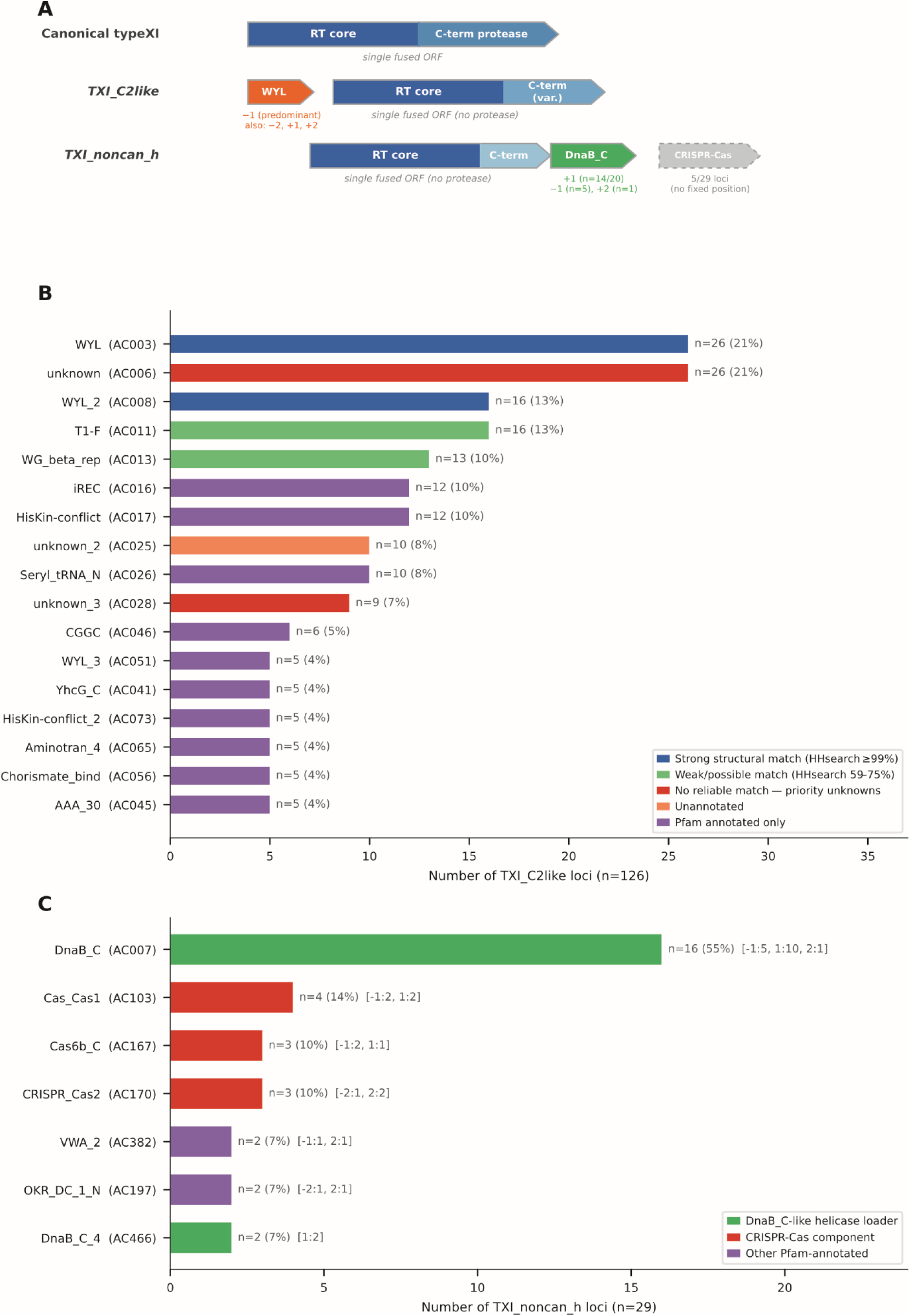
Genomic architecture and accessory protein repertoires of two noncanonical type XI-like lineages. (**A**) Schematic representation of the genomic organization of canonical type XI retrons and the two non-canonical lineages identified in this study. Canonical type XI retrons encode a single open reading frame in which the reverse transcriptase (RT) domain is fused to a C-terminal trypsin-like serine protease. In TXI_C2like, the RT lacks the protease domain and is encoded as a single ORF with a variable C-terminal extension, whereas accessory WYL-domain proteins are encoded as separate ORFs positioned upstream of the RT (offset −1, predominant) or at alternative positions within the ±2-gene neighborhood (offsets −2, +1, and +2). TXI_noncan_h similarly lacks protease fusion and is associated with DnaB_C-containing proteins encoded as separate ORFs, most frequently at the immediate downstream position (offset +1; 14 of 20 loci), although upstream positions also occur (offset −1; 5 of 20 loci). A subset of TXI_noncan_h loci was additionally associated with nearby CRISPR-Cas genes (5 of 29 loci, 17.2%), shown as a dashed arrow indicating absence of a fixed genomic position. Solid arrows indicate the predominant genomic organization. **(B)** Recurrent accessory protein clusters associated with TXI_C2like (n=126). Bars indicate the number of loci in which each cluster was detected. Colors denote annotation confidence: blue, strong structural assignment by HHsearch (probability ≥99%); green, tentative structural assignment (probability 59–75%); red, no reliable structural match (priority novel unknowns); purple, Pfam-based annotation only. The dominant clusters corresponded to WYL-family proteins (AC003, AC008, and AC051; collectively detected in 47 loci, 37.3%) and the uncharacterized cluster AC006 (n=26 loci). **(C)** Recurrent accessory protein clusters associated with TXI_noncan_h (n = 29 curated loci). DnaB_C-containing proteins were detected in 20 loci (69.0%) and were dominated by AC007, with AC466 representing a secondary recurrent cluster. Two additional singleton clusters (AC377 and AC1199), each detected in a single locus, contributed to the overall total but are not shown. Positional offsets relative to the RT are indicated in brackets for each displayed cluster. CRISPR-Cas-associated clusters shown in the panel include AC103, AC167 and AC170; together with additional singleton CRISPR-Cas-associated clusters not shown, these occurred in 5 loci (17.2%). In several loci, multiple Cas components co-occurred, consistent with the presence of partial or complete CRISPR-Cas systems adjacent to the retron RT.

TXI_C2like loci (n = 126; 11 high-confidence and 115 low-confidence candidates; 112 motif-compatible) were characterized by a recurrent association with WYL-domain proteins. WYL-associated modules were detected in 47 loci (37.3%) and grouped into three enriched accessory clusters: AC003 (WYL; n = 26 loci), AC008 (WYL_2; n = 16), and AC051 (WYL_3; n = 5). These proteins were found both upstream and downstream of the RT (offsets −2, −1, +1, and +2), indicating a flexible local organization rather than a fixed operon structure (**Figure 4A**). HHsearch against the pdb70 database confirmed strong structural similarity for the two dominant clusters: AC003 matched 7P5X_AX (probability = 99.6%, E = 5.1 × 10⁻¹⁹) and AC008 matched 7P5X_AY (probability = 99.7%, E = 4.2 × 10⁻²²), both corresponding to the WYL superfamily fold, a nucleic acid-binding regulatory domain frequently associated with bacterial defense (Makarova et al., 2014; Müller et al., 2019; Keller and Weber-Ban, 2023). In addition to WYL proteins, TXI_C2like displayed a broader accessory repertoire comprising eight significantly enriched protein clusters relative to all type XI-like loci (Fisher’s exact test, FDR < 3.6 × 10⁻⁸; **Table 2**). Among these, AC006 (n = 26 loci; OR = ∞) encoded a protein lacking detectable Pfam domains or structural homologs, making it the most abundant uncharacterized accessory module associated with this lineage and a priority target for functional investigation (**Figure 4B**; **Table 2**).

**Table 2.** Accessory protein clusters significantly associated with TXI_C2like and TXI_noncan_h retron sublineages.

| AC | Label | Pfam annotation | Sublineage | N loci | OR | FDR | Note |
| --- | --- | --- | --- | --- | --- | --- | --- |
| AC003 | WYL | WYL | TXI_C2like | 26 | ∞ | $1.7 \times 10^{-21}$ | |
| AC006 | unknown | — | TXI_C2like | 26 | ∞ | $4.8 \times 10^{-20}$ | *a* |
| AC008 | WYL_2 | WYL | TXI_C2like | 16 | ∞ | $5.1 \times 10^{-17}$ | |
| AC011 | T1-F | T1-F | TXI_C2like | 16 | ∞ | $7.8 \times 10^{-15}$ | |
| AC013 | WG_beta_rep | WG_beta_rep | TXI_C2like | 13 | ∞ | $7.5 \times 10^{-12}$ | |
| AC016 | iREC | iREC | TXI_C2like | 12 | 62.4 | $2.5 \times 10^{-9}$ | |
| AC017 | HisKin-conflict | HisKin-conflict | TXI_C2like | 12 | 62.4 | $2.5 \times 10^{-9}$ | |
| AC026 | Seryl_tRNA_N | Seryl_tRNA_N | TXI_C2like | 10 | ∞ | $3.6 \times 10^{-8}$ | |
| AC007 | DnaB_C | DnaB_C | TXI_noncan_h | 16 | 14.7 | $1.6 \times 10^{-9}$ | |
| AC103 | Cas_Cas1 | Cas1 (CRISPR) | TXI_noncan_h | 4 | ∞ | $7.0 \times 10^{-4}$ | *b* |
| AC167 | Cas6b_C | Cas6b (CRISPR) | TXI_noncan_h | 3 | ∞ | $2.0 \times 10^{-3}$ | *b* |
| AC170 | CRISPR_Cas2 | Cas2 (CRISPR) | TXI_noncan_h | 3 | ∞ | $2.0 \times 10^{-3}$ | *b* |

OR = odds ratio from Fisher's exact test with Benjamini-Hochberg FDR correction. ∞ = cluster exclusive to sublineage. N loci = number of typeXI RT loci carrying the accessory cluster within a $\pm 2$ -gene window.
<sup>a</sup> AC006 encodes a protein of unknown function with no detectable Pfam domain or structural homology; priority novel module for functional characterization.
<sup>b</sup> AC103 (Cas1), AC167 (Cas6b), and AC170 (Cas2) co-occur in overlapping sets of TXI\_noncan\_h loci, consistent with components of complete CRISPR-Cas systems. Their association suggests functional coupling between TXI\_noncan\_h retrons and adaptive immunity systems.

TXI_noncan_h loci (n = 29 curated loci; all low-confidence; 22 motif-compatible; n = 24 included in phylogenetic analysis) were dominated by DnaB_C-like helicase-domain proteins encoded as separate ORFs adjacent to the RT, which were detected in 20 loci (69.0%). These proteins showed a strong positional bias toward the immediate downstream gene (offset +1; 14 loci), although upstream occurrences were also observed (offset −1; 5 loci; offset +2; 1 locus), suggesting a conserved functional coupling between RT and the associated helicase-related protein. A subset of loci (5 of 29; 17.2%) was located in proximity to CRISPR-Cas genes without a consistent positional pattern, consistent with their presence within broader defense islands rather than in a conserved operon arrangement. No WYL-domain proteins were detected in TXI_noncan_h, clearly distinguishing this lineage from TXI_C2like (**Figure 4A** and **4C**). Statistical enrichment analysis identified AC007 (DnaB_C; n = 16 loci; OR = 14.7; FDR = 1.6 × 10⁻⁹) as the dominant accessory cluster in the study population. Three additional clusters corresponding to CRISPR-Cas components—AC103 (Cas1; n = 4), AC167 (Cas6b; n = 3), and AC170 (Cas2; n = 3)—were detected exclusively in TXI_noncan_h loci (OR = ∞; FDR < 2 × 10⁻³). The co-occurrence of Cas1, Cas2, and Cas6b in overlapping loci suggests the presence of complete CRISPR-Cas systems adjacent to the retron rather than the isolated recruitment of individual components (**Table 2**).

Together, TXI_C2like and TXI_noncan_h are defined by two distinct patterns of accessory module recruitment around protease-less type XI-like RTs: a WYL-enriched lineage with a diverse and flexible accessory repertoire (TXI_C2like) and a DnaB_C-dominated lineage characterized by preferential downstream positioning and occasional association with CRISPR-Cas systems (TXI_noncan_h).

Phylogenetic analysis showed that TXI_C2like and TXI_noncan_h do not represent sporadic architectural variants scattered throughout the canonical type XI tree but instead form two distinct monophyletic radiations within a broader type XI-like clade (**Figure 5**). TXI_C2like formed a well-supported branch composed exclusively of Bacteroidota sequences, whereas TXI_noncan_h formed a separate lineage comprising both Bacillota and Bacteroidota members. A protease-fused type XI RT from *Lactobacillus paracasei* branched basally to both non-canonical lineages with strong bootstrap support (93%), consistent with a model in which these systems emerged through the loss of the ancestral protease domain, followed by the independent recruitment of distinct stand-alone accessory modules in each lineage.

**Figure 5.**
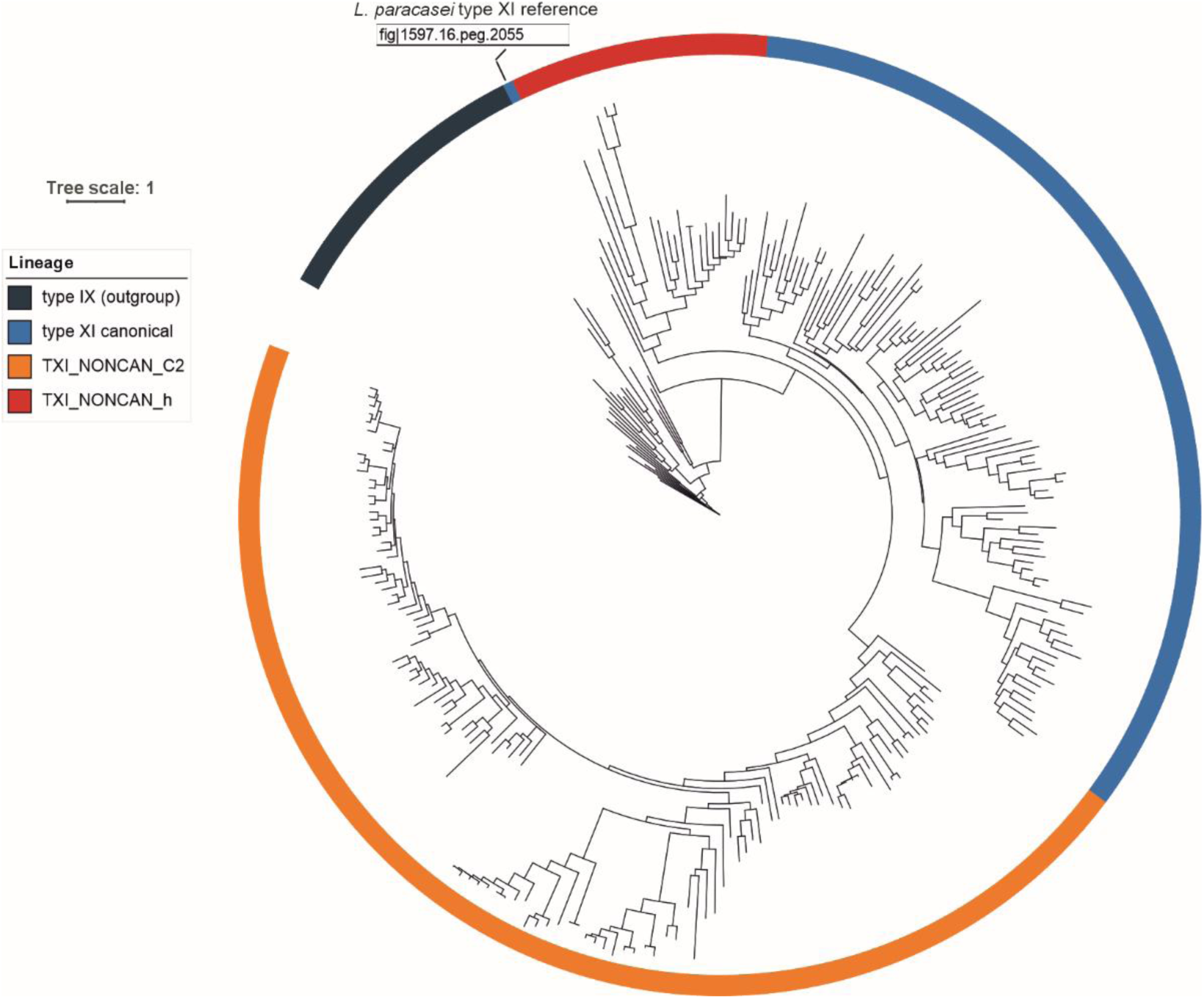
Phylogenetic structure and evolutionary diversification of noncanonical type XI-like retrons. Maximum-likelihood phylogeny of canonical and non-canonical type XI-like retron reverse transcriptases (RTs) inferred with IQ-TREE2 using the LG+I+G4 substitution model and 1,000 ultrafast bootstrap replicates from an alignment of the conserved RT domain. Type IX retron RTs (black) were included as outgroups to root the tree. TXI_C2like (n = 126; orange) and TXI_noncan_h (n = 24; red) formed two distinct monophyletic radiations within the broader type XI clade (blue), rather than representing sporadic architectural variants dispersed among canonical type XI retrons. TXI_C2like was composed exclusively of Bacteroidota sequences, whereas TXI_noncan_h comprised both Bacillota and Bacteroidota members. The position of the protease-fused type XI reference sequence from *L. paracasei* (fig1597.16_peg.2055; Mestre et al., 2020) is indicated in the tree. This sequence branches basally to the canonical and non-canonical type XI-like lineages with strong bootstrap support (93%), consistent with a model in which non-canonical type XI-like retrons arose through loss of the ancestral C-terminal protease fusion followed by independent recruitment of distinct stand-alone accessory modules.

### Identification of candidate msr/msd-like ncRNA structures in TXI_noncan_h and TXI_C2like retrons

Type XI retrons were previously recognized as one of the few retron lineages for which no msr/msd-like ncRNA structure could be identified computationally (Mestre et al., 2020). To determine whether the expanded TXI_noncan_h and TXI_C2like datasets recovered from SPIRE provided sufficient sequence diversity for ncRNA discovery, we performed de novo searches for conserved RNA secondary structures in the genomic regions upstream of RT genes.

For TXI_noncan_h, CMfinder identified a recurrent candidate motif in 20 of the 28 loci. A covariance model built from this motif detected significant matches exclusively upstream of the RT start codon at distances of 22–104 bp. Structure-guided alignment of the 300 bp immediately upstream of the RT start codon, followed by R-scape analysis, yielded a consensus structure comprising 23 base pairs, 13 of which showed statistically significant covariation (observed/expected ratio = 1.23). The predicted RNA adopts a compact double stem-loop architecture containing a conserved sequence motif consistent with the a1/a2 inverted-repeat organization, characteristic of canonical retron msr elements (**Figure 6A**).

**Figure 6.**
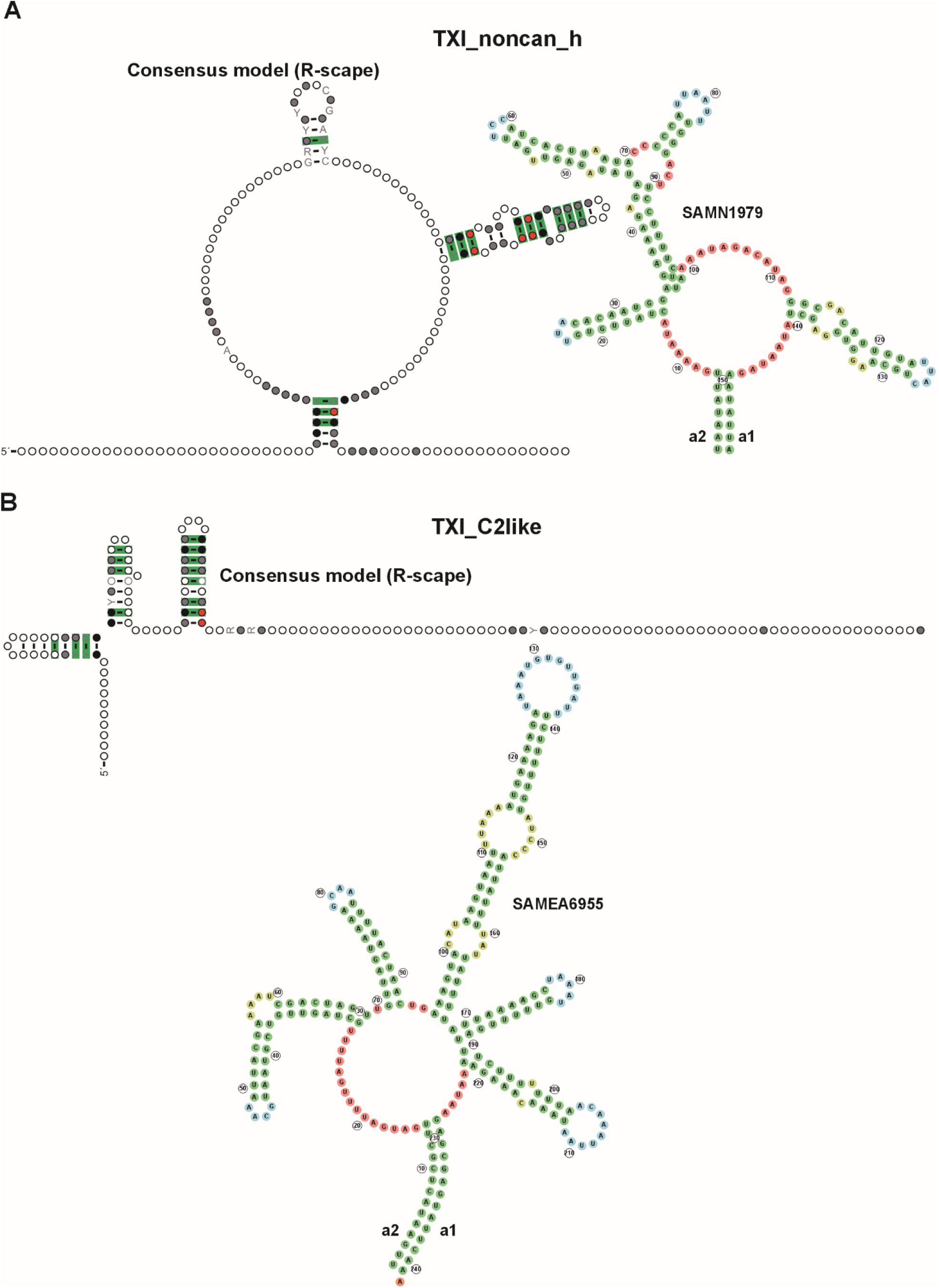
Candidate msr/msd-like ncRNA structures identified in non-canonical type XI-like retrons. (**A**) TXI_noncan_h. Consensus secondary structure of a conserved ncRNA element identified in 28 loci using the 300 bp immediately upstream of the RT start codon. The consensus model was derived from a structure-guided multiple sequence alignment generated using mLocARNA 2.0.1 and evaluated using R-scape 2.0.4. A representative RNAfold/FORNA structure from an individual locus is shown alongside the consensus model and closely recapitulates the predicted double-stem-loop architecture. (**B**) TXI_C2like. Consensus secondary structure of a conserved ncRNA element identified in 39 loci lacking an associated WYL-domain protein, using the 300 bp immediately upstream of the reverse transcriptase (RT) start codon. The consensus model was generated and evaluated as described for panel A. A representative RNAfold/FORNA structure from an individual locus is shown alongside the consensus model and recapitulates the predicted three-stem-loop architecture. For both panels, the circle size indicates positional occupancy (the proportion of sequences containing a nucleotide at each alignment position), and the circle color indicates nucleotide conservation (red, ≥97%; dark gray, ≥90%; light gray, ≥75%; white, ≥50%). Base pairs showing statistically significant covariation above the phylogenetic expectation (R-scape E < 0.05) are highlighted in green. Conserved inverted repeat elements corresponding to the canonical a1 and a2 regions are indicated, and the 5′ end of each structure is marked.

For TXI_C2like, CMfinder identified a recurrent upstream motif in 73 of the 120 loci. Covariance model searches confirmed significant matches in 37 loci, all located 38–388 bp upstream of the RT start codon. Structure-guided alignment and R-scape analyses yielded a consensus structure comprising 25 predicted base pairs, 14 of which showed statistically significant covariation (observed/expected ratio = 1.16). The resulting model displayed a three-stem-loop architecture flanked by inverted-repeat elements consistent with the a1/a2 regions characteristic of canonical retron msr/msd transcripts (**Figure 6B**). Comparable structures were recovered, irrespective of whether the associated WYL-domain protein was encoded upstream or downstream of the RT, indicating that the ncRNA is functionally linked to the RT rather than to the accessory module.

Representative RNAfold structures from individual loci closely matched the corresponding consensus models, supporting the structural conservation of both ncRNA families (**Figure 6A** and **6B**).

Together, these analyses provide the first computational evidence for candidate msr/msd-like ncRNA elements in type XI retrons, addressing a gap highlighted in previous surveys of retron ncRNA diversity (Mestre et al., 2020). Both TXI_noncan_h and TXI_C2like encode distinct conserved RNA structures upstream of their RT genes, indicating that these highly divergent lineages retain the core architectural organization of canonical retrons despite their non-canonical protein architectures.

## Discussion

This study provides the first systematic survey of retron diversity across the SPIRE prokaryotic genome resource, encompassing 107,078 representative genomes and identifying 1,286 retron candidates spanning all 13 canonical retron types, together with additional divergent lineages. In addition to confirming the broad taxonomic and ecological distribution of known retrons, our analysis uncovered two candidate novel type XI-like lineages: TXI_C2like and TXI_noncan_h. These findings demonstrate that large-scale genome-resolved surveys continue to reveal substantial unexplored diversity in retron systems.

Host-associated environments accounted for half of all retron candidates, and type XI-like retrons were strongly enriched in Bacteroidota. The pronounced host association of clade2_Ec107like further suggests that Bacillota-dominated gut communities represent a productive niche for diversification of this lineage. Quantitative analysis of taxonomic breadth using the Gini-Simpson diversity index revealed a continuum ranging from highly specialized groups, such as clade2_Ec107like and type I-C2, to broadly distributed lineages, including types VI and XIII. These contrasting patterns likely reflect distinct evolutionary strategies: specialized lineages may have co-evolved with particular bacterial hosts and their associated phage communities, whereas more cosmopolitan retrons may spread more readily through horizontal gene transfer across diverse bacterial taxa.

The subdivision of the type XI-like clade underscores the importance of resolving biologically meaningful sublineages. The apparent Bacteroidota specificity of the overall type XI-like group was driven primarily by TXI_C2like, which was detected exclusively in Bacteroidota, whereas canonical type XI retrons showed a broader taxonomic distribution, and TXI_noncan_h was recovered from both Bacteroidota and Bacillota. The ecological enrichment of type X in aquatic environments and type V in anthropogenic habitats further indicates that retron distribution is shaped not only by phylogenetic history but also by environmental context, potentially reflecting adaptation to habitat-specific phage communities. Although these patterns likely reflect genuine ecological constraints, they may also be influenced by uneven sampling across environments, emphasizing the need to explore under-represented habitats such as soils, deep marine ecosystems, and extreme environments.

The designation of TXI_C2like and TXI_noncan_h as candidate novel retron lineages is supported by four independent lines of evidence: distinct RT phylogeny, non-canonical domain architectures, lineage-specific accessory modules, and computational identification of candidate msr/msd-like ncRNA elements. Previous surveys were unable to detect ncRNA structures associated with type XI retrons (Mestre et al., 2020), highlighting the unusual divergence of this lineage and the limitations of the sequence data available at the time. By leveraging the expanded SPIRE dataset together with structure-guided alignment using mLocARNA and covariance analysis with R-scape, we identified conserved RNA structures with statistically supported covariation in both lineages. The three stem-loop architecture of TXI_C2like and the compact a1/a2-flanked structure of TXI_noncan_h are consistent with the general organizational principles of canonical retron ncRNAs, indicating that these highly divergent systems likely retain the same core functional framework.

The accessory modules associated with these lineages suggest distinct biological roles. TXI_C2like is recurrently associated with WYL-domain proteins, a class of regulatory factors implicated in nucleic acid sensing and frequently linked to bacterial defense systems (Keller and Weber-Ban, 2023). In contrast, TXI_noncan_h is associated with DnaB_C-containing proteins and, in a subset of loci, with nearby CRISPR-Cas genes, raising the possibility of functional coupling between retrons and other bacterial immune systems, consistent with the modular organization of defense islands in prokaryotic genomes (Makarova et al., 2020; Payne et al., 2021). The significance of the DnaB_C association remains unclear, but it may indicate a replication-related function or structural role within the retron ribonucleoprotein complex.

Several limitations should be considered. The conservative HMM threshold used for candidate identification prioritized specificity at the expense of sensitivity and likely underestimated the abundance of highly divergent retrons, particularly within types VI, I-B1, I-B2, and IV retrons. Searches using relaxed thresholds recovered additional candidates, but their classification was less reliable because of reduced HMM specificity in this score range. The ncRNA structures described here remain computational predictions and require further experimental validation. In addition, the predominance of metagenome-assembled genomes in SPIRE introduces potential assembly fragmentation biases that may affect genomic context analysis.

Overall, the discovery of TXI_C2like and TXI_noncan_h expands the known diversity of retron systems and highlights type XI-like retrons as a particularly dynamic evolutionary group. Experimental characterization of the associated WYL-RT and DnaB_C-RT modules, together with biochemical validation of the predicted ncRNA structures, will be essential to determine whether these systems synthesize msDNA and elucidate their biological functions. Their strong association with gut-associated Bacteroidota and Bacillota suggests that suitable cultured representatives may be available for future studies.

## Acknowledgments

The author thanks the members of the Microbiome Engineering, Reverse-transcriptases, and regulatory RNA (MIRTA) laboratory for helpful discussions.

## Author contributions

N.T. conceived the study, performed all analyses, and wrote the manuscript.

## Competing interests

The authors declare no competing financial interests.

## Funding

This work was supported by grant PID2023-147707NB-I00 from the MCIN/AEI (10.13039/501100011033).

## Notes

### Competing Interest Statement

The authors have declared no competing interest.

## References

Barbera, P., Kozlov, A.M., Czech, L., Morel, B., Darriba, D., Flouri, T. and Stamatakis, A. (2019) EPA-ng: Massively Parallel Evolutionary Placement of Genetic Sequences. Systematic Biology, 68, 365–369. 10.1093/sysbio/syy054

Benjamini, Y. and Hochberg, Y. (1995) Controlling the false discovery rate: a practical and powerful approach to multiple testing. Journal of the Royal Statistical Society Series B, 57, 289–300. 10.1111/j.2517-6161.1995.tb02031.x

Bernheim, A. and Sorek, R. (2020) The pan-immune system of bacteria: antiviral defence as a community resource. Nature Reviews Microbiology, 18, 113–119. 10.1038/s41579-019-0278-2

Bobonis, J., Mitosch, K., Mateus, A., Kritikos, G., Elfenbein, J.R., Savitski, M.M. and Typas, A. (2022) Bacterial retrons encode phage-defending tripartite toxin–antitoxin systems. Nature, 609, 144–150. 10.1038/s41586-022-05091-4

Czech, L., Barbera, P. and Stamatakis, A. (2020) Genesis and Gappa: processing, analyzing and visualizing phylogenetic (placement) data. Bioinformatics, 36, 3263–3265. 10.1093/bioinformatics/btaa070

Eddy, S.R. (2011) Accelerated Profile HMM Searches. PLoS Computational Biology, 7, e1002195. 10.1371/journal.pcbi.1002195

Gao, L., Altae-Tran, H., Böhning, F., Makarova, K.S., Segel, M., Schmid-Burgk, J.L., Koob, J., Wolf, Y.I., Koonin, E.V. and Zhang, F. (2020) Diverse enzymatic activities mediate antiviral immunity in prokaryotes. Science, 369, 1077–1084. 10.1126/science.aba0372

García-Rodríguez, F.M., Martínez-Abarca, F., Gozález-Delgado, A., Wen, D.J., Shipman. S.L., Wilkinson M.E. and Toro, N. (2025) Structurally related phage helicases trigger type III-A3 retron anti-phage defense. Nucleic Acids Research, 53, gkaf1396. 10.1093/nar/gkaf1396

Hyatt, D., Chen, G.-L., LoCascio, P.F., Land, M.L., Larimer, F.W. and Hauser, L.J. (2010) Prodigal: prokaryotic gene recognition and translation initiation site identification. BMC Bioinformatics, 11, 119. 10.1186/1471-2105-11-119

Inouye, M. and Inouye, S. (1991) msDNA and bacterial reverse transcriptase. Annual Review of Microbiology, 45, 163–186. 10.1146/annurev.mi.45.100191.001115

Katoh, K. and Frith, M.C. (2012) Adding unaligned sequences into an existing alignment using MAFFT and LAST. Bioinformatics, 28, 3144–3146. 10.1093/bioinformatics/bts578

Katoh, K. and Standley, D.M. (2013) MAFFT Multiple Sequence Alignment Software Version 7: improvements in performance and usability. Molecular Biology and Evolution, 30, 772–780. 10.1093/molbev/mst010

Keller, L.M.L. and Weber-Ban, E. (2023) An emerging class of nucleic acid-sensing regulators in bacteria: WYL domain-containing proteins. Current Opinion in Microbiology, 74, 102296. 10.1016/j.mib.2023.102296

Lampson, B.C., Inouye, M. and Inouye, S. (1989) Reverse transcriptase in a clinical strain of Escherichia coli: production of branched RNA-linked msDNA. Science, 243, 1033–1038. 10.1126/science.2466332

Letunic, I. and Bork, P. (2024) Interactive Tree of Life (iTOL) v6: recent updates to the phylogenetic tree display and annotation tool. Nucleic Acids Research, 52, W78–W82. 10.1093/nar/gkae268

Makarova, K.S., Anantharaman, V., Grishin, N.V., Koonin, E.V. and Aravind, L. (2014) CARF and WYL domains: ligand-binding regulators of prokaryotic defense systems. Frontiers in Genetics, 5, 102. 10.3389/fgene.2014.00102

Makarova, K.S., Wolf, Y.I., Snir, S. and Koonin, E.V. (2011) Defense islands in bacterial and archaeal genomes and prediction of novel defense systems. Journal of Bacteriology, 193, 6039–6056. 10.1128/jb.05535-11

Mestre, M.R., González-Delgado, A., Gutiérrez-Rus, L.I., Martínez-Abarca, F. and Toro, N. (2020) Systematic prediction of genes functionally associated with bacterial retrons and classification of the encoded tripartite systems. Nucleic Acids Research, 48, 12632–12647. 10.1093/nar/gkaa1149

Millman, A., Bernheim, A., Stokar-Avihail, A., Fedorenko, T., Voichek, M., Leavitt, A., Oppenheimer-Shaanan, Y. and Sorek, R. (2020) Bacterial retrons function in anti-phage defense. Cell, 183, 1551–1561.e12. 10.1016/j.cell.2020.09.065

Minh, B.Q., Schmidt, H.A., Chernomor, O., Schrempf, D., Woodhams, M.D., von Haeseler, A. and Lanfear, R. (2020) IQ-TREE 2: new models and efficient methods for phylogenetic inference in the genomic era. Molecular Biology and Evolution, 37, 1530–1534. 10.1093/molbev/msaa015

Mistry, J., Chuguransky, S., Williams, L., Qureshi, M., Salazar, G.A., Sonnhammer, E.L.L., Tosatto, S.C.E., Paladin, L., Raj, S., Richardson, L.J. et al. (2021) Pfam: The protein families database in 2021. Nucleic Acids Research, 49, D412–D419. 10.1093/nar/gkaa913

Müller, A.U., Leibundgut, M., Ban, N. and Weber-Ban, E. (2019) Structure and functional implications of WYL domain-containing bacterial DNA damage response regulator PafBC. Nature Communications, 10, 4653. 10.1038/s41467-019-12567-x

Nawrocki, E.P. and Eddy, S.R. (2013) Infernal 1.1: 100-fold faster RNA homology searches. Bioinformatics, 29, 2933–2935. 10.1093/bioinformatics/btt509

Payne L.J., Meaden S., Mestre M.R., Palmer C., Toro N., Fineran P.C. and Jackson S.A. (2022) PADLOC: a web server for the identification of antiviral defence systems in microbial genomes. Nucleic Acids Research 50, W541–W550. 10.1093/nar/gkac400

Rivas, E., Clements, J. and Eddy, S.R. (2017) A statistical test for conserved RNA structure shows lack of evidence for structure in lncRNAs. Nature Methods, 14, 45–48. 10.1038/nmeth.4066

Schmidt, T.S.B., Fullam, A., Ferretti, P., Orakov, A., Maistrenko, O.M., Ruscheweyh, H.-J., Letunic, I., Duan, Y., Van Rossum, T., Sunagawa, S. et al. (2024) SPIRE: a Searchable, Planetary-scale mIcrobiome REsource. Nucleic Acids Research, 52, D777–D783. 10.1093/nar/gkad943

Schubert, M.G., Goodman, D.B., Wannier, T.M., Kaur, D., Farzadfard, F., Lu, T.K., Shipman, S.L. and Church, G.M. (2021) High-throughput functional variant screens via in vivo production of single-stranded DNA. Proceedings of the National Academy of Sciences USA, 118, e2018181118. 10.1073/pnas.2018181118

Shen, W. and Ren, H. (2021) TaxonKit: a practical and efficient NCBI taxonomy toolkit. Journal of Genetics and Genomics, 48, 844–850. 10.1016/j.jgg.2021.03.006

Steinegger, M. and Söding, J. (2017) MMseqs2 enables sensitive protein sequence searching for the analysis of massive data sets. Nature Biotechnology, 35, 1026–1028. 10.1038/nbt.3988

Virtanen, P., Gommers, R., Oliphant, T.E., Haberland, M., Reddy, T., Cournapeau, D., Burovski, E., Peterson, P., Weckesser, W., Bright, J. et al. (2020) SciPy 1.0: fundamental algorithms for scientific computing in Python. Nature Methods, 17, 261–272. 10.1038/s41592-020-0772-5

Will, S., Reiche, K., Hofacker, I.L., Stadler, P.F. and Backofen, R. (2007) Inferring noncoding RNA families and classes by means of genome-scale structure-based clustering. PLoS Computational Biology, 3, e65. 10.1371/journal.pcbi.0030065

Yao, Z., Weinberg, Z. and Ruzzo, W.L. (2006) CMfinder—a covariance model based RNA motif finding algorithm. Bioinformatics, 22, 445–452. 10.1093/bioinformatics/btk008

Zhao, B., Chen, S.A., Lee, J. and Fraser, H.B. (2022) Bacterial retrons enable precise gene editing in human cells. The CRISPR Journal, 5, 31–39. 10.1089/crispr.2021.0065

