## Supplementary Figures for "Landscape of retron diversity across the SPIRE prokaryotic metagenome resource reveals candidate novel type XI-like lineages"

### Supplementary Materials

#### Supplementary Figures

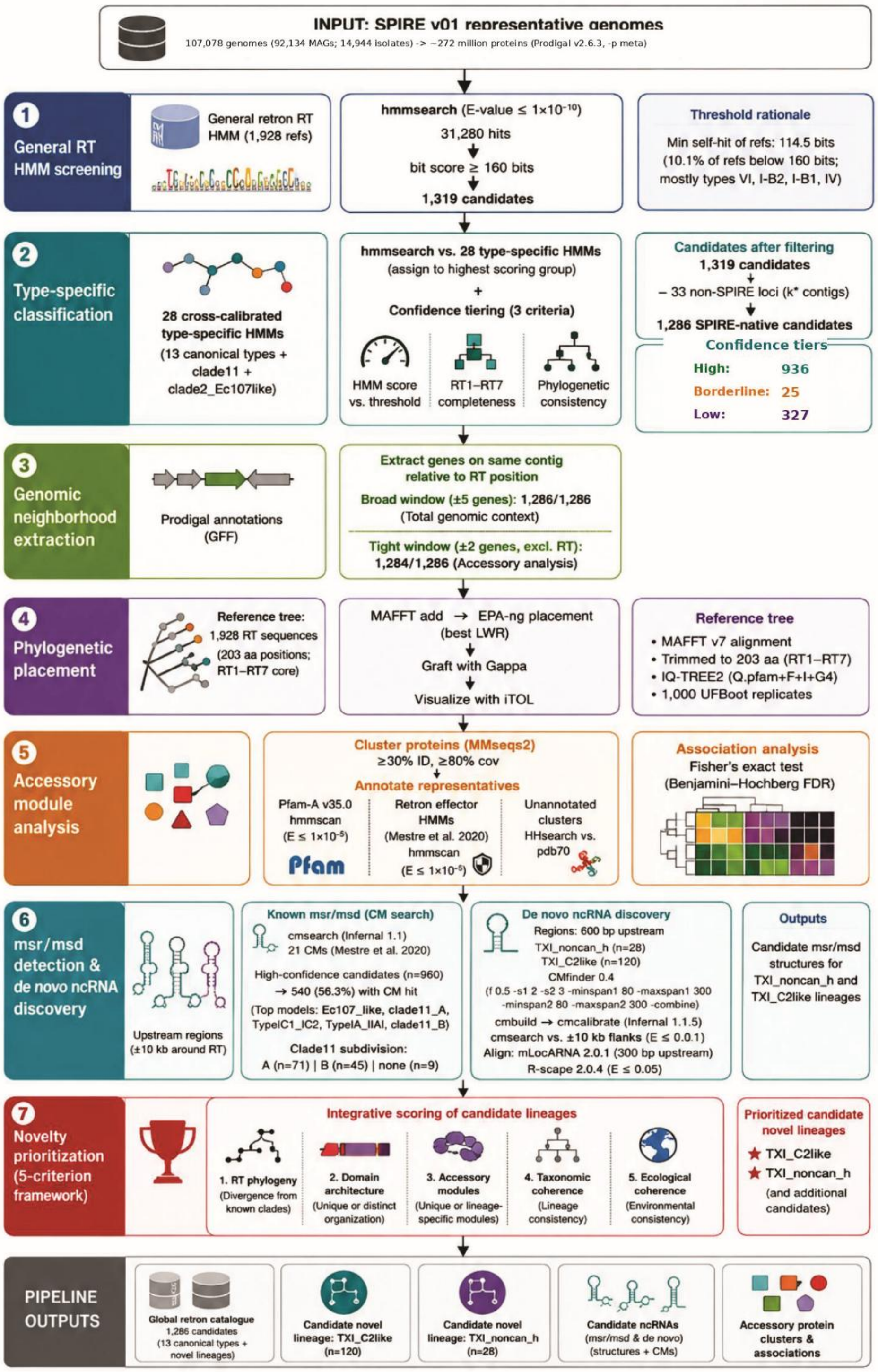

**Supplementary Figure S1. Overview of the computational pipeline used for retron discovery and classification in the SPIRE representative genome collection.** Protein-coding genes predicted from 107,078 representative genomes were screened using a general retron RT HMM, followed by classification with 28 cross-calibrated type-specific HMMs. Candidates were further evaluated through genomic neighbourhood analysis, phylogenetic placement, accessory module annotation, covariance model searches for known msr/msd elements, and de novo ncRNA structure discovery. A five-criterion novelty prioritization framework integrating RT phylogeny, domain architecture, accessory modules, taxonomic coherence, and ecological coherence was used to identify candidate novel retron lineages.

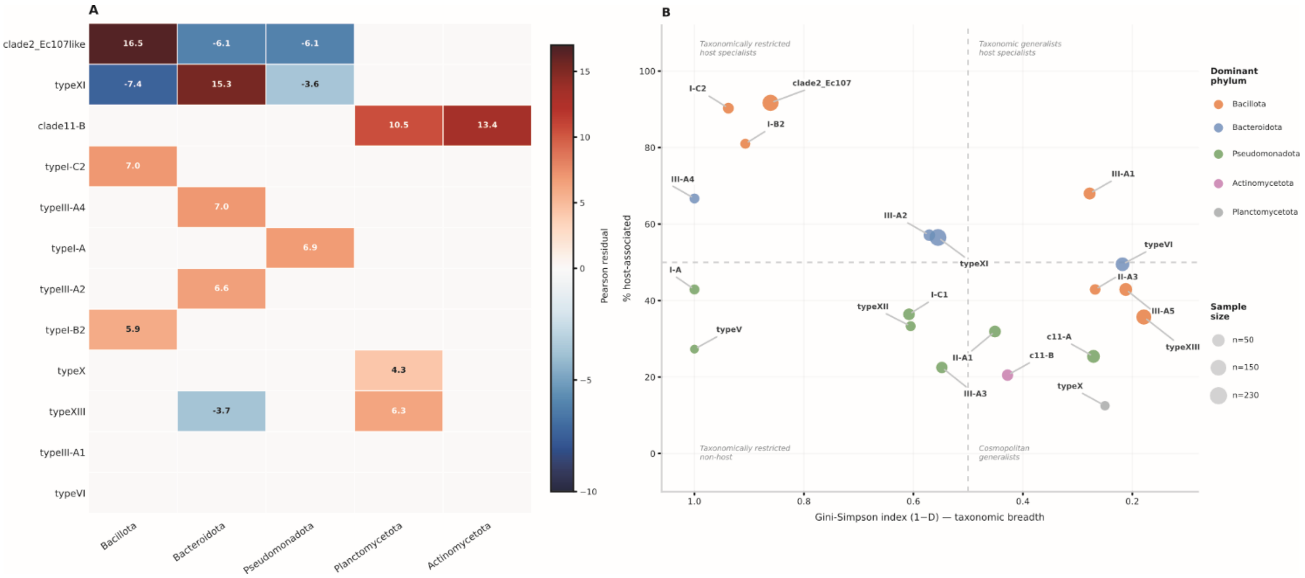


**Supplementary Figure S2. Statistical associations between retron types, phyla, and ecological specialization.** (**A**) Heatmap of Pearson standardized residuals from Chi² contingency analysis of retron group × dominant phylum ($Chi^2$=2,764.8, df=375, $p<10^{-300}$). Red cells indicate significant enrichment, blue cells significant depletion relative to the expected frequency under independence. Only groups with n≥10 and phyla with n≥5 are shown. (**B**) Scatter plot of taxonomic breadth (Gini–Simpson diversity index, 1 − D; x-axis) versus ecological specialization (% host-associated, y-axis)

for each retron group. Bubble size is proportional to the total number of candidates, and colors indicate the dominant phylum. Groups in the lower-left quadrant (low $1 - D$, high host%) are taxonomically restricted and ecologically specialized; groups in the lower-right quadrant (high $1 - D$, low host%) are broadly distributed ecological generalists.

#### Supplementary Tables

**Table S1.** Complete list of 1,286 SPIRE-native retron RT candidates identified in this study. Candidates are sorted by retron group and HMM score

**Table S2**. Complete list of 1,286 SPIRE-native retron RT candidates

**Table S3.** Systematic evidence review of all 28 SPIRE retron classification groups
