## Supplementary Table S1 for "Landscape of retron diversity across the SPIRE prokaryotic metagenome resource reveals candidate novel type XI-like lineages"

**Supplementary Table S1. HMM calibration thresholds for the 29 type-specific retron RT profiles used in SPIRE candidate classification.**

*For each HMM, pos_min = minimum self-hit score (HMM vs its own reference sequences); neg_max = maximum cross-hit score (HMM vs all other groups' reference sequences); threshold = (pos_min + neg_max) / 2. Overlap zone: neg_max > pos_min (classification inherently ambiguous). Clean separation: neg_max < pos_min (groups well-separated). High-confidence: score ≥ threshold. Low-confidence: score ≥ 160 but < threshold. Borderline: score within the overlap zone between distributions.*

| Retron type | pos_min (bits) | neg_max (bits) | threshold (bits) | Separation | Notes | n_ref_seqs_below_160 |
|---|---|---|---|---|---|---|
| **clade11** | 250,2 | 345,7 | 297,9 | **Overlap** | Overlap zone; borderline assignments expected | — |
| **clade11-A** | 290,3 | 344,6 | 317,4 | **Overlap** | Overlap zone; OutgroupA ncRNA sub-clade | — |
| **clade11-B** | 276,7 | 326,2 | 301,4 | **Overlap** | Overlap zone; OutgroupB ncRNA sub-clade | — |
| **clade2_Ec107like** | 230,5 | 258,9 | 244,7 | **Overlap** | Overlap zone; new HMM in pipeline v2 | — |
| **typeI-A** | 276,7 | 301,3 | 289,0 | **Overlap** | Overlap zone | — |
| **typeI-B1** | 236,6 | 144,4 | 190,5 | **Clean** | Clean separation; 34 refs below screening threshold | **34** |
| **typeI-B2** | 216,2 | 138,2 | 177,2 | **Clean** | Clean separation; 36 refs below screening threshold | **36** |
| **typeI-C1** | 249,7 | 311,2 | 280,4 | **Overlap** | Overlap zone | — |
| **typeI-C2** | 232,8 | 295,3 | 264,1 | **Overlap** | Overlap zone | — |
| **typeI-C3** | 310,3 | 285,8 | 298,1 | **Clean** | Clean separation | — |
| **typeII-A1** | 192,8 | 277,2 | 235,0 | **Overlap** | Overlap zone; 2 refs below screening threshold | **2** |
| **typeII-A2** | 292,7 | 216,7 | 254,7 | **Clean** | Clean separation | — |
| **typeII-A3** | 245,0 | 267,9 | 256,4 | **Overlap** | Overlap zone | — |
| **typeIII-A1** | 210,6 | 256,5 | 233,6 | **Overlap** | Overlap zone | — |
| **typeIII-A2** | 239,7 | 252,8 | 246,2 | **Overlap** | Overlap zone | — |
| **typeIII-A3** | 200,9 | 230,5 | 215,7 | **Overlap** | Overlap zone | — |
| **typeIII-A4** | 274,3 | 250,5 | 262,4 | **Clean** | Clean separation | — |
| **typeIII-A5** | 235,3 | 264,9 | 250,1 | **Overlap** | Overlap zone | — |
| **typeIV** | 263,0 | 200,5 | 231,8 | **Clean** | Clean separation; 20 refs below screening threshold | **20** |
| **typeIX** | 253,0 | 187,8 | 220,4 | **Clean** | Clean separation | — |
| **typeV** | 279,8 | 218,9 | 249,3 | **Clean** | Clean separation; 10 refs below screening threshold | **10** |
| **typeVI** | 162,9 | 207,8 | 185,3 | **Overlap** | Overlap zone; 91 refs below screening threshold — most affected | **91** |
| **typeVII-A1** | 303,6 | 177,9 | 240,8 | **Clean** | Clean separation | — |
| **typeVII-A2** | 292,7 | 172,3 | 232,5 | **Clean** | Clean separation | — |
| **typeVIII** | 360,6 | 157,0 | 258,8 | **Clean** | Clean separation; 2 refs below screening threshold | **2** |
| **typeX** | 306,2 | 211,2 | 258,7 | **Clean** | Clean separation | — |
| **typeXI** | 233,3 | 294,2 | 263,8 | **Overlap** | Overlap zone; large low-confidence fraction in SPIRE | — |
| **typeXII** | 263,5 | 302,4 | 282,9 | **Overlap** | Overlap zone | — |
| **typeXIII** | 226,3 | 255,0 | 240,7 | **Overlap** | Overlap zone | — |
| **TOTALS / RANGE (n=29 HMMs)** | | | | **Overlap: 16 \| Clean: 13** | **Threshold range: 177.2 – 317.5 bits** | **Total: 195 refs** |

**Legend:**
■ *Yellow (Overlap): neg_max > pos_min — classification ambiguous; higher borderline/low fraction expected.* ■ *Green (Clean): neg_max < pos_min — groups well-separated; high-confidence classification reliable.* ■ *Red values in column G: number of reference sequences with self-hit score < 160 bits (below the general screening threshold; these would not be recovered in SPIRE).*
