## Supplementary Table S2 for "Landscape of retron diversity across the SPIRE prokaryotic metagenome resource reveals candidate novel type XI-like lineages"

**Supplementary Table S2. Complete list of 1,286 SPIRE-native retron RT candidates identified in this study. Candidates are sorted by retron group and HMM score.  Confidence class: high = score ≥ group-specific calibrated threshold; borderline = score within the overlap zone; low = score ≥ 160 bits but below threshold.**

| Protein ID | Retron group | HMM score | Confidence class | Phylum | Class | Order | Family | Genus | Species | Sample ID | Biome | N neighbors (±5 genes) | Has tight neighbor (±2 genes) | N accessory clusters | Top Pfam annotation (neighbor) | Top effector annotation (neighbor) |
|---|---|---|---|---|---|---|---|---|---|---|---|---|---|---|---|---|
| 1736354.SAMN04151744.LMQA01000( | clade11 | 340,6 | high | Actinomycetota | Actinomycetes | Geodermatophilales | Geodermatophilaceae | Geodermatophilus | Geodermatophilus sp. Leaf369 | SAMN04151744 | host-associated | 11 | Yes | 4 | ABC_tran;Xan_ur_permease;DUF1 | — |
| 1443900.SAMN02569999.JMER01000( | clade11 | 328,1 | high | Actinomycetota | Actinomycetes | Mycobacteriales | Nocardiaceae | Rhodococcoides | Rhodococcoides fascians | SAMN02569999 | aquatic | 11 | Yes | 4 | DeoRC;YrhK;DUF3558;LEDGF | — |
| 2020130.SAMN07346923.CP022462_2 | clade11 | 322,5 | high | Actinomycetota | Actinomycetes | Micrococcales | Micrococcaceae | Arthrobacter | Arthrobacter sp. 7749 | SAMN07346923 | host-associated | 11 | Yes | 4 | HD_assoc;Dus;Bac_luciferase;Acet | — |
| 2026779.SAMN13893644.JAAYAY0100 | clade11 | 299,3 | high | Planctomycetota | Planctomycetia | Planctomycetales | Planctomycetaceae | — | Planctomycetaceae bacterium | SAMN13893644 | temperature | 8 | Yes | 4 | FAD_oxidored;ThuA;PD40;HZS_alp | — |
| 1191521.SAMEA6955893.CAIRTJ0100( | clade11 | 294,4 | high | Verrucomicrobiota | Pedosphaerae | Pedosphaerales | Pedosphaeraceae | Pedosphaera | uncultured Pedosphaera sp. | SAMEA6955893 | aquatic | 11 | Yes | 4 | Beta_helix;UnbV_ASPIC;DabA;Prot | — |
| 904984.SAMEA6956427.CAIMFH0100( | clade11 | 288,8 | high | Planctomycetota | Planctomycetia | Planctomycetales | Planctomycetaceae | Schlesneria | uncultured Schlesneria sp. | SAMEA6956427 | aquatic | 11 | Yes | 4 | Dus;PMT_2;Response_reg;FHA | — |
| 406425.SAMN02598404.CP000960_57 | clade11 | 280,6 | high | Pseudomonadota | Betaproteobacteria | Burkholderiales | Burkholderiaceae | Burkholderia | Burkholderia orbicola | SAMN02598404 | host-associated | 11 | Yes | 4 | MFS_1;Histone_HNS;PFIN;UTRA | — |
| 1062.SAMN16425547.JADJAB0100000 | clade11 | 279,1 | high | Pseudomonadota | Alphaproteobacteria | Rhodobacterales | — | Rhodobacter | Rhodobacter sp. | SAMN16425547 | anthropogenic | 11 | Yes | 4 | Peptidase_S13;CTP_transf_like;AB | — |
| 153809.SAMEA6947388.CAISGC0100( | clade11 | 264,6 | high | Pseudomonadota | — | — | — | — | uncultured Pseudomonadota bacterium | SAMEA6947388 | aquatic | 11 | Yes | 4 | FGE-sulfatase;NYN_YacP;ACR_tra | — |
| 479431.SAMN02598467.CP001737_31 | clade11-A | 349,8 | high | Actinomycetota | Actinomycetes | Nakamurellales | Nakamurellaceae | Nakamurella | Nakamurella multipartita | SAMN02598467 | anthropogenic | 11 | Yes | 4 | CBM_48;Zn_peptidase;LysE | — |
| 1736303.SAMN04151693.LMLU010000( | clade11-A | 341 | high | Actinomycetota | Actinomycetes | Micrococcales | Micrococcaceae | Arthrobacter | Arthrobacter sp. Leaf234 | SAMN04151693 | anthropogenic | 11 | Yes | 4 | BPD_transp_1;DUF5996;ABC_tran | — |
| 1032480.SAMD00061117.AP012204_2 | clade11-A | 337 | high | Actinomycetota | Actinomycetes | Propionibacteriales | Propionibacteriaceae | Microlunatus | Microlunatus phosphovorus | SAMD00061117 | anthropogenic | 11 | Yes | 4 | DUF2252;tRNA-synt_His;ROK;CSI | — |
| 2769423.SAMD00245656.BNCN010000( | clade11-A | 333,1 | high | Pseudomonadota | Betaproteobacteria | Burkholderiales | Comamonadaceae | Comamonas | Comamonas sp. KCTC 72670 | SAMD00245656 | host-associated | 11 | Yes | 4 | HSP70;Phage_holin_3_6;DUF6596 | — |
| 2026658.SAMN14167782.JAAKZI0100( | clade11-A | 332,3 | high | Actinomycetota | Actinomycetes | Micrococcales | Micrococcaceae | Arthrobacter | Arthrobacter silviterrae | SAMN14167782 | host-associated | 11 | Yes | 4 | DSBA;Guanylate_cyc | — |
| 1898745.SAMN05736493.MJGS010000( | clade11-A | 331,7 | high | Actinomycetota | Actinomycetes | Micrococcales | Microbacteriaceae | Frigoribacterium | Frigoribacterium sp. MCBA15_019 | SAMN05736493 | anthropogenic | 11 | Yes | 4 | FlgD;Flg_bbr_C;DUF4190;YitT_mer | — |
| 1522176.SAMN12025139.JAAOYV0100( | clade11-A | 329,3 | high | Actinomycetota | Actinomycetes | Micrococcales | Microbacteriaceae | Frigoribacterium | Frigoribacterium endophyticum | SAMN12025139 | host-associated | 11 | Yes | 4 | YitT_membrane;Mannosyl_trans2;F | — |
| 2175619.SAMN09009058.QKSS010000( | clade11-A | 329,1 | high | Actinomycetota | Actinomycetes | Micrococcales | Microbacteriaceae | Curtobacterium | Curtobacterium sp. MCJR17_020 | SAMN09009058 | anthropogenic | 11 | Yes | 4 | Bac_luciferase;Acyl-CoA_dh_M;Gly | — |
| 1279028.SAMN05660766.OCMI010000 | clade11-A | 329 | high | Actinomycetota | Actinomycetes | Micrococcales | Microbacteriaceae | Curtobacterium | Curtobacterium sp. 314Chir4.1 | SAMN05660766 | host-associated | 11 | Yes | 4 | Bac_luciferase;Acyl-CoA_dh_M;Mg | — |
| 2017486.SAMN07299025.CP022296_4 | clade11-A | 328,7 | high | Actinomycetota | Actinomycetes | Propionibacteriales | Nocardioidaceae | Nocardioides | Nocardioides sp. S5 | SAMN07299025 | aquatic | 11 | Yes | 4 | Acyltransferase;MerR-DNA-bind;Dn | — |
| 1905847.SAMN05854879.CP017580_3 | clade11-A | 328,3 | high | Actinomycetota | Actinomycetes | Micrococcales | Microbacteriaceae | Curtobacterium | Curtobacterium sp. BH-2-1-1 | SAMN05854879 | — | 11 | Yes | 0 | — | — |
| 2485192.SAMN10361296.RKIE0100000( | clade11-A | 328,1 | high | Actinomycetota | Actinomycetes | Micrococcales | Microbacteriaceae | Frigoribacterium | Frigoribacterium sp. PhB160 | SAMN10361296 | host-associated | 7 | Yes | 3 | Apolipoprotein;YitT_membrane;FlbD | — |
| 1909295.SAMN08179254.PLES010001 | clade11-A | 327,1 | high | Actinomycetota | Thermoleophilia | Solirubrobacterales | — | — | Solirubrobacterales bacterium | SAMN08179254 | temperature | 11 | Yes | 4 | Cyt_bd_oxida_II;AHSA1;HTH_20 | — |
| 351215.SAMN14517837.JABEMA0100( | clade11-A | 326,3 | high | Actinomycetota | Actinomycetes | Kineosporiales | Kineosporiaceae | Pseudokineococcus | Pseudokineococcus marinus | SAMN14517837 | anthropogenic | 11 | Yes | 4 | Glyoxalase;Glyoxalase_6;MST;MDN | — |
| 2487580.SAMN16425975.JADJPO0100 | clade11-A | 323,3 | high | Actinomycetota | Actinomycetes | Kineosporiales | Kineosporiaceae | — | Kineosporiaceae bacterium | SAMN16425975 | anthropogenic | 11 | Yes | 4 | Cation_efflux;ROK;AP_endonuc_2 | — |
| 1735682.SAMN04151684.LMKK010000( | clade11-A | 320,6 | high | Actinomycetota | Actinomycetes | Micrococcales | Microbacteriaceae | Agreia | Agreia sp. Leaf210 | SAMN04151684 | host-associated | 11 | Yes | 4 | TMEM260-like;AAA_2 | — |
| 477641.SAMEA2272685.FO203431_10 | clade11-A | 309,9 | high | Actinomycetota | Actinomycetes | Geodermatophilales | Geodermatophilaceae | Modestobacter | Modestobacter marinus | SAMEA2272685 | anthropogenic | 11 | Yes | 4 | Glyco_trans_1_2;LOR;Glyco_hydro | — |
| 100233.SAMEA6946666.CAIVBC01000 | clade11-A | 306,1 | high | Planctomycetota | Planctomycetia | Planctomycetales | Planctomycetaceae | — | uncultured Planctomycetaceae bacteri | SAMEA6946666 | aquatic | 3 | Yes | 2 | WHD_3rd_SelB;ORF6C | — |
| 100233.SAMEA6956128.CAIXVO0100( | clade11-A | 302,3 | high | Planctomycetota | Planctomycetia | Planctomycetales | Planctomycetaceae | — | uncultured Planctomycetaceae bacteri | SAMEA6956128 | aquatic | 8 | Yes | 4 | META;Thil_fer;GBD-like_ZMIZ1_ZI | — |
| 1218077.SAMD00000366.BAYC010000 | clade11-A | 300,9 | high | Pseudomonadota | Betaproteobacteria | Burkholderiales | Burkholderiaceae | Paraburkholderia | Paraburkholderia fungorum | SAMD00000366 | aquatic | 11 | Yes | 4 | Phage_GPD;GyrI-like;AA_permease | — |
| 1884385.SAMN05518669.FNOL010000 | clade11-A | 300,3 | high | Pseudomonadota | Betaproteobacteria | Burkholderiales | Comamonadaceae | Variovorax | Variovorax sp. YR634 | SAMN05518669 | host-associated | 11 | Yes | 4 | FAD_binding_3;SMI1_KNR4;Trp_d | — |
| 2026791.SAMN08179553.PMKM01000 | clade11-A | 300,3 | high | Acidobacteriota | Terriglobia | Bryobacterales | — | — | Bryobacterales bacterium | SAMN08179553 | temperature | 11 | Yes | 4 | CxxCxxCC;ParE_like;SIS_2 | — |
| 2052164.SAMN08179277.PLDW01000( | clade11-A | 299,5 | high | Planctomycetota | Planctomycetia | Gemmatales | Gemmataceae | — | Gemmataceae bacterium | SAMN08179277 | temperature | 11 | Yes | 4 | PGM_PMM_I | — |
| 1191521.SAMEA6956431.CAIYZC0100 | clade11-A | 299,1 | high | Verrucomicrobiota | Pedosphaerae | Pedosphaerales | Pedosphaeraceae | Pedosphaera | uncultured Pedosphaera sp. | SAMEA6956431 | aquatic | 3 | Yes | 2 | Aconitase_C;Glyco_hydro_20b | — |
| 2654182.SAMN17140642.JAFAZI0100( | clade11-A | 298,1 | high | Actinomycetota | Actinomycetes | Frankiales | Frankiaceae | — | Frankiaceae bacterium | SAMN17140642 | host-associated | 11 | Yes | 4 | Response_reg;Peptidase_S11;DUF | — |
| 1913988.SAMN17181029.JAEUML010( | clade11-A | 298 | high | Pseudomonadota | Alphaproteobacteria | — | — | — | Alphaproteobacteria bacterium | SAMN17181029 | anthropogenic | 10 | Yes | 4 | SET;Peptidase_M3;DUF308;Questi | — |
| 1797564.SAMN04316052.MESG01000( | clade11-A | 297,8 | high | Pseudomonadota | Betaproteobacteria | Burkholderiales | — | — | Burkholderiales bacterium RIFCSPLO' | SAMN04316052 | terrestrial | 11 | Yes | 4 | BPD_transp_1;PhyH;Aldedh;Amino | — |
| 2026779.SAMEA104651807.OMIV0100 | clade11-A | 297 | high | Planctomycetota | Planctomycetia | Planctomycetales | Planctomycetaceae | — | Planctomycetaceae bacterium | SAMEA104651807 | aquatic | 7 | Yes | 3 | PGM_PMM_I;GFO_IDH_MocA_C | — |
| 60549.SAMN05444165.FSRU01000002 | clade11-A | 296,3 | high | Pseudomonadota | Betaproteobacteria | Burkholderiales | Burkholderiaceae | Paraburkholderia | Paraburkholderia phenazinium | SAMN05444165 | host-associated | 11 | Yes | 4 | AtuA_ferredoxin;HpsJ;Polysacc_de | — |
| 1882827.SAMN05443579.FOWG01000 | clade11-A | 295,8 | high | Pseudomonadota | Betaproteobacteria | Burkholderiales | Comamonadaceae | Variovorax | Variovorax sp. PDC80 | SAMN05443579 | anthropogenic | 11 | Yes | 4 | FAD_binding_3;SMI1_KNR4;Trp_d | — |
| 1781067.SAMN04320694.LRHV010000 | clade11-A | 295,8 | high | Pseudomonadota | Betaproteobacteria | Burkholderiales | Oxalobacteraceae | Duganella | Duganella sp. HH105 | SAMN04320694 | host-associated | 11 | Yes | 4 | DUF885;Ricin_B_lectin;DUF945;Ep | — |
| 1978231.SAMN08912219.QIAH010004! | clade11-A | 295,8 | high | Acidobacteriota | — | — | — | — | Acidobacteriota bacterium | SAMN08912219 | terrestrial | 10 | Yes | 4 | Cytochrom_C552;CMD;OMP_b-brl_ | — |
| 2026791.SAMN08178866.PMRN01000: | clade11-A | 295,4 | high | Acidobacteriota | Terriglobia | Bryobacterales | — | — | Bryobacterales bacterium | SAMN08178866 | temperature | 8 | Yes | 4 | MacB_PCD;FitA-like_RHH | — |
| 2026748.SAMN16635730.JADMJM010( | clade11-A | 294,5 | high | Pseudomonadota | Alphaproteobacteria | Hyphomonadales | Hyphomonadaceae | — | Hyphomonadaceae bacterium | SAMN16635730 | terrestrial | 11 | Yes | 4 | HD;DAP_epimerase;zf-HC2 | — |
| 2502223.SAMN08712290.SCNV010000( | clade11-A | 294,3 | high | Pseudomonadota | Betaproteobacteria | Burkholderiales | Burkholderiaceae | Burkholderia | Burkholderia sp. 4M9327F10 | SAMN08712290 | aquatic | 6 | Yes | 2 | AtuA_ferredoxin;AtuA | — |
| 2026791.SAMN08180001.PLRJ010000( | clade11-A | 294,2 | high | Acidobacteriota | Terriglobia | Bryobacterales | — | — | Bryobacterales bacterium | SAMN08180001 | temperature | 11 | Yes | 4 | DUF433;Acetyltransf_1;DUF4890;E | — |
| 2854789.SAMN19994155.JAHUWW01( | clade11-A | 291,4 | high | Pseudomonadota | Betaproteobacteria | Burkholderiales | Comamonadaceae | Acidovorax | Acidovorax sp. sif0632 | SAMN19994155 | terrestrial | 11 | Yes | 4 | Ni_hydr_CYTB;Acetyltransf_1;Redc | — |
| 1121035.SAMN02441190.AUCH01000( | clade11-A | 291 | high | Pseudomonadota | Betaproteobacteria | Rhodocyclales | Rhodocyclaceae | Azovibrio | Azovibrio restrictus | SAMN02441190 | host-associated | 11 | Yes | 4 | DXP_synthase_N;GCHY-1;Exo_en | — |
| 2052484.SAMN07200918.NIOE010000( | clade11-A | 290,7 | high | Pseudomonadota | Betaproteobacteria | Burkholderiales | Sphaerotilaceae | Roseateles | Roseateles noduli | SAMN07200918 | host-associated | 11 | Yes | 4 | Response_reg;2CSK_N;LysR_subs | — |
| 48.SAMN07426514.QFQP01000062_1( | clade11-A | 290,7 | high | Myxococcota | Myxococcia | Myxococcales | Archangiaceae | Archangium | Archangium gephyra | SAMN07426514 | anthropogenic | 6 | Yes | 2 | Pentapeptide_4 | — |
| 904984.SAMEA6955890.CAIOKO0100( | clade11-A | 289,3 | low | Planctomycetota | Planctomycetia | Planctomycetales | Planctomycetaceae | Schlesneria | uncultured Schlesneria sp. | SAMEA6955890 | aquatic | 10 | Yes | 4 | GFO_IDH_MocA;NIPSNAP;RtcB;H | — |
| 946333.SAMN04621830.CP015118_73 | clade11-A | 288,8 | low | Pseudomonadota | Betaproteobacteria | Burkholderiales | Sphaerotilaceae | Piscinibacter | Piscinibacter gummiphilus | SAMN04621830 | host-associated | 10 | Yes | 4 | TauE;Hrs_helical;DUF4194 | — |
| 1801619.SAMN04515619.FOLC010000 | clade11-A | 288,3 | low | Pseudomonadota | Betaproteobacteria | Burkholderiales | Oxalobacteraceae | Collimonas | Collimonas sp. OK412 | SAMN04515619 | host-associated | 10 | Yes | 4 | GTP_EFTU;GTP_cyclohydro2;PqiA | — |
| 171953.SAMEA6946981.CAITVS01000 | clade11-A | 288,2 | low | Acidobacteriota | — | — | — | — | uncultured Acidobacteriota bacterium | SAMEA6946981 | aquatic | 9 | Yes | 4 | OMP_b-brl_4;Ank_5;tRNA-synt_1_ | — |
| 2682146.SAMN16425753.JADJHQ010( | clade11-A | 287,8 | low | Pseudomonadota | Betaproteobacteria | Nitrosomonadales | Sterolibacteriaceae | — | Sterolibacteriaceae bacterium | SAMN16425753 | anthropogenic | 10 | Yes | 4 | DUF4372;KH_NucS_shadow;EAL | — |
| 2004485.SAMN07156299.NOIG010000 | clade11-A | 284,8 | low | Pseudomonadota | Betaproteobacteria | Burkholderiales | Comamonadaceae | Acidovorax | Acidovorax kalamii | SAMN07156299 | aquatic | 10 | Yes | 4 | Ni_hydr_CYTB;Acetyltransf_1;Redc | — |
| 2052142.SAMN08179306.PLCU010001 | clade11-A | 284,7 | low | Acidobacteriota | Terriglobia | Terriglobales | Acidobacteriaceae | — | Acidobacteriaceae bacterium | SAMN08179306 | temperature | 10 | Yes | 4 | Lactamase_B;Dus;Ribosomal_L31 | — |
| 2026791.SAMN08178945.PMUN01000: | clade11-A | 284,7 | low | Acidobacteriota | Terriglobia | Bryobacterales | — | — | Bryobacterales bacterium | SAMN08178945 | temperature | 10 | Yes | 4 | PMSR;Aminotran_3;TehB | — |
| 2587122.SAMN12024204.JACHYS010( | clade11-A | 284,2 | low | Pseudomonadota | Betaproteobacteria | Burkholderiales | Sphaerotilaceae | Roseateles | Mitsuaria sp. BK037 | SAMN12024204 | host-associated | 10 | Yes | 4 | DUF488-N3a;LysR_substrate;Abhy | — |
| 2838466.SAMN15816783.DXIO010001 | clade11-A | 284,2 | low | Pseudomonadota | Betaproteobacteria | Burkholderiales | — | Aquabacterium | Candidatus Aquabacterium excrementi | SAMN15816783 | host-associated | 10 | Yes | 4 | DUF5691;SWIM;HTH_18;Isochoris | — |
| 156588.SAMEA6944372.CAIMXY0100( | clade11-A | 282,8 | low | Verrucomicrobiota | — | — | — | — | uncultured Verrucomicrobiota bacteriu | SAMEA6944372 | aquatic | 9 | Yes | 4 | Zn_ribbon_4;Sigma70_r2;FecR | — |
| 259537.SAMN17348697.JAFIHA01000( | clade11-A | 282,3 | low | Pseudomonadota | Betaproteobacteria | Rhodocyclales | Azonexaceae | Dechloromonas | Dechloromonas aromatica (nom. nud.) | SAMN17348697 | host-associated | 6 | Yes | 3 | TerC;tRNA-synt_1b;ThermoDBP-RI | — |
| 1871071.SAMN18061362.JAFKGX010( | clade11-A | 281,5 | low | Pseudomonadota | Betaproteobacteria | Burkholderiales | Comamonadaceae | — | Comamonadaceae bacterium | SAMN18061362 | host-associated | 10 | Yes | 4 | Ribosomal_S17;Redoxin;Acetyltrans | — |
| 2026791.SAMN08178902.PMSX01000( | clade11-A | 281,5 | low | Acidobacteriota | Terriglobia | Bryobacterales | — | — | Bryobacterales bacterium | SAMN08178902 | temperature | 10 | Yes | 4 | Beta-prop_THOC3;PSD4 | — |
| 2026779.SAMN11369404.SYJV010002 | clade11-A | 281,5 | low | Planctomycetota | Planctomycetia | Planctomycetales | Planctomycetaceae | — | Planctomycetaceae bacterium | SAMN11369404 | aquatic | 6 | Yes | 3 | Sulfatase;DPBB_PEX6;SGSH_C | — |
| 2026791.SAMN08178783.PMOK01000( | clade11-A | 280,6 | low | Acidobacteriota | Terriglobia | Bryobacterales | — | — | Bryobacterales bacterium | SAMN08178783 | — | 10 | Yes | 0 | — | — |
| 1978231.SAMN14214674.JABCZI0100( | clade11-A | 277,7 | low | Acidobacteriota | — | — | — | — | Acidobacteriota bacterium | SAMN14214674 | terrestrial | 10 | Yes | 4 | Dus;AIM24;Lactamase_B;AAA_2 | — |
| 2026799.SAMN14913714.JABGQM010 | clade11-A | 277 | low | Verrucomicrobiota | — | — | — | — | Verrucomicrobiota bacterium | SAMN14913714 | aquatic | 10 | Yes | 3 | ABC2_membrane_3;SBP_bac_10;I | — |
| 2053492.SAMN18059897.JAFLDO010( | clade11-A | 272,3 | low | Pseudomonadota | Betaproteobacteria | — | — | Candidatus Accumulib | Accumulibacter sp. | SAMN18059897 | anthropogenic | 10 | Yes | 4 | PacC_helical;Response_reg;HATPa | — |
| 1913988.SAMN18118915.JAGNKE010( | clade11-A | 271,6 | low | Pseudomonadota | Alphaproteobacteria | — | — | — | Alphaproteobacteria bacterium | SAMN18118915 | anthropogenic | 10 | Yes | 4 | GtfA_EBD;Pro_CA;MASE5;UDG | — |
| 1978231.SAMN14214676.JABCZK010( | clade11-A | 271,4 | low | Acidobacteriota | — | — | — | — | Acidobacteriota bacterium | SAMN14214676 | terrestrial | 10 | Yes | 4 | HTH_8;SAICAR_synt;SMC_N | — |
| 2026791.SAMN08178828.PMQB01000! | clade11-A | 269,7 | low | Acidobacteriota | Terriglobia | Bryobacterales | — | — | Bryobacterales bacterium | SAMN08178828 | temperature | 5 | Yes | 3 | Lipase_GDSL_2;HHH_3;Beta-prop_ | — |

| | | | | | | | | | | | | | | | | |
|---|---|---|---|---|---|---|---|---|---|---|---|---|---|---|---|---|
| 86027.SAMEA6956982.CAIMXF01000C | **clade11-A** | 269,1 | low | Pseudomonadota | Betaproteobacteria | — | — | — | uncultured beta proteobacterium | SAMEA6956982 | aquatic | 10 | Yes | 3 | MFS_1;Phenol_MetA_deg | — |
| 1632864.SAMN03417854.CP011270_3 | **clade11-A** | 268,8 | low | Planctomycetota | Planctomycetia | Planctomycetales | Planctomycetaceae | Planctomyces | Planctomyces sp. SH-PL14 | SAMN03417854 | anthropogenic | 10 | Yes | 4 | CENP-F_leu_zip;SIS;Reprolysin_3; | — |
| 2072747.SAMN08366045.RCHM01000 | **clade11-A** | 268,2 | low | Pseudomonadota | Gammaproteobacteria | Pseudomonadales | Ketobacteraceae | Ketobacter | Ketobacter sp. GenoA1 | SAMN08366045 | aquatic | 10 | Yes | 4 | Acyl-CoA_dh_1;ChaC;DUF4234;Al | — |
| 373675.SAMN04488045.FNUZ0100000 | **clade11-A** | 265,1 | low | Pseudomonadota | Alphaproteobacteria | Rhodobacterales | Roseobacteraceae | Thalassococcus | Thalassococcus halodurans | SAMN04488045 | aquatic | 10 | Yes | 4 | Pyr_redox_2;Dioxygenase_C;Sacch | — |
| 2591109.SAMN12116251.CP041352_3 | **clade11-A** | 263,6 | low | Pseudomonadota | Betaproteobacteria | — | Casimicrobiaceae | Casimicrobium | Casimicrobium huifangae | SAMN12116251 | anthropogenic | 10 | Yes | 4 | HMGL-like;2-Hacid_dh_C;CPSase_ | — |
| 100234.SAMEA9694258.CAJXQE01000 | **clade11-A** | 260 | low | Verrucomicrobiota | Verrucomicrobiia | Verrucomicrobiales | — | — | uncultured Verrucomicrobiales bacteriu | SAMEA9694258 | temperature | 7 | Yes | 4 | SASA;Sulfatase;LRR_14;Beta-prop | — |
| 2595004.SAMN12273953.VKKX010000 | **clade11-A** | 253,7 | low | Pseudomonadota | Alphaproteobacteria | Rhodobacterales | Paracoccaceae | Gemmobacter | Gemmobacter caeruleus | SAMN12273953 | anthropogenic | 10 | Yes | 4 | Acetyltransf_1;GFA;ABC_tran;MipZ | — |
| 1891241.SAMN11380348.VBCQ01000 | **clade11-A** | 233,9 | low | Pseudomonadota | Betaproteobacteria | — | — | — | Betaproteobacteria bacterium | SAMN11380348 | terrestrial | 7 | Yes | 4 | GTP_EFTU;UbiA;ELFV_dehydrog | — |
| 1978231.SAMN10966252.VFZW010000 | **clade11-A** | 214,2 | low | Acidobacteriota | — | — | — | — | Acidobacteriota bacterium | SAMN10966252 | aquatic | 5 | Yes | 2 | PQQ_2;Vgb_lyase | — |
| 1314686.SAMN06349878.MVKV01000C | **clade11-B** | 336,2 | high | Actinomycetota | Actinomycetes | Mycobacteriales | Gordoniaceae | Gordonia | Gordonia sp. IITR100 | SAMN06349878 | host-associated | 9 | Yes | 4 | ABC_tran;Acetyltransf_1;Globin;GT | — |
| 1136179.SAMN02603403.CP003761_5 | **clade11-B** | 319 | high | Actinomycetota | Actinomycetes | Mycobacteriales | Nocardiaceae | Rhodococcus | Rhodococcus erythropolis | SAMN02603403 | — | 11 | Yes | 0 | — | — |
| 1112204.SAMN02603285.CP003119_4 | **clade11-B** | 319 | high | Actinomycetota | Actinomycetes | Mycobacteriales | Gordoniaceae | Gordonia | Gordonia polyisoprenivorans | SAMN02603285 | aquatic | 11 | Yes | 4 | Acyl_transf_3;CorA;HNH;Sulfotrans | — |
| 1219016.SAMD00047211.BCWX01000 | **clade11-B** | 318,8 | high | Actinomycetota | Actinomycetes | Mycobacteriales | Nocardiaceae | Rhodococcus | Rhodococcus globerulus | SAMD00047211 | anthropogenic | 11 | Yes | 4 | Aldolase_II;NMT1;TetR_N;ECH_1 | — |
| 2739471.SAMN15016254.JABUDE010C | **clade11-B** | 318,2 | high | Actinomycetota | Actinomycetes | Mycobacteriales | Nocardiaceae | Rhodococcus | Rhodococcus sp. BP-369 | SAMN15016254 | aquatic | 11 | Yes | 4 | Flavin_Reduct;VanW;Thiolase_N;a | — |
| 1051662.SAMN02569993.JMEX010000 | **clade11-B** | 315,3 | high | Actinomycetota | Actinomycetes | Mycobacteriales | Nocardiaceae | Rhodococcoides | Rhodococcoides fascians | SAMN02569993 | host-associated | 11 | Yes | 4 | Beta-lactamase;FAD-oxidase_C;adh | — |
| 1108044.SAMD00041778.BAFB010000 | **clade11-B** | 314,1 | high | Actinomycetota | Actinomycetes | Mycobacteriales | Gordoniaceae | Gordonia | Gordonia otitidis | SAMD00041778 | anthropogenic | 11 | Yes | 4 | EryCIII-like_C;DUF3046;RecA_N;R | — |
| 1219362.SAMD00046772.BDAP010000 | **clade11-B** | 314 | high | Actinomycetota | Actinomycetes | Mycobacteriales | Nocardiaceae | Williamsia | Williamsia muralis | SAMD00046772 | anthropogenic | 11 | Yes | 4 | Ala_racemase_N;DNA_pol3_alpha; | — |
| 2806442.SAMN17526040.CP069242_2 | **clade11-B** | 313,4 | high | Actinomycetota | Actinomycetes | Mycobacteriales | Nocardiaceae | Rhodococcus | Rhodococcus sp. USK13 | SAMN17526040 | host-associated | 11 | Yes | 4 | Polyketide_cyc2;PLDc_N;TetR_N;F | — |
| 498198.SAMN06265174.FXTG0100001 | **clade11-B** | 312,7 | high | Actinomycetota | Actinomycetes | Mycobacteriales | Dietziaceae | Dietzia | Dietzia kunjamensis | SAMN06265174 | anthropogenic | 6 | Yes | 2 | DUF4131;DUF4077 | — |
| 194249.SAMEA9694981.CAJXRT01000 | **clade11-B** | 311,9 | high | Actinomycetota | Actinomycetes | Mycobacteriales | Nocardiaceae | Rhodococcus | uncultured Rhodococcus sp. | SAMEA9694981 | aquatic | 11 | Yes | 4 | GyrI-like;HTH_3;AzlC | — |
| 2838538.SAMN15816639.DWWP0100C | **clade11-B** | 311,4 | high | Actinomycetota | Actinomycetes | Mycobacteriales | Dietziaceae | Dietzia | Candidatus Dietzia intestinipullorum | SAMN15816639 | host-associated | 7 | Yes | 3 | DUF2339;Guanylate_cyc | — |
| 632772.SAMD00060964.AP011115_53 | **clade11-B** | 310,7 | high | Actinomycetota | Actinomycetes | Mycobacteriales | Nocardiaceae | Rhodococcus | Rhodococcus opacus | SAMD00060964 | host-associated | 11 | Yes | 4 | Nitroreductase;adh_short_C2;DUF3 | — |
| 1089453.SAMD00041776.BAFC010000 | **clade11-B** | 309,2 | high | Actinomycetota | Actinomycetes | Mycobacteriales | Gordoniaceae | Gordonia | Gordonia sputi | SAMD00041776 | anthropogenic | 11 | Yes | 4 | EryCIII-like_C;DUF3046;RecA_N;R | — |
| 1220583.SAMD00041806.BANR010000 | **clade11-B** | 308,7 | high | Actinomycetota | Actinomycetes | Mycobacteriales | Gordoniaceae | Gordonia | Gordonia aichiensis | SAMD00041806 | anthropogenic | 11 | Yes | 4 | EryCIII-like_C;DUF3046;RecA_N;R | — |
| 2838537.SAMN15816635.DWWG0100C | **clade11-B** | 304,2 | high | Actinomycetota | Actinomycetes | Mycobacteriales | Dietziaceae | Dietzia | Candidatus Dietzia intestinigallinarum | SAMN15816635 | host-associated | 11 | Yes | 4 | EamA;FUSC_2;PadR;HAAS | — |
| 57704.SAMN04457857.LSRG01000010 | **clade11-B** | 296,5 | high | Actinomycetota | Actinomycetes | Mycobacteriales | Tsukamurellaceae | Tsukamurella | Tsukamurella tyrosinosolvens | SAMN04457857 | host-associated | 11 | Yes | 4 | Isochorismatase;HTH_18;HAD_2;H | — |
| 47312.SAMN04457854.LSRH01000040 | **clade11-B** | 291,9 | high | Actinomycetota | Actinomycetes | Mycobacteriales | Tsukamurellaceae | Tsukamurella | Tsukamurella pulmonis | SAMN04457854 | host-associated | 11 | Yes | 4 | HAD_2;DUF2157;Acetyltransf_1 | — |
| 100233.SAMEA6952416.CAIOTJ01000 | **clade11-B** | 276,6 | low | Planctomycetota | Planctomycetia | Planctomycetales | Planctomycetaceae | — | uncultured Planctomycetaceae bacteri | SAMEA6952416 | aquatic | 10 | Yes | 4 | Response_reg;Sacchrp_dh_NADP; | — |
| 100234.SAMEA9695074.CAJXVQ0100 | **clade11-B** | 273,5 | low | Verrucomicrobiota | Verrucomicrobiia | Verrucomicrobiales | — | — | uncultured Verrucomicrobiales bacteriu | SAMEA9695074 | temperature | 10 | Yes | 4 | 2Fe-2S_thioredx;DNA_pol3_delta;P | — |
| 2026780.SAMN15870312.JACTNB010C | **clade11-B** | 265,4 | low | Planctomycetota | — | — | — | — | Planctomycetota bacterium | SAMN15870312 | anthropogenic | 10 | Yes | 4 | WCX;Abhydrolase_6;PFK;Hydrolas | — |
| 756272.SAMN00103628.CP002546_23 | **clade11-B** | 257,4 | low | Planctomycetota | Planctomycetia | Planctomycetales | Planctomycetaceae | Rubinisphaera | Rubinisphaera brasiliensis | SAMN00103628 | aquatic | 10 | Yes | 4 | IMPDH;RmaAD;PFK;Hydrolase | — |
| 904984.SAMEA6954088.CAIWEX0100 | **clade11-B** | 255,9 | low | Planctomycetota | Planctomycetia | Planctomycetales | Planctomycetaceae | Schlesneria | uncultured Schlesneria sp. | SAMEA6954088 | temperature | 2 | Yes | 2 | DUF1501 | — |
| 904984.SAMEA6951781.CAIKKW0100 | **clade11-B** | 255,5 | low | Planctomycetota | Planctomycetia | Planctomycetales | Planctomycetaceae | Schlesneria | uncultured Schlesneria sp. | SAMEA6951781 | temperature | 5 | Yes | 2 | Glycos_transf_3;Abhydrolase_2 | — |
| 904984.SAMEA6956427.CAIMFH0100 | **clade11-B** | 251,9 | low | Planctomycetota | Planctomycetia | Planctomycetales | Planctomycetaceae | Schlesneria | uncultured Schlesneria sp. | SAMEA6956427 | aquatic | 10 | Yes | 4 | MSMEG_6518_N;Peptidase_M50;l | — |
| 904984.SAMEA6954856.CAIQIR01000 | **clade11-B** | 251,4 | low | Planctomycetota | Planctomycetia | Planctomycetales | Planctomycetaceae | Schlesneria | uncultured Schlesneria sp. | SAMEA6954856 | aquatic | 10 | Yes | 4 | PseudoU_synth_2;Vma12;RsfS | — |
| 2052181.SAMD00166097.BJHP010000 | **clade11-B** | 248,7 | low | Planctomycetota | Planctomycetia | — | — | — | Planctomycetia bacterium | SAMD00166097 | aquatic | 10 | Yes | 4 | HPPK;Peptidase_M16_C;ATP_binc | — |
| 2026780.SAMN15013952.JABTUM010 | **clade11-B** | 248,6 | low | Planctomycetota | — | — | — | — | Planctomycetota bacterium | SAMN15013952 | temperature | 10 | Yes | 3 | TPR_19;Inositol_P;SGL | — |
| 2026780.SAMN10966916.VGZH010000 | **clade11-B** | 248,5 | low | Planctomycetota | — | — | — | — | Planctomycetota bacterium | SAMN10966916 | aquatic | 10 | Yes | 4 | DUF1501;ATP_bind_3;Peptidase_l | — |
| 2026779.SAMN14915429.JABJDJ0100 | **clade11-B** | 245,3 | low | Planctomycetota | Planctomycetia | Planctomycetales | Planctomycetaceae | — | Planctomycetaceae bacterium | SAMN14915429 | aquatic | 10 | Yes | 4 | GFO_IDH_MocA;AP_endonuc_2;S | — |
| 904984.SAMEA6955702.CAIULL01000 | **clade11-B** | 242,9 | low | Planctomycetota | Planctomycetia | Planctomycetales | Planctomycetaceae | Schlesneria | uncultured Schlesneria sp. | SAMEA6955702 | — | 10 | Yes | 0 | — | — |
| 904984.SAMEA6955890.CAIOKO0100 | **clade11-B** | 242,6 | low | Planctomycetota | Planctomycetia | Planctomycetales | Planctomycetaceae | Schlesneria | uncultured Schlesneria sp. | SAMEA6955890 | aquatic | 10 | Yes | 0 | — | — |
| 904984.SAMEA6943704.CAIVEI010000 | **clade11-B** | 240,1 | low | Planctomycetota | Planctomycetia | Planctomycetales | Planctomycetaceae | Schlesneria | uncultured Schlesneria sp. | SAMEA6943704 | aquatic | 6 | Yes | 3 | MJ0013;CHAT;SBP_bac_10 | — |
| 2026778.SAMN13893722.JAAYDY010C | **clade11-B** | 227,4 | low | Planctomycetota | Phycisphaerae | — | — | — | Phycisphaerae bacterium | SAMN13893722 | anthropogenic | 6 | Yes | 3 | Phage_tail_beta;Beta-prop_IFT122_ | — |
| 2026780.SAMN18118927.JAGNKM010 | **clade11-B** | 215,1 | low | Planctomycetota | — | — | — | — | Planctomycetota bacterium | SAMN18118927 | anthropogenic | 10 | Yes | 4 | LRR_14;Beta-prop_IFT122_1st;FG | — |
| 2026778.SAMD00166093.BJHL010000 | **clade11-B** | 188,1 | low | Planctomycetota | Phycisphaerae | — | — | — | Phycisphaerae bacterium | SAMD00166093 | aquatic | 10 | Yes | 4 | GT87;RNase_HII;Thymidylate_kin;E | — |
| 153809.SAMEA6949946.CAIMSU0100 | **clade11-B** | 179,7 | low | Pseudomonadota | — | — | — | — | uncultured Pseudomonadota bacterium | SAMEA6949946 | aquatic | 6 | Yes | 3 | — | — |
| 2026724.SAMN20307496.JAIOAS0100 | **clade11-B** | 174,6 | low | Chloroflexota | — | — | — | — | Chloroflexota bacterium | SAMN20307496 | temperature | 9 | Yes | 4 | Band_7;Peripla_BP_3;Response_re | — |
| 1895807.SAMN05660562.MKTT010000 | **clade11-B** | 158,3 | low | Planctomycetota | Planctomycetia | Planctomycetales | — | — | Planctomycetales bacterium 71-10 | SAMN05660562 | anthropogenic | 10 | Yes | 4 | MMPL;Band_7;DUF6298 | — |
| 2044587.SAMN01731007.KB822441_6 | **clade2_Ec107like** | 322,5 | high | Bacillota | Clostridia | Lachnospirales | Lachnospiraceae | Schaedlerella | Schaedlerella arabinosiphila | SAMN01731007 | host-associated | 11 | Yes | 4 | ECF-ribofla_trS;tRNA-synt_1b;BcrA | — |
| 297314.SAMEA8805696.CAJULL01000 | **clade2_Ec107like** | 321,7 | high | Bacillota | Clostridia | Lachnospirales | Lachnospiraceae | — | uncultured Lachnospiraceae bacterium | SAMEA8805696 | host-associated | 11 | Yes | 4 | DUF3267;tRNA-synt_1b;Glyco_hyc | — |
| 1121114.SAMN02441708.KB892637_1 | **clade2_Ec107like** | 321 | high | Bacillota | Clostridia | Lachnospirales | Lachnospiraceae | Blautia | Blautia producta | SAMN02441708 | host-associated | 11 | Yes | 4 | F420_ligase;Metallophos_2;Ham1p | — |
| 2025493.SAMN07460455.NQOF01000C | **clade2_Ec107like** | 319,1 | high | Bacillota | Clostridia | Lachnospirales | Lachnospiraceae | Blautia | Blautia hominis | SAMN07460455 | host-associated | 11 | Yes | 4 | Response_reg;Lipocalin_4;DUF740 | — |
| 1898203.SAMEA8801244.CAJSZT010C | **clade2_Ec107like** | 319 | high | Bacillota | Clostridia | Lachnospirales | Lachnospiraceae | — | Lachnospiraceae bacterium | SAMEA8801244 | host-associated | 11 | Yes | 4 | ECF-ribofla_trS;tRNA-synt_1b;FtsX | — |
| 1745713.SAMEA3959520.LT574838_1 | **clade2_Ec107like** | 319 | high | Bacillota | Clostridia | Lachnospirales | Lachnospiraceae | Bariatricus | Bariatricus massiliensis | SAMEA3959520 | host-associated | 11 | Yes | 4 | Glycos_transf_2;MazG;adh_short;A | — |
| 1410651.SAMN02743875.JHWJ010000 | **clade2_Ec107like** | 318,8 | high | Bacillota | Clostridia | Lachnospirales | Lachnospiraceae | Lacrimispora | Lacrimispora aerotolerans | SAMN02743875 | host-associated | 11 | Yes | 4 | PDH_N;DAHP_synth_1;Choline_bi | — |
| 1946596.SAMN06450376.DGRE01000C | **clade2_Ec107like** | 318,6 | high | Bacillota | Clostridia | Lachnospirales | Lachnospiraceae | Hungatella | Hungatella sp. UBA4396 | SAMN06450376 | anthropogenic | 11 | Yes | 4 | PDH_N;DAHP_synth_1;Choline_bi | — |
| 94868.SAMN05504968.MCIA01000032 | **clade2_Ec107like** | 317,5 | high | Bacillota | Clostridia | Lachnospirales | Lachnospiraceae | Lacrimispora | Lacrimispora algidixylanolytica | SAMN05504968 | anthropogenic | 11 | Yes | 4 | PDH_N;DAHP_synth_1;Choline_bi | — |
| 1898203.SAMN09901047.QXWY01000 | **clade2_Ec107like** | 317,4 | high | Bacillota | Clostridia | Lachnospirales | Lachnospiraceae | — | Lachnospiraceae bacterium | SAMN09901047 | host-associated | 11 | Yes | 4 | adh_short;tRNA-synt_1b;ECF-ribofl | — |
| 1111728.SAMN02440598.ATYS010000 | **clade2_Ec107like** | 317,3 | high | Pseudomonadota | Gammaproteobacteria | Enterobacterales | Budviciaceae | Budvicia | Budvicia aquatica | SAMN02440598 | host-associated | 11 | Yes | 4 | HD;Diacid_rec;GD_AH_C;MFS_1 | — |
| 1965569.SAMN06473635.NFKT010000 | **clade2_Ec107like** | 315,7 | high | Bacillota | Clostridia | Lachnospirales | Lachnospiraceae | Lachnoclostridium | Lachnoclostridium sp. An169 | SAMN06473635 | host-associated | 11 | Yes | 4 | DegV;DHDPS;FAD_binding_5;Fer2 | — |
| 904190.SAMEA5278550.CAAEFH0100 | **clade2_Ec107like** | 315,6 | high | Bacillota | Clostridia | Lachnospirales | Lachnospiraceae | Robinsoniella | uncultured Robinsoniella sp. | SAMEA5278550 | host-associated | 11 | Yes | 4 | Glyco_hydro_77;Acyltransferase;Fts | — |
| 537007.SAMN07340700.CP022413_28 | **clade2_Ec107like** | 315,3 | high | Bacillota | Clostridia | Lachnospirales | Lachnospiraceae | Blautia | Blautia hansenii | SAMN07340700 | host-associated | 11 | Yes | 4 | CBM_48;TetR_N;TatD_DNase;AI-2 | — |
| 297314.SAMEA7848117.CAJKQE0100 | **clade2_Ec107like** | 315,3 | high | Bacillota | Clostridia | Lachnospirales | Lachnospiraceae | — | uncultured Lachnospiraceae bacterium | SAMEA7848117 | host-associated | 11 | Yes | 0 | — | — |
| 1946599.SAMN06457310.DIJQ010000C | **clade2_Ec107like** | 314,5 | high | Bacillota | Clostridia | Lachnospirales | Lachnospiraceae | Hungatella | Hungatella sp. UBA5640 | SAMN06457310 | anthropogenic | 11 | Yes | 4 | His_kinase;Response_reg;DAHP_s | — |
| 904190.SAMEA5849789.CABPVW010C | **clade2_Ec107like** | 314,4 | high | Bacillota | Clostridia | Lachnospirales | Lachnospiraceae | Robinsoniella | uncultured Robinsoniella sp. | SAMEA5849789 | host-associated | 11 | Yes | 4 | Glyco_hydro_77;Acyltransferase;Fts | — |
| 1232440.SAMD00008809.BAHR02000C | **clade2_Ec107like** | 314 | high | Bacillota | Clostridia | Lachnospirales | Lachnospiraceae | Hungatella | Hungatella hathewayi | SAMD00008809 | host-associated | 11 | Yes | 4 | 2TM_P5A-ATPase;DUF2871;EPSF | — |
| 33035.SAMN02903323.JPJF01000011_ | **clade2_Ec107like** | 314 | high | Bacillota | Clostridia | Lachnospirales | Lachnospiraceae | Blautia | Blautia producta | SAMN02903323 | host-associated | 11 | Yes | 4 | DinB_2;Hydrolase_4;Condensation; | — |
| 742740.SAMN02463857.GL834320_36 | **clade2_Ec107like** | 313,9 | high | Bacillota | Clostridia | Lachnospirales | Lachnospiraceae | Otoolea | [Clostridium] symbiosum | SAMN02463857 | host-associated | 11 | Yes | 4 | GFO_IDH_MocA;LysR_substrate;N | — |
| 904190.SAMEA4890949.UQGT010000 | **clade2_Ec107like** | 313,3 | high | Bacillota | Clostridia | Lachnospirales | Lachnospiraceae | Robinsoniella | uncultured Robinsoniella sp. | SAMEA4890949 | host-associated | 10 | Yes | 4 | ABC_tran;FtsX;Acyltransferase;Gly | — |
| 297314.SAMEA7202538.CAJFUP0100C | **clade2_Ec107like** | 312,1 | high | Bacillota | Clostridia | Lachnospirales | Lachnospiraceae | — | uncultured Lachnospiraceae bacterium | SAMEA7202538 | host-associated | 9 | Yes | 4 | PTS-HPr;DUF6264;Adenylsucc_sy | — |
| 2559709.SAMN10863240.SPHP01000C | **clade2_Ec107like** | 312 | high | Bacillota | Clostridia | Eubacteriales | Clostridiaceae | Clostridium | Clostridium sp. 1001175st1 A10 | SAMN10863240 | host-associated | 11 | Yes | 4 | PDH_N;EPSP_synthase;GDE_C;L | — |
| 297314.SAMEA8805883.CAJUTU0100C | **clade2_Ec107like** | 311,7 | high | Bacillota | Clostridia | Lachnospirales | Lachnospiraceae | — | uncultured Lachnospiraceae bacterium | SAMEA8805883 | host-associated | 11 | Yes | 4 | RraA-like;GFO_IDH_MocA;DUF60 | — |
| 1304881.SAMN02597187.AZUI0100000 | **clade2_Ec107like** | 311,4 | high | Bacillota | Clostridia | Lachnospirales | Lachnospiraceae | Lacrimispora | Lacrimispora indolis | SAMN02597187 | host-associated | 11 | Yes | 4 | His_kinase;Response_reg;DAHP_s | — |
| 1286362.SAMN02745763.PGET010000 | **clade2_Ec107like** | 311 | high | Bacillota | Clostridia | Lachnospirales | Lachnospiraceae | Lacrimispora | Lacrimispora celerecrescens | SAMN02745763 | host-associated | 11 | Yes | 0 | — | — |
| 765821.SAMEA7848047.CAJJVI010000 | **clade2_Ec107like** | 311 | high | Bacillota | Clostridia | Lachnospirales | Lachnospiraceae | Blautia | uncultured Blautia sp. | SAMEA7848047 | host-associated | 7 | Yes | 3 | GWT1;Alpha-amylase;ToxN_toxin | — |
| 500632.SAMN00000730.DS995353_83 | **clade2_Ec107like** | 310,7 | high | Bacillota | Clostridia | Lachnospirales | Lachnospiraceae | Faecalimonas | Faecalimonas nexilis | SAMN00000730 | host-associated | 11 | Yes | 4 | Topoisom_bac;BcrAD_BadFG;Pseu | — |
| 1720294.SAMN09736885.QVLV010000 | **clade2_Ec107like** | 309,8 | high | Bacillota | Clostridia | Lachnospirales | Lachnospiraceae | Eisenbergiella | Eisenbergiella massiliensis | SAMN09736885 | host-associated | 11 | Yes | 4 | Glyoxalase;Metallophos;Flg_new;D/ | — |
| 1291051.SAMD00000437.BBAB010000 | **clade2_Ec107like** | 309,6 | high | Bacillota | Clostridia | Lachnospirales | Lachnospiraceae | Mediterraneibacter | Mediterraneibacter glycyrrhizinilyticus | SAMD00000437 | host-associated | 11 | Yes | 4 | Glyco_hydro_3;Glyco_hydro_79n | — |
| 1898203.SAMEA7848093.CAJKQB010C | **clade2_Ec107like** | 309,4 | high | Bacillota | Clostridia | Lachnospirales | Lachnospiraceae | — | Lachnospiraceae bacterium | SAMEA7848093 | host-associated | 11 | Yes | 4 | HATPase_c;Response_reg;Metallop | — |

| | | | | | | | | | | | | | | | | |
|---|---|---|---|---|---|---|---|---|---|---|---|---|---|---|---|---|
| 59620.SAMEA4890773.UPZT01000002 | **clade2_Ec107like** | 309 | high | Bacillota | Clostridia | Eubacteriales | Clostridiaceae | Clostridium | uncultured Clostridium sp. | SAMEA4890773 | host-associated | 11 | Yes | 4 | Hpt;HD_5;DAPDH_C;tRNA_edit | — |
| 2763667.SAMN15805280.CP060635_1 | **clade2_Ec107like** | 308,9 | high | Bacillota | Clostridia | Lachnospirales | Lachnospiraceae | Wansuia | Wansuia hejianensis | SAMN15805280 | host-associated | 11 | Yes | 4 | HTH_18;adh_short_C2;GFO_IDH_ | — |
| 1232458.SAMD00008692.BAIL0200001 | **clade2_Ec107like** | 307,8 | high | Bacillota | Clostridia | Eubacteriales | — | — | Clostridiales bacterium VE202-29 | SAMD00008692 | host-associated | 11 | Yes | 4 | DAPDH_C;Flg_new;SMAGP;tRNA_ | — |
| 77133.SAMEA7852881.CAJMPJ01000 | **clade2_Ec107like** | 307,7 | high | — | — | — | — | — | uncultured bacterium | SAMEA7852881 | — | 11 | Yes | 4 | PDH_N;DAHP_synth_1;Choline_bi | — |
| 457421.SAMN02463678.DS990263_33 | **clade2_Ec107like** | 306 | high | Bacillota | Clostridia | Eubacteriales | — | — | Clostridiales bacterium 1_7_47FAA | SAMN02463678 | host-associated | 11 | Yes | 0 | — | — |
| 358742.SAMN09734406.QVEX0100000 | **clade2_Ec107like** | 305,8 | high | Bacillota | Clostridia | Lachnospirales | Lachnospiraceae | Enterocloster | Enterocloster aldenensis | SAMN09734406 | host-associated | 11 | Yes | 4 | TetR_N;adh_short;AIM24;Cons_hy | — |
| 297314.SAMEA8805860.CAJUTQ0100 | **clade2_Ec107like** | 305,8 | high | Bacillota | Clostridia | Lachnospirales | Lachnospiraceae | — | uncultured Lachnospiraceae bacterium | SAMEA8805860 | host-associated | 9 | Yes | 3 | Acyl_transf_3;TatD_DNase;RrnaAD | — |
| 1123075.SAMN02441541.AUDP010000 | **clade2_Ec107like** | 305,7 | high | Bacillota | Clostridia | Eubacteriales | Oscillospiraceae | Ruminococcus | Ruminococcus gauvreauii | SAMN02441541 | host-associated | 11 | Yes | 4 | GMP_synt_C;AbiTii;MTTB;AP_end | — |
| 2838502.SAMN15816793.DWUX01000 | **clade2_Ec107like** | 305,3 | high | Bacillota | Clostridia | Lachnospirales | Lachnospiraceae | Blautia | Candidatus Blautia stercoripullorum | SAMN15816793 | host-associated | 11 | Yes | 4 | AI-2E_transport;TatD_DNase;Alpha | — |
| 742733.SAMN02463853.JH376424_86 | **clade2_Ec107like** | 304,2 | high | Bacillota | Clostridia | Lachnospirales | Lachnospiraceae | Enterocloster | Enterocloster citroniae | SAMN02463853 | host-associated | 11 | Yes | 4 | TetR_N;adh_short;AIM24;Cons_hy | — |
| 1796618.SAMN10864764.SGXF010000 | **clade2_Ec107like** | 303,9 | high | Bacillota | Clostridia | Lachnospirales | Lachnospiraceae | Cuneatibacter | Cuneatibacter caecimuris | SAMN10864764 | host-associated | 11 | Yes | 4 | Transketolase_N;MerR_1;APH;DHI | — |
| 411902.SAMN00627070.DS480695_87 | **clade2_Ec107like** | 303,4 | high | Bacillota | Clostridia | Lachnospirales | Lachnospiraceae | Enterocloster | Enterocloster bolteae | SAMN00627070 | host-associated | 11 | Yes | 4 | Flavoprotein;YoaP;BPD_transp_1 | — |
| 2559708.SAMN10863263.SPHO010000 | **clade2_Ec107like** | 303,3 | high | Bacillota | Clostridia | Eubacteriales | Clostridiaceae | Clostridium | Clostridium sp. 1001271st1 H5 | SAMN10863263 | host-associated | 11 | Yes | 4 | BPD_transp_1;Flavoprotein;DpaA_I | — |
| 1912897.SAMN09487132.CP030280_3 | **clade2_Ec107like** | 302,9 | high | Bacillota | Clostridia | Lachnospirales | Lachnospiraceae | Blautia | Blautia argi | SAMN09487132 | host-associated | 11 | Yes | 4 | Glyco_hydro_77;Zn_peptidase_2;C | — |
| 1898203.SAMN17800811.JAGZGI0100 | **clade2_Ec107like** | 302,9 | high | Bacillota | Clostridia | Lachnospirales | Lachnospiraceae | — | Lachnospiraceae bacterium | SAMN17800811 | host-associated | 11 | Yes | 4 | TetR_N;SpoIIID;Glyco_hydro_4 | — |
| 1122155.SAMN02745158.FQVI0100001 | **clade2_Ec107like** | 301,4 | high | Bacillota | Clostridia | Eubacteriales | Clostridiaceae | Lactonifactor | Lactonifactor longoviformis | SAMN02745158 | host-associated | 11 | Yes | 4 | Ldh_1_N;Malic_M;FAA_hydrolase;I | — |
| 562.SAMN14342434.CP050195_3503 | **clade2_Ec107like** | 301,3 | high | Pseudomonadota | Gammaproteobacteria | Enterobacterales | Enterobacteriaceae | Escherichia | Escherichia coli | SAMN14342434 | host-associated | 11 | Yes | 4 | dUTPase;SlmA-like_C;Pribosyltran; | — |
| 165186.SAMEA7202183.CAJFQJ01000 | **clade2_Ec107like** | 300,7 | high | Bacillota | Clostridia | Eubacteriales | Oscillospiraceae | Ruminococcus | uncultured Ruminococcus sp. | SAMEA7202183 | host-associated | 6 | Yes | 2 | Fer2_2;FAD_binding_5 | — |
| 297314.SAMEA8805492.CAJUFN0100 | **clade2_Ec107like** | 300,7 | high | Bacillota | Clostridia | Lachnospirales | Lachnospiraceae | — | uncultured Lachnospiraceae bacterium | SAMEA8805492 | host-associated | 11 | Yes | 4 | Zn_ribbon_2;GyrI-like;GIY-YIG;SseE | — |
| 297314.SAMEA7202275.CAJFIC010000 | **clade2_Ec107like** | 300,5 | high | Bacillota | Clostridia | Lachnospirales | Lachnospiraceae | — | uncultured Lachnospiraceae bacterium | SAMEA7202275 | host-associated | 11 | Yes | 4 | PTS-HPr;DUF6264;Adenylsucc_sy | — |
| 1879010.SAMN17800882.JAGZJB0100 | **clade2_Ec107like** | 300,2 | high | Bacillota | — | — | — | — | Bacillota bacterium | SAMN17800882 | host-associated | 11 | Yes | 4 | Peptidase_M17;X25_BaPul_like;Fo | — |
| 297314.SAMEA7202512.CAJFNX0100 | **clade2_Ec107like** | 299,6 | high | Bacillota | Clostridia | Lachnospirales | Lachnospiraceae | — | uncultured Lachnospiraceae bacterium | SAMEA7202512 | host-associated | 7 | Yes | 3 | AAA-ATPase_like;Trypsin_2;UTRA | — |
| 411463.SAMN00627091.DS264270_16 | **clade2_Ec107like** | 298 | high | Bacillota | Clostridia | Eubacteriales | Eubacteriaceae | Eubacterium | Eubacterium ventriosum | SAMN00627091 | host-associated | 11 | Yes | 4 | DUF2201_N;GmrSD_N;FOXO-TAD | — |
| 1658109.SAMEA3442603.LN852692_3 | **clade2_Ec107like** | 298 | high | Bacillota | Erysipelotrichia | Erysipelotrichales | Erysipelotrichaceae | Candidatus Stoquefich | Candidatus Stoquefichus sp. SB1 | SAMEA3442603 | host-associated | 11 | Yes | 4 | DUF3784;PucR;VanY;RBFA | — |
| 59620.SAMEA7847086.CAJLJW010000 | **clade2_Ec107like** | 297,8 | high | Bacillota | Clostridia | Eubacteriales | Clostridiaceae | Clostridium | uncultured Clostridium sp. | SAMEA7847086 | host-associated | 11 | Yes | 4 | SbcD_C;Cupin_2;Peptidase_M20;A | — |
| 610130.SAMEA7202544.CAJFOY01000 | **clade2_Ec107like** | 297 | high | Bacillota | Clostridia | Lachnospirales | Lachnospiraceae | Lacrimispora | Lacrimispora saccharolytica | SAMEA7202544 | host-associated | 11 | Yes | 4 | Big_3;Cupin_2;PRD | — |
| 224209.SAMEA8804963.CAJTLN01000 | **clade2_Ec107like** | 296,7 | high | Bacillota | Bacilli | — | — | — | uncultured Bacilli bacterium | SAMEA8804963 | host-associated | 11 | Yes | 4 | HTH_5;Hydrolase;APH;DUF1653 | — |
| 297314.SAMEA8805827.CAJURT0100 | **clade2_Ec107like** | 296,5 | high | Bacillota | Clostridia | Lachnospirales | Lachnospiraceae | — | uncultured Lachnospiraceae bacterium | SAMEA8805827 | host-associated | 11 | Yes | 4 | CAP;PfkB;BPD_transp_1 | — |
| 2044939.SAMEA5849627.CABJAP0100 | **clade2_Ec107like** | 296,4 | high | Bacillota | Clostridia | — | — | — | Clostridia bacterium | SAMEA5849627 | host-associated | 11 | Yes | 4 | Peptidase_S8;Acetyltransf_1;DUF1 | — |
| 224209.SAMEA8805129.CAJTRV0100 | **clade2_Ec107like** | 296 | high | Bacillota | Bacilli | — | — | — | uncultured Bacilli bacterium | SAMEA8805129 | host-associated | 9 | Yes | 4 | Ribosomal_L44;Acetyltransf_1;PHP | — |
| 172733.SAMEA7847762.CAJKVZ01000 | **clade2_Ec107like** | 294,9 | high | Bacillota | Clostridia | Eubacteriales | — | — | uncultured Eubacteriales bacterium | SAMEA7847762 | host-associated | 6 | Yes | 2 | FMN_dh;Branch_AA_trans | — |
| 2485925.SAMEA6150724.CACZGD010 | **clade2_Ec107like** | 294,9 | high | Bacillota | Clostridia | Eubacteriales | Oscillospiraceae | — | Oscillospiraceae bacterium | SAMEA6150724 | host-associated | 8 | Yes | 4 | HTH_28;ThiG | — |
| 2485925.SAMN11294983.SVTP010000 | **clade2_Ec107like** | 293,9 | high | Bacillota | Clostridia | Eubacteriales | Oscillospiraceae | — | Oscillospiraceae bacterium | SAMN11294983 | host-associated | 8 | Yes | 4 | RHS_repeat;MerR_1;Glyco_hydro_ | — |
| 2485925.SAMN16345939.JAFVXA0100 | **clade2_Ec107like** | 293,8 | high | Bacillota | Clostridia | Eubacteriales | Oscillospiraceae | — | Oscillospiraceae bacterium | SAMN16345939 | host-associated | 11 | Yes | 4 | Pan_kinase;ECF_trnsprt;DUF7575; | — |
| 172733.SAMEA8805310.CAJUBH0100 | **clade2_Ec107like** | 293,5 | high | Bacillota | Clostridia | Eubacteriales | — | — | uncultured Eubacteriales bacterium | SAMEA8805310 | host-associated | 11 | Yes | 4 | Peptidase_Prp;Ribosomal_L27;HD_ | — |
| 2838555.SAMN15816711.DXDD010000 | **clade2_Ec107like** | 293,3 | high | Bacillota | Clostridia | Lachnospirales | Lachnospiraceae | Eisenbergiella | Candidatus Eisenbergiella pullistercoris | SAMN15816711 | host-associated | 11 | Yes | 4 | DUF3592;Cellulase;DUF2268;DAP | — |
| 1898204.SAMEA8801320.CAJSZV0100 | **clade2_Ec107like** | 293,1 | high | Bacillota | Clostridia | Eubacteriales | Clostridiaceae | — | Clostridiaceae bacterium | SAMEA8801320 | host-associated | 11 | Yes | 4 | zf-IS66;Acyl_transf_3;SDF | — |
| 2044939.SAMN16346710.JAFXAN0100 | **clade2_Ec107like** | 293 | high | Bacillota | Clostridia | — | — | — | Clostridia bacterium | SAMN16346710 | host-associated | 10 | Yes | 4 | DNA_ligase_aden;WHD_RNase_R | — |
| 224209.SAMEA7202252.CAJFHJ01000 | **clade2_Ec107like** | 292,8 | high | Bacillota | Bacilli | — | — | — | uncultured Bacilli bacterium | SAMEA7202252 | host-associated | 8 | Yes | 4 | PIK3CG_ABD;DUF1292;CTP_synt | — |
| 1750510.SAMEA7202319.CAJFVE0100 | **clade2_Ec107like** | 292,8 | high | Bacillota | Clostridia | Lachnospirales | Lachnospiraceae | Eisenbergiella | uncultured Eisenbergiella sp. | SAMEA7202319 | host-associated | 11 | Yes | 4 | DAPDH_C;Cellulase;MFS_1 | — |
| 2049044.SAMN16346208.JAFWHJ0100 | **clade2_Ec107like** | 292,7 | high | Bacillota | Erysipelotrichia | Erysipelotrichales | Erysipelotrichaceae | — | Erysipelotrichaceae bacterium | SAMN16346208 | host-associated | 11 | Yes | 4 | DUF4230;Hydrolase_3;Hydrolase;R | — |
| 2485925.SAMN11294755.SVKV010000 | **clade2_Ec107like** | 292,5 | high | Bacillota | Clostridia | Eubacteriales | Oscillospiraceae | — | Oscillospiraceae bacterium | SAMN11294755 | host-associated | 6 | Yes | 2 | TatD_DNase;UDPGT | — |
| 1849603.SAMEA5278299.CAAFVQ010 | **clade2_Ec107like** | 292,3 | high | Pseudomonadota | Gammaproteobacteria | Enterobacterales | Enterobacteriaceae | — | Enterobacteriaceae bacterium | SAMEA5278299 | host-associated | 11 | Yes | 4 | RNase_PH;Pribosyltran;SlmA-like_( | — |
| 2485925.SAMN11294929.SVRN010000 | **clade2_Ec107like** | 292,3 | high | Bacillota | Clostridia | Eubacteriales | Oscillospiraceae | — | Oscillospiraceae bacterium | SAMN11294929 | host-associated | 10 | Yes | 4 | NADAR-DarT1;TetR_C_33;ABC_tr | — |
| 2838556.SAMN15816624.DXHY010000 | **clade2_Ec107like** | 292,2 | high | Bacillota | Clostridia | Lachnospirales | Lachnospiraceae | Eisenbergiella | Candidatus Eisenbergiella stercoraviur | SAMN15816624 | host-associated | 11 | Yes | 4 | Target_ACGX;DUF3592;Cellulase;I | — |
| 2763047.SAMN15808687.JACOPN0100 | **clade2_Ec107like** | 292,1 | high | Bacillota | Clostridia | Eubacteriales | — | Flintibacter | Flintibacter faecis | SAMN15808687 | host-associated | 11 | Yes | 4 | CobA_CobO_BtuR;PEP-utilizers_C | — |
| 224209.SAMEA8805448.CAJUBJ01000 | **clade2_Ec107like** | 291,7 | high | Bacillota | Bacilli | — | — | — | uncultured Bacilli bacterium | SAMEA8805448 | host-associated | 10 | Yes | 4 | ADP_ribosyl_GH;HH_RND_rel;APH | — |
| 297314.SAMEA4891325.UQVK0100001 | **clade2_Ec107like** | 291,5 | high | Bacillota | Clostridia | Lachnospirales | Lachnospiraceae | — | uncultured Lachnospiraceae bacterium | SAMEA4891325 | host-associated | 7 | Yes | 3 | DAPDH_C;DUF3592;MFS_1 | — |
| 297314.SAMEA8805098.CAJTPQ0100 | **clade2_Ec107like** | 291,5 | high | Bacillota | Clostridia | Lachnospirales | Lachnospiraceae | — | uncultured Lachnospiraceae bacterium | SAMEA8805098 | host-associated | 11 | Yes | 4 | HTH_18;Glyco_hydro_2_C;DUF53 | — |
| 157472.SAMEA7846648.CAJJRY01000 | **clade2_Ec107like** | 291,5 | high | Bacillota | Bacilli | Bacillales | — | — | uncultured Bacillales bacterium | SAMEA7846648 | host-associated | 7 | Yes | 3 | Adenine_glyco;FapA;Epimerase | — |
| 2720819.SAMEA7202238.CAJFGS0100 | **clade2_Ec107like** | 290,7 | high | Bacillota | Clostridia | — | — | Candidatus Avimonas | Candidatus Avimonas merdigallinarum | SAMEA7202238 | host-associated | 11 | Yes | 4 | PAD_porph;DUF7620;DUF2201_N | — |
| 172733.SAMEA5850774.CABKLX0100 | **clade2_Ec107like** | 290,7 | high | Bacillota | Clostridia | Eubacteriales | — | — | uncultured Eubacteriales bacterium | SAMEA5850774 | host-associated | 7 | Yes | 3 | HTH_3;La;GFO_IDH_MocA | — |
| 59620.SAMEA7202323.CAJFJI0100000 | **clade2_Ec107like** | 290,6 | high | Bacillota | Clostridia | Eubacteriales | Clostridiaceae | Clostridium | uncultured Clostridium sp. | SAMEA7202323 | host-associated | 11 | Yes | 4 | Radical_SAM;DUF4130;C_GCAxx( | — |
| 1924109.SAMN18359857.JAGHID0100 | **clade2_Ec107like** | 290,5 | high | Bacillota | Clostridia | Lachnospirales | Lachnospiraceae | Eisenbergiella | Eisenbergiella sp. | SAMN18359857 | host-associated | 5 | Yes | 3 | PfkB;AbiEii;DUF6088 | — |
| 59620.SAMEA7847304.CAJJPT010000 | **clade2_Ec107like** | 290,3 | high | Bacillota | Clostridia | Eubacteriales | Clostridiaceae | Clostridium | uncultured Clostridium sp. | SAMEA7847304 | host-associated | 7 | Yes | 3 | SMC_N;Fzo_mitofusin | — |
| 297314.SAMEA8805487.CAJUHR0100 | **clade2_Ec107like** | 290,2 | high | Bacillota | Clostridia | Lachnospirales | Lachnospiraceae | — | uncultured Lachnospiraceae bacterium | SAMEA8805487 | host-associated | 11 | Yes | 4 | BPD_transp_1;Peptidase_M20;Hep | — |
| 1879010.SAMN11294487.SVAM010000 | **clade2_Ec107like** | 290,1 | high | Bacillota | — | — | — | — | Bacillota bacterium | SAMN11294487 | host-associated | 11 | Yes | 4 | DUF951;ABG_transport;YchF-GTP | — |
| 2840815.SAMN15817011.DVGI010000 | **clade2_Ec107like** | 290 | high | Bacillota | Clostridia | — | — | Candidatus Fimenecus | Candidatus Fimenecus excrementaviu | SAMN15817011 | — | 7 | Yes | 0 | — | — |
| 2005360.SAMN18359906.JAGHKA0100 | **clade2_Ec107like** | 289,8 | high | Bacillota | Clostridia | Lachnospirales | Lachnospiraceae | Faecalicatena | Faecalicatena sp. | SAMN18359906 | host-associated | 11 | Yes | 4 | DegV;DHDPS;HTH_11;PP-binding | — |
| 2044939.SAMN16349910.JAGBSO0100 | **clade2_Ec107like** | 289,8 | high | Bacillota | Clostridia | — | — | — | Clostridia bacterium | SAMN16349910 | host-associated | 11 | Yes | 4 | Aldolase_II;SCP2;LbH_EIF2B;Pept | — |
| 165185.SAMEA4891232.UQRM010000 | **clade2_Ec107like** | 289,3 | high | Bacillota | Clostridia | Eubacteriales | Eubacteriaceae | Eubacterium | uncultured Eubacterium sp. | SAMEA4891232 | host-associated | 8 | Yes | 3 | HTH_3;PIN_8;Thioredoxin_3 | — |
| 1262908.PRJEB1041.HF993109_93 | **clade2_Ec107like** | 289,3 | high | Mycoplasmatota | Mollicutes | Mycoplasmatales | Mycoplasmataceae | Mycoplasma | Mycoplasma sp. CAG:956 | PRJEB1041 | host-associated | 11 | Yes | 4 | LRR_At5g56370;Transglut_core;HT | — |
| 2485925.SAMN11294845.SVOH010000 | **clade2_Ec107like** | 288,6 | high | Bacillota | Clostridia | Eubacteriales | Oscillospiraceae | — | Oscillospiraceae bacterium | SAMN11294845 | host-associated | 11 | Yes | 4 | His_Phos_1;RuvX;Acetyltransf_1;G | — |
| 658143.SAMEA4891343.UQVP0100001 | **clade2_Ec107like** | 288,5 | high | Mycoplasmatota | — | — | — | — | uncultured Mycoplasmatota bacterium | SAMEA4891343 | host-associated | 8 | Yes | 4 | DUF4006;Flg_new;HD_3 | — |
| 1903720.SAMEA5278229.CAAGCO010 | **clade2_Ec107like** | 288,4 | high | Bacillota | Bacilli | — | — | — | Bacilli bacterium | SAMEA5278229 | host-associated | 8 | Yes | 4 | B3_4;H-kinase_dim;NlpE | — |
| 707003.SAMEA7846686.CAJJLK01000 | **clade2_Ec107like** | 287,9 | high | Bacillota | Clostridia | Eubacteriales | Oscillospiraceae | — | uncultured Oscillospiraceae bacterium | SAMEA7846686 | host-associated | 6 | Yes | 2 | GGDEF | — |
| 2320105.SAMN10024802.RAYY010000 | **clade2_Ec107like** | 287,9 | high | — | — | — | — | — | bacterium D16-56 | SAMN10024802 | host-associated | 11 | Yes | 4 | AraC_binding;DUF2809;Peptidase_ | — |
| 1903720.SAMEA5278509.CAAEXI0100 | **clade2_Ec107like** | 287,8 | high | Bacillota | Bacilli | — | — | — | Bacilli bacterium | SAMEA5278509 | host-associated | 11 | Yes | 4 | Mur_ligase_M;DNA_binding_1;SLT | — |
| 1965578.SAMN06473650.NFKG010000 | **clade2_Ec107like** | 287,4 | high | Bacillota | Clostridia | Eubacteriales | Oscillospiraceae | Pseudoflavonifractor | Pseudoflavonifractor sp. An187 | SAMN06473650 | host-associated | 11 | Yes | 4 | PAD_porph;Pur_DNA_glyco;Sel1 | — |
| 1898207.SAMN19224892.JAHZBC0100 | **clade2_Ec107like** | 286,6 | high | Bacillota | Clostridia | Eubacteriales | — | — | Clostridiales bacterium | SAMN19224892 | host-associated | 11 | Yes | 4 | GntP_permease;PAS_6;2-Hacid_dh | — |
| 297314.SAMEA8805573.CAJUGD0100 | **clade2_Ec107like** | 286,6 | high | Bacillota | Clostridia | Lachnospirales | Lachnospiraceae | — | uncultured Lachnospiraceae bacterium | SAMEA8805573 | host-associated | 11 | Yes | 4 | Cas12f1-like_TNB;DUF2809;Peptic | — |
| 1751881.SAMEA8805622.CAJUJV0100 | **clade2_Ec107like** | 286,5 | high | Bacillota | Clostridia | Lachnospirales | Lachnospiraceae | Hungatella | uncultured Hungatella sp. | SAMEA8805622 | host-associated | 11 | Yes | 4 | Flagellin_C;DUF6510;CyanoTRADI | — |
| 41978.SAMN16345046.JAFUKO010000 | **clade2_Ec107like** | 286 | high | Bacillota | Clostridia | Eubacteriales | Oscillospiraceae | Ruminococcus | Ruminococcus sp. | SAMN16345046 | host-associated | 11 | Yes | 4 | Dockerin_1;PepSY;FIK;Y1_Tnp | — |
| 28121.SAMN16349382.JAGAYD010000 | **clade2_Ec107like** | 286 | high | Bacillota | Clostridia | Lachnospirales | Lachnospiraceae | Butyrivibrio | Butyrivibrio sp. | SAMN16349382 | host-associated | 11 | Yes | 4 | GtfA_EBD;MS_TIM-barrel;PPR_3 | — |
| 157472.SAMEA7847028.CAJLIR010000 | **clade2_Ec107like** | 285,9 | high | Bacillota | Bacilli | Bacillales | — | — | uncultured Bacillales bacterium | SAMEA7847028 | host-associated | 11 | Yes | 4 | Sporozoite_P67;ETRAMP;SesA | — |
| 2840820.SAMN15817229.DVML010000 | **clade2_Ec107like** | 285,6 | high | Mycoplasmatota | — | — | — | Candidatus Fimihabitar | Candidatus Fimihabitans intestinipullor | SAMN15817229 | host-associated | 11 | Yes | 4 | SmpB;RNB;NDK | — |
| 658143.SAMEA4891688.URIY0100001 | **clade2_Ec107like** | 285,6 | high | Mycoplasmatota | — | — | — | — | uncultured Mycoplasmatota bacterium | SAMEA4891688 | host-associated | 6 | Yes | 2 | LytR_cpsA_psr;DDE_Tnp_ISL3 | — |
| 59620.SAMEA7202342.CAJFSI0100000 | **clade2_Ec107like** | 285,4 | high | Bacillota | Clostridia | Eubacteriales | Clostridiaceae | Clostridium | uncultured Clostridium sp. | SAMEA7202342 | host-associated | 11 | Yes | 4 | FA_synthesis;C_GCAxxG_C_C;Lys | — |
| 172733.SAMEA4891262.UQSP0100000 | **clade2_Ec107like** | 285,3 | high | Bacillota | Clostridia | Eubacteriales | — | — | uncultured Eubacteriales bacterium | SAMEA4891262 | host-associated | 11 | Yes | 4 | O-antigen_lig;HTH_19;RNA_pol_I_ | — |

| | | | | | | | | | | | | | | | | |
|---|---|---|---|---|---|---|---|---|---|---|---|---|---|---|---|---|
| 2044939.SAMN16342067.JAFQCR0100 | clade2_Ec107like | 285 | high | Bacillota | Clostridia | — | — | — | Clostridia bacterium | SAMN16342067 | host-associated | 11 | Yes | 4 | TPP_enzyme_C;POR;GtaA_6_Hair | — |
| 331630.SAMEA6149063.CACWUS0100 | clade2_Ec107like | 284,8 | high | Bacillota | Erysipelotrichia | Erysipelotrichales | Erysipelotrichaceae | — | uncultured Erysipelotrichaceae bacteri | SAMEA6149063 | host-associated | 11 | Yes | 4 | PedC-like;KTI12;Peptidase_M19 | — |
| 2840670.SAMN15817049.DVMT010000 | clade2_Ec107like | 284,7 | high | Bacillota | Bacilli | — | — | Candidatus Aphodocol | Candidatus Aphodocola excrementigal | SAMN15817049 | host-associated | 11 | Yes | 4 | SLT;HMGL-like;Cyclophil_like2;Pep | — |
| 297314.SAMEA8805848.CAJUSJ01000 | clade2_Ec107like | 284,4 | high | Bacillota | Clostridia | Lachnospirales | Lachnospiraceae | — | uncultured Lachnospiraceae bacterium | SAMEA8805848 | host-associated | 11 | Yes | 4 | YoeB_toxin;Hydrolase_3;HemY_N | — |
| 2044939.SAMEA5278674.CAAFVV010 | clade2_Ec107like | 284,3 | high | Bacillota | Clostridia | — | — | — | Clostridia bacterium | SAMEA5278674 | host-associated | 9 | Yes | 4 | PMI_typeI_cat;SpoU_methylase;Na | — |
| 297314.SAMEA8805495.CAJUDH0100 | clade2_Ec107like | 283,7 | high | Bacillota | Clostridia | Lachnospirales | Lachnospiraceae | — | uncultured Lachnospiraceae bacterium | SAMEA8805495 | host-associated | 11 | Yes | 4 | PfkB;ROK;MobB;MoeA_N | — |
| 1263032.PRJEB1017.HF990861_60 | clade2_Ec107like | 283,3 | high | Bacillota | — | — | — | — | Firmicutes bacterium CAG:822 | PRJEB1017 | host-associated | 11 | Yes | 4 | Aldo_ket_red;HSP20;tRNA-synt_1c | — |
| 1265.SAMEA6148936.CACWPR010000 | clade2_Ec107like | 283,1 | high | Bacillota | Clostridia | Eubacteriales | Oscillospiraceae | Ruminococcus | Ruminococcus flavefaciens | SAMEA6148936 | host-associated | 7 | Yes | 3 | SdpI;DinB_2;PGK | — |
| 297314.SAMEA7202416.CAJFLQ01000 | clade2_Ec107like | 282,9 | high | Bacillota | Clostridia | Lachnospirales | Lachnospiraceae | — | uncultured Lachnospiraceae bacterium | SAMEA7202416 | host-associated | 11 | Yes | 4 | Fer4_22;HTH_Crp_2;Form_Nir_trar | — |
| 59620.SAMEA7846780.CAJJLQ010000 | clade2_Ec107like | 282,7 | high | Bacillota | Clostridia | Eubacteriales | Clostridiaceae | Clostridium | uncultured Clostridium sp. | SAMEA7846780 | host-associated | 11 | Yes | 4 | Methyltransf_4;tRNA-synt_1b;DUF | — |
| 2485925.SAMN11294846.SVOI0100000 | clade2_Ec107like | 282,4 | high | Bacillota | Clostridia | Eubacteriales | Oscillospiraceae | — | Oscillospiraceae bacterium | SAMN11294846 | host-associated | 11 | Yes | 4 | Helicase_dom;DUF2201_N;AAA_5 | — |
| 2485925.SAMEA7847870.CAJKFU0100 | clade2_Ec107like | 282,1 | high | Bacillota | Clostridia | Eubacteriales | Oscillospiraceae | — | Oscillospiraceae bacterium | SAMEA7847870 | host-associated | 11 | Yes | 4 | Helicase_C_2;ClpB_D2-small;ABC_ | — |
| 41978.SAMN16340732.JAFOED010000 | clade2_Ec107like | 281,8 | high | Bacillota | Clostridia | Eubacteriales | Oscillospiraceae | Ruminococcus | Ruminococcus sp. | SAMN16340732 | host-associated | 8 | Yes | 4 | DUF8255;PepSY;Acyl_transf_3 | — |
| 297314.SAMEA7202269.CAJFIE010000 | clade2_Ec107like | 281,7 | high | Bacillota | Clostridia | Lachnospirales | Lachnospiraceae | — | uncultured Lachnospiraceae bacterium | SAMEA7202269 | host-associated | 11 | Yes | 4 | MarR_2;Form_Nir_trans;HTH_Crp_ | — |
| 658143.SAMEA4891289.UQTP0100000 | clade2_Ec107like | 281,4 | high | Mycoplasmatota | — | — | — | — | uncultured Mycoplasmatota bacterium | SAMEA4891289 | host-associated | 11 | Yes | 4 | CofD;Peptidase_S55;Epimerase | — |
| 2485925.SAMN16927241.JAEDFW010 | clade2_Ec107like | 280,9 | high | Bacillota | Clostridia | Eubacteriales | Oscillospiraceae | — | Oscillospiraceae bacterium | SAMN16927241 | host-associated | 3 | Yes | 0 | — | — |
| 2721134.SAMEA7202493.CAJFNQ010 | clade2_Ec107like | 280,9 | high | Bacillota | Clostridia | — | — | Candidatus Metarumin | Candidatus Metaruminococcus galliste | SAMEA7202493 | host-associated | 11 | Yes | 4 | AcrF2;Flg_new_2;Fe-ADH;HrcA | — |
| 2060878.SAMN16344967.JAFUHN010 | clade2_Ec107like | 280,8 | high | Bacillota | Erysipelotrichia | Erysipelotrichales | Erysipelotrichaceae | Solobacterium | Solobacterium sp. | SAMN16344967 | host-associated | 11 | Yes | 4 | MerR_1;Imp-YgjV;Gate;Lactamase_ | — |
| 1265.SAMN04487832.FPJT01000001_ | clade2_Ec107like | 280,3 | high | Bacillota | Clostridia | Eubacteriales | Oscillospiraceae | Ruminococcus | Ruminococcus flavefaciens | SAMN04487832 | anthropogenic | 11 | Yes | 4 | HipA_C;Resolvase;GATase_6;Forn | — |
| 1898207.SAMN10183766.SFHJ010000 | clade2_Ec107like | 280,1 | high | Bacillota | Clostridia | Eubacteriales | — | — | Clostridiales bacterium | SAMN10183766 | host-associated | 11 | Yes | 4 | DUF4015;DUF554;Iso_dh;Aconitas | — |
| 331630.SAMEA6150187.CACYLN0100 | clade2_Ec107like | 280,1 | high | Bacillota | Erysipelotrichia | Erysipelotrichales | Erysipelotrichaceae | — | uncultured Erysipelotrichaceae bacteri | SAMEA6150187 | host-associated | 11 | Yes | 4 | Glyco_hydro_1;MFS_1_like;TelA;D | — |
| 297314.SAMEA8805230.CAJTTU01000 | clade2_Ec107like | 279,9 | high | Bacillota | Clostridia | Lachnospirales | Lachnospiraceae | — | uncultured Lachnospiraceae bacterium | SAMEA8805230 | host-associated | 11 | Yes | 4 | AAA_2;ClpS;Wzy_C | — |
| 1751881.SAMEA8805086.CAJTOT010 | clade2_Ec107like | 279,8 | high | Bacillota | Clostridia | Lachnospirales | Lachnospiraceae | Hungatella | uncultured Hungatella sp. | SAMEA8805086 | host-associated | 11 | Yes | 4 | LysR_substrate;Aldo_ket_red;MreB | — |
| 658143.SAMEA4892001.URUT0100002 | clade2_Ec107like | 279,7 | high | Mycoplasmatota | — | — | — | — | uncultured Mycoplasmatota bacterium | SAMEA4892001 | host-associated | 11 | Yes | 4 | Peptidase_C39_2;dUTPase_2;3HC | — |
| 1898203.SAMN17800722.JAGZCX010 | clade2_Ec107like | 279,4 | high | Bacillota | Clostridia | Lachnospirales | Lachnospiraceae | — | Lachnospiraceae bacterium | SAMN17800722 | host-associated | 11 | Yes | 4 | Mg_chelatase;C_GCAxxG_C_C;DU | — |
| 1898204.SAMN09901061.QXWR01000 | clade2_Ec107like | 279,3 | high | Bacillota | Clostridia | Eubacteriales | Clostridiaceae | — | Clostridiaceae bacterium | SAMN09901061 | host-associated | 11 | Yes | 4 | Lactamase_B;Radical_SAM;HATPa | — |
| 707003.SAMEA8805751.CAJUOA0100 | clade2_Ec107like | 278,6 | high | Bacillota | Clostridia | Eubacteriales | Oscillospiraceae | — | uncultured Oscillospiraceae bacterium | SAMEA8805751 | host-associated | 6 | Yes | 2 | PAPS_reduct;DUF6110 | — |
| 2044939.SAMN16344038.JAFSXU0100 | clade2_Ec107like | 277,9 | high | Bacillota | Clostridia | — | — | — | Clostridia bacterium | SAMN16344038 | host-associated | 11 | Yes | 4 | Y1_Tnp;PrsW-protease;GATase;His | — |
| 172733.SAMEA7846782.CAJLHB01000 | clade2_Ec107like | 277,8 | high | Bacillota | Clostridia | Eubacteriales | — | — | uncultured Eubacteriales bacterium | SAMEA7846782 | host-associated | 7 | Yes | 3 | IU_nuc_hydro;HTH_18;Transpeptid | — |
| 707003.SAMEA5278737.CAAGBT0100 | clade2_Ec107like | 277,7 | high | Bacillota | Clostridia | Eubacteriales | Oscillospiraceae | — | uncultured Oscillospiraceae bacterium | SAMEA5278737 | host-associated | 9 | Yes | 4 | PDDEXK_4;DUF2203;GTP_EFTU | — |
| 1898207.SAMN19225273.DXYE010001 | clade2_Ec107like | 277,3 | high | Bacillota | Clostridia | Eubacteriales | — | — | Clostridiales bacterium | SAMN19225273 | — | 4 | Yes | 0 | — | — |
| 707003.SAMEA5278732.CAAFCU0100 | clade2_Ec107like | 276,9 | high | Bacillota | Clostridia | Eubacteriales | Oscillospiraceae | — | uncultured Oscillospiraceae bacterium | SAMEA5278732 | — | 11 | Yes | 0 | — | — |
| 1965629.SAMN06473716.NFIQ010000 | clade2_Ec107like | 276,7 | high | Bacillota | Clostridia | Eubacteriales | Oscillospiraceae | Flavonifractor | Flavonifractor sp. An306 | SAMN06473716 | host-associated | 11 | Yes | 4 | TP_0679;zf-like;DUF4367 | — |
| 165186.SAMEA4891392.UQXS010001 | clade2_Ec107like | 276,3 | high | Bacillota | Clostridia | Eubacteriales | Oscillospiraceae | Ruminococcus | uncultured Ruminococcus sp. | SAMEA4891392 | host-associated | 7 | Yes | 4 | Form-deh_trans;GATase_6;Phage_ | — |
| 224209.SAMEA8805494.CAJUCD0100 | clade2_Ec107like | 276,1 | high | Bacillota | Bacilli | — | — | — | uncultured Bacilli bacterium | SAMEA8805494 | host-associated | 11 | Yes | 4 | FXR_C3;Acetyltransf_7;YchF-GTP | — |
| 172733.SAMEA4891435.UQZJ0100003 | clade2_Ec107like | 275,4 | high | Bacillota | Clostridia | Eubacteriales | — | — | uncultured Eubacteriales bacterium | SAMEA4891435 | host-associated | 10 | Yes | 4 | PAD_porph;DSPc;DUF1722 | — |
| 1193532.SAMEA7848332.CAJKYR010 | clade2_Ec107like | 275,2 | high | Bacillota | Clostridia | Eubacteriales | Butyricicoccaceae | Butyricicoccus | uncultured Butyricicoccus sp. | SAMEA7848332 | host-associated | 11 | Yes | 4 | HTH_3;Pyrophosphatase;MFS_2;B | — |
| 2044939.SAMN16345817.JAFVSI01000 | clade2_Ec107like | 275,2 | high | Bacillota | Clostridia | — | — | — | Clostridia bacterium | SAMN16345817 | host-associated | 7 | Yes | 3 | TMEM164;SASA;HTH_18 | — |
| 707003.SAMEA7847196.CAJLTB01000 | clade2_Ec107like | 275,1 | high | Bacillota | Clostridia | Eubacteriales | Oscillospiraceae | — | uncultured Oscillospiraceae bacterium | SAMEA7847196 | — | 11 | Yes | 0 | — | — |
| 1506.SAMN17801316.JAGZZT0100000 | clade2_Ec107like | 275,1 | high | Bacillota | Clostridia | Eubacteriales | Clostridiaceae | Clostridium | Clostridium sp. | SAMN17801316 | host-associated | 11 | Yes | 4 | SbcD_C;Cupin_2;Peptidase_M20;A | — |
| 41978.SAMN16346810.JAFXEJ010000 | clade2_Ec107like | 274,7 | high | Bacillota | Clostridia | Eubacteriales | Oscillospiraceae | Ruminococcus | Ruminococcus sp. | SAMN16346810 | host-associated | 7 | Yes | 3 | AGA-YXIM_GBD | — |
| 165186.SAMEA8805374.CAJTWZ0100 | clade2_Ec107like | 274,6 | high | Bacillota | Clostridia | Eubacteriales | Oscillospiraceae | Ruminococcus | uncultured Ruminococcus sp. | SAMEA8805374 | host-associated | 11 | Yes | 4 | NikA-like;Relaxase;SH3_3;DUF417 | — |
| 297314.SAMEA6152050.CADBFE0100 | clade2_Ec107like | 274,5 | high | Bacillota | Clostridia | Lachnospirales | Lachnospiraceae | — | uncultured Lachnospiraceae bacterium | SAMEA6152050 | host-associated | 11 | Yes | 4 | AAA_5;DUF2201_N;Dpy19;Zn_Rib | — |
| 1898207.SAMN10183782.SFIY0100002 | clade2_Ec107like | 274,4 | high | Bacillota | Clostridia | Eubacteriales | — | — | Clostridiales bacterium | SAMN10183782 | — | 11 | Yes | 0 | — | — |
| 1410672.SAMN02744011.JHXI0100001 | clade2_Ec107like | 274,2 | high | Bacillota | Clostridia | Eubacteriales | Oscillospiraceae | Ruminococcus | Ruminococcus flavefaciens | SAMN02744011 | host-associated | 11 | Yes | 4 | Sigma70_r4_2;Hydrolase_4;Pro_C | — |
| 707003.SAMEA5851660.CABLZZ01000 | clade2_Ec107like | 274,2 | high | Bacillota | Clostridia | Eubacteriales | Oscillospiraceae | — | uncultured Oscillospiraceae bacterium | SAMEA5851660 | — | 11 | Yes | 0 | — | — |
| 297314.SAMEA8805828.CAJUPT01000 | clade2_Ec107like | 273,9 | high | Bacillota | Clostridia | Lachnospirales | Lachnospiraceae | — | uncultured Lachnospiraceae bacterium | SAMEA8805828 | host-associated | 11 | Yes | 4 | AAA_2;ClpS;VanZ;Wzy_C | — |
| 876091.SAMEA8805491.CAJUDV0100 | clade2_Ec107like | 273,8 | high | Bacillota | Clostridia | Eubacteriales | Oscillospiraceae | Oscillibacter | uncultured Oscillibacter sp. | SAMEA8805491 | host-associated | 10 | Yes | 4 | PCRF;SLH;Sigma70_r2;zf-HC2 | — |
| 1262910.PRJEB727.FR890955_127 | clade2_Ec107like | 273,8 | high | Bacillota | Clostridia | Eubacteriales | Oscillospiraceae | Oscillibacter | Oscillibacter sp. CAG:155 | PRJEB727 | host-associated | 8 | Yes | 4 | Rhomboid;S-AdoMet_synt_C;CSD; | — |
| 1265.SAMEA6150373.CACYSQ010000 | clade2_Ec107like | 273,6 | high | Bacillota | Clostridia | Eubacteriales | Oscillospiraceae | Ruminococcus | Ruminococcus flavefaciens | SAMEA6150373 | host-associated | 11 | Yes | 4 | Spore_permease;GerA;Phage_integ | — |
| 142586.SAMN18359832.JAGHHE0100 | clade2_Ec107like | 273,3 | high | Bacillota | Clostridia | Eubacteriales | Eubacteriaceae | Eubacterium | Eubacterium sp. | SAMN18359832 | host-associated | 7 | Yes | 3 | Acetyltransf_7;AAA_30;Methyltrans | — |
| 1689270.SAMEA5138309.LR130816_6 | clade2_Ec107like | 272,8 | high | Bacillota | Clostridia | Eubacteriales | — | Intestinimonas | Intestinimonas timonensis | SAMEA5138309 | host-associated | 11 | Yes | 4 | Lyx_isomer;PfkB;MarR;ABC_memb | — |
| 876091.SAMEA4891362.UWSF010000 | clade2_Ec107like | 272,7 | high | Bacillota | Clostridia | Eubacteriales | Oscillospiraceae | Oscillibacter | uncultured Oscillibacter sp. | SAMEA4891362 | — | 11 | Yes | 0 | — | — |
| 59620.SAMEA7847350.CAJLVU010000 | clade2_Ec107like | 272,7 | high | Bacillota | Clostridia | Eubacteriales | Clostridiaceae | Clostridium | uncultured Clostridium sp. | SAMEA7847350 | host-associated | 11 | Yes | 4 | AMP-binding;3HCDH_N;FerA;Pepti | — |
| 707003.SAMEA4890827.UQBX0100000 | clade2_Ec107like | 272,5 | high | Bacillota | Clostridia | Eubacteriales | Oscillospiraceae | — | uncultured Oscillospiraceae bacterium | SAMEA4890827 | host-associated | 11 | Yes | 4 | Cupin_2;Pectate_lyase_5;VCA004 | — |
| 172733.SAMEA7847389.CAJKPX01000 | clade2_Ec107like | 272,4 | high | Bacillota | Clostridia | Eubacteriales | — | — | uncultured Eubacteriales bacterium | SAMEA7847389 | — | 11 | Yes | 4 | BNR_2;HTH_18;Cellulase | — |
| 876091.SAMEA8805735.CAJULS01000 | clade2_Ec107like | 271,1 | high | Bacillota | Clostridia | Eubacteriales | Oscillospiraceae | Oscillibacter | uncultured Oscillibacter sp. | SAMEA8805735 | host-associated | 8 | Yes | 4 | RBR;SLH;Sigma70_r2;Ribosomal_ | — |
| 2049044.SAMN11294514.SVBN010000 | clade2_Ec107like | 270,7 | high | Bacillota | Erysipelotrichia | Erysipelotrichales | Erysipelotrichaceae | — | Erysipelotrichaceae bacterium | SAMN11294514 | host-associated | 10 | Yes | 4 | Peptidase_C69;FMN_red;FMN_dh | — |
| 707003.SAMEA7202338.CAJFJV01000 | clade2_Ec107like | 270 | high | Bacillota | Clostridia | Eubacteriales | Oscillospiraceae | — | uncultured Oscillospiraceae bacterium | SAMEA7202338 | host-associated | 11 | Yes | 4 | Asparaginase;BetaGal_ABD_1;Bcr | — |
| 2044939.SAMN16342657.JAFQVO010 | clade2_Ec107like | 269,1 | high | Bacillota | Clostridia | — | — | — | Clostridia bacterium | SAMN16342657 | host-associated | 5 | Yes | 3 | Pribosyltran;Transket_pyr;Transket | — |
| 876091.SAMEA8805797.CAJUQM0100 | clade2_Ec107like | 268,2 | high | Bacillota | Clostridia | Eubacteriales | Oscillospiraceae | Oscillibacter | uncultured Oscillibacter sp. | SAMEA8805797 | host-associated | 10 | Yes | 4 | PTPS_related;GDPD;SLH;PCRF | — |
| 411475.SAMN00627089.JH417816_7 | clade2_Ec107like | 268 | high | Bacillota | Clostridia | Eubacteriales | Oscillospiraceae | Flavonifractor | Flavonifractor plautii | SAMN00627089 | host-associated | 11 | Yes | 4 | AIM24;AMP-binding;LuxC | — |
| 2049044.SAMN16343168.JAFRQF010 | clade2_Ec107like | 267,9 | high | Bacillota | Erysipelotrichia | Erysipelotrichales | Erysipelotrichaceae | — | Erysipelotrichaceae bacterium | SAMN16343168 | host-associated | 10 | Yes | 4 | AAA_14;DSPc;DUF4371;PerC | — |
| 876091.SAMEA5279433.CAAGJQ0100 | clade2_Ec107like | 267,1 | high | Bacillota | Clostridia | Eubacteriales | Oscillospiraceae | Oscillibacter | uncultured Oscillibacter sp. | SAMEA5279433 | host-associated | 7 | Yes | 4 | MatE;HerA_C;SLH;PCRF | — |
| 2584680.SAMN11946324.WNAN01000 | clade2_Ec107like | 267,1 | high | Bacillota | Clostridia | Eubacteriales | Oscillospiraceae | Pseudoflavonifractor | Pseudoflavonifractor sp. BIOML-A18 | SAMN11946324 | host-associated | 11 | Yes | 4 | Molybdopterin;FdhD-NarQ;CutC;Tr | — |
| 2763673.SAMN15805297.JACRTB010 | clade2_Ec107like | 266,7 | high | Bacillota | Clostridia | Eubacteriales | Oscillospiraceae | Yanshouia | Yanshouia hominis | SAMN15805297 | host-associated | 11 | Yes | 4 | ECF_trnsprt;ATP_bind_3;RNase_Y | — |
| 1265.SAMEA6148735.CACWHY010000 | clade2_Ec107like | 263,8 | high | Bacillota | Clostridia | Eubacteriales | Oscillospiraceae | Ruminococcus | Ruminococcus flavefaciens | SAMEA6148735 | anthropogenic | 11 | Yes | 4 | DUF6198;DUF4832;HATPase_c;R | — |
| 41978.SAMN16344534.JAFTQW01000 | clade2_Ec107like | 262,6 | high | Bacillota | Clostridia | Eubacteriales | Oscillospiraceae | Ruminococcus | Ruminococcus sp. | SAMN16344534 | host-associated | 11 | Yes | 4 | GATase_6;PlyB_C;Response_reg; | — |
| 1898207.SAMN19225266.DYCV010000 | clade2_Ec107like | 262,1 | high | Bacillota | Clostridia | Eubacteriales | — | — | Clostridiales bacterium | SAMN19225266 | host-associated | 11 | Yes | 4 | PDDEXK_4;Metallophos;AAA_lid_ | — |
| 2049044.SAMN16343007.JAFRKA010 | clade2_Ec107like | 261,7 | high | Bacillota | Erysipelotrichia | Erysipelotrichales | Erysipelotrichaceae | — | Erysipelotrichaceae bacterium | SAMN16343007 | host-associated | 11 | Yes | 4 | SepF;DSPc;DUF4379;MazG-like | — |
| 165186.SAMEA6150165.CACYKN0100 | clade2_Ec107like | 258,4 | high | Bacillota | Clostridia | Eubacteriales | Oscillospiraceae | Ruminococcus | uncultured Ruminococcus sp. | SAMEA6150165 | host-associated | 11 | Yes | 4 | DUF6311;IU_nuc_hydro;DUF308 | — |
| 2838570.SAMN15816904.DXDN010000 | clade2_Ec107like | 253,1 | high | Bacillota | Clostridia | Eubacteriales | Eubacteriaceae | Eubacterium | Candidatus Eubacterium faecigallinaru | SAMN15816904 | host-associated | 11 | Yes | 4 | EamA;Zn_ribbon_NrdR;PsbP_2;Sig | — |
| 165186.SAMEA6151767.CADAUE0100 | clade2_Ec107like | 250,3 | high | Bacillota | Clostridia | Eubacteriales | Oscillospiraceae | Ruminococcus | uncultured Ruminococcus sp. | SAMEA6151767 | host-associated | 11 | Yes | 4 | Nmad3;Nmad2;WYL;HI_0787_N | — |
| 41978.SAMN16342822.JAFRBX010000 | clade2_Ec107like | 248 | high | Bacillota | Clostridia | Eubacteriales | Oscillospiraceae | Ruminococcus | Ruminococcus sp. | SAMN16342822 | host-associated | 11 | Yes | 4 | Virul_fac_BrkB;PDDEXK_4;WYL;P | — |
| 165186.SAMEA6152540.CADBYC0100 | clade2_Ec107like | 248 | high | Bacillota | Clostridia | Eubacteriales | Oscillospiraceae | Ruminococcus | uncultured Ruminococcus sp. | SAMEA6152540 | host-associated | 11 | Yes | 4 | WYL;HI_0787_N | — |
| 165186.SAMEA6149501.CACXLM0100 | clade2_Ec107like | 242,6 | high | Bacillota | Clostridia | Eubacteriales | Oscillospiraceae | Ruminococcus | uncultured Ruminococcus sp. | SAMEA6149501 | host-associated | 7 | Yes | 3 | Zn_dep_PLPC;RIFIN;Sigma70_r3 | — |
| 41978.SAMN16342855.JAFRDE010000 | clade2_Ec107like | 242,5 | high | Bacillota | Clostridia | Eubacteriales | Oscillospiraceae | Ruminococcus | Ruminococcus sp. | SAMN16342855 | host-associated | 11 | Yes | 4 | Peptidase_M15;S-AdoMet_lyase;Fe | — |
| 1689265.SAMEA7202227.CAJFGQ010 | clade2_Ec107like | 241,1 | high | Bacillota | Clostridia | Eubacteriales | — | Intestinimonas | uncultured Intestinimonas sp. | SAMEA7202227 | host-associated | 11 | Yes | 4 | Peptidase_M56;Penicillinase_R;bP | — |
| 1002368.SAMN02299454.JH417862_4 | clade2_Ec107like | 224,9 | low | Pseudomonadota | Gammaproteobacteria | Enterobacterales | Enterobacteriaceae | Yokenella | Yokenella regensburgei | SAMN02299454 | host-associated | 10 | Yes | 4 | FGGY_C;AraC_binding;VTT_dom | — |

| | | | | | | | | | | | | | | | | |
|---|---|---|---|---|---|---|---|---|---|---|---|---|---|---|---|---|
| 2485925.SAMEA6150216.CACYMP010 | **clade2_Ec107like** | 210,2 | low | Bacillota | Clostridia | Eubacteriales | Oscillospiraceae | — | Oscillospiraceae bacterium | SAMEA6150216 | host-associated | 10 | Yes | 4 | OGG_N;Abhydrolase_3;Acetyltrans | — |
| 2026780.SAMN10222782.RFON01000( | **clade2_Ec107like** | 192 | low | Planctomycetota | — | — | — | — | Planctomycetota bacterium | SAMN10222782 | aquatic | 4 | Yes | 3 | N6_N4_Mtase;TauE | — |
| 296830.SAMEA6951688.CAIOSQ0100( | **clade2_Ec107like** | 186,1 | low | Pseudomonadota | Alphaproteobacteria | Rhodospirillales | — | — | uncultured Rhodospirillales bacterium | SAMEA6951688 | aquatic | 5 | Yes | 2 | AAA_5 | — |
| 945550.SAMN02952919.AEVT0100002 | **typeI-A** | 341,6 | high | Pseudomonadota | Gammaproteobacteria | Vibrionales | Vibrionaceae | Vibrio | Vibrio sinaloensis | SAMN02952919 | aquatic | 11 | Yes | 4 | SMC_N;Ribosomal_L25p;Glycos_tr | — |
| 1806667.SAMEA4029000.FLRA010000 | **typeI-A** | 338 | high | Pseudomonadota | Gammaproteobacteria | Oceanospirillales | Oceanospirillaceae | Marinomonas | Marinomonas gallaica | SAMEA4029000 | aquatic | 11 | Yes | 4 | SMC_N;TrbL;AF_1314_C | — |
| 349965.SAMN00005354.AALF0200000 | **typeI-A** | 333,5 | high | Pseudomonadota | Gammaproteobacteria | Enterobacterales | Yersiniaceae | Yersinia | Yersinia intermedia | SAMN00005354 | anthropogenic | 11 | Yes | 4 | SF0329;SMC_N;Wzy_C;FaeA | — |
| 1122212.SAMN02440684.AULO01000( | **typeI-A** | 330,9 | high | Pseudomonadota | Gammaproteobacteria | Oceanospirillales | Oceanospirillaceae | Marinospirillum | Marinospirillum minutulum | SAMN02440684 | anthropogenic | 11 | Yes | 4 | MmeI_Mtase;MmeI_N;SMC_N | — |
| 1196094.SAMN03081540.CP007446_1 | **typeI-A** | 328,7 | high | Pseudomonadota | Betaproteobacteria | Neisseriales | Neisseriaceae | Snodgrassella | Snodgrassella alvi | SAMN03081540 | host-associated | 11 | Yes | 4 | SMC_N;FTHFS;TAL_FSA | — |
| 181674.SAMEA6943824.CAIOMN0100( | **typeI-A** | 325,7 | high | Pseudomonadota | Gammaproteobacteria | Methylococcales | Methylococcaceae | — | uncultured Methylococcaceae bacteriu | SAMEA6943824 | — | 4 | Yes | 0 | — | — |
| 181674.SAMEA6955042.CAIMEB0100( | **typeI-A** | 324,5 | high | Pseudomonadota | Gammaproteobacteria | Methylococcales | Methylococcaceae | — | uncultured Methylococcaceae bacteriu | SAMEA6955042 | aquatic | 3 | Yes | 2 | SMC_N;HNH_5 | — |
| 181674.SAMEA6949539.CAIPDT01000 | **typeI-A** | 320,8 | high | Pseudomonadota | Gammaproteobacteria | Methylococcales | Methylococcaceae | — | uncultured Methylococcaceae bacteriu | SAMEA6949539 | temperature | 11 | Yes | 4 | RepD-like_N;SMC_N;ESAG1;GmrS | — |
| 154336.SAMEA9694770.CAJXQM0100 | **typeI-A** | 318,8 | high | Pseudomonadota | Gammaproteobacteria | Alteromonadales | Colwelliaceae | Colwellia | uncultured Colwellia sp. | SAMEA9694770 | temperature | 7 | Yes | 3 | Phage_GPA;HTH_29 | — |
| 326537.SAMN06626649.NBOE010000( | **typeI-A** | 318,8 | high | Pseudomonadota | Gammaproteobacteria | Alteromonadales | Colwelliaceae | Colwellia | Colwellia polaris | SAMN06626649 | — | 11 | Yes | 0 | — | — |
| 2136183.SAMN08772518.PYIX020000( | **typeI-A** | 318 | high | Pseudomonadota | Gammaproteobacteria | Moraxellales | Moraxellaceae | Acinetobacter | Acinetobacter sichuanensis | SAMN08772518 | host-associated | 11 | Yes | 4 | RepD-like_N;SMC_N;NAD_kinase_ | — |
| 243277.SAMN02603969.AE003852_22 | **typeI-A** | 316,8 | high | Pseudomonadota | Gammaproteobacteria | Vibrionales | Vibrionaceae | Vibrio | Vibrio cholerae | SAMN02603969 | host-associated | 11 | Yes | 4 | LysR_substrate;TauE;SMC_N;HNH | — |
| 1122619.SAMN02441372.KB892326_9 | **typeI-A** | 315,5 | high | Pseudomonadota | Betaproteobacteria | Burkholderiales | Alcaligenaceae | Oligella | Oligella ureolytica | SAMN02441372 | host-associated | 10 | Yes | 4 | SMC_N;Arm-DNA-bind_3;Phage_ir | — |
| 412437.SAMN08324222.PNRD010000( | **typeI-A** | 313,4 | high | Pseudomonadota | Gammaproteobacteria | Chromatiales | Chromatiaceae | Rheinheimera | Rheinheimera aquimaris | SAMN08324222 | aquatic | 11 | Yes | 4 | RepD-like_N;SMC_N;TetR_C_13;A | — |
| 659.SAMN07327695.PYMQ01000073_ | **typeI-A** | 311,9 | high | Pseudomonadota | Gammaproteobacteria | Vibrionales | Vibrionaceae | Photobacterium | Photobacterium phosphoreum | SAMN07327695 | host-associated | 11 | Yes | 4 | Cas_Csy4;PDDEXK_11;SMC_N;R | — |
| 156424.SAMEA6595137.CADEDJ0100( | **typeI-A** | 294,9 | high | Pseudomonadota | Betaproteobacteria | Nitrosomonadales | Nitrosomonadaceae | Nitrosomonas | uncultured Nitrosomonas sp. | SAMEA6595137 | anthropogenic | 6 | Yes | 2 | Csf1_N;STM3845 | — |
| 1095744.SAMN00761805.AJSW01000( | **typeI-A** | 286 | high | Pseudomonadota | Gammaproteobacteria | Pasteurellales | Pasteurellaceae | Haemophilus | Haemophilus parahaemolyticus | SAMN00761805 | host-associated | 3 | Yes | 2 | RepD-like_N;AAA_15 | — |
| 707232.SAMN00113594.GG770443_1 | **typeI-A** | 282,8 | high | Pseudomonadota | Gammaproteobacteria | Moraxellales | Moraxellaceae | Acinetobacter | Acinetobacter haemolyticus | SAMN00113594 | anthropogenic | 6 | Yes | 2 | Csp | — |
| 762966.SAMN00189152.GL883764_17 | **typeI-A** | 275,2 | low | Pseudomonadota | Betaproteobacteria | Burkholderiales | Sutterellaceae | Parasutterella | Parasutterella excrementihominis | SAMN00189152 | host-associated | 10 | Yes | 4 | AAA_21;AAA_19;SMC_N | — |
| 1035188.SAMN00621708.AFUV010000 | **typeI-A** | 265,7 | low | Pseudomonadota | Gammaproteobacteria | Pasteurellales | Pasteurellaceae | Haemophilus | Haemophilus pittmaniae | SAMN00621708 | host-associated | 10 | Yes | 4 | Importin_rep;GDPD;DUF417;MIP | — |
| 286133.SAMEA7846561.CAJJSA01000 | **typeI-A** | 259,3 | low | Pseudomonadota | Betaproteobacteria | Burkholderiales | Sutterellaceae | Sutterella | uncultured Sutterella sp. | SAMEA7846561 | host-associated | 5 | Yes | 2 | SMC_N | — |
| 2796373.SAMN17054785.JAEKED010( | **typeI-B1** | 282,8 | high | Pseudomonadota | Gammaproteobacteria | Pseudomonadales | Pseudomonadaceae | Pseudomonas | Pseudomonas sp. TH07 | SAMN17054785 | host-associated | 11 | Yes | 4 | Mob_Pre;AAA_21;SHQ1;DUF7982 | — |
| 2760933.SAMN15642185.JACJDS0100 | **typeI-B1** | 280,4 | high | Pseudomonadota | Gammaproteobacteria | Alteromonadales | Pseudoalteromonadaceae | Pseudoalteromonas | Pseudoalteromonas sp. SR44-8 | SAMN15642185 | aquatic | 7 | Yes | 3 | Peptidase_S66;Phage_integrase;AA | — |
| 2778430.SAMN16816195.JADPIE0100( | **typeI-B2** | 241 | high | Bacillota | Clostridia | Halanaerobiales | Halarsenatibacteraceae | Halonatronomonas | Halonatronomonas betaini | SAMN16816195 | aquatic | 11 | Yes | 4 | PglD_N;AAA_15;TolA_bind_tri;DEA | — |
| 165186.SAMEA6149038.CACWTQ010( | **typeI-B2** | 239,9 | high | Bacillota | Clostridia | Eubacteriales | Oscillospiraceae | Ruminococcus | uncultured Ruminococcus sp. | SAMEA6149038 | host-associated | 11 | Yes | 4 | YhcG_C;AAA_15;Eco57I;ResIII | — |
| 1306154.SAMN17155236.JAEMBY010( | **typeI-B2** | 236,2 | high | Bacillota | Bacilli | Bacillales | Caryophanaceae | Rummeliibacillus | Rummeliibacillus suwonensis | SAMN17155236 | anthropogenic | 11 | Yes | 4 | Phage_integrase;AAA_15;NLRC4_ | — |
| 56779.SAMN05421834.FTNC01000003 | **typeI-B2** | 234,3 | high | Bacillota | Clostridia | Halanaerobiales | Halanaerobiaceae | Halanaerobium | Halanaerobium kushneri | SAMN05421834 | terrestrial | 11 | Yes | 4 | DivIC;AAA_15;AAA | — |
| 853.SAMN07350508.NMTY01000012_3 | **typeI-B2** | 232,8 | high | Bacillota | Clostridia | Eubacteriales | Oscillospiraceae | Faecalibacterium | Faecalibacterium prausnitzii | SAMN07350508 | host-associated | 8 | Yes | 4 | RepA_N;AAA_21;DALR_1;DUF193 | — |
| 2841523.SAMN19374015.JAHLPV0100 | **typeI-B2** | 231,7 | high | Bacillota | Clostridia | Acetivibrionales | Acetivibrionaceae | Acetivibrio | Acetivibrio sp. MSJd-27 | SAMN19374015 | host-associated | 11 | Yes | 4 | HTH_17;AAA_21;RE_LlaJI | — |
| 2044939.SAMEA5278378.CAAGAI010( | **typeI-B2** | 231,2 | high | Bacillota | Clostridia | — | — | — | Clostridia bacterium | SAMEA5278378 | host-associated | 11 | Yes | 4 | VWA_4;DUF6809;AAA_21;RRM_D | — |
| 214851.SAMN10183615.SFKI01000027 | **typeI-B2** | 228,3 | high | Bacillota | Clostridia | Eubacteriales | Oscillospiraceae | Subdoligranulum | Subdoligranulum variabile | SAMN10183615 | host-associated | 6 | Yes | 2 | ALP_N | — |
| 1798168.SAMN04487829.FOZB010000 | **typeI-B2** | 222,2 | high | Bacillota | Clostridia | Lachnospirales | Lachnospiraceae | Pseudobutyrivibrio | Pseudobutyrivibrio sp. NOR37 | SAMN04487829 | host-associated | 11 | Yes | 4 | ORF6C;AAA_15;PSS;WD40_RLD | — |
| 1898203.SAMEA8801389.CAJTEV010( | **typeI-B2** | 215,1 | borderline | Bacillota | Clostridia | Lachnospirales | Lachnospiraceae | — | Lachnospiraceae bacterium | SAMEA8801389 | host-associated | 10 | Yes | 4 | RmlD_sub_bind;DUF6602;AAA_21 | — |
| 610130.SAMEA7202544.CAJFOY0100( | **typeI-B2** | 214,5 | borderline | Bacillota | Clostridia | Lachnospirales | Lachnospiraceae | Lacrimispora | Lacrimispora saccharolytica | SAMEA7202544 | host-associated | 7 | Yes | 4 | ADH_Fe_C;AAA_21;DUF6602;Tsp | — |
| 588581.SAMN00002727.ACXX020000( | **typeI-B2** | 213,4 | borderline | Bacillota | Clostridia | Eubacteriales | Oscillospiraceae | Ruminiclostridium | Ruminiclostridium papyrosolvens | SAMN00002727 | host-associated | 10 | Yes | 4 | Pentapeptide_4;AAA_21;tRNA-synt | — |
| 195049.SAMEA7847077.CAJLTX01000 | **typeI-B2** | 210,9 | borderline | Bacillota | Clostridia | Eubacteriales | Clostridiaceae | — | uncultured Clostridiaceae bacterium | SAMEA7847077 | — | 10 | Yes | 0 | — | — |
| 297314.SAMEA8805400.CAJUBW0100 | **typeI-B2** | 210 | borderline | Bacillota | Clostridia | Lachnospirales | Lachnospiraceae | — | uncultured Lachnospiraceae bacterium | SAMEA8805400 | host-associated | 10 | Yes | 4 | Na_Ala_symp;DUF6602;AAA_21;R | — |
| 876091.SAMEA8805500.CAJUDW010( | **typeI-B2** | 208 | borderline | Bacillota | Clostridia | Eubacteriales | Oscillospiraceae | Oscillibacter | uncultured Oscillibacter sp. | SAMEA8805500 | host-associated | 10 | Yes | 4 | HhH-GPD;AAA_21;AbrB | — |
| 297314.SAMEA8805089.CAJTOV0100( | **typeI-B2** | 206,3 | borderline | Bacillota | Clostridia | Lachnospirales | Lachnospiraceae | — | uncultured Lachnospiraceae bacterium | SAMEA8805089 | host-associated | 6 | Yes | 3 | DUF5630;AAA_21;DUF6602 | — |
| 77133.SAMEA7852836.CAJMQQ01000 | **typeI-B2** | 203,8 | borderline | — | — | — | — | — | uncultured bacterium | SAMEA7852836 | host-associated | 7 | Yes | 4 | AAA_21;DUF6602;HTH_3 | — |
| 2823316.SAMN18603874.CP073692_1 | **typeI-B2** | 200,7 | borderline | Bacillota | Clostridia | Lachnospirales | Lachnospiraceae | Faecalicatena | Faecalicatena sp. Marseille-Q4148 | SAMN18603874 | host-associated | 10 | Yes | 4 | DUF6602;Lyase_8_N;AAA_21 | — |
| 297314.SAMEA8805250.CAJTUZ0100( | **typeI-B2** | 198,2 | borderline | Bacillota | Clostridia | Lachnospirales | Lachnospiraceae | — | uncultured Lachnospiraceae bacterium | SAMEA8805250 | host-associated | 10 | Yes | 4 | Metallophos_2;AAA_21;DUF6602;D | — |
| 1898203.SAMN10878285.SRYA010000 | **typeI-B2** | 194,1 | borderline | Bacillota | Clostridia | Lachnospirales | Lachnospiraceae | — | Lachnospiraceae bacterium | SAMN10878285 | host-associated | 10 | Yes | 4 | AAA_21;Fic;Uma2 | — |
| 297314.SAMEA8805667.CAJUQJ0100( | **typeI-B2** | 181,5 | borderline | Bacillota | Clostridia | Lachnospirales | Lachnospiraceae | — | uncultured Lachnospiraceae bacterium | SAMEA8805667 | host-associated | 7 | Yes | 4 | Big_2;AAA_21;DUF6602;YchF-GTI | — |
| 2013716.SAMN06767617.PHCH01000( | **typeI-C1** | 349,7 | high | Pseudomonadota | Betaproteobacteria | — | — | — | Betaproteobacteria bacterium HGW-Be | SAMN06767617 | terrestrial | 11 | Yes | 4 | NttA;DHOR;TOBE | — |
| 1948756.SAMN06451412.DCUV01000( | **typeI-C1** | 348 | high | Spirochaetota | Spirochaetia | — | — | — | Spirochaetia bacterium UBA2205 | SAMN06451412 | anthropogenic | 10 | Yes | 4 | PD40;Na_Pi_cotrans | — |
| 1619952.SAMN03340298.JYOD010000 | **typeI-C1** | 344,2 | high | Pseudomonadota | Betaproteobacteria | Burkholderiales | Burkholderiaceae | — | Burkholderiaceae bacterium 16 | SAMN03340298 | terrestrial | 11 | Yes | 4 | Resolvase;Abi_C;DUF6320;UvrD-h | — |
| 1951640.SAMN06454975.DKFJ010000 | **typeI-C1** | 343,8 | high | Deferribacterota | Deferribacteres | Deferribacterales | Deferribacteraceae | — | Deferribacteraceae bacterium UBA679 | SAMN06454975 | anthropogenic | 11 | Yes | 4 | WYL;DLH;DAHP_synth_1;SIS_2 | — |
| 1743159.SAMN03400177.LOJI0100004 | **typeI-C1** | 342,2 | high | Pseudomonadota | Betaproteobacteria | Burkholderiales | Burkholderiaceae | Polynucleobacter | Polynucleobacter yangtzensis | SAMN03400177 | salinity | 11 | Yes | 4 | Asp_decarbox;dGTP_diPhyd_N | — |
| 2052184.SAMN16056138.JAHDPY010( | **typeI-C1** | 341,7 | high | Thermodesulfobacteriota | Syntrophia | Syntrophales | Syntrophaceae | — | Syntrophaceae bacterium | SAMN16056138 | host-associated | 6 | Yes | 2 | PglZ;BrxL_ATPase | — |
| 1792290.SAMN17124794.JAEMUI0100 | **typeI-C1** | 340,8 | high | Pseudomonadota | Gammaproteobacteria | Oceanospirillales | Oceanospirillaceae | Marinomonas | Marinomonas spartinae | SAMN17124794 | aquatic | 11 | Yes | 4 | tRNA-synt_1;ABC_tran;KARI_N | — |
| 2081523.SAMN20253999.JAIQFH0100( | **typeI-C1** | 340,8 | high | Acidobacteriota | Terriglobia | — | — | — | Terriglobia bacterium | SAMN20253999 | aquatic | 11 | Yes | 4 | Big_3_5;Phage_integrase;HSDR_N | — |
| 1315271.SAMD00169750.BJXY010000 | **typeI-C1** | 340,7 | high | Pseudomonadota | Gammaproteobacteria | Alteromonadales | Pseudoalteromonadaceae | Pseudoalteromonas | Pseudoalteromonas tetraodonis | SAMD00169750 | aquatic | 11 | Yes | 4 | Lipid_DES;SprT-like;HTH_26;Tn7_ | — |
| 1948417.SAMN06454857.DITJ0100004 | **typeI-C1** | 339,5 | high | Pseudomonadota | Alphaproteobacteria | — | — | — | Alphaproteobacteria bacterium UBA61 | SAMN06454857 | anthropogenic | 11 | Yes | 4 | DarT;Macro;Response_reg;HTH_3 | — |
| 1977087.SAMN17574143.JAFEDJ0100 | **typeI-C1** | 336,5 | high | Pseudomonadota | — | — | — | — | Pseudomonadota bacterium | SAMN17574143 | host-associated | 7 | Yes | 3 | EAL;Acetyltransf_10 | — |
| 1898104.SAMN19298592.JAHJTN0100 | **typeI-C1** | 336,3 | high | Bacteroidota | — | — | — | — | Bacteroidota bacterium | SAMN19298592 | terrestrial | 7 | Yes | 3 | UvrD_C_2;DUF2813 | — |
| 208544.SAMEA6949913.CAIJIT010000 | **typeI-C1** | 336,2 | high | Pseudomonadota | Betaproteobacteria | Burkholderiales | — | — | uncultured Burkholderiales bacterium | SAMEA6949913 | aquatic | 2 | Yes | 1 | DDE_Tnp_ISAZ013 | — |
| 316.SAMN03352191.JYHV01000007_9 | **typeI-C1** | 336,2 | high | Pseudomonadota | Gammaproteobacteria | Pseudomonadales | Pseudomonadaceae | Stutzerimonas | Stutzerimonas stutzeri | SAMN03352191 | anthropogenic | 11 | Yes | 4 | MFS_1;Gas_vesicle;Reovirus_L2_7 | — |
| 1265490.SAMN02952889.JHVY010000 | **typeI-C1** | 336 | high | Pseudomonadota | Gammaproteobacteria | Pseudomonadales | Pseudomonadaceae | Pseudomonas | Pseudomonas sp. URMO17WK12:I8 | SAMN02952889 | anthropogenic | 11 | Yes | 4 | RES;Reovirus_L2_7th;AsnC_trans_ | — |
| 129578.SAMN06648058.NQJF0100000 | **typeI-C1** | 336 | high | Pseudomonadota | Gammaproteobacteria | Aeromonadales | Aeromonadaceae | Oceanimonas | Oceanimonas baumannii | SAMN06648058 | anthropogenic | 9 | Yes | 4 | adh_short;HTH_23 | — |
| 562.SAMN14342453.CP050214_1061 | **typeI-C1** | 331,6 | high | Pseudomonadota | Gammaproteobacteria | Enterobacterales | Enterobacteriaceae | Escherichia | Escherichia coli | SAMN14342453 | host-associated | 11 | Yes | 4 | Sigma54_activat;CTD6 | — |
| 562.SAMN14342455.CP050216_4924 | **typeI-C1** | 331,6 | high | Pseudomonadota | Gammaproteobacteria | Enterobacterales | Enterobacteriaceae | Escherichia | Escherichia coli | SAMN14342455 | — | 11 | Yes | 0 | — | — |
| 883078.SAMN02596753.KB375285_44 | **typeI-C1** | 331,1 | high | Pseudomonadota | Alphaproteobacteria | Hyphomicrobiales | Nitrobacteraceae | Afipia | Afipia broomeae | SAMN02596753 | host-associated | 11 | Yes | 4 | AAA_23;UvrD-helicase;Metallophos | — |
| 91915.SAMN20927460.CP081941_236 | **typeI-C1** | 329,9 | high | Pseudomonadota | Alphaproteobacteria | Acetobacterales | Acetobacteraceae | Asaia | Asaia bogorensis | SAMN20927460 | anthropogenic | 11 | Yes | 4 | MC2;CTD6;Resolvase | — |
| 2065379.SAMN08211554.CP025583_1 | **typeI-C1** | 329,3 | high | Pseudomonadota | Alphaproteobacteria | Rhodobacterales | Paracoccaceae | Paracoccus | Paracoccus jeotgali | SAMN08211554 | anthropogenic | 11 | Yes | 4 | PSK_trans_fac;DAO;TrbL | — |
| 562.SAMN04563602.CP017061_4253 | **typeI-C1** | 327,4 | high | Pseudomonadota | Gammaproteobacteria | Enterobacterales | Enterobacteriaceae | Escherichia | Escherichia coli | SAMN04563602 | host-associated | 11 | Yes | 4 | DinI;DUF8087;Phage_portal;Termin | — |
| 192812.SAMN18440232.JAGHQS0100( | **typeI-C1** | 326,3 | high | Pseudomonadota | Alphaproteobacteria | Sphingomonadales | Erythrobacteraceae | Qipengyuania | Qipengyuania flava | SAMN18440232 | aquatic | 11 | Yes | 4 | Peptidase_S9;ThiC_Rad_SAM;Maz | — |
| 2779529.SAMN16427092.JADCNH010( | **typeI-C1** | 325,8 | high | Pseudomonadota | Betaproteobacteria | Burkholderiales | Alcaligenaceae | Verticiella | Verticiella alkaliphila | SAMN16427092 | host-associated | 11 | Yes | 4 | N6_Mtase;RadC;Phage_integrase | — |
| 1124991.SAMN02603113.CP004345_2 | **typeI-C1** | 324,7 | high | Pseudomonadota | Gammaproteobacteria | Enterobacterales | Morganellaceae | Morganella | Morganella morganii | SAMN02603113 | host-associated | 11 | Yes | 4 | Peptidase_S24;Herpes_IE1;Phage_ | — |
| 2099680.SAMN19791121.JAIBBO0100( | **typeI-C1** | 324,7 | high | Methanobacteriota | Methanobacteria | Methanobacteriales | Methanobacteriaceae | — | Methanobacteriaceae archaeon | SAMN19791121 | anthropogenic | 2 | Yes | 1 | NES_C_h | — |
| 60549.SAMN05444165.FSRU01000001 | **typeI-C1** | 324,3 | high | Pseudomonadota | Betaproteobacteria | Burkholderiales | Burkholderiaceae | Paraburkholderia | Paraburkholderia phenazinium | SAMN05444165 | host-associated | 11 | Yes | 4 | DUF2268;Glyoxalase;TPR_5 | — |
| 1660129.SAMN03652473.MEEX01000( | **typeI-C1** | 323 | high | Pseudomonadota | Alphaproteobacteria | Caulobacterales | Caulobacteraceae | Phenylobacterium | Phenylobacterium sp. SCN 70-31 | SAMN03652473 | anthropogenic | 8 | Yes | 4 | PilM_2;Piwi;Sf6_terminase;PhyH | — |
| 1381123.SAMN02299295.AYOD01000( | **typeI-C1** | 322,4 | high | Pseudomonadota | Alphaproteobacteria | Hyphomicrobiales | Phyllobacteriaceae | Aliihoeflea | Aliihoeflea sp. 2WW | SAMN02299295 | host-associated | 11 | Yes | 4 | EamA;QueF;PALP | — |
| 1947379.SAMN06454808.DHXT010001 | **typeI-C1** | 320,9 | high | Pseudomonadota | Betaproteobacteria | Burkholderiales | Comamonadaceae | Rhodoferax | Rhodoferax sp. UBA5149 | SAMN06454808 | terrestrial | 11 | Yes | 4 | RHH_6;Phage_integrase;HK;TMP-T | — |
| 2053307.SAMN16091041.JADQBJ010( | **typeI-C1** | 320,9 | high | Thermodesulfobacteriota | Desulfobulbia | Desulfobulbales | Desulfobulbaceae | — | Desulfobulbaceae bacterium | SAMN16091041 | aquatic | 11 | Yes | 4 | Thy1;SNase;RHH_7 | — |
| 2060068.SAMN08166491.PJZH010000 | **typeI-C1** | 320,2 | high | Pseudomonadota | Gammaproteobacteria | Enterobacterales | Yersiniaceae | Chimaeribacter | Chimaeribacter coloradensis | SAMN08166491 | host-associated | 11 | Yes | 4 | Bac_export_2;DUF5076;Peripla_BP | — |

| | | | | | | | | | | | | | | | | |
|---|---|---|---|---|---|---|---|---|---|---|---|---|---|---|---|---|
| 153809.SAMEA6151226.CACZZK0100( | **typeI-C1** | 318,3 | high | Pseudomonadota | — | — | — | — | uncultured Pseudomonadota bacteriun | SAMEA6151226 | host-associated | 11 | Yes | 4 | Resolvase;CdiA_C_tRNase;Inositol | — |
| 2053615.SAMEA5279326.CAAGBC010 | **typeI-C1** | 316,6 | high | Spirochaetota | Spirochaetia | — | — | — | Spirochaetia bacterium | SAMEA5279326 | host-associated | 11 | Yes | 4 | YtrI_C;JetD_C;Macro | — |
| 29522.SAMN10696518.SAXU01000001 | **typeI-C1** | 315,1 | high | Spirochaetota | Spirochaetia | Brachyspirales | Brachyspiraceae | Brachyspira | Brachyspira aalborgi | SAMN10696518 | host-associated | 11 | Yes | 4 | DUF4160;DUF2442;Eco57I;DUF78 | — |
| 470934.SAMN03068907.CP011427_15 | **typeI-C1** | 307,7 | high | Pseudomonadota | Gammaproteobacteria | Enterobacterales | Erwiniaceae | Pantoea | Pantoea vagans | SAMN03068907 | aquatic | 11 | Yes | 4 | Terminase_6N;Phage_portal;PPR;F | — |
| 1830124.SAMN04620766.LWHH01000( | **typeI-C1** | 304,8 | high | Pseudomonadota | Alphaproteobacteria | Hyphomicrobiales | Brucellaceae | Ochrobactrum | Ochrobactrum sp. 3-3 | SAMN04620766 | terrestrial | 11 | Yes | 4 | Alpha-amy_C_pro;TM2;GSH-S_AT | — |
| 1981031.SAMEA8805184.CAJTRX010( | **typeI-C1** | 297,1 | high | Thermodesulfobacteriota | Desulfovibrionia | Desulfovibrionales | Desulfovibrionaceae | Mailhella | uncultured Mailhella sp. | SAMEA8805184 | host-associated | 11 | Yes | 4 | ADP_ribosyl_GH;DUF3801;T1RH-I | — |
| 1395944.SAMN11959953.VDMN01000( | **typeI-C1** | 296,9 | high | Pseudomonadota | Alphaproteobacteria | Hyphomicrobiales | Rhizobiaceae | Aliirhizobium | Aliirhizobium smilacinae | SAMN11959953 | anthropogenic | 11 | Yes | 4 | Resolvase;DUF7354;PAP2_3;Phag | — |
| 1940281.SAMN19405844.JAHRFE010( | **typeI-C1** | 292,7 | high | Pseudomonadota | Alphaproteobacteria | Hyphomicrobiales | Rhizobiaceae | Hoeflea | Hoeflea sp. | SAMN19405844 | terrestrial | 6 | Yes | 2 | Resolvase;DUF302 | — |
| 440079.SAMN07680236.NXBQ0100000( | **typeI-C1** | 285,9 | high | Pseudomonadota | Gammaproteobacteria | Aeromonadales | Aeromonadaceae | Aeromonas | Aeromonas bivalvium | SAMN07680236 | host-associated | 11 | Yes | 4 | CENP-U;CrtC;FtsX | — |
| 2529835.SAMN10994878.SJOH010000 | **typeI-C1** | 267,4 | high | Pseudomonadota | Gammaproteobacteria | Moraxellales | Moraxellaceae | Acinetobacter | Acinetobacter sp. ANC 3781 | SAMN10994878 | anthropogenic | 11 | Yes | 4 | HTH_3;Flk_N;UCP010056;Phage_ | — |
| 2529840.SAMN10994883.SJOC010000 | **typeI-C1** | 258,5 | high | Pseudomonadota | Gammaproteobacteria | Moraxellales | Moraxellaceae | Acinetobacter | Acinetobacter sp. ANC 4216 | SAMN10994883 | anthropogenic | 6 | Yes | 3 | Thiolase_N;3HCDH_N | — |
| 1648519.SAMEA8395269.CAJPST010( | **typeI-C1** | 253,1 | high | Pseudomonadota | Gammaproteobacteria | Pasteurellales | Pasteurellaceae | Mesocricetibacter | uncultured Mesocricetibacter sp. | SAMEA8395269 | host-associated | 6 | Yes | 2 | PsiE;Y1_Tnp | — |
| 2840708.SAMN15816948.DVFN010001 | **typeI-C2** | 355,8 | high | Bacillota | Clostridia | Eubacteriales | Oscillospiraceae | Candidatus Avoscillosp | Candidatus Avoscillospira stercorigallir | SAMN15816948 | host-associated | 6 | Yes | 2 | HSP90;Methyltransf_4 | — |
| 512298.SAMEA4891445.UQZO010000 | **typeI-C2** | 353,5 | high | Bacillota | Clostridia | Eubacteriales | Oscillospiraceae | Subdoligranulum | uncultured Subdoligranulum sp. | SAMEA4891445 | host-associated | 9 | Yes | 4 | IstB_IS21;DUF3231;Zn_Ribbon_1;I | — |
| 1586779.SAMEA4891222.UWSG01000 | **typeI-C2** | 353,2 | high | Bacillota | Clostridia | Lachnospirales | Lachnospiraceae | Lachnoclostridium | uncultured Lachnoclostridium sp. | SAMEA4891222 | — | 11 | Yes | 0 | — | — |
| 1673721.SAMEA3481237.LN869529_3 | **typeI-C2** | 352,8 | high | Bacillota | Clostridia | Eubacteriales | — | Intestinimonas | Intestinimonas massiliensis (ex Afouda | SAMEA3481237 | — | 11 | Yes | 0 | — | — |
| 1898207.SAMN13894142.JAAYPT0100 | **typeI-C2** | 351 | high | Bacillota | Clostridia | Eubacteriales | — | — | Clostridiales bacterium | SAMN13894142 | host-associated | 11 | Yes | 4 | Hydrolase;HTH_3;WYL | — |
| 1349765.SAMD00046312.BCQJ010000 | **typeI-C2** | 350,5 | high | Bacillota | Bacilli | Lactobacillales | Enterococcaceae | Enterococcus | Enterococcus canis | SAMD00046312 | host-associated | 8 | Yes | 4 | SieB;DUF6095;MamF_MmsF | — |
| 1139219.SAMN02596962.ASWK01000( | **typeI-C2** | 350,5 | high | Bacillota | Bacilli | Lactobacillales | Enterococcaceae | Enterococcus | Enterococcus dispar | SAMN02596962 | host-associated | 11 | Yes | 0 | — | — |
| 297314.SAMEA8805450.CAJUDY0100( | **typeI-C2** | 348,5 | high | Bacillota | Clostridia | Lachnospirales | Lachnospiraceae | — | uncultured Lachnospiraceae bacteriun | SAMEA8805450 | host-associated | 11 | Yes | 4 | DUF6491;DUF3795;DUF4096 | — |
| 165185.SAMEA4891893.URQO010001 | **typeI-C2** | 347,6 | high | Bacillota | Clostridia | Eubacteriales | Eubacteriaceae | Eubacterium | uncultured Eubacterium sp. | SAMEA4891893 | host-associated | 3 | Yes | 2 | Fer4_5;TfoX_C | — |
| 2292270.SAMN09736896.QUKQ01000( | **typeI-C2** | 346,7 | high | Bacillota | Clostridia | Eubacteriales | Oscillospiraceae | — | Ruminococcaceae bacterium TF06-43 | SAMN09736896 | host-associated | 11 | Yes | 4 | 4HBT;DUF3783;Phage_integrase;F | — |
| 1965575.SAMN06473646.NFKJ010000 | **typeI-C2** | 345,7 | high | Bacillota | Clostridia | Lachnospirales | Lachnospiraceae | Lachnoclostridium | Lachnoclostridium sp. An181 | SAMN06473646 | host-associated | 11 | Yes | 4 | RelB;Fic;DnaB_C;YicC-like_C | — |
| 904996.SAMEA4891869.URPY0100000( | **typeI-C2** | 342,1 | high | Bacillota | Clostridia | Eubacteriales | Eubacteriaceae | Anaerofustis | uncultured Anaerofustis sp. | SAMEA4891869 | host-associated | 11 | Yes | 4 | S-AdoMet_synt_C;EIIA-man;Dak2;I | — |
| 344338.SAMEA7847122.CAJLJV01000 | **typeI-C2** | 341,6 | high | Bacillota | — | — | — | — | uncultured Bacillota bacterium | SAMEA7847122 | host-associated | 11 | Yes | 4 | Cadherin-like;DUF4430;Glyco_hydr | — |
| 199.SAMEA5905261.CABPUP0100000 | **typeI-C2** | 336,6 | high | Campylobacterota | Epsilonproteobacteria | Campylobacterales | Campylobacteraceae | Campylobacter | Campylobacter concisus | SAMEA5905261 | host-associated | 11 | Yes | 4 | ACR_tran;CinA;tRNA-synt_1 | — |
| 1123489.SAMN02441545.KE386795_1 | **typeI-C2** | 334,4 | high | Bacillota | Negativicutes | Veillonellales | Veillonellaceae | Veillonella | Veillonella magna | SAMN02441545 | host-associated | 11 | Yes | 4 | UvrD-helicase;HTH_38;KAP_NTPa | — |
| 1898207.SAMN17800681.JAGZBI0100( | **typeI-C2** | 333,4 | high | Bacillota | Clostridia | Eubacteriales | — | — | Clostridiales bacterium | SAMN17800681 | host-associated | 11 | Yes | 4 | ABC_membrane;DUF3810;DUF11; | — |
| 707003.SAMEA7847144.CAJLUT0100( | **typeI-C2** | 331,7 | high | Bacillota | Clostridia | Eubacteriales | Oscillospiraceae | — | uncultured Oscillospiraceae bacterium | SAMEA7847144 | host-associated | 11 | Yes | 4 | PadR;DUF4131;NLPC_P60;AAA_1 | — |
| 876091.SAMEA8805797.CAJUQM0100 | **typeI-C2** | 331,6 | high | Bacillota | Clostridia | Eubacteriales | Oscillospiraceae | Oscillibacter | uncultured Oscillibacter sp. | SAMEA8805797 | host-associated | 6 | Yes | 3 | Phage_integrase;Zn_Ribbon_1;Pkir | — |
| 2044939.SAMN16342949.JAFRHU010( | **typeI-C2** | 328,8 | high | Bacillota | Clostridia | — | — | — | Clostridia bacterium | SAMN16342949 | host-associated | 11 | Yes | 4 | AAA_25;Sigma70_r4_2;Response_ | — |
| 2485925.SAMEA6152165.CADBJK010( | **typeI-C2** | 328,6 | high | Bacillota | Clostridia | Eubacteriales | Oscillospiraceae | — | Oscillospiraceae bacterium | SAMEA6152165 | host-associated | 11 | Yes | 4 | DUF1979;Arg_tRNA_synt_N;DUF8 | — |
| 1161902.SAMN00829151.AZKM010000 | **typeI-C2** | 327,6 | high | Bacillota | Clostridia | Peptostreptococcales | Anaerovoracaceae | — | [Eubacterium] nodatum | SAMN00829151 | host-associated | 9 | Yes | 4 | HTH_28;MinE;HTH_29 | — |
| 1328.SAMEA7846664.CAJLPW010000 | **typeI-C2** | 327 | high | Bacillota | Bacilli | Lactobacillales | Streptococcaceae | Streptococcus | Streptococcus anginosus | SAMEA7846664 | host-associated | 11 | Yes | 4 | RrnaAD;DUF7822;adh_short;DUF4 | — |
| 888721.SAMN03897724.CP016202_58 | **typeI-C2** | 325,2 | high | Bacillota | Clostridia | Peptostreptococcales | Anaerovoracaceae | — | [Eubacterium] minutum | SAMN03897724 | host-associated | 11 | Yes | 4 | Methylase_S;N6_Mtase;HTH_21;D | — |
| 172733.SAMEA7846885.CAJKUA0100( | **typeI-C2** | 318,7 | high | Bacillota | Clostridia | Eubacteriales | — | — | uncultured Eubacteriales bacterium | SAMEA7846885 | host-associated | 10 | Yes | 4 | AAA_12;AAA_11 | — |
| 1123304.SAMN02441723.AQYA01000( | **typeI-C2** | 316,1 | high | Bacillota | Bacilli | Lactobacillales | Streptococcaceae | Streptococcus | Streptococcus henryi | SAMN02441723 | anthropogenic | 11 | Yes | 4 | Methyltransf_5;RrnaAD;Abi_C;DUF | — |
| 1715105.SAMN04477515.KV797949_8 | **typeI-C2** | 314,8 | high | Bacillota | Bacilli | Lactobacillales | Aerococcaceae | Aerococcus | Aerococcus sp. HMSC062A02 | SAMN04477515 | host-associated | 11 | Yes | 4 | SWI2_SNF2;EF_EFCAB10_C;ZinT | — |
| 2044939.SAMEA5278349.CAAGAO010 | **typeI-C2** | 314,8 | high | Bacillota | Clostridia | — | — | — | Clostridia bacterium | SAMEA5278349 | host-associated | 11 | Yes | 4 | P-mevalo_kinase;DUF4467;Resolv | — |
| 1965587.SAMN06473663.NFJU010000 | **typeI-C2** | 313,4 | high | Bacillota | Clostridia | Eubacteriales | Acutalibacteraceae | Anaeromassilibacillus | Anaeromassilibacillus sp. An200 | SAMN06473663 | host-associated | 7 | Yes | 3 | MatE;Y1_Tnp | — |
| 1123312.SAMN02256411.KB904582_1 | **typeI-C2** | 312,4 | high | Bacillota | Bacilli | Lactobacillales | Streptococcaceae | Streptococcus | Streptococcus ovis | SAMN02256411 | host-associated | 11 | Yes | 4 | Peptidase_MA_2;RrnaAD;Abi_C;HI | — |
| 83427.SAMEA7848085.CAJMDY01000( | **typeI-C2** | 279,8 | high | Bacillota | Bacilli | Lactobacillales | Streptococcaceae | Streptococcus | uncultured Streptococcus sp. | SAMEA7848085 | host-associated | 11 | Yes | 4 | BPD_transp_1;CBM_11;tRNA-Thr_ | — |
| 331679.SAMN11653953.VBTH0100002 | **typeI-C2** | 266,2 | high | Bacillota | Bacilli | Lactobacillales | Lactobacillaceae | Pediococcus | Pediococcus stilesii | SAMN11653953 | host-associated | 6 | Yes | 2 | CoV_S2;VanZ | — |
| 1965645.SAMN06473737.NFIA0100002 | **typeI-C3** | 357,5 | high | Bacteroidota | Bacteroidia | Bacteroidales | Rikenellaceae | Alistipes | Alistipes sp. An54 | SAMN06473737 | host-associated | 11 | Yes | 4 | Metallophos;HTH_17 | — |
| 1872444.SAMN16346988.JAFXLG0100 | **typeI-C3** | 325,9 | high | Bacteroidota | Bacteroidia | Bacteroidales | Rikenellaceae | Alistipes | Alistipes sp. | SAMN16346988 | host-associated | 11 | Yes | 4 | tRNA_edit;NTP_transf_5;Peptidase | — |
| 2840757.SAMN15817056.JADILV0100( | **typeI-C3** | 318,1 | high | Bacteroidota | Bacteroidia | Bacteroidales | — | Candidatus Cryptobact | Candidatus Cryptobacteroides avicola | SAMN15817056 | host-associated | 8 | Yes | 4 | TatA_B_E;GFO_IDH_MocA;Aminot | — |
| 505249.SAMN07360937.NWVX010000 | **typeI-C3** | 308,9 | borderline | Campylobacterota | Epsilonproteobacteria | Campylobacterales | Arcobacteraceae | Malaciobacter | Malaciobacter marinus | SAMN07360937 | terrestrial | 10 | Yes | 4 | DUF808;AhpC-TSA;TRAP_alpha;M | — |
| 1802259.SAMN04314634.MICF010000( | **typeI-C3** | 305,8 | borderline | Campylobacterota | Epsilonproteobacteria | Campylobacterales | Sulfurimonadaceae | Sulfurimonas | Sulfurimonas sp. RIFOXYD12_FULL_ | SAMN04314634 | terrestrial | 7 | Yes | 4 | Putative_G5P;zf-CRD;ParE_toxin | — |
| 2030927.SAMN11294446.SUYX010000 | **typeI-C3** | 293,1 | low | Bacteroidota | Bacteroidia | Bacteroidales | — | — | Bacteroidales bacterium | SAMN11294446 | host-associated | 10 | Yes | 4 | LPD3;UDPG_MGDP_dh_N;NTP_t | — |
| 2590021.SAMN12085941.CP041166_5 | **typeI-C3** | 282,3 | low | Campylobacterota | Epsilonproteobacteria | Campylobacterales | Sulfurimonadaceae | Sulfurimonas | Sulfurimonas xiamenensis | SAMN12085941 | anthropogenic | 10 | Yes | 4 | Lipase_GDSL_2;Phage_integrase;F | — |
| 315271.SAMN14567401.CP051206_10 | **typeI-C3** | 278 | low | Cyanobacteriota | Cyanophyceae | Nostocales | Aphanizomenonaceae | Dolichospermum | Dolichospermum flos-aquae | SAMN14567401 | aquatic | 10 | Yes | 4 | DNA_pol_B;PDDEXK_7;DNA_pol3 | — |
| 2052166.SAMN11164354.DUDU01000( | **typeI-C3** | 264,3 | low | Candidatus Melainabacte | — | — | — | — | Candidatus Melainabacteria bacterium | SAMN11164354 | aquatic | 10 | Yes | 4 | MinC_C;DivIVA;ABC2_membrane; | — |
| 152509.SAMEA6946680.CAIMGI01000( | **typeI-C3** | 228,7 | low | Bacteroidota | — | — | — | — | uncultured Bacteroidota bacterium | SAMEA6946680 | aquatic | 6 | Yes | 3 | SdpI;DnaB_C | — |
| 152509.SAMEA6951679.CAIKXD01000 | **typeI-C3** | 220,9 | low | Bacteroidota | — | — | — | — | uncultured Bacteroidota bacterium | SAMEA6951679 | aquatic | 10 | Yes | 4 | DnaB_C;EppA_BapA;Zeta_toxin | — |
| 152509.SAMEA6953983.CAINDA0100( | **typeI-C3** | 218,5 | low | Bacteroidota | — | — | — | — | uncultured Bacteroidota bacterium | SAMEA6953983 | aquatic | 10 | Yes | 4 | WCX;UDG;DnaB_C | — |
| 194843.SAMEA6951804.CAIQUS0100( | **typeI-C3** | 214,5 | low | Bacteroidota | Bacteroidia | Bacteroidales | — | — | uncultured Bacteroidales bacterium | SAMEA6951804 | aquatic | 10 | Yes | 4 | Metallophos;DnaB_C;Y_phosphatas | — |
| 2026728.SAMN16425675.JADJEU0100 | **typeI-C3** | 212 | low | Bacteroidota | Flavobacteriia | Flavobacteriales | Crocinitomicaceae | — | Crocinitomicaceae bacterium | SAMN16425675 | anthropogenic | 10 | Yes | 4 | DnaB_C;Ubiquitin_RHG40_C;Ocnu | — |
| 1198309.SAMN04102813.LKEF010000 | **typeII-A1** | 291,5 | high | Pseudomonadota | Gammaproteobacteria | Pseudomonadales | Pseudomonadaceae | Pseudomonas | Pseudomonas fluorescens | SAMN04102813 | host-associated | 11 | Yes | 4 | AAA_13;STM3845 | — |
| 2789216.SAMN12641115.CP043489_8 | **typeII-A1** | 284,2 | high | Pseudomonadota | Alphaproteobacteria | Hyphomicrobiales | Xanthobacteraceae | Labrys | Labrys sp. KNU-23 | SAMN12641115 | host-associated | 11 | Yes | 4 | UvrB_inter;STM3845;Polysacc_lyas | — |
| 2013740.SAMN06767641.PHBJ010000 | **typeII-A1** | 280,2 | high | — | Deltaproteobacteria | — | — | — | Deltaproteobacteria bacterium HGW-D | SAMN06767641 | terrestrial | 7 | Yes | 3 | Transposase_20;STM3845;DUF31 | — |
| 1969471.SAMN08180044.PLPS010000 | **typeII-A1** | 274,9 | high | Acidobacteriota | Terriglobia | Terriglobales | Acidobacteriaceae | Granulicella | Granulicella sp. | SAMN08180044 | temperature | 7 | Yes | 3 | STM3845 | — |
| 42354.SAMN05216300.FNOE01000035 | **typeII-A1** | 274,4 | high | Pseudomonadota | Betaproteobacteria | Nitrosomonadales | Nitrosomonadaceae | Nitrosomonas | Nitrosomonas oligotropha | SAMN05216300 | aquatic | 11 | Yes | 4 | DUF4179;DUF3653;STM3845 | — |
| 1737357.SAMN14908366.JACHEG010( | **typeII-A1** | 272,9 | high | Pseudomonadota | Alphaproteobacteria | Hyphomicrobiales | Rhizobiaceae | Rhizobium | Rhizobium wenxiniae | SAMN14908366 | host-associated | 11 | Yes | 4 | Phage_integrase;STM3845 | — |
| 358.SAMN16786093.CP072309_21 | **typeII-A1** | 272,7 | high | Pseudomonadota | Alphaproteobacteria | Hyphomicrobiales | Rhizobiaceae | Agrobacterium | Agrobacterium tumefaciens | SAMN16786093 | anthropogenic | 11 | Yes | 4 | DUF8208;STM3845;Phage_integra | — |
| 1954207.SAMN06299364.MVDH01000( | **typeII-A1** | 271,4 | high | Pseudomonadota | Gammaproteobacteria | Cellvibrionales | Cellvibrionaceae | Cellvibrio | Cellvibrio sp. 79 | SAMN06299364 | host-associated | 11 | Yes | 4 | AAA_31;STM3845;DUF3037;HipA_ | — |
| 2026735.SAMN18119946.JAGOGO010 | **typeII-A1** | 271,4 | high | Myxococcota | Myxococcia | — | — | — | Deltaproteobacteria bacterium | SAMN18119946 | anthropogenic | 11 | Yes | 4 | PPV_E1_C;STM3845;DUF499 | — |
| 1173101.SAMN18839544.CP073754_2 | **typeII-A1** | 271,2 | high | Pseudomonadota | Gammaproteobacteria | Methylococcales | Methylococcaceae | Methylomonas | Methylomonas paludis | SAMN18839544 | aquatic | 11 | Yes | 4 | Trypco2;STM3845;Virulence_RhuM | — |
| 2569093.SAMEA6954955.CAILKY0100( | **typeII-A1** | 270,4 | high | Acidobacteriota | Terriglobia | — | — | — | uncultured Acidobacteriia bacterium | SAMEA6954955 | aquatic | 11 | Yes | 4 | AAA_15;STM3845;Pkinase | — |
| 58169.SAMN10095182.RAHG0100000 | **typeII-A1** | 268,4 | high | Pseudomonadota | Gammaproteobacteria | Enterobacterales | Yersiniaceae | Rahnella | Rahnella inusitata | SAMN10095182 | host-associated | 11 | Yes | 4 | Ank;Glyco_transf_15;STM3845;Og | — |
| 1978525.SAMN16426060.JADJSP0100 | **typeII-A1** | 267,6 | high | Pseudomonadota | Alphaproteobacteria | Sphingomonadales | — | — | Sphingomonadales bacterium | SAMN16426060 | anthropogenic | 11 | Yes | 4 | DUF4189;STM3845;Aida_N | — |
| 2735259.SAMN14884351.JABJWH010( | **typeII-A1** | 266,9 | high | Pseudomonadota | Gammaproteobacteria | Oceanospirillales | Alcanivoracaceae | Alloalcanivorax | Alloalcanivorax profundimaris | SAMN14884351 | aquatic | 11 | Yes | 4 | NERD;DUF7042;STM3845;ParD_a | — |
| 1895729.SAMN05660584.MKUP01000( | **typeII-A1** | 266,8 | high | Pseudomonadota | Betaproteobacteria | Burkholderiales | — | — | Burkholderiales bacterium 67-32 | SAMN05660584 | anthropogenic | 11 | Yes | 4 | LysR_substrate;STM3845;Sigma54 | — |
| 2571749.SAMN11479196.CP039695_1 | **typeII-A1** | 266,2 | high | Pseudomonadota | Alphaproteobacteria | Sphingomonadales | Sphingomonadaceae | Novosphingobium | Novosphingobium sp. EMRT-2 | SAMN11479196 | aquatic | 11 | Yes | 4 | NAD_binding_10;Phage_int_M;STM | — |
| 57320.SAMEA104288281.LT907975_1 | **typeII-A1** | 265 | high | Thermodesulfobacteriota | Desulfovibrionia | Desulfovibrionales | Desulfovibrionaceae | Pseudodesulfovibrio | Pseudodesulfovibrio profundus | SAMEA104288281 | aquatic | 11 | Yes | 4 | rve;STM3845;DDE_Tnp_ISL3 | — |
| 1292034.SAMN02470743.APMP01000( | **typeII-A1** | 264,2 | high | Pseudomonadota | Alphaproteobacteria | Caulobacterales | Caulobacteraceae | Caulobacter | Caulobacter vibrioides | SAMN02470743 | anthropogenic | 11 | Yes | 4 | AAA_33;STM3845;EcsC;DNA_gyr | — |
| 34002.SAMN04489859.FODE01000022 | **typeII-A1** | 262,9 | high | Pseudomonadota | Alphaproteobacteria | Rhodobacterales | Paracoccaceae | Paracoccus | Paracoccus alcaliphilus | SAMN04489859 | host-associated | 11 | Yes | 4 | UDG;STM3845;Iso_dh;VirionAssen | — |
| 1947551.SAMN06455167.DDVW01000 | **typeII-A1** | 257,6 | high | Pseudomonadota | Gammaproteobacteria | Lysobacterales | Lysobacteraceae | Stenotrophomonas | Stenotrophomonas sp. UBA2302 | SAMN06455167 | host-associated | 6 | Yes | 2 | ANT;DDE_Tnp_ISL3 | — |
| 2485107.SAMN10361094.SMBO01000( | **typeII-A1** | 257,5 | high | Pseudomonadota | Alphaproteobacteria | Sphingomonadales | Sphingomonadaceae | Novosphingobium | Novosphingobium sp. PhB57 | SAMN10361094 | terrestrial | 7 | Yes | 3 | STM3845;DUF3800;MmcB-like | — |
| 1122188.SAMN02745674.FUXP010000 | **typeII-A1** | 256,8 | high | Pseudomonadota | Gammaproteobacteria | Lysobacterales | Lysobacteraceae | Novilysobacter | Novilysobacter spongiicola | SAMN02745674 | aquatic | 11 | Yes | 4 | PIN7;STM3845;PDH_E1_M | — |

| | | | | | | | | | | | | | | | | |
|---|---|---|---|---|---|---|---|---|---|---|---|---|---|---|---|---|
| 1909294.SAMEA4707559.UBPT010000 | **typeII-A1** | 255,3 | high | Pseudomonadota | Alphaproteobacteria | Hyphomicrobiales | — | — | Hyphomicrobiales bacterium | SAMEA4707559 | anthropogenic | 6 | Yes | 2 | Aminotran_5 | — |
| 2769289.SAMN15951386.JACVAP010 | **typeII-A1** | 253,3 | high | Pseudomonadota | Gammaproteobacteria | Lysobacterales | Lysobacteraceae | Xanthomonas | Xanthomonas sp. XNM01 | SAMN15951386 | host-associated | 11 | Yes | 4 | MM_CoA_mutase;STM3845;C2c1_ | — |
| 1953413.SAMN06450843.DLMJ010000 | **typeII-A1** | 252,9 | high | Candidatus Binatota | Candidatus Binatia | Candidatus Binatales | Candidatus Binataceae | Candidatus Binatus | Candidatus Binatus soli | SAMN06450843 | salinity | 11 | Yes | 4 | DUF4140;STM3845;Methyltransf_2 | — |
| 2032568.SAMN07572155.NTJF010000 | **typeII-A1** | 252,8 | high | Pseudomonadota | Gammaproteobacteria | Lysobacterales | Lysobacteraceae | — | Lysobacteraceae bacterium NML93-0 | SAMN07572155 | anthropogenic | 11 | Yes | 4 | PDH_E1_M;GvpG | — |
| 1914330.SAMN07620071.PBYA010000 | **typeII-A1** | 252,5 | high | Pseudomonadota | Gammaproteobacteria | Salinisphaerales | Salinisphaeraceae | Salinisphaera | Salinisphaera sp. | SAMN07620071 | aquatic | 11 | Yes | 4 | Ku;STM3845;LigD_Prim-Pol;tRNA_ | — |
| 2268192.SAMN10966423.VGGK010000 | **typeII-A1** | 252,1 | high | Chlorobiota | — | — | — | — | Chlorobiota bacterium | SAMN10966423 | aquatic | 11 | Yes | 4 | BH4;M_Eco57I_C;STM3845;YfhO | — |
| 1978525.SAMN18061488.JAFKLT0100 | **typeII-A1** | 251,8 | high | Pseudomonadota | Alphaproteobacteria | Sphingomonadales | — | — | Sphingomonadales bacterium | SAMN18061488 | anthropogenic | 9 | Yes | 4 | Aminotran_4;Phage_integrase;STM | — |
| 1648404.SAMN03565637.CP011310_2 | **typeII-A1** | 247,5 | high | Pseudomonadota | Alphaproteobacteria | Sphingomonadales | Erythrobacteraceae | Aurantiacibacter | Aurantiacibacter atlanticus | SAMN03565637 | temperature | 11 | Yes | 4 | Phage_int_M;Nal1_C;HTH_Tnp_1 | — |
| 86027.SAMEA5364554.CAADGS01000 | **typeII-A1** | 244,4 | high | Pseudomonadota | Betaproteobacteria | — | — | — | uncultured beta proteobacterium | SAMEA5364554 | anthropogenic | 8 | Yes | 4 | Arm-DNA-bind_3;STM3845;Tag1_ | — |
| 157176.SAMEA5954144.CABVRX0100 | **typeII-A1** | 241,5 | high | Nitrospirota | Nitrospiria | Nitrospirales | Nitrospiraceae | Nitrospira | uncultured Nitrospira sp. | SAMEA5954144 | anthropogenic | 11 | Yes | 4 | DUF7315;STM3845;GST_N_3;PilZ | — |
| 1898112.SAMN11532965.VBWB010000 | **typeII-A1** | 239,2 | high | Pseudomonadota | Alphaproteobacteria | Rhodospirillales | Rhodospirillaceae | — | Rhodospirillaceae bacterium | SAMN11532965 | terrestrial | 8 | Yes | 4 | dnstrm_HI1420;STM3845;Phage_i | — |
| 2052142.SAMN08179164.PLIC0100000 | **typeII-A1** | 234 | high | Acidobacteriota | Terriglobia | Terriglobales | Acidobacteriaceae | — | Acidobacteriaceae bacterium | SAMN08179164 | temperature | 11 | Yes | 4 | Radical_SAM;STM3845;DEAD;Ba | — |
| 2780381.SAMN16434398.JADCLP010 | **typeII-A1** | 232,7 | high | Pseudomonadota | Alphaproteobacteria | Sphingomonadales | Erythrobacteraceae | — | Erythrobacteraceae bacterium E2-1 Ye | SAMN16434398 | host-associated | 11 | Yes | 4 | STM3845;ABC_tran;Peptidase_M3 | — |
| 1946060.SAMN06455422.DFTV010000 | **typeII-A1** | 229 | high | Pseudomonadota | Gammaproteobacteria | Chromatiales | Chromatiaceae | Arsukibacterium | Arsukibacterium sp. UBA4203 | SAMN06455422 | aquatic | 11 | Yes | 4 | DUF6500;DUF4440;STM3845;AA | — |
| 2026749.SAMN07619267.PAAX010000 | **typeII-A1** | 228,4 | high | Ignavibacteriota | — | — | — | — | Ignavibacteriota bacterium | SAMN07619267 | anthropogenic | 11 | Yes | 4 | Peptidase_M1;STM3845;PhdYeFM | — |
| 1977087.SAMN19298580.JAHJTB0100 | **typeII-A1** | 225,3 | high | Pseudomonadota | — | — | — | — | Pseudomonadota bacterium | SAMN19298580 | terrestrial | 11 | Yes | 4 | MerR_1;Competence;Virul_fac_Brk | — |
| 172733.SAMEA7846713.CAJJME0100 | **typeII-A1** | 211,1 | high | Bacillota | Clostridia | Eubacteriales | — | — | uncultured Eubacteriales bacterium | SAMEA7846713 | host-associated | 4 | Yes | 3 | NTP_transferase;Flg_new_2 | — |
| 2721131.SAMEA7202576.CAJFVX010 | **typeII-A1** | 209,4 | high | Bacillota | Clostridia | — | — | Candidatus Howiella | Candidatus Howiella intestinavium | SAMEA7202576 | host-associated | 8 | Yes | 4 | Mem_trans;HTH_3;STM3845;AAA_ | — |
| 1898203.SAMN11294595.SVEQ010000 | **typeII-A1** | 206,5 | high | Bacillota | Clostridia | Lachnospirales | Lachnospiraceae | — | Lachnospiraceae bacterium | SAMN11294595 | host-associated | 11 | Yes | 4 | PglD_N;Bac_transf;STM3845;DUF | — |
| 253830.SAMEA6954216.CAIMLR01000 | **typeII-A1** | 205,6 | high | Myxococcota | Myxococcia | Myxococcales | — | — | uncultured Myxococcales bacterium | SAMEA6954216 | aquatic | 11 | Yes | 4 | Methylase_S;Phage_capsid_4;STM | — |
| 1938870.SAMN06273570.OCMY010000 | **typeII-A1** | 200 | high | Pseudomonadota | Gammaproteobacteria | Enterobacterales | Erwiniaceae | Pantoea | Candidatus Pantoea floridensis | SAMN06273570 | host-associated | 11 | Yes | 4 | GPW_gp25;Phage_spike;STM384 | — |
| 717962.SAMEA2272068.FP929038_45 | **typeII-A1** | 198,1 | high | Bacillota | Clostridia | Lachnospirales | Lachnospiraceae | Pseudocoprococcus | Pseudocoprococcus catus | SAMEA2272068 | host-associated | 11 | Yes | 4 | ThsA_Macro;STM3845 | — |
| 2315688.SAMN09976312.QXJC010000 | **typeII-A1** | 196,5 | high | Pseudomonadota | Betaproteobacteria | Burkholderiales | Comamonadaceae | Simplicispira | Simplicispira hankyongi | SAMN09976312 | host-associated | 8 | Yes | 4 | AAA_13;TPP_enzyme_N;STM384 | — |
| 1218077.SAMD00000366.BAYC010000 | **typeII-A1** | 196,2 | high | Pseudomonadota | Betaproteobacteria | Burkholderiales | Burkholderiaceae | Paraburkholderia | Paraburkholderia fungorum | SAMD00000366 | aquatic | 11 | Yes | 4 | BPD_transp_1;STM3845;AAA_15 | — |
| 61635.SAMEA4532333.FO681348_167 | **typeII-A1** | 157,3 | low | Mycoplasmatota | Mollicutes | Acholeplasmatales | Acholeplasmataceae | Acholeplasma | Acholeplasma brassicae | SAMEA4532333 | anthropogenic | 10 | Yes | 4 | P5CR_dimer;UPF0158;STM3845; | — |
| 1173724.SAMN00990758.AAXWDZ010 | **typeII-A2** | 303,6 | high | Pseudomonadota | Gammaproteobacteria | Enterobacterales | Enterobacteriaceae | Citrobacter | Citrobacter freundii | SAMN00990758 | host-associated | 11 | Yes | 4 | Membra_charge;Psu;Ogr_Delta | — |
| 1913989.SAMN07620381.PALV010000 | **typeII-A2** | 301,5 | high | Pseudomonadota | Gammaproteobacteria | — | — | — | Gammaproteobacteria bacterium | SAMN07620381 | aquatic | 11 | Yes | 4 | Peptidase_S8;AAA;STM3845;Meth | — |
| 172194.SAMEA9694482.CAJXTU0100 | **typeII-A2** | 292,9 | high | Pseudomonadota | Gammaproteobacteria | Thiotrichales | Piscirickettsiaceae | Cycloclasticus | uncultured Cycloclasticus sp. | SAMEA9694482 | temperature | 11 | Yes | 4 | Methylase_S;STM3845 | — |
| 1768806.SAMN04299486.LSKJ010001 | **typeII-A2** | 289 | borderline | Pseudomonadota | Alphaproteobacteria | Rhodospirillales | Rhodospirillaceae | — | Rhodospirillaceae bacterium CCH5-H1 | SAMN04299486 | aquatic | 10 | Yes | 4 | 2-Hacid_dh_C;Arabinose_bd | — |
| 331696.SAMN09742692.CP031417_26 | **typeII-A2** | 258,9 | borderline | Pseudomonadota | Alphaproteobacteria | Hyphomicrobiales | Xanthobacteraceae | Pseudolabrys | Pseudolabrys taiwanensis | SAMN09742692 | terrestrial | 10 | Yes | 4 | Avidin;PMSR;DUF2442;DUF4160 | — |
| 2282150.SAMN18059875.JAFLCS0100 | **typeII-A2** | 258,1 | borderline | Pseudomonadota | Alphaproteobacteria | Caulobacterales | — | — | Caulobacterales bacterium | SAMN18059875 | anthropogenic | 10 | Yes | 4 | ABC2_membrane_4;ThiC_Rad_SA | — |
| 154336.SAMEA9694234.CAJXUT0100 | **typeII-A2** | 244,5 | low | Pseudomonadota | Gammaproteobacteria | Alteromonadales | Colwelliaceae | Colwellia | uncultured Colwellia sp. | SAMEA9694234 | temperature | 4 | Yes | 3 | Zn_ribbon_15;DUF6622;Pentapepti | — |
| 1898207.SAMN13894018.JAAYKZ0100 | **typeII-A3** | 308,6 | high | Bacillota | Clostridia | Eubacteriales | — | — | Clostridiales bacterium | SAMN13894018 | temperature | 2 | Yes | 1 | STM3845 | — |
| 720554.SAMN02261428.CP003065_17 | **typeII-A3** | 307,2 | high | Bacillota | Clostridia | Acetivibrionales | Acetivibrionaceae | Acetivibrio | Acetivibrio clariflavus | SAMN02261428 | terrestrial | 11 | Yes | 0 | — | — |
| 1125747.SAMD00036597.BAEK010000 | **typeII-A3** | 304,2 | high | Pseudomonadota | Gammaproteobacteria | Alteromonadales | Alteromonadaceae | Paraglaciecola | Paraglaciecola chathamensis | SAMD00036597 | aquatic | 6 | Yes | 2 | STM3845;Macro | — |
| 400153.SAMN12559023.VRLR0100001 | **typeII-A3** | 301,7 | high | Pseudomonadota | Gammaproteobacteria | Chromatiales | Chromatiaceae | Rheinheimera | Rheinheimera tangshanensis | SAMN12559023 | anthropogenic | 11 | Yes | 4 | Zn_ribbon_6;RVT_1;STM3845;GD | — |
| 2026887.SAMN11533002.WPIF010000 | **typeII-A3** | 300,6 | high | Nitrospirota | — | — | — | — | Nitrospirota bacterium | SAMN11533002 | terrestrial | 7 | Yes | 3 | HTH_3;HigB-like_toxin;STM3845 | — |
| 1879010.SAMEA7848008.CAJLNZ010 | **typeII-A3** | 297,7 | high | Bacillota | — | — | — | — | Bacillota bacterium | SAMEA7848008 | host-associated | 11 | Yes | 4 | DUF6594;STM3845;Transposase_ | — |
| 297314.SAMEA8805156.CAJTUB0100 | **typeII-A3** | 293,8 | high | Bacillota | Clostridia | Lachnospirales | Lachnospiraceae | — | uncultured Lachnospiraceae bacterium | SAMEA8805156 | host-associated | 11 | Yes | 4 | GyrI-like;STM3845;DUF2229;LysM | — |
| 1286170.SAMN02603584.CP004142_2 | **typeII-A3** | 293,1 | high | Pseudomonadota | Gammaproteobacteria | Enterobacterales | Enterobacteriaceae | Raoultella | Raoultella ornithinolytica | SAMN02603584 | host-associated | 11 | Yes | 4 | Resolvase;STM3845;Adeno_E3_C | — |
| 297314.SAMEA6151505.CADAKA0100 | **typeII-A3** | 290,9 | high | Bacillota | Clostridia | Lachnospirales | Lachnospiraceae | — | uncultured Lachnospiraceae bacterium | SAMEA6151505 | host-associated | 8 | Yes | 4 | BD-FAE;ABC_tran;STM3845;DUF | — |
| 1898203.SAMN16342483.JAFQOW010 | **typeII-A3** | 288,2 | high | Bacillota | Clostridia | Lachnospirales | Lachnospiraceae | — | Lachnospiraceae bacterium | SAMN16342483 | host-associated | 11 | Yes | 4 | Acetyltransf_3;Acetyltransf_1;STM3 | — |
| 1129257.SAMEA6955832.CAIYPA0100 | **typeII-A3** | 286,9 | high | Bacteroidota | Bacteroidia | — | — | — | uncultured Bacteroidia bacterium | SAMEA6955832 | temperature | 11 | Yes | 4 | PIN_3;STM3845;CBP_BcsR | — |
| 59620.SAMEA8805740.CAJUMP01000 | **typeII-A3** | 286,6 | high | Bacillota | Clostridia | Eubacteriales | Clostridiaceae | Clostridium | uncultured Clostridium sp. | SAMEA8805740 | — | 7 | Yes | 0 | — | — |
| 59620.SAMEA8805516.CAJUHD01000 | **typeII-A3** | 286,3 | high | Bacillota | Clostridia | Eubacteriales | Clostridiaceae | Clostridium | uncultured Clostridium sp. | SAMEA8805516 | host-associated | 11 | Yes | 4 | DUF8318;Asn_synthase;STM3845 | — |
| 2800330.SAMN17189439.JAENGP010 | **typeII-A3** | 285,7 | high | Pseudomonadota | Betaproteobacteria | Burkholderiales | Alcaligenaceae | Advenella | Advenella mandrilli | SAMN17189439 | anthropogenic | 11 | Yes | 4 | GmrSD_N;STM3845;Type_ISP_C; | — |
| 2024844.SAMN14914534.JABHWA010 | **typeII-A3** | 283,9 | high | Nitrospinota | Nitrospinia | Nitrospinales | Nitrospinaceae | Nitrospina | Nitrospina sp. | SAMN14914534 | aquatic | 11 | Yes | 4 | FlgM;Peptidase_M28;STM3845 | — |
| 1926307.SAMN18120172.JAGOOQ010 | **typeII-A3** | 283,4 | high | Bacillota | Negativicutes | Veillonellales | Veillonellaceae | Veillonella | Veillonella sp. | SAMN18120172 | anthropogenic | 7 | Yes | 3 | SLFN-g3_helicase;MazG-like;STM3 | — |
| 2026760.SAMN15049710.JACDNW010 | **typeII-A3** | 281,5 | high | Fidelibacterota | — | — | — | — | Fidelibacterota bacterium | SAMN15049710 | aquatic | 11 | Yes | 4 | STM3845;Tospov_NS-S_N | — |
| 1849603.SAMEA5278299.CAAFVQ010 | **typeII-A3** | 281,1 | high | Pseudomonadota | Gammaproteobacteria | Enterobacterales | Enterobacteriaceae | — | Enterobacteriaceae bacterium | SAMEA5278299 | host-associated | 11 | Yes | 4 | Phage_int_M;Phage_integrase;STM | — |
| 1797955.SAMN04314391.MGVA010000 | **typeII-A3** | 281 | high | Elusimicrobiota | — | — | — | — | Elusimicrobia bacterium RIFOXYA12_ | SAMN04314391 | terrestrial | 11 | Yes | 4 | AAA_15;STM3845;Mal_decarbox_ | — |
| 2126740.SAMN08731216.PYLO010000 | **typeII-A3** | 280,2 | high | Bacillota | Clostridia | Eubacteriales | Clostridiaceae | Clostridium | Clostridium fessum | SAMN08731216 | host-associated | 11 | Yes | 4 | OST3_OST6;STM3845;KAP_NTP | — |
| 1898203.SAMEA7848093.CAJKQB010 | **typeII-A3** | 277,7 | high | Bacillota | Clostridia | Lachnospirales | Lachnospiraceae | — | Lachnospiraceae bacterium | SAMEA7848093 | host-associated | 11 | Yes | 4 | HicB_lk_antitox;HicA_toxin;STM38 | — |
| 1121132.SAMN02745247.FRDH010000 | **typeII-A3** | 277,3 | high | Bacillota | Clostridia | Lachnospirales | Lachnospiraceae | Butyrivibrio | Butyrivibrio hungatei | SAMN02745247 | host-associated | 11 | Yes | 4 | FRG;STM3845;ATLF;Glycos_trans | — |
| 1898104.SAMN14914016.JABHCC010 | **typeII-A3** | 276,7 | high | Bacteroidota | — | — | — | — | Bacteroidota bacterium | SAMN14914016 | aquatic | 11 | Yes | 4 | HI_1054_N;STM3845;AAA_15;Ph | — |
| 2013811.SAMN06767730.PGYR010000 | **typeII-A3** | 271,7 | high | Ignavibacteriota | — | — | — | — | Ignavibacteriae bacterium HGW-Ignavi | SAMN06767730 | anthropogenic | 10 | Yes | 4 | DUF3788;STM3845 | — |
| 1898104.SAMN19298609.JAHJUE0100 | **typeII-A3** | 264,6 | high | Bacteroidota | — | — | — | — | Bacteroidota bacterium | SAMN19298609 | terrestrial | 11 | Yes | 4 | Mrr_cat;STM3845;LiaF-TM;Adaptin | — |
| 1953111.SAMN06455312.DLMI010001 | **typeII-A3** | 264 | high | Acidobacteriota | — | — | — | — | Acidobacteria bacterium UBA7540 | SAMN06455312 | salinity | 7 | Yes | 3 | NMT1;STM3845;Y1_Tnp | — |
| 1274384.SAMD00244887.BMDT010000 | **typeII-A3** | 250 | high | Bacillota | Bacilli | Lactobacillales | Enterococcaceae | Enterococcus | Enterococcus alcedinis | SAMD00244887 | host-associated | 11 | Yes | 4 | AAA_15;STM3845;PH_alsin;SIR2_ | — |
| 221953.SAMEA5851657.CABLZX0100 | **typeII-A3** | 230,5 | low | Spirochaetota | Spirochaetia | Brachyspirales | Brachyspiraceae | Brachyspira | uncultured Brachyspira sp. | SAMEA5851657 | host-associated | 10 | Yes | 4 | Radical_SAM;SF-assemblin;STM3 | — |
| 1907659.SAMEA4555082.LT635469_6 | **typeIII-A1** | 308,4 | high | Bacillota | Clostridia | Lachnospirales | Lachnospiraceae | Blautia | Blautia sp. Marseille-P3201T | SAMEA4555082 | host-associated | 11 | Yes | 4 | PRTase-CE;DUF7761;DDE_Tnp_I | — |
| 1953166.SAMN06454853.DITN010000 | **typeIII-A1** | 307,4 | high | Bacteroidota | — | — | — | — | Bacteroidetes bacterium UBA6183 | SAMN06454853 | aquatic | 11 | Yes | 4 | YjcN;HTH_45;PRTase-CE | — |
| 1946545.SAMN06453218.DFXA010000 | **typeIII-A1** | 306,4 | high | Bacteroidota | Flavobacteriia | Flavobacteriales | Flavobacteriaceae | Flavobacterium | Flavobacterium sp. UBA4120 | SAMN06453218 | terrestrial | 11 | Yes | 4 | PLDc_2;AhpC-TSA;DnaJ_C | — |
| 2293108.SAMN09734653.QTVG010000 | **typeIII-A1** | 304,8 | high | Bacillota | Clostridia | Eubacteriales | Eubacteriaceae | Eubacterium | Eubacterium sp. AF36-5BH | SAMN09734653 | host-associated | 11 | Yes | 4 | TMEM62_C;DIP2311-like_C;DUF7 | — |
| 152509.SAMEA6951052.CAIMBK0100 | **typeIII-A1** | 303,5 | high | Bacteroidota | — | — | — | — | uncultured Bacteroidota bacterium | SAMEA6951052 | aquatic | 8 | Yes | 4 | Ribosomal_L35p;Ribosomal_L20;H | — |
| 239.SAMN18120117.JAGOJA01000001 | **typeIII-A1** | 300,8 | high | Bacteroidota | Flavobacteriia | Flavobacteriales | Flavobacteriaceae | Flavobacterium | Flavobacterium sp. | SAMN18120117 | anthropogenic | 11 | Yes | 4 | Peptidase_M50;PLDc_2;YceI;DnaJ | — |
| 1898203.SAMEA8801244.CAJSZT010 | **typeIII-A1** | 300,5 | high | Bacillota | Clostridia | Lachnospirales | Lachnospiraceae | — | Lachnospiraceae bacterium | SAMEA8801244 | host-associated | 11 | Yes | 4 | HTH_3;PNP_UDP_1;DUF7761;PR | — |
| 2763671.SAMN15805278.JACRSZ010 | **typeIII-A1** | 298,1 | high | Bacillota | Clostridia | Lachnospirales | Lachnospiraceae | Jingyaoa | Jingyaoa shaoxingensis | SAMN15805278 | host-associated | 10 | Yes | 4 | PRTase-CE;DUF7761;LytR_cpsA_ | — |
| 190304.SAMN02603417.AE009951_72 | **typeIII-A1** | 298,1 | high | Fusobacteriota | Fusobacteriia | Fusobacteriales | Fusobacteriaceae | Fusobacterium | Fusobacterium nucleatum | SAMN02603417 | host-associated | 11 | Yes | 4 | PRTase-CE;DUF7761;MatE;S-met | — |
| 213322.SAMEA9694813.CAJXTY0100 | **typeIII-A1** | 297,8 | high | Bacteroidota | Flavobacteriia | Flavobacteriales | — | — | uncultured Flavobacteriales bacterium | SAMEA9694813 | temperature | 11 | Yes | 4 | NLPC_P60;Thiolase_N;HTH_45;P | — |
| 297314.SAMEA8805636.CAJUKA0100 | **typeIII-A1** | 296,6 | high | Bacillota | Clostridia | Lachnospirales | Lachnospiraceae | — | uncultured Lachnospiraceae bacterium | SAMEA8805636 | host-associated | 6 | Yes | 2 | DUF3173;Phage_int_SAM_3 | — |
| 1898204.SAMEA5849616.CABIZX0100 | **typeIII-A1** | 296,3 | high | Bacillota | Clostridia | Eubacteriales | Clostridiaceae | — | Clostridiaceae bacterium | SAMEA5849616 | host-associated | 6 | Yes | 2 | PRTase-CE;HTH_45 | — |
| 297314.SAMEA8805562.CAJUHN0100 | **typeIII-A1** | 296,3 | high | Bacillota | Clostridia | Lachnospirales | Lachnospiraceae | — | uncultured Lachnospiraceae bacterium | SAMEA8805562 | host-associated | 4 | Yes | 0 | — | — |
| 2485925.SAMN16342043.JAFQBT010 | **typeIII-A1** | 296,1 | high | Bacillota | Clostridia | Eubacteriales | Oscillospiraceae | — | Oscillospiraceae bacterium | SAMN16342043 | host-associated | 9 | Yes | 4 | PRTase-CE;wHTH-PRTase_assc; | — |
| 165185.SAMEA8805504.CAJUEE0100 | **typeIII-A1** | 294,7 | high | Bacillota | Clostridia | Eubacteriales | Eubacteriaceae | Eubacterium | uncultured Eubacterium sp. | SAMEA8805504 | host-associated | 11 | Yes | 0 | — | — |
| 1977054.SAMN07619280.PAPI010000 | **typeIII-A1** | 290,9 | high | Bacteroidota | Flavobacteriia | Flavobacteriales | Flavobacteriaceae | Leeuwenhoekiella | Leeuwenhoekiella sp. | SAMN07619280 | aquatic | 9 | Yes | 4 | AbiEii;DUF6922;HTH_45;PRTase- | — |
| 1869212.SAMN16425552.JADJAG010 | **typeIII-A1** | 290,7 | high | Bacteroidota | Chitinophagia | Chitinophagales | Chitinophagaceae | — | Chitinophagaceae bacterium | SAMN16425552 | anthropogenic | 11 | Yes | 4 | DUF2188;HTH_45;PRTase-CE | — |
| 192149.SAMN14492348.CP051244_14 | **typeIII-A1** | 289,6 | high | Bacteroidota | Flavobacteriia | Flavobacteriales | Flavobacteriaceae | Allomuricauda | Allomuricauda sp. | SAMN14492348 | aquatic | 11 | Yes | 2 | DUF2683 | — |
| 259315.SAMEA7847295.CAJMAQ0100 | **typeIII-A1** | 288,1 | high | Bacillota | Clostridia | Eubacteriales | Oscillospiraceae | Faecalibacterium | uncultured Faecalibacterium sp. | SAMEA7847295 | host-associated | 7 | Yes | 3 | Relaxase;PRTase-CE;DUF6061 | — |

| | | | | | | | | | | | | | | | | |
|---|---|---|---|---|---|---|---|---|---|---|---|---|---|---|---|---|
| 558021.SAMN11512334.VFPD0100000 | **typeIII-A1** | 284,3 | high | Bacteroidota | Flavobacteriia | Flavobacteriales | Weeksellaceae | Chryseobacterium | Chryseobacterium aquifrigidense | SAMN11512334 | host-associated | 11 | Yes | 4 | TfoX_N;RlaP;DUF7225;PRTase-C | — |
| 152509.SAMEA6947882.CAIRES01000 | **typeIII-A1** | 284,3 | high | Bacteroidota | — | — | — | — | uncultured Bacteroidota bacterium | SAMEA6947882 | aquatic | 11 | Yes | 4 | Epimerase;RuvB_N;HTH_45;PRTa | — |
| 370804.SAMEA6152641.CADCBZ0100 | **typeIII-A1** | 283,4 | high | Bacteroidota | Bacteroidia | Bacteroidales | Prevotellaceae | — | uncultured Prevotellaceae bacterium | SAMEA6152641 | host-associated | 7 | Yes | 3 | DUF4372;wHTH-PRTase_assc;PR | — |
| 2766537.SAMN15783688.CP060696_2 | **typeIII-A1** | 277 | high | Bacillota | Clostridia | Eubacteriales | Oscillospiraceae | Caproicibacterium | Caproicibacterium amylolyticum | SAMN15783688 | host-associated | 11 | Yes | 4 | PRTase-CE;NADH_dhqG_C;Beta- | — |
| 1045867.SAMN18262562.JAGGDU010 | **typeIII-A1** | 274,5 | high | Cyanobacteriota | Cyanophyceae | Oscillatoriales | Microcoleaceae | Trichodesmium | Trichodesmium erythraeum | SAMN18262562 | aquatic | 4 | Yes | 3 | PRTase-CE;DUF2683 | — |
| 2044939.SAMN16347727.JAFYNR0100 | **typeIII-A1** | 273,1 | high | Bacillota | Clostridia | — | — | — | Clostridia bacterium | SAMN16347727 | host-associated | 9 | Yes | 4 | PRTase-CE;wHTH-PRTase_assc; | — |
| 1898204.SAMN13893815.DUPM01000 | **typeIII-A1** | 272,6 | high | Bacillota | Clostridia | Eubacteriales | Clostridiaceae | — | Clostridiaceae bacterium | SAMN13893815 | host-associated | 6 | Yes | 3 | FtsX;HTH_45;Metallophos | — |
| 194843.SAMEA6148840.CACWMD010 | **typeIII-A1** | 272 | high | Bacteroidota | Bacteroidia | Bacteroidales | — | — | uncultured Bacteroidales bacterium | SAMEA6148840 | host-associated | 11 | Yes | 4 | 23S_rRNA_IVP;DegT_DnrJ_EryC1 | — |
| 1898104.SAMN16426528.JADKJP0100 | **typeIII-A1** | 268,1 | high | Bacteroidota | — | — | — | — | Bacteroidota bacterium | SAMN16426528 | anthropogenic | 11 | Yes | 4 | Transposase_mut;HTH_45;PRTase | — |
| 2292300.SAMN09736784.QULN010000 | **typeIII-A1** | 265,2 | high | Bacillota | Clostridia | Eubacteriales | Butyricicoccaceae | Butyricicoccus | Butyricicoccus sp. OM04-18BH | SAMN09736784 | host-associated | 11 | Yes | 4 | Glycos_transf_2;NTP_transferase; | — |
| 2301481.SAMEA8805439.CAJUDL010 | **typeIII-A1** | 263,5 | high | Bacteroidota | Bacteroidia | Bacteroidales | Muribaculaceae | — | uncultured Muribaculaceae bacterium | SAMEA8805439 | host-associated | 11 | Yes | 4 | Glycos_transf_1;HTH_45;PRTase- | — |
| 1898206.SAMN16342349.JAFQNN010 | **typeIII-A1** | 261,9 | high | Spirochaetota | Spirochaetia | Spirochaetales | Spirochaetaceae | — | Spirochaetaceae bacterium | SAMN16342349 | host-associated | 9 | Yes | 4 | IFN-gamma;GmrSD_N;HTH_45;PR | — |
| 2838877.SAMN19314831.CP075682_2 | **typeIII-A1** | 261,5 | high | Bacteroidota | Flavobacteriia | Flavobacteriales | Weeksellaceae | Chryseobacterium | Chryseobacterium sp. ZHDP1 | SAMN19314831 | anthropogenic | 11 | Yes | 4 | Septknot;DUF2911;HTH_45;PRTa | — |
| 1912772.SAMN13567726.JADMLB010 | **typeIII-A1** | 261,3 | high | Thermodesulfobacteriota | Desulfovibrionia | Desulfovibrionales | Desulfovibrionaceae | Halodesulfovibrio | Halodesulfovibrio sp. | SAMN13567726 | host-associated | 11 | Yes | 4 | PRTase-CE;HTH_45;EIF3E_C | — |
| 2056868.SAMN08111093.QEHP010000 | **typeIII-A1** | 260,3 | high | Bacteroidota | Flavobacteriia | Flavobacteriales | Weeksellaceae | Chryseobacterium | Chryseobacterium sp. HMWF035 | SAMN08111093 | anthropogenic | 11 | Yes | 0 | — | — |
| 331630.SAMEA6149455.CACXJP0100 | **typeIII-A1** | 244,9 | high | Bacillota | Erysipelotrichia | Erysipelotrichales | Erysipelotrichaceae | — | uncultured Erysipelotrichaceae bacteri | SAMEA6149455 | host-associated | 11 | Yes | 4 | YidD;Ribonucleas_3_3 | — |
| 2203212.SAMN09090847.QGNY010000 | **typeIII-A1** | 242,3 | high | Bacteroidota | Sphingobacteriia | Sphingobacteriales | Sphingobacteriaceae | Pedobacter | Pedobacter paludis | SAMN09090847 | host-associated | 11 | Yes | 4 | DUF4365;Aminotran_1_2 | — |
| 2053569.SAMN16344304.JAFTIA01000 | **typeIII-A1** | 241,2 | high | Lentisphaerota | Lentisphaeria | — | — | — | Lentisphaeria bacterium | SAMN16344304 | host-associated | 11 | Yes | 4 | Pput2613-deam;Phage_holin_3_6 | — |
| 172901.SAMN08775280.QEKH0100002 | **typeIII-A1** | 238,4 | high | Lentisphaerota | Lentisphaeria | Victivallales | Victivallaceae | Victivallis | Victivallis vadensis | SAMN08775280 | host-associated | 9 | Yes | 4 | SmpB;Phage_holin_3_3;FAD_oxid | — |
| 2053569.SAMN16344634.JAFTUS0100 | **typeIII-A1** | 237 | high | Lentisphaerota | Lentisphaeria | — | — | — | Lentisphaeria bacterium | SAMN16344634 | host-associated | 11 | Yes | 4 | DUF1232;Pput2613-deam;Prometh | — |
| 1673717.SAMEA3750678.LN868536_1 | **typeIII-A1** | 233,6 | high | Bacillota | Clostridia | Eubacteriales | Acutalibacteraceae | Anaeromassilibacillus | Anaeromassilibacillus senegalensis | SAMEA3750678 | host-associated | 11 | Yes | 4 | Aminoglyc_resit;Glyco_hydro_25;G | — |
| 2053569.SAMEA5278474.CAAGES010 | **typeIII-A1** | 223,1 | high | Lentisphaerota | Lentisphaeria | — | — | — | Lentisphaeria bacterium | SAMEA5278474 | host-associated | 11 | Yes | 4 | DUF6273;SQHop_cyclase_C;Histo | — |
| 1948697.SAMN06452111.DHMO01000 | **typeIII-A1** | 222,6 | high | Lentisphaerota | Lentisphaeria | — | — | — | Lentisphaeria bacterium UBA4640 | SAMN06452111 | — | 11 | Yes | 0 | — | — |
| 1965604.SAMN06473682.NFJK010000 | **typeIII-A1** | 219,9 | high | Bacillota | Clostridia | Eubacteriales | Acutalibacteraceae | Anaeromassilibacillus | Anaeromassilibacillus sp. An250 | SAMN06473682 | host-associated | 11 | Yes | 4 | FG-GAP;GH97_N;Glyco_hydro_25 | — |
| 1120746.SAMEA2602734.CCFG01000 | **typeIII-A1** | 208,6 | low | — | — | — | — | — | bacterium MS4 | SAMEA2602734 | host-associated | 10 | Yes | 4 | FGGY_N;UTRA;DeoRC | — |
| 707003.SAMEA7848199.CAJMFC0100 | **typeIII-A1** | 203 | low | Bacillota | Clostridia | Eubacteriales | Oscillospiraceae | — | uncultured Oscillospiraceae bacterium | SAMEA7848199 | host-associated | 2 | Yes | 2 | Glyco_hydro_25;GH97_N | — |
| 905011.SAMEA7202205.CAJFQP0100 | **typeIII-A1** | 197,7 | low | Bacillota | Clostridia | Eubacteriales | Oscillospiraceae | Anaerotruncus | uncultured Anaerotruncus sp. | SAMEA7202205 | host-associated | 7 | Yes | 4 | DUF3089;tRNA_U5-meth_tr;YgaP- | — |
| 77133.SAMEA6946128.CAIOCH010000 | **typeIII-A1** | 189,2 | low | — | — | — | — | — | uncultured bacterium | SAMEA6946128 | aquatic | 1 | Yes | 1 | — | — |
| 286133.SAMEA7846561.CAJJSA01000 | **typeIII-A1** | 170 | low | Pseudomonadota | Betaproteobacteria | Burkholderiales | Sutterellaceae | Sutterella | uncultured Sutterella sp. | SAMEA7846561 | host-associated | 10 | Yes | 4 | Peptidase_M20;DUF2201_N | — |
| 1932692.SAMN11294377.SUWG01000 | **typeIII-A1** | 169 | low | Lentisphaerota | — | — | — | — | Lentisphaerota bacterium | SAMN11294377 | host-associated | 10 | Yes | 4 | DUF6530;zf-C2H2_ZNF462_11;H | — |
| 153809.SAMEA6947789.CAIKRR01000 | **typeIII-A1** | 168,4 | low | Pseudomonadota | — | — | — | — | uncultured Pseudomonadota bacteriun | SAMEA6947789 | aquatic | 8 | Yes | 4 | DUF6036;MarR;N6_N4_Mtase | — |
| 194843.SAMEA6954344.CAIRHD01000 | **typeIII-A2** | 310,2 | high | Bacteroidota | Bacteroidia | Bacteroidales | — | — | uncultured Bacteroidales bacterium | SAMEA6954344 | aquatic | 10 | Yes | 4 | Fic;HemN_C;PRTase-CE | — |
| 595494.SAMN02598508.CP001616_71 | **typeIII-A2** | 309,3 | high | Pseudomonadota | Gammaproteobacteria | Aeromonadales | Aeromonadaceae | Tolumonas | Tolumonas auensis | SAMN02598508 | anthropogenic | 11 | Yes | 4 | rve;HTH_Tnp_1;DNApolII_N;PRTa | — |
| 1262921.PRJEB697.HF992628_8 | **typeIII-A2** | 308,9 | high | Bacteroidota | Bacteroidia | Bacteroidales | Prevotellaceae | Prevotella | Prevotella sp. CAG:1185 | PRJEB697 | host-associated | 11 | Yes | 4 | DNA_methylase;Phage_RepB;Hem | — |
| 1946060.SAMN06455422.DFTV010000 | **typeIII-A2** | 303,9 | high | Pseudomonadota | Gammaproteobacteria | Chromatiales | Chromatiaceae | Arsukibacterium | Arsukibacterium sp. UBA4203 | SAMN06455422 | aquatic | 8 | Yes | 4 | HTH_Tnp_1;rve;HTH_17;Dynactin_ | — |
| 1950669.SAMN06457402.DITE010000 | **typeIII-A2** | 302 | high | Bacteroidota | Bacteroidia | Bacteroidales | — | — | Bacteroidales bacterium UBA6192 | SAMN06457402 | anthropogenic | 8 | Yes | 4 | PRTase-CE;HemN_C;Zn_ribbon_1 | — |
| 983548.SAMN00713609.CP002528_28 | **typeIII-A2** | 300,5 | high | Bacteroidota | Flavobacteriia | Flavobacteriales | Flavobacteriaceae | Dokdonia | Dokdonia sp. 4H-3-7-5 | SAMN00713609 | temperature | 11 | Yes | 4 | PRTase-CE;HemN_C;DUF4369;H | — |
| 162156.SAMEA7848044.CAJJYT01000 | **typeIII-A2** | 297,7 | high | Bacteroidota | Bacteroidia | Bacteroidales | Bacteroidaceae | Bacteroides | uncultured Bacteroides sp. | SAMEA7848044 | host-associated | 11 | Yes | 4 | HemN_C;RteC;DUF1062 | — |
| 1137281.SAMN01940371.ANLA010000 | **typeIII-A2** | 292 | high | Bacteroidota | Flavobacteriia | Flavobacteriales | Flavobacteriaceae | Xanthomarina | Xanthomarina gelatinilytica | SAMN01940371 | aquatic | 6 | Yes | 2 | Mga;PRTase-CE | — |
| 159272.SAMEA8030120.CAJOCR0100 | **typeIII-A2** | 291,5 | high | Bacteroidota | Bacteroidia | Bacteroidales | Prevotellaceae | Prevotella | uncultured Prevotella sp. | SAMEA8030120 | host-associated | 1 | No | 0 | — | — |
| 2136182.SAMN08772519.CP035934_8 | **typeIII-A2** | 276,8 | high | Pseudomonadota | Gammaproteobacteria | Moraxellales | Moraxellaceae | Acinetobacter | Acinetobacter cumulans | SAMN08772519 | aquatic | 11 | Yes | 4 | PRTase-CE;tRNA-synt_1;DUF648 | — |
| 2136182.SAMN08772519.CP035936_6 | **typeIII-A2** | 276,8 | high | Pseudomonadota | Gammaproteobacteria | Moraxellales | Moraxellaceae | Acinetobacter | Acinetobacter cumulans | SAMN08772519 | aquatic | 11 | Yes | 0 | — | — |
| 2040292.SAMEA104414344.OEPV010 | **typeIII-A2** | 276,6 | high | Bacteroidota | Bacteroidia | Bacteroidales | Dysgonomonadaceae | Pseudodysgonomonas | Pseudodysgonomonas massiliensis | SAMEA104414344 | host-associated | 11 | Yes | 4 | PRTase-CE;Staph_reg_Sar_Rot;D | — |
| 1953167.SAMN06455210.DISB010000 | **typeIII-A2** | 269,9 | high | Bacteroidota | — | — | — | — | Bacteroidetes bacterium UBA6221 | SAMN06455210 | anthropogenic | 11 | Yes | 4 | PRTase-CE;HTH_45;Bac_globin | — |
| 2601176.SAMN12492949.VPDZ010000 | **typeIII-A2** | 269,3 | high | Pseudomonadota | Gammaproteobacteria | Moraxellales | Moraxellaceae | Acinetobacter | Acinetobacter sp. YH12251 | SAMN12492949 | anthropogenic | 11 | Yes | 4 | PRTase-CE;tRNA-synt_1;DUF648 | — |
| 2040292.SAMEA104414344.OEPV010 | **typeIII-A2** | 252,6 | high | Bacteroidota | Bacteroidia | Bacteroidales | Dysgonomonadaceae | Pseudodysgonomonas | Pseudodysgonomonas massiliensis | SAMEA104414344 | host-associated | 11 | Yes | 4 | ASD1_dom;DUF4011;Lyase_8 | — |
| 162156.SAMEA7848044.CAJJYT01000 | **typeIII-A2** | 252 | high | Bacteroidota | Bacteroidia | Bacteroidales | Bacteroidaceae | Bacteroides | uncultured Bacteroides sp. | SAMEA7848044 | host-associated | 11 | Yes | 4 | SusD_RagB;CarbopepD_reg_2;HT | — |
| 162156.SAMEA5279210.CAAFNQ0100 | **typeIII-A2** | 251,2 | high | Bacteroidota | Bacteroidia | Bacteroidales | Bacteroidaceae | Bacteroides | uncultured Bacteroides sp. | SAMEA5279210 | host-associated | 11 | Yes | 4 | HATPase_c_4;DUF5686;DUF4011 | — |
| 742766.SAMN02463862.GL891979_45 | **typeIII-A2** | 250,9 | high | Bacteroidota | Bacteroidia | Bacteroidales | Dysgonomonadaceae | Dysgonomonas | Dysgonomonas gade | SAMN02463862 | anthropogenic | 11 | Yes | 4 | ABC_tran;DUF4011;Sigma70_r2;A | — |
| 77133.SAMEA6951857.CAIQON010000 | **typeIII-A2** | 249,3 | high | — | — | — | — | — | uncultured bacterium | SAMEA6951857 | aquatic | 6 | Yes | 2 | NHase_alpha;HD | — |
| 1073388.SAMN02463954.JH815524_3 | **typeIII-A2** | 246,9 | high | Bacteroidota | Bacteroidia | Bacteroidales | Bacteroidaceae | Bacteroides | Bacteroides fragilis | SAMN02463954 | host-associated | 11 | Yes | 4 | DUF5686;S1;Anti-Pycsar_Apyc1 | — |
| 226186.SAMN02604314.AE015928_44 | **typeIII-A2** | 244,1 | high | Bacteroidota | Bacteroidia | Bacteroidales | Bacteroidaceae | Bacteroides | Bacteroides thetaiotaomicron | SAMN02604314 | host-associated | 11 | Yes | 4 | HEF_HK;DUF5686;MTES_1575;D | — |
| 742727.SAMN02463851.JH992944_94 | **typeIII-A2** | 242,4 | high | Bacteroidota | Bacteroidia | Bacteroidales | Bacteroidaceae | Bacteroides | Bacteroides oleiciplenus | SAMN02463851 | host-associated | 11 | Yes | 0 | — | — |
| 1236514.SAMD00010222.BAKL010000 | **typeIII-A2** | 242,2 | high | Bacteroidota | Bacteroidia | Bacteroidales | Bacteroidaceae | Bacteroides | Bacteroides stercorirosoris | SAMD00010222 | host-associated | 11 | Yes | 0 | — | — |
| 411901.SAMN00627055.AAVM020000 | **typeIII-A2** | 241,9 | high | Bacteroidota | Bacteroidia | Bacteroidales | Bacteroidaceae | Bacteroides | Bacteroides caccae | SAMN00627055 | host-associated | 11 | Yes | 4 | GSIII_N;DUF5686;DUF4011;S1 | — |
| 338188.SAMEA5849530.CABIXA01000 | **typeIII-A2** | 241,4 | high | Bacteroidota | Bacteroidia | Bacteroidales | Bacteroidaceae | Bacteroides | Bacteroides finegoldii | SAMEA5849530 | host-associated | 11 | Yes | 4 | S1;DUF4011;DUF5686;GSIII_N | — |
| 162156.SAMEA7847312.CAJLKP01000 | **typeIII-A2** | 241 | high | Bacteroidota | Bacteroidia | Bacteroidales | Bacteroidaceae | Bacteroides | uncultured Bacteroides sp. | SAMEA7847312 | — | 6 | Yes | 2 | Carb_kinase;SecDF_P1_head | — |
| 29523.SAMN19224740.DXOY0100016 | **typeIII-A2** | 240,7 | high | Bacteroidota | Bacteroidia | Bacteroidales | Bacteroidaceae | Bacteroides | Bacteroides sp. | SAMN19224740 | host-associated | 5 | Yes | 4 | PDDEXK_2;DUF5686;DUF4011;S | — |
| 295405.SAMD00061068.AP006841_98 | **typeIII-A2** | 240,3 | high | Bacteroidota | Bacteroidia | Bacteroidales | Bacteroidaceae | Bacteroides | Bacteroides fragilis | SAMD00061068 | host-associated | 11 | Yes | 4 | GSIII_N;DUF5686;S1;Anti-Pycsar_ | — |
| 1796613.SAMN04621613.CP015401_6 | **typeIII-A2** | 240,2 | high | Bacteroidota | Bacteroidia | Bacteroidales | Bacteroidaceae | Bacteroides | Bacteroides caecimuris | SAMN04621613 | host-associated | 11 | Yes | 0 | — | — |
| 471870.SAMN00000015.ABJL0200000 | **typeIII-A2** | 240 | high | Bacteroidota | Bacteroidia | Bacteroidales | Bacteroidaceae | Bacteroides | Bacteroides intestinalis | SAMN00000015 | host-associated | 11 | Yes | 4 | Pep_deformylase;SecDF_P1_head; | — |
| 702447.SAMN00007489.ADKP010000 | **typeIII-A2** | 239,5 | low | Bacteroidota | Bacteroidia | Bacteroidales | Bacteroidaceae | Bacteroides | Bacteroides xylanisolvens | SAMN00007489 | host-associated | 8 | Yes | 0 | — | — |
| 512312.SAMEA8805346.CAJUAD0100 | **typeIII-A2** | 239,4 | low | Bacteroidota | Bacteroidia | Bacteroidales | Tannerellaceae | Parabacteroides | uncultured Parabacteroides sp. | SAMEA8805346 | host-associated | 2 | Yes | 0 | — | — |
| 997874.SAMN02463917.JH724085_33 | **typeIII-A2** | 239,1 | low | Bacteroidota | Bacteroidia | Bacteroidales | Bacteroidaceae | Bacteroides | Bacteroides cellulosilyticus | SAMN02463917 | host-associated | 10 | Yes | 0 | — | — |
| 2723058.SAMN14490823.JAAXGE010 | **typeIII-A2** | 238,3 | low | Bacteroidota | Bacteroidia | Bacteroidales | Bacteroidaceae | Bacteroides | Bacteroides sp. GM023 | SAMN14490823 | host-associated | 10 | Yes | 4 | Selenoprotein_S;DUF5686;DUF73 | — |
| 1871006.SAMN15533020.JADMZD010 | **typeIII-A2** | 238,1 | low | Bacteroidota | Bacteroidia | Bacteroidales | Bacteroidaceae | Bacteroides | Bacteroides congonensis | SAMN15533020 | host-associated | 10 | Yes | 4 | LRR_5;DUF5686;DUF4011;S1 | — |
| 85831.SAMEA6831314.CAEUHO01000 | **typeIII-A2** | 237,2 | low | Bacteroidota | Bacteroidia | Bacteroidales | Bacteroidaceae | Bacteroides | Bacteroides acidifaciens | SAMEA6831314 | host-associated | 10 | Yes | 4 | S1;DUF4011;DUF5686;AAA-ATPa | — |
| 712711.SAMN05761448.MIQB0100001 | **typeIII-A2** | 217,7 | low | Bacteroidota | Bacteroidia | Bacteroidales | Tannerellaceae | Tannerella | Tannerella sp. oral taxon 808 | SAMN05761448 | host-associated | 5 | Yes | 2 | NAD_Gly3P_dh_C;tRNA-synt_2 | — |
| 1798573.SAMN04313869.MHBH01000 | **typeIII-A2** | 192,6 | low | Lentisphaerota | — | — | — | — | Lentisphaerae bacterium GWF2_49_2 | SAMN04313869 | terrestrial | 10 | Yes | 4 | Prok-E2_D;SBP_bac_8;N_methyl | — |
| 1506601.SAMEA6956220.CAIMFA010 | **typeIII-A2** | 189,2 | low | Lentisphaerota | Lentisphaeria | — | — | — | uncultured Lentisphaeria bacterium | SAMEA6956220 | aquatic | 10 | Yes | 0 | — | — |
| 1506601.SAMEA6956826.CAIPAT0100 | **typeIII-A2** | 187,3 | low | Lentisphaerota | Lentisphaeria | — | — | — | uncultured Lentisphaeria bacterium | SAMEA6956826 | aquatic | 4 | Yes | 4 | DUF6908;DUF932 | — |
| 195913.SAMN04488004.FOTF0100001 | **typeIII-A2** | 175,4 | low | Pseudomonadota | Alphaproteobacteria | Rhodobacterales | Roseobacteraceae | Loktanella | Loktanella salsilacus | SAMN04488004 | salinity | 5 | Yes | 2 | STM3845;Toprim | — |
| 2024855.SAMN13893731.JAAYEH010 | **typeIII-A2** | 170,4 | low | Planctomycetota | Planctomycetia | Pirellulales | Pirellulaceae | Rhodopirellula | Rhodopirellula sp. | SAMN13893731 | temperature | 8 | Yes | 4 | AAA_19;BD-FAE;Band_7;AP_end | — |
| 1192759.SAMN02470887.AKIB010000 | **typeIII-A3** | 320,2 | high | Pseudomonadota | Alphaproteobacteria | Sphingomonadales | Sphingobiaceae | Sphingobium | Sphingobium xenophagum | SAMN02470887 | terrestrial | 7 | Yes | 3 | PRTase-CE;SRAP;DUF7547 | — |
| 546367.SAMN08019857.DPJS0100004 | **typeIII-A3** | 319,6 | high | Pseudomonadota | Gammaproteobacteria | Enterobacterales | Hafniaceae | Hafnia | Hafnia paralvei | SAMN08019857 | anthropogenic | 11 | Yes | 4 | HTH_18;Ni_hydr_CYTB;PRTase-C | — |
| 1219052.SAMD00046752.BCYZ010000 | **typeIII-A3** | 318,1 | high | Pseudomonadota | Alphaproteobacteria | Sphingomonadales | Sphingomonadaceae | Sphingomonas | Sphingomonas pruni | SAMD00046752 | host-associated | 8 | Yes | 4 | MobA_MobL;DUF6118;PRTase-C | — |
| 1946868.SAMN06455384.DCRO01000 | **typeIII-A3** | 314,7 | high | Pseudomonadota | Gammaproteobacteria | Thiotrichales | Piscirickettsiaceae | Methylophaga | Methylophaga sp. UBA1490 | SAMN06455384 | aquatic | 11 | Yes | 4 | MOSC;PFF1_TM;Host_attach | — |
| 1125979.SAMN02470738.AHZC010001 | **typeIII-A3** | 314,6 | high | Pseudomonadota | Alphaproteobacteria | Hyphomicrobiales | Rhizobiaceae | Rhizobium | Rhizobium sp. PDO1-076 | SAMN02470738 | host-associated | 3 | Yes | 2 | Gamma_PGA_hydro;PRTase-CE | — |
| 2163011.SAMN08912484.CP028901_2 | **typeIII-A3** | 310,8 | high | Pseudomonadota | Betaproteobacteria | Burkholderiales | Alcaligenaceae | Orrella | Orrella marina | SAMN08912484 | anthropogenic | 11 | Yes | 4 | Metallophos;PRTase-CE | — |

| | | | | | | | | | | | | | | | | |
|---|---|---|---|---|---|---|---|---|---|---|---|---|---|---|---|---|
| 1909294.SAMEA4707468.UBQB01000( | typeIII-A3 | 309,5 | high | Pseudomonadota | Alphaproteobacteria | Hyphomicrobiales | — | — | Hyphomicrobiales bacterium | SAMEA4707468 | host-associated | 11 | Yes | 4 | PRTase-CE;Penicillinase_R | — |
| 1909294.SAMEA4707564.UBOO01000( | typeIII-A3 | 309,5 | high | Pseudomonadota | Alphaproteobacteria | Hyphomicrobiales | — | — | Hyphomicrobiales bacterium | SAMEA4707564 | — | 11 | Yes | 0 | — | — |
| 2723087.SAMN14515540.JACHCP010( | typeIII-A3 | 308 | high | Pseudomonadota | Gammaproteobacteria | Lysobacterales | Rhodanobacteraceae | Rhodanobacter | Rhodanobacter sp. A1T4 | SAMN14515540 | terrestrial | 11 | Yes | 4 | MM_CoA_mutase;PRTase-CE;RE_ | — |
| 216595.SAMEA2272316.AM181176_18 | typeIII-A3 | 306,6 | high | Pseudomonadota | Gammaproteobacteria | Pseudomonadales | Pseudomonadaceae | Pseudomonas | Pseudomonas fluorescens | SAMEA2272316 | host-associated | 11 | Yes | 4 | TnpB_IS66;DDE_Tnp_IS66;PRTas | — |
| 400153.SAMN12559023.VRLR0100002 | typeIII-A3 | 305,8 | high | Pseudomonadota | Gammaproteobacteria | Chromatiales | Chromatiaceae | Rheinheimera | Rheinheimera tangshanensis | SAMN12559023 | anthropogenic | 6 | Yes | 2 | PRTase-CE;UvrD-helicase | — |
| 1195246.SAMN02470206.AKKU01000( | typeIII-A3 | 304,8 | high | Pseudomonadota | Gammaproteobacteria | Alteromonadales | Alteromonadaceae | Alishewanella | Alishewanella agri | SAMN02470206 | — | 11 | Yes | 0 | — | — |
| 1417228.SAMN04498477.CP014579_7 | typeIII-A3 | 304,7 | high | Pseudomonadota | Betaproteobacteria | Burkholderiales | Burkholderiaceae | Paraburkholderia | Paraburkholderia phytofirmans | SAMN04498477 | temperature | 11 | Yes | 4 | PRTase-CE;Glyoxalase_8;Metalloth | — |
| 49181.SAMN16425591.JADJBR010000 | typeIII-A3 | 303,4 | high | Pseudomonadota | Betaproteobacteria | Rhodocyclales | Zoogloeaceae | Zoogloea | Zoogloea sp. | SAMN16425591 | anthropogenic | 11 | Yes | 4 | HTH_3;NERD;PRTase-CE;Virulen( | — |
| 1330531.SAMN02472181.KE166411_9 | typeIII-A3 | 300,1 | high | Pseudomonadota | Gammaproteobacteria | Pseudomonadales | Pseudomonadaceae | Pseudomonas | Pseudomonas plecoglossicida | SAMN02472181 | host-associated | 6 | Yes | 2 | PRTase-CE | — |
| 1400867.SAMN02641530.CP006768_2 | typeIII-A3 | 299,7 | high | Pseudomonadota | Gammaproteobacteria | Moraxellales | Moraxellaceae | Acinetobacter | Acinetobacter baumannii | SAMN02641530 | anthropogenic | 11 | Yes | 4 | PRTase-CE;DUF6741;4HBT;Hexa | — |
| 1946058.SAMN06451192.DEYP010000 | typeIII-A3 | 298,8 | high | Pseudomonadota | Gammaproteobacteria | Chromatiales | Chromatiaceae | Arsukibacterium | Arsukibacterium sp. UBA3155 | SAMN06451192 | aquatic | 11 | Yes | 4 | Phage_AlpA;PRTase-CE;WHD_Br | — |
| 314271.SAMN12291826.VNHV0100001 | typeIII-A3 | 298,2 | high | Pseudomonadota | Alphaproteobacteria | Rhodobacterales | Roseobacteraceae | Maritimibacter | Maritimibacter alkaliphilus | SAMN12291826 | aquatic | 11 | Yes | 4 | PRiA4_ORF3;PRTase-CE;AbiJ_NT | — |
| 373675.SAMN04488045.FNUZ0100000 | typeIII-A3 | 298,2 | high | Pseudomonadota | Alphaproteobacteria | Rhodobacterales | Roseobacteraceae | Thalassococcus | Thalassococcus halodurans | SAMN04488045 | aquatic | 9 | Yes | 0 | — | — |
| 1848702.SAMN07581400.PCOM01000( | typeIII-A3 | 296 | high | Pseudomonadota | Alphaproteobacteria | Hyphomicrobiales | Brucellaceae | Ochrobactrum | Ochrobactrum sp. MYb19 | SAMN07581400 | host-associated | 7 | Yes | 3 | YobI-ATPase;PRTase-CE | — |
| 304208.SAMD00018367.DF820596_6 | typeIII-A3 | 294,5 | high | Pseudomonadota | Gammaproteobacteria | Alteromonadales | Pseudoalteromonadacea | Pseudoalteromonas | Pseudoalteromonas sp. '520P1 No. 41 | SAMD00018367 | aquatic | 11 | Yes | 4 | DUF7662;PRTase-CE;YrhC;tSH3-I | — |
| 1122616.SAMN02441143.KE383828_2 | typeIII-A3 | 294,4 | high | Pseudomonadota | Gammaproteobacteria | Oceanospirillales | Oceanospirillaceae | Oceanospirillum | Oceanospirillum beijerinckii | SAMN02441143 | aquatic | 11 | Yes | 4 | HTH_3;PRTase-CE | — |
| 314276.SAMN02436083.CH672406_16 | typeIII-A3 | 293,4 | high | Pseudomonadota | Gammaproteobacteria | Alteromonadales | Idiomarinaceae | Idiomarina | Idiomarina baltica | SAMN02436083 | aquatic | 11 | Yes | 4 | Acetyltransf_7;PRTase-CE;Phage_ | — |
| 409.SAMN20165186.JAIETD010000002 | typeIII-A3 | 293 | high | Pseudomonadota | Alphaproteobacteria | Hyphomicrobiales | Methylobacteriaceae | Methylobacterium | Methylobacterium sp. | SAMN20165186 | aquatic | 6 | Yes | 4 | Phage_tube_2;PRTase-CE;Phage_ | — |
| 2502232.SAMN08712299.SCOE01000( | typeIII-A3 | 288,9 | high | Pseudomonadota | Betaproteobacteria | Burkholderiales | Comamonadaceae | — | Comamonadaceae bacterium ALPHA2 | SAMN08712299 | anthropogenic | 11 | Yes | 4 | Astacin;Peptidase_C14;PRTase-CE | — |
| 2502219.SAMN08712286.SCNR01000( | typeIII-A3 | 284,7 | high | Pseudomonadota | Alphaproteobacteria | Caulobacterales | Caulobacteraceae | Caulobacter | Caulobacter sp. 3R27C2-B | SAMN08712286 | aquatic | 9 | Yes | 4 | DUF6088;AbiEii;PRTase-CE;DUF3 | — |
| 2597224.SAMN10849879.SHMQ01000( | typeIII-A3 | 283,4 | high | — | Deltaproteobacteria | Candidatus Acidulodesulf | — | Candidatus Acidulodes | Candidatus Acidulodesulfobacterium a | SAMN10849879 | anthropogenic | 11 | Yes | 4 | BAR_4;Phage_integrase;PRTase-C | — |
| 68895.SAMD00023459.BBQM0100000 | typeIII-A3 | 283,1 | high | Pseudomonadota | Betaproteobacteria | Burkholderiales | Burkholderiaceae | Cupriavidus | Cupriavidus basilensis | SAMD00023459 | terrestrial | 11 | Yes | 4 | AimR;PRTase-CE;DUF4411 | — |
| 2033014.SAMN09639989.DTKD010000 | typeIII-A3 | 281,5 | high | Armatimonadota | — | — | — | — | Armatimonadota bacterium | SAMN09639989 | aquatic | 11 | Yes | 4 | HTH_3;PRTase-CE;Phage_portal | — |
| 1670.SAMN12638718.VTFV01000006_ | typeIII-A3 | 278,5 | high | Actinomycetota | Actinomycetes | Micrococcales | Micrococcaceae | Arthrobacter | Arthrobacter citreus | SAMN12638718 | host-associated | 11 | Yes | 4 | AIPR;Cecropin;PRTase-CE;DUF33 | — |
| 57320.SAMEA104288281.LT907975_1( | typeIII-A3 | 277,4 | high | Thermodesulfobacteriota | Desulfovibrionia | Desulfovibrionales | Desulfovibrionaceae | Pseudodesulfovibrio | Pseudodesulfovibrio profundus | SAMEA104288281 | aquatic | 11 | Yes | 4 | HTH_Tnp_1;PRTase-CE;HTH_3;N | — |
| 2052182.SAMN16426408.JADKFD010( | typeIII-A3 | 275,8 | high | Pseudomonadota | Gammaproteobacteria | Lysobacterales | Rhodanobacteraceae | — | Rhodanobacteraceae bacterium | SAMN16426408 | anthropogenic | 11 | Yes | 4 | dnstrm_HI1420;SLFN-g3_helicase; | — |
| 1552759.SAMN12660232.CP043474_2 | typeIII-A3 | 270,6 | high | Actinomycetota | Actinomycetes | Mycobacteriales | Mycobacteriaceae | Mycolicibacterium | Mycolicibacterium grossiae | SAMN12660232 | anthropogenic | 11 | Yes | 4 | HTH_5;Hydrolase;PRTase-CE;Gm | — |
| 2740298.SAMN15042369.CP054861_9 | typeIII-A3 | 270,2 | high | Pseudomonadota | Alphaproteobacteria | Hyphomicrobiales | Aurantimonadaceae | Martelella | Martelella soudanensis | SAMN15042369 | aquatic | 11 | Yes | 4 | DDE_Tnp_1;DUF4096;RovC_DNA | — |
| 2562705.SAMEA6944266.CAIQXC010( | typeIII-A3 | 267,9 | high | Verrucomicrobiota | Verrucomicrobiia | Verrucomicrobiales | Akkermansiaceae | — | Akkermansiaceae bacterium | SAMEA6944266 | aquatic | 5 | Yes | 4 | Methyltr_RsmB-F;PRTase-CE;SWI | — |
| 184870.SAMN08193735.PKHX0100001 | typeIII-A3 | 237,8 | high | Actinomycetota | Actinomycetes | Actinomycetales | Actinomycetaceae | Varibaculum | Varibaculum cambriense | SAMN08193735 | host-associated | 11 | Yes | 4 | ABC2_membrane_3;N6_Mtase;Me | — |
| 1852372.SAMEA4521270.FNWI010000 | typeIII-A3 | 236,6 | high | Actinomycetota | Actinomycetes | Actinomycetales | Actinomycetaceae | Varibaculum | Varibaculum massiliense | SAMEA4521270 | — | 11 | Yes | 0 | — | — |
| 563192.SAMN02463824.KE150238_24 | typeIII-A3 | 233,7 | high | Thermodesulfobacteriota | Desulfovibrionia | Desulfovibrionales | Desulfovibrionaceae | Bilophila | Bilophila wadsworthia | SAMN02463824 | host-associated | 11 | Yes | 4 | Methylase_S;PRTase-CE;Fic;Cupi | — |
| 1981031.SAMEA7202140.CAJFUZ010( | typeIII-A3 | 233,4 | high | Thermodesulfobacteriota | Desulfovibrionia | Desulfovibrionales | Desulfovibrionaceae | Mailhella | uncultured Mailhella sp. | SAMEA7202140 | — | 11 | Yes | 0 | — | — |
| 158897.SAMD00245121.BMKV010000( | typeIII-A3 | 229,5 | high | Actinomycetota | Actinomycetes | Micrococcales | Micrococcaceae | Pseudarthrobacter | Pseudarthrobacter scleromae | SAMD00245121 | anthropogenic | 11 | Yes | 4 | DNA_methylase;Helicase_C | — |
| 1121890.SAMN02440631.AUDO01000( | typeIII-A4 | 348,7 | high | Bacteroidota | Flavobacteriia | Flavobacteriales | Flavobacteriaceae | Flavobacterium | Flavobacterium frigidarium | SAMN02440631 | salinity | 9 | Yes | 4 | DUF4172;Lipase_3;PRTase-CE;Ec | — |
| 651561.SAMN06453202.DCFI0100000 | typeIII-A4 | 345,8 | high | Bacteroidota | Flavobacteriia | Flavobacteriales | Weeksellaceae | Chryseobacterium | Chryseobacterium arthrosphaerae | SAMN06453202 | host-associated | 11 | Yes | 4 | TraG_N;GtfB_M;PRTase-CE;Acety | — |
| 194843.SAMEA6945835.CAJAXK0100( | typeIII-A4 | 327,3 | high | Bacteroidota | Bacteroidia | Bacteroidales | — | — | uncultured Bacteroidales bacterium | SAMEA6945835 | aquatic | 11 | Yes | 4 | Ntox30;PRTase-CE;N6_Mtase;Met | — |
| 1932669.SAMEA6565344.LR824569_2 | typeIII-A4 | 327,3 | high | Bacteroidota | Flavobacteriia | Flavobacteriales | Weeksellaceae | Chryseobacterium | Chryseobacterium sp. JV274 | SAMEA6565344 | host-associated | 11 | Yes | 4 | PRTase-CE;HTH_17 | — |
| 1871037.SAMN10967439.DTTW01000( | typeIII-A4 | 324,8 | high | Bacteroidota | Flavobacteriia | Flavobacteriales | Flavobacteriaceae | — | Flavobacteriaceae bacterium | SAMN10967439 | aquatic | 11 | Yes | 4 | HTH_17;WHD_lin-66;PRTase-CE; | — |
| 2212467.SAMN11294424.SUYB010000 | typeIII-A4 | 318,6 | high | Bacteroidota | Bacteroidia | Bacteroidales | Bacteroidaceae | — | Bacteroidaceae bacterium | SAMN11294424 | host-associated | 11 | Yes | 4 | PRTase-CE;NttA;PSI_induc_2 | — |
| 194843.SAMEA6150768.CACZHT0100 | typeIII-A4 | 316,6 | high | Bacteroidota | Bacteroidia | Bacteroidales | — | — | uncultured Bacteroidales bacterium | SAMEA6150768 | host-associated | 11 | Yes | 0 | — | — |
| 194843.SAMEA6150337.CACYRK0100 | typeIII-A4 | 315,2 | high | Bacteroidota | Bacteroidia | Bacteroidales | — | — | uncultured Bacteroidales bacterium | SAMEA6150337 | host-associated | 7 | Yes | 0 | — | — |
| 1965624.SAMN06473711.NFIV010000 | typeIII-A4 | 314,5 | high | Bacteroidota | Bacteroidia | Bacteroidales | Muribaculaceae | Muribaculum | Muribaculum sp. An289 | SAMN06473711 | host-associated | 7 | Yes | 3 | Transposase_mut;PRTase-CE;Arm | — |
| 2683263.SAMN13494107.JAAALB0100 | typeIII-A4 | 307,5 | high | Bacteroidota | Flavobacteriia | Flavobacteriales | Flavobacteriaceae | Cellulophaga | Cellulophaga sp. BC115SP | SAMN13494107 | aquatic | 9 | Yes | 4 | SH3_3;PRTase-CE;S1;DUF7660 | — |
| 2249356.SAMN09487226.CP030261_2 | typeIII-A4 | 305,5 | high | Bacteroidota | Flavobacteriia | Flavobacteriales | Flavobacteriaceae | Flavobacterium | Flavobacterium fluviale | SAMN09487226 | terrestrial | 11 | Yes | 4 | MerR_1;Abhydrolase_1;PRTase-CE | — |
| 2748031.SAMN16350081.JAGBZD010( | typeIII-A4 | 304,7 | high | Bacteroidota | Bacteroidia | Bacteroidales | Paludibacteraceae | — | Paludibacteraceae bacterium | SAMN16350081 | host-associated | 7 | Yes | 3 | HTH_17;PRTase-CE | — |
| 1236517.SAMN03704035.CP012074_1 | typeIII-A4 | 300,6 | high | Bacteroidota | Bacteroidia | Bacteroidales | Prevotellaceae | Prevotella | Prevotella fusca | SAMN03704035 | — | 11 | Yes | 0 | — | — |
| 702438.SAMN02463847.JH114217_20 | typeIII-A4 | 300,1 | high | Bacteroidota | Bacteroidia | Bacteroidales | Prevotellaceae | Segatella | Segatella oulorum | SAMN02463847 | host-associated | 11 | Yes | 4 | PRTase-CE;Fijivirus_P9-2;DUF270 | — |
| 2030927.SAMN18472452.JAGHRT010( | typeIII-A4 | 299,4 | high | Bacteroidota | Bacteroidia | Bacteroidales | — | — | Bacteroidales bacterium | SAMN18472452 | host-associated | 11 | Yes | 4 | Bac_transf;PRTase-CE;Hypoth_Ym | — |
| 370804.SAMEA6151246.CADAAG0100 | typeIII-A4 | 298,1 | high | Bacteroidota | Bacteroidia | Bacteroidales | Prevotellaceae | — | uncultured Prevotellaceae bacterium | SAMEA6151246 | host-associated | 5 | Yes | 0 | — | — |
| 370804.SAMEA6152641.CADCBZ0100 | typeIII-A4 | 295,1 | high | Bacteroidota | Bacteroidia | Bacteroidales | Prevotellaceae | — | uncultured Prevotellaceae bacterium | SAMEA6152641 | host-associated | 11 | Yes | 4 | Med9;DEDD_Tnp_IS110;PRTase-( | — |
| 2030927.SAMN14407244.JAAVFK0100 | typeIII-A4 | 284,4 | high | Bacteroidota | Bacteroidia | Bacteroidales | — | — | Bacteroidales bacterium | SAMN14407244 | host-associated | 8 | Yes | 4 | AbiEii;Metallophos;PRTase-CE | — |
| 194843.SAMEA6954344.CAIRHD0100( | typeIII-A4 | 282,2 | high | Bacteroidota | Bacteroidia | Bacteroidales | — | — | uncultured Bacteroidales bacterium | SAMEA6954344 | aquatic | 11 | Yes | 4 | PHB_acc;N6_Mtase;PRTase-CE;M | — |
| 194843.SAMEA6149846.CACXYM0100 | typeIII-A4 | 272,9 | borderline | Bacteroidota | Bacteroidia | Bacteroidales | — | — | uncultured Bacteroidales bacterium | SAMEA6149846 | host-associated | 10 | Yes | 4 | Pox_polyA_pol_N;KAP_NTPase;PI | — |
| 194843.SAMEA6152515.CADBWX010( | typeIII-A4 | 272,4 | borderline | Bacteroidota | Bacteroidia | Bacteroidales | — | — | uncultured Bacteroidales bacterium | SAMEA6152515 | host-associated | 7 | Yes | 4 | PRTase-CE;KAP_NTPase | — |
| 2293831.SAMN09736732.QUMM01000 | typeIII-A5 | 298,8 | high | Bacillota | Clostridia | Lachnospirales | Lachnospiraceae | — | Lachnospiraceae bacterium OF09-6 | SAMN09736732 | host-associated | 7 | Yes | 3 | PRTase-CE;HTH_45 | — |
| 1681184.SAMN04123205.KQ465162_8 | typeIII-A5 | 294 | high | Bacillota | Bacilli | Bacillales | Bacillaceae | Lysinibacillus | Lysinibacillus sp. ZYM-1 | SAMN04123205 | terrestrial | 11 | Yes | 4 | YhjD;PRTase-CE;IMS_C;YolD | — |
| 1834198.SAMN04621621.CP015400_2 | typeIII-A5 | 289,3 | high | Bacillota | Clostridia | Eubacteriales | Oscillospiraceae | — | Hungateiclostridiaceae bacterium KB1 | SAMN04621621 | host-associated | 11 | Yes | 4 | DUF6537;GP57;DnaD_N;PRTase- | — |
| 796937.SAMN00828788.ALNK0100002 | typeIII-A5 | 288,9 | high | Bacillota | Clostridia | Peptostreptococcales | Filifactoraceae | Peptoanaerobacter | Peptoanaerobacter stomatis | SAMN00828788 | host-associated | 11 | Yes | 4 | PRTase-CE;wHTH-PRTase_assc; | — |
| 1898203.SAMEA5278693.CAAFYN010( | typeIII-A5 | 283,4 | high | Bacillota | Clostridia | Lachnospirales | Lachnospiraceae | — | Lachnospiraceae bacterium | SAMEA5278693 | host-associated | 11 | Yes | 4 | PRTase-CE;wHTH-PRTase_assc | — |
| 1898207.SAMN19225207.DXWH01000 | typeIII-A5 | 283 | high | Bacillota | Clostridia | Eubacteriales | — | — | Clostridiales bacterium | SAMN19225207 | host-associated | 10 | Yes | 4 | DUF6809;PRTase-CE | — |
| 1280.SAMN05853508.CP017807_2376 | typeIII-A5 | 282 | high | Bacillota | Bacilli | Bacillales | Staphylococcaceae | Staphylococcus | Staphylococcus aureus | SAMN05853508 | host-associated | 11 | Yes | 4 | TPP_enzyme_C;PRTase-CE;HNH | — |
| 2028282.SAMN18359877.JAGHIX0100 | typeIII-A5 | 281,3 | high | Bacillota | Clostridia | Lachnospirales | Lachnospiraceae | Lachnoclostridium | Lachnoclostridium sp. | SAMN18359877 | host-associated | 4 | Yes | 2 | wHTH-PRTase_assc;PRTase-CE | — |
| 297314.SAMEA8805636.CAJUKA0100( | typeIII-A5 | 279,5 | high | Bacillota | Clostridia | Lachnospirales | Lachnospiraceae | — | uncultured Lachnospiraceae bacterium | SAMEA8805636 | host-associated | 10 | Yes | 4 | DUF5481;wHTH-PRTase_assc;PR | — |
| 286138.SAMEA8805337.CAJUBG0100( | typeIII-A5 | 279,4 | high | Bacillota | Clostridia | Lachnospirales | Lachnospiraceae | Dorea | uncultured Dorea sp. | SAMEA8805337 | — | 10 | Yes | 0 | — | — |
| 2044939.SAMN16345800.JAFVRR010( | typeIII-A5 | 275,3 | high | Bacillota | Clostridia | — | — | — | Clostridia bacterium | SAMN16345800 | host-associated | 11 | Yes | 4 | HNH;SLFN-g3_helicase;PRTase-C | — |
| 1946771.SAMN06451782.DDLI010000 | typeIII-A5 | 274,8 | high | Bacillota | Bacilli | Bacillales | Bacillaceae | Lysinibacillus | Lysinibacillus sp. UBA2576 | SAMN06451782 | anthropogenic | 11 | Yes | 4 | Flavin_Reduct;MerR_1;wHTH-PRT | — |
| 1948087.SAMN06454066.DIWF010000 | typeIII-A5 | 272,9 | high | Bacillota | — | — | — | — | Firmicutes bacterium UBA6113 | SAMN06454066 | terrestrial | 6 | Yes | 2 | wHTH-PRTase_assc;PRTase-CE | — |
| 1898207.SAMN19225191.DYCF010000 | typeIII-A5 | 271,9 | high | Bacillota | Clostridia | Eubacteriales | — | — | Clostridiales bacterium | SAMN19225191 | host-associated | 11 | Yes | 4 | PNP_UDP_1;RVT_1;wHTH-PRTas | — |
| 1950840.SAMN06452039.DDQW01000 | typeIII-A5 | 271,8 | high | Bacillota | Clostridia | Eubacteriales | — | — | Clostridiales bacterium UBA2432 | SAMN06452039 | terrestrial | 8 | Yes | 4 | CC4_CEP85;HTH_3;PRTase-CE;I | — |
| 642492.SAMN00013959.CP002582_28 | typeIII-A5 | 271,6 | high | Bacillota | Clostridia | Lachnospirales | Cellulosilyticaceae | Cellulosilyticum | Cellulosilyticum lentocellum | SAMN00013959 | host-associated | 11 | Yes | 4 | wHTH-PRTase_assc;PRTase-CE | — |
| 1898207.SAMN14407303.JAAVHR010( | typeIII-A5 | 267,8 | high | Bacillota | Clostridia | Eubacteriales | — | — | Clostridiales bacterium | SAMN14407303 | host-associated | 11 | Yes | 4 | PRTase-CE;HTH_28;MCPsignal | — |
| 1915574.SAMN06027062.DAKJ010000 | typeIII-A5 | 267,8 | high | Methanobacteriota | Methanomicrobia | Methanosarcinales | Methanosarcinaceae | Methanosarcina | Methanosarcina sp. UBA289 | SAMN06027062 | anthropogenic | 11 | Yes | 4 | YqaJ;Gly_zipper;RsgA_GTPase | — |
| 1410670.SAMN02744016.JHXF010000 | typeIII-A5 | 263,4 | high | Bacillota | Clostridia | Eubacteriales | Oscillospiraceae | Ruminococcus | Ruminococcus flavefaciens | SAMN02744016 | host-associated | 8 | Yes | 4 | HTH_Tnp_1;SH3_3;PRTase-CE | — |
| 2250255.SAMEA8236582.CAJPFJ0100 | typeIII-A5 | 261,2 | high | Methanobacteriota | Methanomicrobia | Methanosarcinales | — | — | Methanosarcinales archaeon | SAMEA8236582 | terrestrial | 11 | Yes | 4 | GIIM;tRNA-Thr_ED | — |
| 297314.SAMEA8805271.CAJTZG0100( | typeIII-A5 | 261 | high | Bacillota | Clostridia | Lachnospirales | Lachnospiraceae | — | uncultured Lachnospiraceae bacterium | SAMEA8805271 | host-associated | 11 | Yes | 4 | NikA-like;HD_5;wHTH-PRTase_as | — |
| 1751881.SAMEA8805505.CAJUEJ010( | typeIII-A5 | 259,7 | high | Bacillota | Clostridia | Lachnospirales | Lachnospiraceae | Hungatella | uncultured Hungatella sp. | SAMEA8805505 | host-associated | 11 | Yes | 4 | His_kinase;HTH_34;PRTase-CE | — |
| 2800128.SAMN17181733.CP067364_2 | typeIII-A5 | 258,8 | high | Bacillota | Bacilli | Lactobacillales | Carnobacteriaceae | Carnobacterium | Carnobacterium sp. CS13 | SAMN17181733 | host-associated | 11 | Yes | 4 | Band_7;PRTase-CE;Upx-like | — |
| 2485925.SAMN11294765.SVLF010000 | typeIII-A5 | 252 | high | Bacillota | Clostridia | Eubacteriales | Oscillospiraceae | — | Oscillospiraceae bacterium | SAMN11294765 | host-associated | 10 | Yes | 4 | SBP_bac_1;Agd3_C;PRTase-CE | — |

| | | | | | | | | | | | | | | | | |
|---|---|---|---|---|---|---|---|---|---|---|---|---|---|---|---|---|
| 297314.SAMEA8805790.CAJUQB0100 | **typeIII-A5** | 251,6 | high | Bacillota | Clostridia | Lachnospirales | Lachnospiraceae | — | uncultured Lachnospiraceae bacterium | SAMEA8805790 | host-associated | 10 | Yes | 4 | Penicillinase_R;DUF4180;HTH_34; | — |
| 2293093.SAMN09736788.QULM01000 | **typeIII-A5** | 247,8 | high | Bacillota | Clostridia | Lachnospirales | Lachnospiraceae | Coprococcus | Coprococcus sp. OM04-5BH | SAMN09736788 | host-associated | 11 | Yes | 4 | Peptidase_S41;GMP_synt_C;PRTa | — |
| 286138.SAMEA8805309.CAJUAM0100 | **typeIII-A5** | 243,9 | high | Bacillota | Clostridia | Lachnospirales | Lachnospiraceae | Dorea | uncultured Dorea sp. | SAMEA8805309 | host-associated | 11 | Yes | 4 | HxlR;tRNA_edit;PRTase-CE;ROK | — |
| 169435.SAMN15533212.JADNIE01000 | **typeIII-A5** | 243,6 | high | Bacillota | Clostridia | Eubacteriales | Oscillospiraceae | Anaerotruncus | Anaerotruncus colihominis | SAMN15533212 | host-associated | 9 | Yes | 4 | PRTase-CE;HTH_AsnC-type;CstA; | — |
| 1903720.SAMN18359853.JAGHHZ010 | **typeIII-A5** | 242 | high | Bacillota | Bacilli | — | — | — | Bacilli bacterium | SAMN18359853 | host-associated | 9 | Yes | 4 | ZW10_C2;LysM;PRTase-CE;Resol | — |
| 224209.SAMEA8804963.CAJTLN0100 | **typeIII-A5** | 241,7 | high | Bacillota | Bacilli | — | — | — | uncultured Bacilli bacterium | SAMEA8804963 | host-associated | 11 | Yes | 4 | RE_LlaJI;PRTase-CE;Acetyltransf_ | — |
| 2026735.SAMN17179375.JAFDGY010 | **typeIII-A5** | 241 | high | Myxococcota | Myxococcia | — | — | — | Deltaproteobacteria bacterium | SAMN17179375 | temperature | 6 | Yes | 2 | PNP_UDP_1;TPR_6 | — |
| 2049433.SAMN10607086.SKZT010000 | **typeIII-A5** | 236,4 | high | Thermodesulfobacteriota | Desulfobacteria | Desulfobacterales | Desulfobacteraceae | — | Desulfobacteraceae bacterium | SAMN10607086 | salinity | 9 | Yes | 4 | TPR_6;PNP_UDP_1;DUF983;DUF | — |
| 2026735.SAMN18120041.JAGOOG010 | **typeIII-A5** | 233,9 | low | Myxococcota | Myxococcia | — | — | — | Deltaproteobacteria bacterium | SAMN18120041 | anthropogenic | 10 | Yes | 4 | DUF1848;N6_N4_Mtase;Peptidase | — |
| 2137878.SAMN16056095.JAHDOI0100 | **typeIII-A5** | 232,8 | low | Bacillota | Negativicutes | Selenomonadales | — | — | Selenomonadales bacterium | SAMN16056095 | anthropogenic | 10 | Yes | 4 | HTH_17;HTH_Tnp_1_2;RH3_dom; | — |
| 156588.SAMEA6955620.CAIMWS0100 | **typeIII-A5** | 227,8 | low | Verrucomicrobiota | — | — | — | — | uncultured Verrucomicrobiota bacteriu | SAMEA6955620 | ph | 7 | Yes | 4 | HEAT_2;N6_N4_Mtase;DUF1848 | — |
| 282683.SAMN04488105.FNAV0100001 | **typeIII-A5** | 227,4 | low | Pseudomonadota | Alphaproteobacteria | Rhodobacterales | Roseobacteraceae | Salipiger | Salipiger thiooxidans | SAMN04488105 | aquatic | 10 | Yes | 4 | Mmel_Mtase;Mrr_N;Sigma70_r4;Pe | — |
| 1895720.SAMN05660546.MKTD01000 | **typeIII-A5** | 226 | low | Bacteroidota | Bacteroidia | — | — | — | Bacteroidia bacterium 43-41 | SAMN05660546 | anthropogenic | 10 | Yes | 4 | GMT-wHTH;Phage_integrase;PRTa | — |
| 1798573.SAMN04313869.MHBH01000 | **typeIII-A5** | 222,4 | low | Lentisphaerota | — | — | — | — | Lentisphaerae bacterium GWF2_49_2 | SAMN04313869 | terrestrial | 10 | Yes | 4 | DUF983;ResIII;TPR_19;CBM_4_9_ | — |
| 1798574.SAMN04313866.MHBI010000 | **typeIII-A5** | 221,2 | low | Lentisphaerota | — | — | — | — | Lentisphaerae bacterium GWF2_50_9 | SAMN04313866 | terrestrial | 10 | Yes | 4 | BSH_RND;ACR_tran;ResIII;PUMA | — |
| 2044936.SAMEA8395272.CAJPTC010 | **typeIII-A5** | 219,7 | low | Bacteroidota | Bacteroidia | — | — | — | Bacteroidia bacterium | SAMEA8395272 | host-associated | 10 | Yes | 4 | PatX;PRTase-CE;Phage_int_SAM_ | — |
| 194843.SAMEA104666353.ONFO0100 | **typeIII-A5** | 218,8 | low | Bacteroidota | Bacteroidia | Bacteroidales | — | — | uncultured Bacteroidales bacterium | SAMEA104666353 | host-associated | 10 | Yes | 4 | AAA_19;OLD-like_TOPRIM;PRTas | — |
| 2024855.SAMN07619947.NZVD01000 | **typeIII-A5** | 218,5 | low | Planctomycetota | Planctomycetia | Pirellulales | Pirellulaceae | Rhodopirellula | Rhodopirellula sp. | SAMN07619947 | aquatic | 10 | Yes | 4 | SGL;Lum_binding;DUF6009 | — |
| 1349822.SAMN05444348.RBXN01000 | **typeIII-A5** | 217,6 | low | Bacteroidota | Bacteroidia | Bacteroidales | Barnesiellaceae | Coprobacter | Coprobacter fastidiosus | SAMN05444348 | host-associated | 10 | Yes | 4 | HTH_17;Phage_int_SAM_5;PRTas | — |
| 59823.SAMN11294431.SUYI01000044_ | **typeIII-A5** | 216,8 | low | Bacteroidota | Bacteroidia | Bacteroidales | Prevotellaceae | Prevotella | Prevotella sp. | SAMN11294431 | host-associated | 10 | Yes | 4 | UvrD-helicase;AAA_15;PRTase-CE | — |
| 1129257.SAMEA6152838.CADCJI0100 | **typeIII-A5** | 216,7 | low | Bacteroidota | Bacteroidia | — | — | — | uncultured Bacteroidia bacterium | SAMEA6152838 | — | 10 | Yes | 0 | — | — |
| 2026779.SAMN07619720.PAPJ010000 | **typeIII-A5** | 214,3 | low | Planctomycetota | Planctomycetia | Planctomycetales | Planctomycetaceae | — | Planctomycetaceae bacterium | SAMN07619720 | aquatic | 10 | Yes | 4 | SGL;Lum_binding;DUF6009 | — |
| 2005381.SAMN14533040.JABACA010 | **typeIII-A5** | 213,9 | low | Planctomycetota | Planctomycetia | Pirellulales | Pirellulaceae | Mariniblastus | Mariniblastus sp. | SAMN14533040 | aquatic | 6 | Yes | 3 | PS_pyruv_trans;DUF5413 | — |
| 1952733.SAMN06454406.DEAW01000 | **typeIII-A5** | 213,2 | low | Planctomycetota | Planctomycetia | Planctomycetales | Planctomycetaceae | — | Planctomycetaceae bacterium UBA29 | SAMN06454406 | aquatic | 5 | Yes | 2 | SGL | — |
| 2026780.SAMN10966917.VGZI010001 | **typeIII-A5** | 208 | low | Planctomycetota | — | — | — | — | Planctomycetota bacterium | SAMN10966917 | aquatic | 6 | Yes | 3 | PAPS_reduct;AhpC-TSA;PDDEXK | — |
| 2026779.SAMN14914450.JABHSU010 | **typeIII-A5** | 206,3 | low | Planctomycetota | Planctomycetia | Planctomycetales | Planctomycetaceae | — | Planctomycetaceae bacterium | SAMN14914450 | aquatic | 1 | Yes | 1 | Lum_binding | — |
| 1798574.SAMN04313866.MHBI010000 | **typeIII-A5** | 196,2 | low | Lentisphaerota | — | — | — | — | Lentisphaerae bacterium GWF2_50_9 | SAMN04313866 | terrestrial | 10 | Yes | 4 | Prok-E2_D;DUF4942 | — |
| 2053306.SAMN18059852.JAFLBV0100 | **typeIII-A5** | 195,3 | low | Ignavibacteriota | Ignavibacteria | — | — | — | Ignavibacteria bacterium | SAMN18059852 | anthropogenic | 10 | Yes | 4 | Aminotran_5;MS_channel_2nd;Uvrl | — |
| 2052166.SAMN15436385.JACRFZ010 | **typeIII-A5** | 194,8 | low | Candidatus Melainabacte | — | — | — | — | Candidatus Melainabacteria bacterium | SAMN15436385 | anthropogenic | 10 | Yes | 4 | HigB_toxin;SMI1_KNR4 | — |
| 278095.SAMEA6946262.CAILFO01000 | **typeIII-A5** | 193,7 | low | Lentisphaerota | — | — | — | — | uncultured Lentisphaerota bacterium | SAMEA6946262 | aquatic | 1 | Yes | 0 | — | — |
| 278095.SAMEA6946262.CAILFO01000 | **typeIII-A5** | 193,4 | low | Lentisphaerota | — | — | — | — | uncultured Lentisphaerota bacterium | SAMEA6946262 | aquatic | 3 | Yes | 3 | Cation_efflux | — |
| 77133.SAMEA9695134.CAJXZK01000 | **typeIII-A5** | 192,4 | low | — | — | — | — | — | uncultured bacterium | SAMEA9695134 | anthropogenic | 5 | Yes | 2 | ATLF;Thioredoxin_2 | — |
| 2364082.SAMN18076853.JAGVJU010 | **typeIII-A5** | 191,1 | low | Candidatus Melainabacte | — | Candidatus Obscuribacte | — | — | Candidatus Obscuribacterales bacteriu | SAMN18076853 | aquatic | 10 | Yes | 4 | NLPC_P60;DUF4240;TPR_MalT | — |
| 2026780.SAMN14533038.JABAAE010 | **typeIII-A5** | 188,6 | low | Planctomycetota | — | — | — | — | Planctomycetota bacterium | SAMN14533038 | aquatic | 10 | Yes | 3 | DUF1579;Sulfatase | — |
| 2035772.SAMN19297679.JAHIOU0100 | **typeIII-A5** | 187,4 | low | Candidatus Omnitrophota | — | — | — | — | Candidatus Omnitrophota bacterium | SAMN19297679 | terrestrial | 5 | Yes | 4 | IZUMO;PDDEXK_1;WDR62-MABI | — |
| 1430440.SAMEA3138838.HG794546_3 | **typeIII-A5** | 185,2 | low | Pseudomonadota | Alphaproteobacteria | Rhodospirillales | Rhodospirillaceae | Magnetospirillum | Magnetospirillum gryphiswaldense | SAMEA3138838 | aquatic | 10 | Yes | 4 | SBP_bac_3;DDE_Tnp_IS66;DUF7 | — |
| 194843.SAMEA6151249.CADAAI01000 | **typeIII-A5** | 182 | low | Bacteroidota | Bacteroidia | Bacteroidales | — | — | uncultured Bacteroidales bacterium | SAMEA6151249 | host-associated | 9 | Yes | 4 | DUF6254;SH3_3;Flavin_Reduct | — |
| 2597224.SAMN10849879.SHMQ01000 | **typeIII-A5** | 179,7 | low | — | Deltaproteobacteria | Candidatus Acidulodesulf | — | Candidatus Acidulodes | Candidatus Acidulodesulfobacterium a | SAMN10849879 | anthropogenic | 10 | Yes | 4 | UBA_3;Phage_ORF5 | — |
| 1936112.SAMN14533044.JABABI0100 | **typeIII-A5** | 178,2 | low | Planctomycetota | Planctomycetia | Planctomycetales | Planctomycetaceae | Fuerstiella | Fuerstiella sp. | SAMN14533044 | aquatic | 10 | Yes | 4 | Flg_hook;DUF6702 | — |
| 2597224.SAMN10849879.SHMQ01000 | **typeIII-A5** | 173,9 | low | — | Deltaproteobacteria | Candidatus Acidulodesulf | — | Candidatus Acidulodes | Candidatus Acidulodesulfobacterium a | SAMN10849879 | anthropogenic | 10 | Yes | 4 | UBA_3;Phage_ORF5;Ribosomal_L | — |
| 370804.SAMEA104667047.ONJX0100 | **typeIII-A5** | 173,6 | low | Bacteroidota | Bacteroidia | Bacteroidales | Prevotellaceae | — | uncultured Prevotellaceae bacterium | SAMEA104667047 | host-associated | 10 | Yes | 4 | MapZ_EC1;DUF6254;HepT-like | — |
| 194843.SAMEA6151907.CADAZL0100 | **typeIII-A5** | 170 | low | Bacteroidota | Bacteroidia | Bacteroidales | — | — | uncultured Bacteroidales bacterium | SAMEA6151907 | host-associated | 10 | Yes | 4 | MapZ_EC1;DUF6254;OMP_b-brl | — |
| 2778646.SAMN16384127.JADMNM010 | **typeIII-A5** | 169,7 | low | Pseudomonadota | Alphaproteobacteria | Sphingomonadales | Erythrobacteraceae | Qipengyuania | Qipengyuania sp. YIM B01966 | SAMN16384127 | anthropogenic | 6 | Yes | 3 | MarR_2;ABC1;Phage_int_SAM_5 | — |
| 1961416.SAMN06454634.DBZJ010002 | **typeIII-A5** | 160 | low | Thermodesulfobacteriota | Syntrophia | Syntrophales | Syntrophaceae | — | Syntrophaceae bacterium UBA1163 | SAMN06454634 | salinity | 10 | Yes | 4 | DUF7683;HTH_3;AGTRAP | — |
| 2268206.SAMN13894326.JAAYWD010 | **typeIII-A5** | 159,5 | low | Thermotogota | Thermotogae | Thermotogales | Thermotogaceae | — | Thermotogaceae bacterium | SAMN13894326 | anthropogenic | 10 | Yes | 4 | AAA_2;OrfB_IS605;TM2;DUF4350 | — |
| 2212472.SAMN12876714.WGDB01000 | **typeIII-A5** | 131,6 | low | Campylobacterota | Epsilonproteobacteria | Campylobacterales | Helicobacteraceae | — | Helicobacteraceae bacterium | SAMN12876714 | temperature | 10 | Yes | 4 | Nitro_FeMo-Co;Rhodanese;Ham1p | — |
| 129578.SAMN06648058.NQJF0100000 | **typeIV** | 345,1 | high | Pseudomonadota | Gammaproteobacteria | Aeromonadales | Aeromonadaceae | Oceanimonas | Oceanimonas baumannii | SAMN06648058 | anthropogenic | 6 | Yes | 2 | ANC1_spectrin;Zn_ribbon_double | — |
| 153809.SAMEA6953771.CAITHT01000 | **typeIV** | 337,2 | high | Pseudomonadota | — | — | — | — | uncultured Pseudomonadota bacterium | SAMEA6953771 | aquatic | 11 | Yes | 4 | DUF2730;Retron_Ec48_antiviral;Re | — |
| 1236544.SAMD00046904.BCZT010001 | **typeIV** | 333,7 | high | Pseudomonadota | Gammaproteobacteria | Alteromonadales | Shewanellaceae | Shewanella | Shewanella algae | SAMD00046904 | host-associated | 2 | Yes | 1 | NERD | — |
| 114715.SAMEA9694635.CAJXMR0100 | **typeIV** | 332,2 | high | Pseudomonadota | Betaproteobacteria | Burkholderiales | Burkholderiaceae | Ralstonia | uncultured Ralstonia sp. | SAMEA9694635 | aquatic | 8 | Yes | 4 | Mrr_cat;Retron_Ec48_antiviral | — |
| 472181.SAMN09638568.DRHT010000 | **typeIV** | 314,7 | high | Pseudomonadota | Gammaproteobacteria | Pseudomonadales | Pseudomonadaceae | Halopseudomonas | Halopseudomonas sabulinigri | SAMN09638568 | aquatic | 7 | Yes | 3 | EamA;Iso_dh;Retron_Ec48_antivira | — |
| 1406866.SAMN02732283.AXZR010000 | **typeIX** | 292,3 | high | Pseudomonadota | Alphaproteobacteria | Rhodobacterales | Roseobacteraceae | Sulfitobacter | Sulfitobacter pontiacus | SAMN02732283 | aquatic | 10 | Yes | 4 | DUF6602;MarR;GIY-YIG | — |
| 1898112.SAMN14915609.JABJKH0100 | **typeIX** | 289,5 | high | Pseudomonadota | Alphaproteobacteria | Rhodospirillales | Rhodospirillaceae | — | Rhodospirillaceae bacterium | SAMN14915609 | aquatic | 9 | Yes | 4 | DUF2779;ARMET_N | — |
| 1977087.SAMN19298580.JAHJTB0100 | **typeIX** | 288,4 | high | Pseudomonadota | — | — | — | — | Pseudomonadota bacterium | SAMN19298580 | terrestrial | 8 | Yes | 4 | HDOD;HD_5;GGDEF_2;TrkA_N | — |
| 383381.SAMN02737000.JMIV0100000 | **typeIX** | 281,3 | high | Pseudomonadota | Alphaproteobacteria | Sphingomonadales | Erythrobacteraceae | Erythrobacter | Erythrobacter sp. JL475 | SAMN02737000 | aquatic | 11 | Yes | 4 | NMB0537_N;Phage_integrase;ARM | — |
| 2026779.SAMN10607091.SKZY010003 | **typeIX** | 259,4 | high | Planctomycetota | Planctomycetia | Planctomycetales | Planctomycetaceae | — | Planctomycetaceae bacterium | SAMN10607091 | aquatic | 7 | Yes | 3 | GGDEF_2;MauE;PIN | — |
| 998088.SAMN02603940.CP002607_39 | **typeV** | 335,4 | high | Pseudomonadota | Gammaproteobacteria | Aeromonadales | Aeromonadaceae | Aeromonas | Aeromonas veronii | SAMN02603940 | host-associated | 11 | Yes | 4 | CSD | — |
| 1196835.SAMN02603608.CP003677_4 | **typeV** | 328,3 | high | Pseudomonadota | Gammaproteobacteria | Pseudomonadales | Pseudomonadaceae | Stutzerimonas | Stutzerimonas stutzeri | SAMN02603608 | terrestrial | 11 | Yes | 4 | GTP_EFTU;Tom5;CSD;VirionAsse | — |
| 56192.SAMN07327741.PYLU01000035 | **typeV** | 321,7 | high | Pseudomonadota | Gammaproteobacteria | Vibrionales | Vibrionaceae | Photobacterium | Photobacterium iliopiscarium | SAMN07327741 | host-associated | 8 | Yes | 4 | Clathrin-link;CSD;CbiA | — |
| 518738.SAMN13287348.JAACSM0100 | **typeV** | 320,3 | high | Pseudomonadota | Gammaproteobacteria | Alteromonadales | Shewanellaceae | Shewanella | Shewanella vesiculosa | SAMN13287348 | terrestrial | 11 | Yes | 4 | DUF6037;CSD;Clathrin-link;ASA5 | — |
| 1283291.SAMN11792376.VEDF010000 | **typeV** | 319,6 | high | Pseudomonadota | Gammaproteobacteria | Pseudomonadales | Pseudomonadaceae | Pseudomonas | Pseudomonas sp. URMO17WK12:I11 | SAMN11792376 | anthropogenic | 11 | Yes | 4 | LysR_substrate;CSD | — |
| 1882827.SAMN05443579.FOWG01000 | **typeV** | 298,2 | high | Pseudomonadota | Betaproteobacteria | Burkholderiales | Comamonadaceae | Variovorax | Variovorax sp. PDC80 | SAMN05443579 | anthropogenic | 11 | Yes | 4 | tRNA-synt_1c;Peptidase_M23;AstE | — |
| 83655.SAMN13547356.WVWB010000 | **typeV** | 296,2 | high | Pseudomonadota | Gammaproteobacteria | Enterobacterales | Enterobacteriaceae | Leclercia | Leclercia adecarboxylata | SAMN13547356 | anthropogenic | 11 | Yes | 4 | T6SS_HCP;SymE_toxin;DUF443;M | — |
| 1566294.SAMN03159417.FOWV01000 | **typeV** | 294,9 | high | Pseudomonadota | Betaproteobacteria | Burkholderiales | Burkholderiaceae | Ralstonia | Ralstonia sp. NFACC01 | SAMN03159417 | host-associated | 11 | Yes | 4 | Mrr_cat;CSD | — |
| 180197.SAMN02982919.FOGD010000 | **typeV** | 294,4 | high | Pseudomonadota | Betaproteobacteria | Burkholderiales | Comamonadaceae | Giesbergeria | Giesbergeria anulus | SAMN02982919 | anthropogenic | 11 | Yes | 4 | CBS;CSD;LYTB | — |
| 2775292.SAMN16237464.JACYUQ010 | **typeV** | 284,7 | high | Pseudomonadota | Betaproteobacteria | Burkholderiales | Oxalobacteraceae | — | Oxalobacteraceae sp. CFBP 13730 | SAMN16237464 | anthropogenic | 3 | Yes | 2 | TGS | — |
| 63.SAMN07313362.CP022423_1311 | **typeV** | 268 | borderline | Pseudomonadota | Betaproteobacteria | Neisseriales | Neisseriaceae | Vitreoscilla | Vitreoscilla filiformis | SAMN07313362 | anthropogenic | 10 | Yes | 4 | Transket_pyr;CSD;LysR_substrate | — |
| 1004785.SAMN02604118.CP003845_2 | **typeVI** | 266,7 | high | Pseudomonadota | Gammaproteobacteria | Alteromonadales | Alteromonadaceae | Alteromonas | Alteromonas macleodii | SAMN02604118 | aquatic | 11 | Yes | 4 | LCAT;DUF7379;HTH_3 | — |
| 562.SAMN13545722.CP047609_747 | **typeVI** | 260,8 | high | Pseudomonadota | Gammaproteobacteria | Enterobacterales | Enterobacteriaceae | Escherichia | Escherichia coli | SAMN13545722 | host-associated | 11 | Yes | 4 | HTH_3;FRG;Phage_portal | — |
| 748224.SAMN00189147.GL538343_33 | **typeVI** | 239,5 | high | Bacillota | Clostridia | Eubacteriales | Oscillospiraceae | Faecalibacterium | Faecalibacterium prausnitzii | SAMN00189147 | host-associated | 11 | Yes | 4 | HTH_3;DUF6050;Relaxase | — |
| 862719.PRJEA50367.FQ311872_41 | **typeVI** | 236,5 | high | Pseudomonadota | Alphaproteobacteria | Rhodospirillales | Azospirillaceae | Azospirillum | Azospirillum lipoferum | PRJEA50367 | aquatic | 11 | Yes | 4 | UPF0020;Amanitin;Beta_elim_lyase | — |
| 169435.SAMN15533212.JADNIE01000 | **typeVI** | 236,3 | high | Bacillota | Clostridia | Eubacteriales | Oscillospiraceae | Anaerotruncus | Anaerotruncus colihominis | SAMN15533212 | host-associated | 11 | Yes | 0 | — | — |
| 1979412.SAMN14944538.JABMOK010 | **typeVI** | 235,6 | high | Pseudomonadota | Gammaproteobacteria | Pseudomonadales | Pseudohongiellaceae | Pseudohongiella | Pseudohongiella sp. | SAMN14944538 | temperature | 11 | Yes | 4 | HTH_3;DUF2951;Acetyltransf_10 | — |
| 2044939.SAMN16344672.JAFTWE010 | **typeVI** | 235,3 | high | Bacillota | Clostridia | — | — | — | Clostridia bacterium | SAMN16344672 | host-associated | 11 | Yes | 4 | HTH_3;ADPrib_exo_Tox | — |
| 2485925.SAMN16343178.JAFRQP010 | **typeVI** | 235,1 | high | Bacillota | Clostridia | Eubacteriales | Oscillospiraceae | — | Oscillospiraceae bacterium | SAMN16343178 | host-associated | 11 | Yes | 4 | Mob_Pre;SlyX;Response_reg;HATI | — |
| 556261.SAMN02463701.GL870812_31 | **typeVI** | 234,3 | high | Bacillota | Clostridia | Eubacteriales | Clostridiaceae | Clostridium | Clostridium sp. D5 | SAMN02463701 | host-associated | 11 | Yes | 4 | HTH_3;KilA-N | — |
| 153809.SAMEA6946215.CAIXHP01000 | **typeVI** | 232,9 | high | Pseudomonadota | — | — | — | — | uncultured Pseudomonadota bacterium | SAMEA6946215 | aquatic | 11 | Yes | 4 | DNA_methylase;RE_SacI;HTH_3 | — |
| 2109688.SAMD00113750.AP018533_2 | **typeVI** | 232,4 | high | Bacillota | Clostridia | Eubacteriales | — | — | Clostridiales bacterium CCNA10 | SAMD00113750 | host-associated | 11 | Yes | 4 | Relaxase;DUF6050;HTH_3 | — |
| 1895845.SAMN05660629.MKWI01000 | **typeVI** | 231,2 | high | Pseudomonadota | Alphaproteobacteria | Sphingomonadales | Sphingobiaceae | Sphingobium | Sphingobium sp. 66-54 | SAMN05660629 | anthropogenic | 11 | Yes | 4 | ThsB_TIR;Resolvase;HTH_3;Dicer_ | — |

| | | | | | | | | | | | | | | | | |
|---|---|---|---|---|---|---|---|---|---|---|---|---|---|---|---|---|
| 1930536.SAMN17140639.JAFAZF0100 | **typeVI** | 230,1 | high | Pseudomonadota | Alphaproteobacteria | Sphingomonadales | Sphingomonadaceae | — | Sphingomonadaceae bacterium | SAMN17140639 | host-associated | 11 | Yes | 4 | HTH_3;DUF2278;His_Phos_1 | — |
| 759821.SAMN09630405.QOHO010000 | **typeVI** | 228,7 | high | Bacillota | Clostridia | Lachnospirales | Lachnospiraceae | Lacrimispora | Lacrimispora amygdalina | SAMN09630405 | anthropogenic | 6 | Yes | 2 | HTH_3 | — |
| 1410652.SAMN02744012.JHXK010000 | **typeVI** | 227,2 | high | Bacillota | Clostridia | Eubacteriales | Clostridiaceae | Clostridium | Clostridium beijerinckii | SAMN02744012 | host-associated | 9 | Yes | 4 | Imp-YgjV;HTH_3;HSDR_N_2;TelK_ | — |
| 395358.SAMEA9694658.CAJXPX0100 | **typeVI** | 226,8 | high | Pseudomonadota | Alphaproteobacteria | Hyphomonadales | Hyphomonadaceae | — | uncultured Hyphomonadaceae bacteriu | SAMEA9694658 | temperature | 10 | Yes | 4 | HEPN_Toprim_N;HTH_3 | — |
| 1031594.SAMN09074687.QPJU010000 | **typeVI** | 225,5 | high | Pseudomonadota | Betaproteobacteria | Burkholderiales | Comamonadaceae | Extensimonas | Extensimonas vulgaris | SAMN09074687 | anthropogenic | 11 | Yes | 4 | QatB;KAP_NTPase;HTH_3;ARM_L | — |
| 266264.SAMN02598450.CP000352_14 | **typeVI** | 225,5 | high | Pseudomonadota | Betaproteobacteria | Burkholderiales | Burkholderiaceae | Cupriavidus | Cupriavidus metallidurans | SAMN02598450 | — | 11 | Yes | 0 | — | — |
| 1895859.SAMN05660633.MKWM0100 | **typeVI** | 225,5 | high | Pseudomonadota | Betaproteobacteria | Nitrosomonadales | Thiobacillaceae | Thiobacillus | Thiobacillus sp. 63-78 | SAMN05660633 | — | 11 | Yes | 0 | — | — |
| 1940281.SAMN19405844.JAHRFE010 | **typeVI** | 224,3 | high | Pseudomonadota | Alphaproteobacteria | Hyphomicrobiales | Rhizobiaceae | Hoeflea | Hoeflea sp. | SAMN19405844 | terrestrial | 11 | Yes | 4 | Phage_integrase;Resolvase;HTH_3 | — |
| 1798255.SAMN04315987.MGXA01000 | **typeVI** | 223,8 | high | Pseudomonadota | Betaproteobacteria | Nitrosomonadales | — | — | Gallionellales bacterium RIFCSPLOW | SAMN04315987 | terrestrial | 11 | Yes | 4 | Rhodanese;ARM_LIN;DUF4325;HT | — |
| 1122616.SAMN02441143.AULT010000 | **typeVI** | 223,5 | high | Pseudomonadota | Gammaproteobacteria | Oceanospirillales | Oceanospirillaceae | Oceanospirillum | Oceanospirillum beijerinckii | SAMN02441143 | aquatic | 11 | Yes | 4 | YdiH;HTH_3;PIN;MazE_antitoxin | — |
| 2052162.SAMN11532980.VIKH010000 | **typeVI** | 222,3 | high | Pseudomonadota | Betaproteobacteria | Nitrosomonadales | Gallionellaceae | — | Gallionellaceae bacterium | SAMN11532980 | terrestrial | 11 | Yes | 4 | Bac_export_3;Bac_export_1;Chor_I | — |
| 1913988.SAMN19298773.JAHKAM010 | **typeVI** | 222,2 | high | Pseudomonadota | Alphaproteobacteria | — | — | — | Alphaproteobacteria bacterium | SAMN19298773 | anthropogenic | 11 | Yes | 4 | TonB_dep_Rec_b-barrel;UvrB_inter | — |
| 2779351.SAMN16401875.JADCKL010 | **typeVI** | 221,6 | high | Bacillota | Clostridia | Lachnospirales | Lachnospiraceae | Claveliimonas | Claveliimonas monacensis | SAMN16401875 | host-associated | 11 | Yes | 4 | BsuBI_PstI_RE;Eco57I;HTH_3 | — |
| 1236514.SAMD00010222.BAKL010001 | **typeVI** | 220,2 | high | Bacteroidota | Bacteroidia | Bacteroidales | Bacteroidaceae | Bacteroides | Bacteroides stercorirosoris | SAMD00010222 | host-associated | 9 | Yes | 4 | DUF5347;HTH_3;Fib_succ_major | — |
| 1321819.SAMN02436886.KE993103_2 | **typeVI** | 219 | high | Bacteroidota | Bacteroidia | Bacteroidales | Bacteroidaceae | Bacteroides | Bacteroides pyogenes | SAMN02436886 | host-associated | 7 | Yes | 3 | DDE_Tnp_ISL3;HTH_3;DUF5347 | — |
| 1232436.SAMEA2272249.CAPF010000 | **typeVI** | 218,8 | high | Actinomycetota | Coriobacteriia | Coriobacteriales | Coriobacteriaceae | Enorma | Enorma timonensis | SAMEA2272249 | host-associated | 11 | Yes | 4 | ResIII;N6_N4_Mtase;HTH_3 | — |
| 1123279.SAMN02440570.ATUS010000 | **typeVI** | 217,9 | high | Pseudomonadota | Gammaproteobacteria | Cellvibrionales | Spongiibacteraceae | Spongiibacter | Spongiibacter tropicus | SAMN02440570 | temperature | 11 | Yes | 4 | AAA_13;Apo-VLDL-II;HTH_3;ARM_ | — |
| 2137880.SAMEA8805625.CAJUJH010 | **typeVI** | 217,5 | high | Candidatus Melainabacte | — | Candidatus Gastranaerop | — | — | Candidatus Gastranaerophilales bacte | SAMEA8805625 | host-associated | 11 | Yes | 4 | Methylase_S;N6_Mtase;HTH_3;Mla | — |
| 2212467.SAMN16344105.JAFTAJ0100 | **typeVI** | 216 | high | Bacteroidota | Bacteroidia | Bacteroidales | Bacteroidaceae | — | Bacteroidaceae bacterium | SAMN16344105 | — | 8 | Yes | 0 | — | — |
| 370804.SAMEA6152912.CADCMC010 | **typeVI** | 216 | high | Bacteroidota | Bacteroidia | Bacteroidales | Prevotellaceae | — | uncultured Prevotellaceae bacterium | SAMEA6152912 | host-associated | 8 | Yes | 0 | — | — |
| 876091.SAMEA8805256.CAJTZK0100 | **typeVI** | 215,6 | high | Bacillota | Clostridia | Eubacteriales | Oscillospiraceae | Oscillibacter | uncultured Oscillibacter sp. | SAMEA8805256 | host-associated | 11 | Yes | 4 | DegV;Phage_integrase;DUF6809 | — |
| 194843.SAMEA6150337.CACYRK0100 | **typeVI** | 215,2 | high | Bacteroidota | Bacteroidia | Bacteroidales | — | — | uncultured Bacteroidales bacterium | SAMEA6150337 | host-associated | 9 | Yes | 4 | HATPase_c;Response_reg;TerC;A | — |
| 317.SAMN16237900.JACYWI01000001 | **typeVI** | 215,2 | high | Pseudomonadota | Gammaproteobacteria | Pseudomonadales | Pseudomonadaceae | Pseudomonas | Pseudomonas syringae | SAMN16237900 | host-associated | 11 | Yes | 4 | DUF4172;GFA | — |
| 853.SAMN07350508.NMTY01000004_ | **typeVI** | 215,2 | high | Bacillota | Clostridia | Eubacteriales | Oscillospiraceae | Faecalibacterium | Faecalibacterium prausnitzii | SAMN07350508 | host-associated | 11 | Yes | 4 | HD_5;Response_reg;DUF4300 | — |
| 718252.SAMEA3138378.FP929045_22 | **typeVI** | 215,1 | high | Bacillota | Clostridia | Eubacteriales | Oscillospiraceae | Faecalibacterium | Faecalibacterium prausnitzii | SAMEA3138378 | host-associated | 11 | Yes | 4 | RuvC;Relaxase | — |
| 748224.SAMN00189147.GL538328_11 | **typeVI** | 214,9 | high | Bacillota | Clostridia | Eubacteriales | Oscillospiraceae | Faecalibacterium | Faecalibacterium prausnitzii | SAMN00189147 | host-associated | 7 | Yes | 0 | — | — |
| 445971.SAMN02299418.DS560015_20 | **typeVI** | 214,7 | high | Bacillota | Clostridia | Eubacteriales | Eubacteriaceae | Anaerofustis | Anaerofustis stercorihominis | SAMN02299418 | host-associated | 11 | Yes | 4 | BtrH_N;Fungus-induced;Peptidase_ | — |
| 551459.SAMN06049055.MUGO010000 | **typeVI** | 214,2 | high | Bacteroidota | Flavobacteriia | Flavobacteriales | Weeksellaceae | Chryseobacterium | Chryseobacterium piscicola | SAMN06049055 | anthropogenic | 11 | Yes | 4 | DUF5347;HTH_3;NERD;Pdc1_Ge | — |
| 59823.SAMN16344881.JAFUEF010000 | **typeVI** | 214,1 | high | Bacteroidota | Bacteroidia | Bacteroidales | Prevotellaceae | Prevotella | Prevotella sp. | SAMN16344881 | — | 6 | Yes | 0 | — | — |
| 59620.SAMEA4892316.USGX0100008 | **typeVI** | 213,8 | high | Bacillota | Clostridia | Eubacteriales | Clostridiaceae | Clostridium | uncultured Clostridium sp. | SAMEA4892316 | host-associated | 2 | Yes | 1 | YnfE | — |
| 1776379.SAMEA3932115.LT556050_3 | **typeVI** | 213,6 | high | Bacteroidota | Bacteroidia | Bacteroidales | Prevotellaceae | Leyella | Leyella lascolaii | SAMEA3932115 | — | 11 | Yes | 0 | — | — |
| 537013.SAMN00008812.EQ973344_64 | **typeVI** | 213,5 | high | Bacillota | Clostridia | Eubacteriales | Oscillospiraceae | — | [Clostridium] methylpentosum | SAMN00008812 | — | 11 | Yes | 0 | — | — |
| 2485925.SAMN16342071.JAFQCV010 | **typeVI** | 212,5 | high | Bacillota | Clostridia | Eubacteriales | Oscillospiraceae | — | Oscillospiraceae bacterium | SAMN16342071 | host-associated | 11 | Yes | 4 | zf_PCFS4;Recombinase;Zona_CL2 | — |
| 2030927.SAMN14407242.JAAVFI0100 | **typeVI** | 211,7 | high | Bacteroidota | Bacteroidia | Bacteroidales | — | — | Bacteroidales bacterium | SAMN14407242 | host-associated | 11 | Yes | 4 | ResIII;Reeler | — |
| 2748031.SAMN16347300.JAFXXG010 | **typeVI** | 211,6 | high | Bacteroidota | Bacteroidia | Bacteroidales | Paludibacteraceae | — | Paludibacteraceae bacterium | SAMN16347300 | — | 5 | Yes | 0 | — | — |
| 59823.SAMN16343164.JAFRQB01000 | **typeVI** | 211,6 | high | Bacteroidota | Bacteroidia | Bacteroidales | Prevotellaceae | Prevotella | Prevotella sp. | SAMN16343164 | host-associated | 8 | Yes | 0 | — | — |
| 370804.SAMEA6149422.CACXIP01000 | **typeVI** | 211,2 | high | Bacteroidota | Bacteroidia | Bacteroidales | Prevotellaceae | — | uncultured Prevotellaceae bacterium | SAMEA6149422 | host-associated | 7 | Yes | 0 | — | — |
| 2044939.SAMN16347922.JAFYVE0100 | **typeVI** | 211,2 | high | Bacillota | Clostridia | — | — | — | Clostridia bacterium | SAMN16347922 | — | 11 | Yes | 0 | — | — |
| 2720829.SAMEA7202527.CAJFOI0100 | **typeVI** | 211,2 | high | Bacillota | Clostridia | — | — | Candidatus Avispirillum | Candidatus Avispirillum faecium | SAMEA7202527 | host-associated | 11 | Yes | 4 | Rhomboid;RE_MjaI | — |
| 1161415.SAMN02597443.AZYV010000 | **typeVI** | 211,2 | high | Bacillota | Bacilli | Lactobacillales | Streptococcaceae | Streptococcus | Streptococcus sp. BS29a | SAMN02597443 | host-associated | 11 | Yes | 4 | DUF4435;Methylase_S;DUF6341;D | — |
| 871327.SAMN05444001.FNVS0100002 | **typeVI** | 211 | high | Bacteroidota | Bacteroidia | Bacteroidales | Tannerellaceae | Parabacteroides | Parabacteroides chinchillae | SAMN05444001 | host-associated | 11 | Yes | 4 | HTH_17;FliS_cochap;YnfE;Virulenc | — |
| 2718.SAMEA8395222.CAJPRL010000 | **typeVI** | 210,6 | high | Pseudomonadota | Gammaproteobacteria | Cardiobacteriales | Cardiobacteriaceae | Cardiobacterium | Cardiobacterium hominis | SAMEA8395222 | host-associated | 9 | Yes | 4 | FUSC-like;Competence;YdiH | — |
| 1121870.SAMN02440607.AUAA01000 | **typeVI** | 210,3 | high | Bacteroidota | Flavobacteriia | Flavobacteriales | Weeksellaceae | Epilithonimonas | Epilithonimonas tenax | SAMN02440607 | — | 10 | Yes | 0 | — | — |
| 2485925.SAMEA6152165.CADBJK010 | **typeVI** | 209,7 | high | Bacillota | Clostridia | Eubacteriales | Oscillospiraceae | — | Oscillospiraceae bacterium | SAMEA6152165 | host-associated | 11 | Yes | 4 | Flg_new_2;Zn-C2H2_12 | — |
| 1796613.SAMN04621613.CP015401_8 | **typeVI** | 209,3 | high | Bacteroidota | Bacteroidia | Bacteroidales | Bacteroidaceae | Bacteroides | Bacteroides caecimuris | SAMN04621613 | host-associated | 11 | Yes | 0 | — | — |
| 194843.SAMEA7847407.CAJLWQ0100 | **typeVI** | 209 | high | Bacteroidota | Bacteroidia | Bacteroidales | — | — | uncultured Bacteroidales bacterium | SAMEA7847407 | host-associated | 7 | Yes | 3 | LMBR1;Fic | — |
| 1122147.SAMN02369451.AZFW01000 | **typeVI** | 207,9 | high | Bacillota | Bacilli | Lactobacillales | Lactobacillaceae | Schleiferilactobacillus | Schleiferilactobacillus harbinensis | SAMN02369451 | host-associated | 11 | Yes | 4 | GT2_TM_C;Peptidase_M78;HTH_ | — |
| 186571.SAMN10864689.SLXM0100001 | **typeVI** | 207,8 | high | Bacteroidota | Flavobacteriia | Flavobacteriales | Flavobacteriaceae | Tenacibaculum | Tenacibaculum skagerrakense | SAMN10864689 | aquatic | 11 | Yes | 4 | DUF5347;HTH_3;Peptidase_S58;V | — |
| 2795120.SAMN17004923.JAELUC010 | **typeVI** | 207,5 | high | Bacteroidota | Sphingobacteriia | Sphingobacteriales | Sphingobacteriaceae | Pedobacter | Pedobacter sp. ASV12 | SAMN17004923 | host-associated | 1 | No | 0 | — | — |
| 2048242.SAMEA104588875.LT985749_ | **typeVI** | 206,9 | high | Bacteroidota | Bacteroidia | Bacteroidales | Rikenellaceae | Alistipes | Alistipes sp. Marseille-P5061 | SAMEA104588875 | host-associated | 11 | Yes | 4 | ParB_N;HTH_3 | — |
| 165185.SAMEA7847541.CAJMER0100 | **typeVI** | 206,9 | high | Bacillota | Clostridia | Eubacteriales | Eubacteriaceae | Eubacterium | uncultured Eubacterium sp. | SAMEA7847541 | host-associated | 8 | Yes | 4 | PFam54_60;SseF;Fic;rve | — |
| 658143.SAMEA4891723.URKB010000 | **typeVI** | 206,3 | high | Mycoplasmatota | — | — | — | — | uncultured Mycoplasmatota bacterium | SAMEA4891723 | host-associated | 10 | Yes | 4 | DUF4010;Helicase_C;DUF4391 | — |
| 449673.SAMN00000004.DS499667_48 | **typeVI** | 205,9 | high | Bacteroidota | Bacteroidia | Bacteroidales | Bacteroidaceae | Bacteroides | Bacteroides stercoris | SAMN00000004 | — | 11 | Yes | 0 | — | — |
| 1898104.SAMN08179015.PMXD01000 | **typeVI** | 205,6 | high | Bacteroidota | — | — | — | — | Bacteroidota bacterium | SAMN08179015 | temperature | 11 | Yes | 4 | DUF5347;HTH_3;WG_beta_rep;PH | — |
| 1975676.SAMN05954931.MRCM01000 | **typeVI** | 205,5 | high | Bacteroidota | Flavobacteriia | Flavobacteriales | Flavobacteriaceae | Flavobacterium | Flavobacterium sp. ACN2 | SAMN05954931 | host-associated | 11 | Yes | 4 | DUF5347;HTH_3;Lipase_GDSL_2 | — |
| 290398.SAMN02598275.CP000285_12 | **typeVI** | 205,1 | high | Pseudomonadota | Gammaproteobacteria | Oceanospirillales | Halomonadaceae | Chromohalobacter | Chromohalobacter israelensis | SAMN02598275 | aquatic | 11 | Yes | 4 | Phage_integrase;PyocinActivator | — |
| 1298593.SAMEA2272589.HF680312_3 | **typeVI** | 205,1 | high | Pseudomonadota | Gammaproteobacteria | Oceanospirillales | Oceanospirillaceae | Thalassolituus | Thalassolituus oleivorans | SAMEA2272589 | aquatic | 11 | Yes | 4 | Peptidase_M19;Resolvase | — |
| 1191309.SAMN02470478.AJZH020000 | **typeVI** | 204,1 | high | Pseudomonadota | Gammaproteobacteria | Vibrionales | Vibrionaceae | Vibrio | Vibrio splendidus | SAMN02470478 | aquatic | 11 | Yes | 4 | SIS;Cys_Met_Meta_PP;HTH_3 | — |
| 2320122.SAMN10024804.RAYW01000 | **typeVI** | 203,8 | high | — | — | — | — | — | bacterium J10(2018) | SAMN10024804 | host-associated | 5 | Yes | 3 | ATPase_2;APC_r;DDE_Tnp_ISL3 | — |
| 1898104.SAMN19298609.JAHJUE0100 | **typeVI** | 203,4 | high | Bacteroidota | — | — | — | — | Bacteroidota bacterium | SAMN19298609 | terrestrial | 11 | Yes | 4 | Bul1_C;HTH_3;DUF2326;MC6 | — |
| 25.SAMN12272628.VKGK01000009_11 | **typeVI** | 202,8 | high | Pseudomonadota | Gammaproteobacteria | Alteromonadales | Shewanellaceae | Shewanella | Shewanella hanedai | SAMN12272628 | temperature | 11 | Yes | 4 | HTH_3;Fapy_DNA_glyco | — |
| 2049046.SAMN09989679.QYQM01000 | **typeVI** | 202,6 | high | Bacteroidota | Bacteroidia | Bacteroidales | Porphyromonadaceae | — | Porphyromonadaceae bacterium | SAMN09989679 | aquatic | 11 | Yes | 4 | DUF5347;HTH_3;Trypsin_2 | — |
| 1898104.SAMN21435362.JAIUNW010 | **typeVI** | 198,9 | high | Bacteroidota | — | — | — | — | Bacteroidota bacterium | SAMN21435362 | anthropogenic | 10 | Yes | 4 | YafQ_toxin;HTH_3;HTH_Tnp_1;rve | — |
| 2049048.SAMN11294459.SUZK010000 | **typeVI** | 194,2 | high | Bacteroidota | Bacteroidia | Bacteroidales | Rikenellaceae | — | Rikenellaceae bacterium | SAMN11294459 | host-associated | 11 | Yes | 4 | KAP_NTPase;GloB_C | — |
| 2026760.SAMN10744134.SCKI010000 | **typeVI** | 192,5 | high | Fidelibacterota | — | — | — | — | Fidelibacterota bacterium | SAMN10744134 | aquatic | 11 | Yes | 4 | Fer4_21;DsrE;Esterase | — |
| 880526.SAMN02441234.KE386488_19 | **typeVI** | 192,2 | high | Bacteroidota | Bacteroidia | Bacteroidales | Rikenellaceae | Rikenella | Rikenella microfusus | SAMN02441234 | host-associated | 11 | Yes | 4 | DUF6236 | — |
| 2052142.SAMN08179812.PMAT01000 | **typeVI** | 191,9 | high | Acidobacteriota | Terriglobia | Terriglobales | Acidobacteriaceae | — | Acidobacteriaceae bacterium | SAMN08179812 | terrestrial | 11 | Yes | 4 | DDE_Tnp_1_assoc;CBM_48;HTH_ | — |
| 2052142.SAMN08179208.PLGM01000 | **typeVI** | 190,6 | high | Acidobacteriota | Terriglobia | Terriglobales | Acidobacteriaceae | — | Acidobacteriaceae bacterium | SAMN08179208 | temperature | 6 | Yes | 2 | type_II_gspD_N0;HTH_3 | — |
| 1895771.SAMN05660602.MKVH01000 | **typeVI** | 189,2 | high | Candidatus Kapaibacteric | Candidatus Kapaibacte | Candidatus Kapaibacteria | Candidatus Kapaibacteria | Candidatus Kapaibacte | Candidatus Kapaibacterium thiocyanat | SAMN05660602 | anthropogenic | 11 | Yes | 4 | rve;HTH_Tnp_1;Nuc_deoxyrib_tr | — |
| 2282154.SAMN11533020.WSYB01000 | **typeVI** | 188,3 | high | Thermodesulfobacteriota | Desulfuromonadia | Geobacterales | Geobacteraceae | — | Geobacteraceae bacterium | SAMN11533020 | terrestrial | 11 | Yes | 4 | MTES_1575;SNase;HTH_3;DUF67 | — |
| 1978231.SAMN08912159.QHZN01000 | **typeVI** | 183,8 | high | Acidobacteriota | — | — | — | — | Acidobacteriota bacterium | SAMN08912159 | terrestrial | 11 | Yes | 4 | HTH_37;DUF5955;Big_2;STN | — |
| 2035772.SAMEA6955419.CAIQIN0100 | **typeVI** | 175,5 | high | Candidatus Omnitrophota | — | — | — | — | Candidatus Omnitrophota bacterium | SAMEA6955419 | aquatic | 11 | Yes | 4 | SPC25;PDDEXK_1;HTH_3;TPR_I | — |
| 2026778.SAMN13893689.JAAYCR010 | **typeVI** | 174,2 | high | Planctomycetota | Phycisphaerae | — | — | — | Phycisphaerae bacterium | SAMN13893689 | temperature | 11 | Yes | 4 | Resolvase;MarR_2;HTH_3;DAXX_ | — |
| 2528828.SAMN11124364.SOIB010001 | **typeVI** | 173,1 | high | Candidatus Stahliibacteric | — | — | — | — | Candidatus Stahliibacteriota bacterium | SAMN11124364 | aquatic | 3 | Yes | 2 | HTH_3;DUF7683 | — |
| 876.SAMN09779309.CP036295_2579 | **typeVI** | 172 | high | Thermodesulfobacteriota | Desulfovibrionia | Desulfovibrionales | Desulfovibrionaceae | Desulfovibrio | Desulfovibrio desulfuricans | SAMN09779309 | anthropogenic | 11 | Yes | 4 | HTH_3;Phage_integrase;Phage_co | — |
| 152509.SAMEA6955447.CAIVKC01000 | **typeVI** | 170,3 | high | Bacteroidota | — | — | — | — | uncultured Bacteroidota bacterium | SAMEA6955447 | aquatic | 11 | Yes | 4 | ABC2_membrane_3;Transglut_prok | — |
| 529385.SAMEA8805564.CAJUHZ0100 | **typeVI** | 167,7 | high | Thermodesulfobacteriota | Desulfovibrionia | Desulfovibrionales | Desulfovibrionaceae | Bilophila | uncultured Bilophila sp. | SAMEA8805564 | host-associated | 7 | Yes | 3 | Flagellin_C;HTH_3;Glyco_transf_9 | — |
| 58621.SAMN14908373.JACHGO01000 | **typeVI** | 163,5 | high | Thermodesulfobacteriota | Desulfovibrionia | Desulfovibrionales | Desulfovibrionaceae | Desulfovibrio | Desulfovibrio intestinalis | SAMN14908373 | host-associated | 11 | Yes | 4 | HTH_3;SEC-C;IncFII_repA | — |
| 2049043.SAMN19225054.JAHZEL0100 | **typeVI** | 162,8 | low | Thermodesulfobacteriota | Desulfovibrionia | Desulfovibrionales | Desulfovibrionaceae | — | Desulfovibrionaceae bacterium | SAMN19225054 | host-associated | 10 | Yes | 4 | Peptidase_S58;HATPase_c | — |

| | | | | | | | | | | | | | | | | |
|---|---|---|---|---|---|---|---|---|---|---|---|---|---|---|---|---|
| 2049428.SAMN16425662.JADJEI01000( | **typeVI** | 160,3 | low | Ignavibacteriota | Ignavibacteria | Ignavibacteriales | — | — | Ignavibacteriales bacterium | SAMN16425662 | anthropogenic | 10 | Yes | 4 | Y1_Tnp;HTH_3;LETM1_C | — |
| 2026779.SAMN13893644.JAAYAY0100 | **typeVI** | 157,9 | low | Planctomycetota | Planctomycetia | Planctomycetales | Planctomycetaceae | — | Planctomycetaceae bacterium | SAMN13893644 | temperature | 10 | Yes | 4 | DUF2461;HTH_31;HTH_PafC | — |
| 576117.SAMN04432131.LRUD0100000 | **typeVII-A1** | 351 | high | Pseudomonadota | Alphaproteobacteria | Rhodobacterales | Roseobacteraceae | Celeribacter | Celeribacter halophilus | SAMN04432131 | aquatic | 11 | Yes | 4 | MerR-DNA-bind;Hydrolase;Lsr2_DN | — |
| 1912891.SAMN07426584.QFOA010000( | **typeVII-A1** | 327,6 | high | Pseudomonadota | Alphaproteobacteria | Sphingomonadales | Sphingobiaceae | Sphingobium | Sphingobium sp. | SAMN07426584 | anthropogenic | 11 | Yes | 4 | TPR_14 | — |
| 1977087.SAMN21435408.JAIUOU0100 | **typeVII-A1** | 321,3 | high | Pseudomonadota | — | — | — | — | Pseudomonadota bacterium | SAMN21435408 | — | 7 | Yes | 0 | — | — |
| 1035191.SAMN02436690.KB291762_8 | **typeVII-A1** | 320 | high | Pseudomonadota | Alphaproteobacteria | Caulobacterales | Caulobacteraceae | Brevundimonas | Brevundimonas diminuta | SAMN02436690 | terrestrial | 9 | Yes | 4 | DUF2200;MPTase-PolyVal;DUF29 | — |
| 751586.SAMN02469913.GL883086_93 | **typeVII-A1** | 320 | high | Pseudomonadota | Alphaproteobacteria | Caulobacterales | Caulobacteraceae | Brevundimonas | Brevundimonas diminuta | SAMN02469913 | — | 11 | Yes | 0 | — | — |
| 1977087.SAMN08158042.PKDC010000( | **typeVII-A1** | 319,1 | high | Pseudomonadota | — | — | — | — | Pseudomonadota bacterium | SAMN08158042 | terrestrial | 11 | Yes | 4 | Hydrolase;Thioredoxin_4;SseB | — |
| 2282150.SAMN18059876.JAFLCT0100 | **typeVII-A1** | 294,3 | borderline | Pseudomonadota | Alphaproteobacteria | Caulobacterales | — | — | Caulobacterales bacterium | SAMN18059876 | anthropogenic | 4 | Yes | 3 | Cpn10;Cpn60_TCP1;DUF4168 | — |
| 1797342.SAMN04313819.MEOF010000( | **typeVII-A2** | 303,1 | high | Bacteroidota | — | — | — | — | Bacteroidetes bacterium GWF2_33_3 | SAMN04313819 | terrestrial | 11 | Yes | 4 | DUF6602;DUF3800;PFK;PfkB | — |
| 1948560.SAMN06451549.DIWM010000( | **typeVII-A2** | 291,3 | borderline | — | Deltaproteobacteria | — | — | — | Deltaproteobacteria bacterium UBA61( | SAMN06451549 | anthropogenic | 10 | Yes | 4 | Trypsin_2;DUF3800;AlaDh_PNT_C | — |
| 1977087.SAMN15717994.JAGDMU010 | **typeVII-A2** | 174,9 | low | Pseudomonadota | — | — | — | — | Pseudomonadota bacterium | SAMN15717994 | aquatic | 10 | Yes | 4 | Polyketide_cyc;Phage_integrase;Cp | — |
| 1032480.SAMD00061117.AP012204_4 | **typeVII-A2** | 167,2 | low | Actinomycetota | Actinomycetes | Propionibacteriales | Propionibacteriaceae | Microlunatus | Microlunatus phosphovorus | SAMD00061117 | anthropogenic | 10 | Yes | 4 | DDE_Tnp_1_assoc;Complex1_LYR | — |
| 1948743.SAMN06451839.DLRY010000 | **typeX** | 326,4 | high | Verrucomicrobiota | Opitutia | — | — | — | Opitutia bacterium UBA7876 | SAMN06451839 | aquatic | 8 | Yes | 4 | 3keto-disac_hyd;MTES_1575;Bac_ | — |
| 2026801.SAMN13495009.WTHK01000 | **typeX** | 323,6 | high | Verrucomicrobiota | Verrucomicrobiia | Verrucomicrobiales | — | — | Verrucomicrobiales bacterium | SAMN13495009 | aquatic | 6 | Yes | 2 | TRAP-delta;Abhydrolase_7 | — |
| 2026801.SAMN07620217.PAPL010000 | **typeX** | 310,1 | high | Verrucomicrobiota | Verrucomicrobiia | Verrucomicrobiales | — | — | Verrucomicrobiales bacterium | SAMN07620217 | aquatic | 11 | Yes | 4 | ATLF;Bac_DNA_binding;Abhydrola | — |
| 2026779.SAMN07619713.PBOJ010001 | **typeX** | 308,1 | high | Planctomycetota | Planctomycetia | Planctomycetales | Planctomycetaceae | — | Planctomycetaceae bacterium | SAMN07619713 | aquatic | 8 | Yes | 4 | HTH_17;DUF1501;Sulfatase | — |
| 2026801.SAMN10967627.DUBC010000( | **typeX** | 306,7 | high | Verrucomicrobiota | Verrucomicrobiia | Verrucomicrobiales | — | — | Verrucomicrobiales bacterium | SAMN10967627 | aquatic | 11 | Yes | 4 | Abhydrolase_7;Bac_DNA_binding;A | — |
| 2026779.SAMN08017441.DLVN010000 | **typeX** | 306,6 | high | Planctomycetota | Planctomycetia | Planctomycetales | Planctomycetaceae | — | Planctomycetaceae bacterium | SAMN08017441 | aquatic | 7 | Yes | 3 | DRL_cat | — |
| 2026779.SAMN07619712.PBOU010000( | **typeX** | 304,3 | borderline | Planctomycetota | Planctomycetia | Planctomycetales | Planctomycetaceae | — | Planctomycetaceae bacterium | SAMN07619712 | aquatic | 7 | Yes | 4 | GlpR;DUF1501;Sulfatase | — |
| 2026771.SAMN14914005.JABHBR010( | **typeX** | 303 | borderline | Verrucomicrobiota | Opitutia | — | — | — | Opitutia bacterium | SAMN14914005 | aquatic | 3 | Yes | 3 | PA14;GidB;Bac_DNA_binding | — |
| 2026779.SAMN14915429.JABJDJ0100( | **typeX** | 283,6 | borderline | Planctomycetota | Planctomycetia | Planctomycetales | Planctomycetaceae | — | Planctomycetaceae bacterium | SAMN14915429 | aquatic | 10 | Yes | 4 | DUF4231;Ferritin | — |
| 153809.SAMEA6953877.CAINMA0100( | **typeX** | 237,6 | low | Pseudomonadota | — | — | — | — | uncultured Pseudomonadota bacterium | SAMEA6953877 | aquatic | 8 | Yes | 4 | FGE-sulfatase;Bac_DNA_binding | — |
| 1506601.SAMEA7846325.CAJKAR010( | **typeX** | 224,1 | low | Lentisphaerota | Lentisphaeria | — | — | — | uncultured Lentisphaeria bacterium | SAMEA7846325 | host-associated | 4 | Yes | 3 | HMGL-like;CPSase_L_D2;Lipocalin | — |
| 1506601.SAMEA7847370.CAJMAW01( | **typeX** | 212,7 | low | Lentisphaerota | Lentisphaeria | — | — | — | uncultured Lentisphaeria bacterium | SAMEA7847370 | — | 10 | Yes | 4 | NRDD;PGA_cap;N_methyl;Glycos_ | — |
| 2026780.SAMN09638790.DRQF010000( | **typeX** | 187,5 | low | Planctomycetota | — | — | — | — | Planctomycetota bacterium | SAMN09638790 | temperature | 9 | Yes | 4 | ALGX;Peptidase_M28;Lipase_GDS | — |
| 490829.SAMN05421850.FNEB0100002 | **typeX** | 173 | low | Pseudomonadota | Alphaproteobacteria | Rhodobacterales | Roseobacteraceae | Lutimaribacter | Lutimaribacter saemankumensis | SAMN05421850 | aquatic | 10 | Yes | 4 | OPT;Hydrolase;TetR_N | — |
| 208544.SAMEA6951753.CAIVGK0100( | **typeX** | 168,3 | low | Pseudomonadota | Betaproteobacteria | Burkholderiales | — | — | uncultured Burkholderiales bacterium | SAMEA6951753 | aquatic | 2 | Yes | 2 | rve;HTH_Tnp_1 | — |
| 1674809.SAMN03837491.LUYM010000( | **typeX** | 150,6 | low | Actinomycetota | Actinomycetes | Actinomycetales | — | — | Actinomycetales bacterium Actino_02 | SAMN03837491 | host-associated | 10 | Yes | 4 | Nitro_FeMo-Co;DUF1919;AAA_15; | — |
| 1215112.SAMD00046934.BDAI0100000 | **typeXI** | 317,3 | high | Pseudomonadota | Gammaproteobacteria | Pseudomonadales | Pseudomonadaceae | Pseudomonas | Pseudomonas nitroreducens | SAMD00046934 | host-associated | 11 | Yes | 4 | DUF3732;HEPN_RiboL-PSP;YagK | — |
| 425022.SAMEA6956137.CAIWDD0100 | **typeXI** | 310,9 | high | Cyanobacteriota | Cyanophyceae | Nostocales | Nodulariaceae | Nodularia | uncultured Nodularia sp. | SAMEA6956137 | aquatic | 11 | Yes | 4 | Ferritin;ParB_N;T1-F;Sarcoglycan_ | — |
| 1211.SAMEA6946765.CAINGU0100000 | **typeXI** | 307,4 | high | Cyanobacteriota | — | — | — | — | uncultured cyanobacterium | SAMEA6946765 | aquatic | 11 | Yes | 4 | KIX_2;PIN_3;DUF6972 | — |
| 2024841.SAMN10967645.DUBU010000( | **typeXI** | 304,8 | high | Bdellovibrionota | Bdellovibrionia | Bdellovibrionales | Pseudobdellovibrionacea | Micavibrio | Micavibrio sp. | SAMN10967645 | aquatic | 7 | Yes | 3 | Resolvase;HATPase_c_3 | — |
| 2692823.SAMN13702774.JACJPD0100 | **typeXI** | 304,1 | high | Cyanobacteriota | Cyanophyceae | Oscillatoriales | Microcoleaceae | Microcoleus | Microcoleus sp. FACHB-53 | SAMN13702774 | terrestrial | 11 | Yes | 4 | Acetyltransf_1;DOPA_dioxygen;RB | — |
| 1919144.SAMN06020721.PXPI0100010 | **typeXI** | 303,9 | high | Cyanobacteriota | — | — | — | — | Cyanobacteria bacterium SW_9_44_5 | SAMN06020721 | salinity | 6 | Yes | 2 | adh_short;GTP_cyclohydrol | — |
| 32049.SAMN01081740.CP000955_26 | **typeXI** | 303,3 | high | Cyanobacteriota | Cyanophyceae | Chroococcales | Geminocystaceae | Picosynechococcus | Picosynechococcus sp. PCC 7002 | SAMN01081740 | aquatic | 11 | Yes | 4 | DUF5516;Endonuclease_NS;AAA_ | — |
| 1419583.SAMN02770242.AZQQ010000( | **typeXI** | 302,9 | high | Pseudomonadota | Gammaproteobacteria | Pseudomonadales | Pseudomonadaceae | Pseudomonas | Pseudomonas mandelii | SAMN02770242 | terrestrial | 11 | Yes | 4 | Fic;DUF4238;MatE | — |
| 1850361.SAMN05003984.LXYR010000 | **typeXI** | 302,1 | high | Cyanobacteriota | Cyanophyceae | Leptolyngbyales | Leptolyngbyaceae | Phormidesmis | Phormidesmis priestleyi | SAMN05003984 | temperature | 8 | Yes | 4 | Amidase;MazE_antitoxin;SLFN_Alb | — |
| 1961416.SAMN06454634.DBZJ010002 | **typeXI** | 301,5 | high | Thermodesulfobacteriota | Syntrophia | Syntrophales | Syntrophaceae | — | Syntrophaceae bacterium UBA1163 | SAMN06454634 | salinity | 6 | Yes | 2 | Adhesin_E;SPICE | — |
| 2773166.SAMN13287367.JAACTD010( | **typeXI** | 299,6 | high | Cyanobacteriota | — | — | — | — | Cyanobacteria bacterium CG_2015-16 | SAMN13287367 | terrestrial | 11 | Yes | 4 | YgjP-like;TUSC1;Phasin;AAA_33 | — |
| 312306.SAMN10523265.CP034337_76 | **typeXI** | 299,1 | high | Pseudomonadota | Gammaproteobacteria | Pseudomonadales | Pseudomonadaceae | Pseudomonas | Pseudomonas entomophila | SAMN10523265 | host-associated | 11 | Yes | 4 | Transposase_mut;HATPase_c_3 | — |
| 1916131.SAMN05959523.MPNC010000 | **typeXI** | 293,9 | high | Chloroflexota | Dehalococcoidia | — | — | — | SAR202 cluster bacterium bin87 | SAMN05959523 | host-associated | 11 | Yes | 3 | gag-asp_proteas;CoA_transf_3;adh | — |
| 1850361.SAMN05003984.LXYR010000 | **typeXI** | 292,7 | high | Cyanobacteriota | Cyanophyceae | Leptolyngbyales | Leptolyngbyaceae | Phormidesmis | Phormidesmis priestleyi | SAMN05003984 | temperature | 11 | Yes | 4 | AF2212-like;TPR_16;DUF4953 | — |
| 2013652.SAMN06767553.PHET010000( | **typeXI** | 289,5 | high | Actinomycetota | — | — | — | — | Actinobacteria bacterium HGW-Actinol | SAMN06767553 | terrestrial | 8 | Yes | 4 | MC8;DUF2326;MazG-like;SLFN-g3 | — |
| 2035772.SAMN19298168.JAHJHP0100 | **typeXI** | 287,5 | high | Candidatus Omnitrophota | — | — | — | — | Candidatus Omnitrophota bacterium | SAMN19298168 | terrestrial | 9 | Yes | 4 | HEPN_SAV_6107;Phage_integrase | — |
| 1656094.SAMN05511237.MDHN01000 | **typeXI** | 285,9 | high | Pseudomonadota | Gammaproteobacteria | Alteromonadales | Alteromonadaceae | Alteromonas | Alteromonas confluentis | SAMN05511237 | aquatic | 8 | Yes | 4 | HATPase_c;dCMP_cyt_deam_1;Ca | — |
| 1310370.SAMN15791687.JACKZK010( | **typeXI** | 285,4 | high | Pseudomonadota | Gammaproteobacteria | Pseudomonadales | Pseudomonadaceae | Pseudomonas | Pseudomonas sp. MSSRFD41 | SAMN15791687 | anthropogenic | 8 | Yes | 4 | Glyco_hydro_19;Phage_integrase;H | — |
| 195105.SAMN04279595.NIPU0100001 | **typeXI** | 282,5 | high | Pseudomonadota | Alphaproteobacteria | Rhodobacterales | Paracoccaceae | Haematobacter | Haematobacter massiliensis | SAMN04279595 | host-associated | 9 | Yes | 4 | Zn_ribbon_recom;AAA_15;UvrD-he | — |
| 1881068.SAMN05428950.FNZB010000 | **typeXI** | 282,3 | high | Pseudomonadota | Alphaproteobacteria | Sphingomonadales | Sphingomonadaceae | Sphingomonas | Sphingomonas sp. OV641 | SAMN05428950 | anthropogenic | 11 | Yes | 4 | HATPase_c;DCD;AGS_C;TniB | — |
| 2722791.SAMN14402108.JAAVJH0100 | **typeXI** | 282,3 | high | Pseudomonadota | Alphaproteobacteria | Sphingomonadales | Sphingomonadaceae | Sphingomonas | Sphingomonas corticis | SAMN14402108 | — | 11 | Yes | 0 | — | — |
| 481743.SAMN00191238.CP001793_26 | **typeXI** | 278,9 | high | Bacillota | Bacilli | Bacillales | Paenibacillaceae | Paenibacillus | Paenibacillus sp. Y412MC10 | SAMN00191238 | host-associated | 11 | Yes | 4 | RNB;SmpB;TIR_2;nSTAND3 | — |
| 640205.SAMN05216381.FNBM0100000 | **typeXI** | 278,6 | high | Pseudomonadota | Gammaproteobacteria | Pseudomonadales | Pseudomonadaceae | Phytopseudomonas | Phytopseudomonas seleniipraecipitans | SAMN05216381 | host-associated | 11 | Yes | 4 | Bflower;YobI-ATPase;Acetyltransf_ | — |
| 2583453.SAMN11812194.VCIG010000 | **typeXI** | 278,4 | high | Pseudomonadota | Alphaproteobacteria | Hyphomicrobiales | Brucellaceae | Ochrobactrum | Ochrobactrum sp. CGA5 | SAMN11812194 | host-associated | 11 | Yes | 4 | Resolvase;FRG | — |
| 367825.SAMN05904704.CP017754_18 | **typeXI** | 278,4 | high | Pseudomonadota | Betaproteobacteria | Burkholderiales | Burkholderiaceae | Cupriavidus | Cupriavidus malaysiensis | SAMN05904704 | anthropogenic | 11 | Yes | 4 | PAAR_motif;SRAP;HTH_31;T5orf1 | — |
| 1230476.SAMN01893863.KE747856_1 | **typeXI** | 275,8 | high | Pseudomonadota | Alphaproteobacteria | Hyphomicrobiales | Nitrobacteraceae | Bradyrhizobium | Bradyrhizobium sp. DFCI-1 | SAMN01893863 | anthropogenic | 11 | Yes | 4 | GST_N_2;AMNp_N | — |
| 1874826.SAMN16426229.JADJYM010( | **typeXI** | 275,1 | high | Pseudomonadota | Alphaproteobacteria | Sphingomonadales | Sphingomonadaceae | Novosphingobium | Novosphingobium sp. | SAMN16426229 | anthropogenic | 11 | Yes | 4 | DUF7007;Trypsin_2;Trep_dent_lipo | — |
| 1802172.SAMN04315629.MIAM010000 | **typeXI** | 275 | high | Pseudomonadota | Alphaproteobacteria | Sphingomonadales | Sphingopyxidaceae | Sphingopyxis | Sphingopyxis sp. RIFCSPHIGHO2_12 | SAMN04315629 | anthropogenic | 11 | Yes | 4 | Glutaredoxin;DUF1178;KAP_NTPa | — |
| 1913989.SAMN19297468.JAHIGR0100 | **typeXI** | 274,8 | high | Pseudomonadota | Gammaproteobacteria | — | — | — | Gammaproteobacteria bacterium | SAMN19297468 | terrestrial | 10 | Yes | 4 | Oxidored_FMN;HTH_60;NAD_bind | — |
| 277961.SAMN11791491.VCGW010000 | **typeXI** | 274,6 | high | Pseudomonadota | Gammaproteobacteria | Pseudomonadales | Marinobacteraceae | Marinobacter | Marinobacter maritimus | SAMN11791491 | aquatic | 11 | Yes | 4 | TIR_2;PhiKZ_VTX;Thioredoxin;Tau | — |
| 1660088.SAMN03652433.MEDN010000 | **typeXI** | 274,1 | high | Pseudomonadota | Alphaproteobacteria | Hyphomicrobiales | Rhizobiaceae | Agrobacterium | Agrobacterium sp. SCN 61-19 | SAMN03652433 | anthropogenic | 11 | Yes | 4 | Pribosyltran;Dak2;APH;GCV_T | — |
| 1903720.SAMEA5278229.CAAGCO010 | **typeXI** | 273,2 | high | Bacillota | Bacilli | — | — | — | Bacilli bacterium | SAMEA5278229 | host-associated | 11 | Yes | 4 | Thr_synth_N;Hydrolase;PLK4_bind | — |
| 2013842.SAMN06767759.PGXO010000( | **typeXI** | 273,2 | high | Synergistota | — | — | — | — | Synergistetes bacterium HGW-Synerg | SAMN06767759 | aquatic | 11 | Yes | 4 | NOV_C;Peptidase_S24;HTH_3 | — |
| 2508294.SAMEA5537675.CABPRY010( | **typeXI** | 270,5 | high | Pseudomonadota | Betaproteobacteria | Burkholderiales | Burkholderiaceae | Pandoraea | Pandoraea cepalis | SAMEA5537675 | anthropogenic | 7 | Yes | 3 | AAA_15;GIY-YIG;AMP-binding | — |
| 2762296.SAMN15732460.JACOGB010( | **typeXI** | 270 | high | Pseudomonadota | Betaproteobacteria | Burkholderiales | Oxalobacteraceae | Undibacterium | Undibacterium sp. FT79W | SAMN15732460 | terrestrial | 11 | Yes | 4 | Cys_rich_CPCC;TniQ | — |
| 2775282.SAMN16237746.JACYUG010( | **typeXI** | 269,4 | high | Pseudomonadota | Alphaproteobacteria | Sphingomonadales | Sphingomonadaceae | Sphingomonas | Sphingomonas sp. CFBP 8760 | SAMN16237746 | host-associated | 11 | Yes | 4 | AAA_15;DUF5131 | — |
| 1886787.SAMN11366401.SSTI0100002 | **typeXI** | 264,2 | high | Pseudomonadota | Alphaproteobacteria | Sphingomonadales | Sphingomonadaceae | Sphingomonas | Sphingomonas olei | SAMN11366401 | host-associated | 8 | Yes | 4 | MarR_2;SIR2_2;Piwi | — |
| 409.SAMN20165186.JAIETD01000006 | **typeXI** | 261,2 | high | Pseudomonadota | Alphaproteobacteria | Hyphomicrobiales | Methylobacteriaceae | Methylobacterium | Methylobacterium sp. | SAMN20165186 | aquatic | 3 | Yes | 2 | DUF2958 | — |
| 331632.SAMEA6152175.CADBJR0100( | **typeXI** | 261,1 | high | Actinomycetota | Coriobacteriia | Coriobacteriales | Coriobacteriaceae | — | uncultured Coriobacteriaceae bacteriu | SAMEA6152175 | host-associated | 6 | Yes | 2 | FRG | — |
| 208549.SAMEA6954897.CAIZEI010000 | **typeXI** | 259 | high | Pseudomonadota | Alphaproteobacteria | Hyphomicrobiales | — | — | uncultured Hyphomicrobiales bacteriu | SAMEA6954897 | aquatic | 4 | Yes | 3 | RNase_PH;Ham1p_like | — |
| 458233.SAMD00060930.AP009484_36 | **typeXI** | 258,8 | high | Bacillota | Bacilli | Bacillales | Staphylococcaceae | Macrococcoides | Macrococcoides caseolyticum | SAMD00060930 | host-associated | 11 | Yes | 4 | SLFN-g3_helicase;MazG-like;Metall | — |
| 1660160.SAMN03652429.MEDJ010000 | **typeXI** | 257,5 | high | Acidobacteriota | — | — | — | — | Acidobacteria bacterium SCN 69-37 | SAMN03652429 | anthropogenic | 8 | Yes | 4 | IstB_IS21;HEPN_Swt1;Helicase_C | — |
| 2795124.SAMN17004917.JAELUI0100( | **typeXI** | 253,3 | high | Pseudomonadota | Betaproteobacteria | Burkholderiales | Burkholderiaceae | Ralstonia | Ralstonia sp. ASV6 | SAMN17004917 | host-associated | 9 | Yes | 4 | AAA;Hydrolase;HTH_Tnp_1;TnpB_ | — |
| 2282150.SAMN09639277.DSIT0100009 | **typeXI** | 253,2 | high | Pseudomonadota | Alphaproteobacteria | Caulobacterales | — | — | Caulobacterales bacterium | SAMN09639277 | temperature | 11 | Yes | 4 | TetR_C_28;UvrB_inter;TbpB_B_D | — |
| 1961564.SAMN06450404.DING010000 | **typeXI** | 253,2 | high | Thermodesulfobacteriota | Desulfovibrionia | Desulfovibrionales | Desulfovibrionaceae | — | Desulfovibrionaceae bacterium UBA5 | SAMN06450404 | terrestrial | 11 | Yes | 4 | 4HBT;SIS;GATase_7;CPSase_L_D | — |
| 1345023.SAMN02471612.ATCL010000 | **typeXI** | 251,3 | high | Bacillota | Bacilli | Bacillales | — | Exiguobacterium | Exiguobacterium chiriqhucha | SAMN02471612 | host-associated | 11 | Yes | 4 | Methyltrans_RNA;DUF4435;Peptid | — |
| 152509.SAMEA6152703.CADCEC0100 | **typeXI** | 249,2 | high | Bacteroidota | — | — | — | — | uncultured Bacteroidota bacterium | SAMEA6152703 | host-associated | 11 | Yes | 4 | iREC;XendoU;WYL | — |
| 1529005.SAMN04075706.LJSC010000 | **typeXI** | 249,2 | high | Pseudomonadota | Alphaproteobacteria | Rhodobacterales | Paracoccaceae | Gemmobacter | Gemmobacter sp. LW-1 | SAMN04075706 | anthropogenic | 11 | Yes | 4 | Exonuc_VII_L;Arm-DNA-bind_3;Bs | — |
| 1916129.SAMN05959503.MPMM01000 | **typeXI** | 247,4 | high | Chloroflexota | Dehalococcoidia | — | — | — | SAR202 cluster bacterium bin22 | SAMN05959503 | host-associated | 11 | Yes | 4 | FRG | — |
| 1895921.SAMN05660545.MKTC010000( | **typeXI** | 244 | high | Bacteroidota | — | — | — | — | Bacteroidetes bacterium 37-13 | SAMN05660545 | anthropogenic | 8 | Yes | 4 | KilA-N;HEPN_MAE_28990;AAA_3 | — |

| | | | | | | | | | | | | | | | | |
|---|---|---|---|---|---|---|---|---|---|---|---|---|---|---|---|---|
| 370804.SAMEA6150200.CACYME0100 | typeXI | 242 | high | Bacteroidota | Bacteroidia | Bacteroidales | Prevotellaceae | — | uncultured Prevotellaceae bacterium | SAMEA6150200 | host-associated | 11 | Yes | 4 | CGGC;HisKin-conflict;MTUS1_CCI | — |
| 2030927.SAMN16349853.JAGBQJ0100 | typeXI | 240,6 | high | Bacteroidota | Bacteroidia | Bacteroidales | — | — | Bacteroidales bacterium | SAMN16349853 | host-associated | 8 | Yes | 4 | WYL;YhcG_C;WG_beta_rep | — |
| 196367.SAMN06075480.MTHB0100004 | typeXI | 240,6 | high | Pseudomonadota | Betaproteobacteria | Burkholderiales | Burkholderiaceae | Caballeronia | Caballeronia sordidicola | SAMN06075480 | terrestrial | 7 | Yes | 3 | DUF6933;Trypsin_2;Phage_integra | — |
| 2030927.SAMN16347308.JAFXXO0100 | typeXI | 239,9 | high | Bacteroidota | Bacteroidia | Bacteroidales | — | — | Bacteroidales bacterium | SAMN16347308 | host-associated | 11 | Yes | 4 | WYL;HisKin-conflict;Tenui_NCP | — |
| 286133.SAMEA5852096.CABMQD0100 | typeXI | 239,5 | high | Pseudomonadota | Betaproteobacteria | Burkholderiales | Sutterellaceae | Sutterella | uncultured Sutterella sp. | SAMEA5852096 | host-associated | 11 | Yes | 4 | MrcB_N;ATPase_2;DUF927 | — |
| 1129257.SAMEA6152766.CADCGK010 | typeXI | 237,8 | high | Bacteroidota | Bacteroidia | — | — | — | uncultured Bacteroidia bacterium | SAMEA6152766 | host-associated | 11 | Yes | 4 | HD;SACS | — |
| 2030927.SAMN16349994.JAGBVU0100 | typeXI | 237,3 | high | Bacteroidota | Bacteroidia | Bacteroidales | — | — | Bacteroidales bacterium | SAMN16349994 | host-associated | 11 | Yes | 4 | GerT_N;DUF7002;Beta-prop_CAF1 | — |
| 1916129.SAMN05959503.MPMM01000 | typeXI | 236,1 | high | Chloroflexota | Dehalococcoidia | — | — | — | SAR202 cluster bacterium bin22 | SAMN05959503 | host-associated | 11 | Yes | 4 | Phage_integrase;HNH;FRG;DUF69 | — |
| 2748031.SAMN16348336.JAFZLC0100 | typeXI | 235,8 | high | Bacteroidota | Bacteroidia | Bacteroidales | Paludibacteraceae | — | Paludibacteraceae bacterium | SAMN16348336 | host-associated | 8 | Yes | 4 | Pribosyltran;DNA_processg_A;WHI | — |
| 152509.SAMEA6150143.CACYJU0100 | typeXI | 235,6 | high | Bacteroidota | — | — | — | — | uncultured Bacteroidota bacterium | SAMEA6150143 | host-associated | 9 | Yes | 4 | DnaB_C;T1-F;DUF2846;WYL | — |
| 2030927.SAMN16347413.JAFYBP0100 | typeXI | 235,2 | high | Bacteroidota | Bacteroidia | Bacteroidales | — | — | Bacteroidales bacterium | SAMN16347413 | host-associated | 9 | Yes | 4 | Sulfotransfer_1;DnaB_C | — |
| 59823.SAMN19224964.DXTA01000038 | typeXI | 235 | high | Bacteroidota | Bacteroidia | Bacteroidales | Prevotellaceae | Prevotella | Prevotella sp. | SAMN19224964 | host-associated | 8 | Yes | 4 | iREC;HisKin-conflict;SufE | — |
| 152509.SAMEA6151326.CADADB0100 | typeXI | 234,7 | high | Bacteroidota | — | — | — | — | uncultured Bacteroidota bacterium | SAMEA6151326 | host-associated | 11 | Yes | 4 | GVIN1_C;WYL;T1-F;Cyano_EgtBD | — |
| 2026756.SAMN11294391.SUWU01000 | typeXI | 234,6 | high | Bacteroidota | Bacteroidia | Bacteroidales | Lentimicrobiaceae | — | Lentimicrobiaceae bacterium | SAMN11294391 | host-associated | 10 | Yes | 4 | WYL;Asp_protease_2;T1-F | — |
| 194843.SAMEA6150145.CACYJZ0100 | typeXI | 233,4 | high | Bacteroidota | Bacteroidia | Bacteroidales | — | — | uncultured Bacteroidales bacterium | SAMEA6150145 | host-associated | 11 | Yes | 4 | WYL;HisKin-conflict;WHD_eIF2D | — |
| 754429.SAMN06133352.CP019333_12 | typeXI | 232,7 | low | Bacteroidota | Flavobacteriia | Flavobacteriales | Flavobacteriaceae | Gilvibacter | Gilvibacter sp. SZ-19 | SAMN06133352 | temperature | 10 | Yes | 4 | AAA_30;Seryl_tRNA_N;MRP-S26 | — |
| 59823.SAMN11294429.SUYG01000053 | typeXI | 232,6 | low | Bacteroidota | Bacteroidia | Bacteroidales | Prevotellaceae | Prevotella | Prevotella sp. | SAMN11294429 | host-associated | 10 | Yes | 4 | Plasmid_parti_N;CGGC;T1-F;EMY | — |
| 2030927.SAMN16341878.JAFPVK0100 | typeXI | 232,6 | low | Bacteroidota | Bacteroidia | Bacteroidales | — | — | Bacteroidales bacterium | SAMN16341878 | host-associated | 10 | Yes | 4 | iREC;WHD_eIF2D;WG_beta_rep | — |
| 2049048.SAMN16344483.JAFTOX0100 | typeXI | 231,9 | low | Bacteroidota | Bacteroidia | Bacteroidales | Rikenellaceae | — | Rikenellaceae bacterium | SAMN16344483 | host-associated | 10 | Yes | 4 | AAA_30;WYL;Seryl_tRNA_N | — |
| 2030927.SAMN16343841.JAFSQF0100 | typeXI | 231,2 | low | Bacteroidota | Bacteroidia | Bacteroidales | — | — | Bacteroidales bacterium | SAMN16343841 | host-associated | 10 | Yes | 4 | DnaJ;DnaB_C;Bflower_2 | — |
| 2030927.SAMN19225073.JAHZEX0100 | typeXI | 230,9 | low | Bacteroidota | Bacteroidia | Bacteroidales | — | — | Bacteroidales bacterium | SAMN19225073 | host-associated | 10 | Yes | 4 | Ykzl;Seryl_tRNA_N;WYL;TM225 | — |
| 59823.SAMN16343164.JAFRQB01000 | typeXI | 230,8 | low | Bacteroidota | Bacteroidia | Bacteroidales | Prevotellaceae | Prevotella | Prevotella sp. | SAMN16343164 | host-associated | 10 | Yes | 4 | PLDc_2;CGGC;AAA_30 | — |
| 1236515.SAMD00010076.BAKM010000 | typeXI | 230,7 | low | Bacteroidota | Bacteroidia | Bacteroidales | Bacteroidaceae | Phocaeicola | Phocaeicola sartorii | SAMD00010076 | host-associated | 10 | Yes | 4 | DUF1972;YjbH;MTUS1_CCDC69_ | — |
| 1129257.SAMEA6152677.CADCCZ010 | typeXI | 229,7 | low | Bacteroidota | Bacteroidia | — | — | — | uncultured Bacteroidia bacterium | SAMEA6152677 | host-associated | 10 | Yes | 4 | iREC;MTUS1_CCDC69_CC1;Asp_ | — |
| 54062.SAMD00166260.BJWE01000002 | typeXI | 229,5 | low | Bacillota | Bacilli | Lactobacillales | Lactobacillaceae | Pediococcus | Pediococcus parvulus | SAMD00166260 | anthropogenic | 10 | Yes | 4 | rve;HTH_28;VanZ;DUF2715 | — |
| 370804.SAMEA6151525.CADAKX0100 | typeXI | 229,1 | low | Bacteroidota | Bacteroidia | Bacteroidales | Prevotellaceae | — | uncultured Prevotellaceae bacterium | SAMEA6151525 | host-associated | 7 | Yes | 4 | CGGC;PLDc_2 | — |
| 194843.SAMEA6150768.CACZHT0100 | typeXI | 228,9 | low | Bacteroidota | Bacteroidia | Bacteroidales | — | — | uncultured Bacteroidales bacterium | SAMEA6150768 | host-associated | 10 | Yes | 4 | Cytochrome_C7;WG_beta_rep;WY | — |
| 1952463.SAMN06456358.DDQG01000 | typeXI | 228,9 | low | Bacteroidota | Bacteroidia | Bacteroidales | Prevotellaceae | — | Prevotellaceae bacterium UBA2448 | SAMN06456358 | host-associated | 10 | Yes | 4 | AAA_30;DUF7002;DUF5496;YhcG | — |
| 370804.SAMEA6150222.CACYNE0100 | typeXI | 228,8 | low | Bacteroidota | Bacteroidia | Bacteroidales | Prevotellaceae | — | uncultured Prevotellaceae bacterium | SAMEA6150222 | host-associated | 10 | Yes | 4 | SufE;HisKin-conflict;iREC | — |
| 1704277.SAMN04002371.LIYN0200000 | typeXI | 228,8 | low | Deinococcota | Deinococci | Deinococcales | Deinococcaceae | Deinococcus | Deinococcus sp. UR1 | SAMN04002371 | terrestrial | 10 | Yes | 4 | AIRS_C;GATase_7;DUF2196 | — |
| 370804.SAMEA6149422.CACXIP0100 | typeXI | 228,6 | low | Bacteroidota | Bacteroidia | Bacteroidales | Prevotellaceae | — | uncultured Prevotellaceae bacterium | SAMEA6149422 | host-associated | 10 | Yes | 4 | Beta-prop_IFT122_1st;WYL;HisKin | — |
| 2079531.SAMEA104588899.LT985324_ | typeXI | 228,3 | low | Bacteroidota | Bacteroidia | Bacteroidales | Prevotellaceae | Prevotella | Prevotella merdae | SAMEA104588899 | host-associated | 10 | Yes | 4 | PIN;iREC;Ribosomal_L27 | — |
| 370804.SAMEA6152573.CADBYY0100 | typeXI | 228 | low | Bacteroidota | Bacteroidia | Bacteroidales | Prevotellaceae | — | uncultured Prevotellaceae bacterium | SAMEA6152573 | host-associated | 10 | Yes | 4 | AAA_30;PLDc_2 | — |
| 1129257.SAMEA6151604.CADANY010 | typeXI | 227,8 | low | Bacteroidota | Bacteroidia | — | — | — | uncultured Bacteroidia bacterium | SAMEA6151604 | host-associated | 9 | Yes | 4 | MreB_Mbl;Orthopox_A43R;DUF42 | — |
| 2030927.SAMN16347328.JAFXYI0100 | typeXI | 227,6 | low | Bacteroidota | Bacteroidia | Bacteroidales | — | — | Bacteroidales bacterium | SAMN16347328 | host-associated | 10 | Yes | 4 | iREC;HisKin-conflict;WYL;HATPas | — |
| 1929085.SAMN16350086.JAGBZI0100 | typeXI | 227,3 | low | Bacteroidota | Bacteroidia | Bacteroidales | Rikenellaceae | Tidjanibacter | Tidjanibacter sp. | SAMN16350086 | host-associated | 6 | Yes | 3 | LRR_5;FTHFS;CE2_N | — |
| 159272.SAMEA7847924.CAJMCE0100 | typeXI | 227 | low | Bacteroidota | Bacteroidia | Bacteroidales | Prevotellaceae | Prevotella | uncultured Prevotella sp. | SAMEA7847924 | host-associated | 8 | Yes | 4 | T1-F;WYL | — |
| 1129257.SAMEA6148842.CACWMH01 | typeXI | 226,5 | low | Bacteroidota | Bacteroidia | — | — | — | uncultured Bacteroidia bacterium | SAMEA6148842 | host-associated | 10 | Yes | 4 | AAA_11;T1-F;WG_beta_rep;WYL | — |
| 194843.SAMEA6151427.CADAGY0100 | typeXI | 226,3 | low | Bacteroidota | Bacteroidia | Bacteroidales | — | — | uncultured Bacteroidales bacterium | SAMEA6151427 | host-associated | 10 | Yes | 4 | MTUS1_CCDC69_CC1;HisKin-cor | — |
| 1929085.SAMN16344624.JAFTUI0100 | typeXI | 226,1 | low | Bacteroidota | Bacteroidia | Bacteroidales | Rikenellaceae | Tidjanibacter | Tidjanibacter sp. | SAMN16344624 | host-associated | 10 | Yes | 4 | DUF6261;T1-F;MIT_C;AAA_15 | — |
| 370804.SAMEA6150912.CACZNL0100 | typeXI | 226,1 | low | Bacteroidota | Bacteroidia | Bacteroidales | Prevotellaceae | — | uncultured Prevotellaceae bacterium | SAMEA6150912 | host-associated | 10 | Yes | 0 | — | — |
| 2212467.SAMN16349935.JAGBTN0100 | typeXI | 226 | low | Bacteroidota | Bacteroidia | Bacteroidales | Bacteroidaceae | — | Bacteroidaceae bacterium | SAMN16349935 | host-associated | 10 | Yes | 4 | WG_beta_rep;T1-F | — |
| 194843.SAMEA6149915.CACYBD0100 | typeXI | 225,7 | low | Bacteroidota | Bacteroidia | Bacteroidales | — | — | uncultured Bacteroidales bacterium | SAMEA6149915 | host-associated | 10 | Yes | 4 | PLDc_2;CGGC | — |
| 2486470.SAMN10365287.RIBI0100000 | typeXI | 225,6 | low | Bacteroidota | Bacteroidia | Bacteroidales | Muribaculaceae | — | Muribaculaceae bacterium Isolate-102 | SAMN10365287 | host-associated | 10 | Yes | 4 | WYL;T1-F;HAD_2 | — |
| 2030927.SAMN13893983.JAAYJQ0100 | typeXI | 225,3 | low | Bacteroidota | Bacteroidia | Bacteroidales | — | — | Bacteroidales bacterium | SAMN13893983 | temperature | 10 | Yes | 4 | WYL;T1-F;DUF3695 | — |
| 194843.SAMEA6150745.CACZHD0100 | typeXI | 224,7 | low | Bacteroidota | Bacteroidia | Bacteroidales | — | — | uncultured Bacteroidales bacterium | SAMEA6150745 | host-associated | 10 | Yes | 4 | WYL;HisKin-conflict;iREC | — |
| 2212467.SAMN16346420.JAFWPN010 | typeXI | 224,7 | low | Bacteroidota | Bacteroidia | Bacteroidales | Bacteroidaceae | — | Bacteroidaceae bacterium | SAMN16346420 | host-associated | 10 | Yes | 4 | YhcG_C;Plasmid_parti_N;DUF7002 | — |
| 2301481.SAMEA8805426.CAJUAJ0100 | typeXI | 224,6 | low | Bacteroidota | Bacteroidia | Bacteroidales | Muribaculaceae | — | uncultured Muribaculaceae bacterium | SAMEA8805426 | host-associated | 10 | Yes | 4 | BAR_4;HisKin-conflict;Por_Secre_t | — |
| 194843.SAMEA6150050.CACYGD0100 | typeXI | 224,3 | low | Bacteroidota | Bacteroidia | Bacteroidales | — | — | uncultured Bacteroidales bacterium | SAMEA6150050 | host-associated | 10 | Yes | 4 | WYL;WG_beta_rep;T1-F;LRR_5 | — |
| 194843.SAMEA6150348.CACYRT0100 | typeXI | 224,3 | low | Bacteroidota | Bacteroidia | Bacteroidales | — | — | uncultured Bacteroidales bacterium | SAMEA6150348 | host-associated | 10 | Yes | 4 | iREC;HisKin-conflict;WYL;GP57 | — |
| 2133944.SAMN08763369.CP033459_2 | typeXI | 224 | low | Bacteroidota | Bacteroidia | Bacteroidales | Prevotellaceae | Pseudoprevotella | Pseudoprevotella muciniphila | SAMN08763369 | host-associated | 10 | Yes | 4 | HisKin-conflict;iREC | — |
| 2212467.SAMN16243600.JAEELS0100 | typeXI | 223,4 | low | Bacteroidota | Bacteroidia | Bacteroidales | Bacteroidaceae | — | Bacteroidaceae bacterium | SAMN16243600 | host-associated | 10 | Yes | 4 | DUF5496;CGGC;DUF1848 | — |
| 370804.SAMEA6150573.CACZAB0100 | typeXI | 223,2 | low | Bacteroidota | Bacteroidia | Bacteroidales | Prevotellaceae | — | uncultured Prevotellaceae bacterium | SAMEA6150573 | host-associated | 10 | Yes | 4 | UvrD-helicase;LRR_5 | — |
| 2030927.SAMN16243923.JAEEXD0100 | typeXI | 223,1 | low | Bacteroidota | Bacteroidia | Bacteroidales | — | — | Bacteroidales bacterium | SAMN16243923 | host-associated | 10 | Yes | 4 | SLFN_AlbA_2;DNA_processg_A | — |
| 194843.SAMEA6152128.CADBHU0100 | typeXI | 223,1 | low | Bacteroidota | Bacteroidia | Bacteroidales | — | — | uncultured Bacteroidales bacterium | SAMEA6152128 | host-associated | 10 | Yes | 4 | AMP_deaminase | — |
| 370804.SAMEA6151246.CADAAG0100 | typeXI | 222,2 | low | Bacteroidota | Bacteroidia | Bacteroidales | Prevotellaceae | — | uncultured Prevotellaceae bacterium | SAMEA6151246 | host-associated | 10 | Yes | 4 | iREC;HisKin-conflict;WYL;DUF701 | — |
| 152509.SAMEA6954497.CAINMV0100 | typeXI | 222,1 | low | Bacteroidota | — | — | — | — | uncultured Bacteroidota bacterium | SAMEA6954497 | temperature | 10 | Yes | 4 | Por_Secre_tail;DnaB_C;VWA_2 | — |
| 2498093.SAMN16342757.JAFQZK0100 | typeXI | 222 | low | Bacteroidota | Bacteroidia | Bacteroidales | Muribaculaceae | — | Muribaculaceae bacterium | SAMN16342757 | host-associated | 10 | Yes | 4 | WYL;T1-F;HAD_2 | — |
| 152509.SAMEA6954080.CAIWCG0100 | typeXI | 221,8 | low | Bacteroidota | — | — | — | — | uncultured Bacteroidota bacterium | SAMEA6954080 | aquatic | 10 | Yes | 4 | UvrD_C;VWA_2;DnaB_C;AAA | — |
| 2033407.SAMN14407237.JAAVFD0100 | typeXI | 221,6 | low | Bacteroidota | Bacteroidia | Bacteroidales | Barnesiellaceae | Barnesiella | Barnesiella sp. | SAMN14407237 | host-associated | 10 | Yes | 4 | BAR_4;HisKin-conflict;Glyco_hydro | — |
| 2212467.SAMN16243652.JAEENS0100 | typeXI | 221,5 | low | Bacteroidota | Bacteroidia | Bacteroidales | Bacteroidaceae | — | Bacteroidaceae bacterium | SAMN16243652 | host-associated | 10 | Yes | 4 | YhcG_C;Beta-prop_IFT122_1st;DU | — |
| 152509.SAMEA6953637.CAIUSH0100 | typeXI | 221,5 | low | Bacteroidota | — | — | — | — | uncultured Bacteroidota bacterium | SAMEA6953637 | aquatic | 3 | Yes | 3 | iREC;HisKin-conflict;WYL | — |
| 370804.SAMEA104667055.ONMA0100 | typeXI | 221,5 | low | Bacteroidota | Bacteroidia | Bacteroidales | Prevotellaceae | — | uncultured Prevotellaceae bacterium | SAMEA104667055 | host-associated | 10 | Yes | 4 | WYL;WG_beta_rep;HisKin-conflict; | — |
| 1841864.SAMN14913643.JACJJG0100 | typeXI | 221,3 | low | Bacteroidota | Bacteroidia | Bacteroidales | Prevotellaceae | Marseilla | Marseilla massiliensis | SAMN14913643 | host-associated | 10 | Yes | 4 | WYL;WG_beta_rep;Seryl_tRNA_N; | — |
| 2030927.SAMN16346686.JAFWZP0100 | typeXI | 221,2 | low | Bacteroidota | Bacteroidia | Bacteroidales | — | — | Bacteroidales bacterium | SAMN16346686 | host-associated | 8 | Yes | 4 | WG_beta_rep;T1-F;TPR_7 | — |
| 194843.SAMEA6152634.CADCBN0100 | typeXI | 221 | low | Bacteroidota | Bacteroidia | Bacteroidales | — | — | uncultured Bacteroidales bacterium | SAMEA6152634 | host-associated | 10 | Yes | 4 | CarbopepD_reg_2;DUF7002;DuOB | — |
| 194843.SAMEA6150745.CACZHD0100 | typeXI | 220,5 | low | Bacteroidota | Bacteroidia | Bacteroidales | — | — | uncultured Bacteroidales bacterium | SAMEA6150745 | host-associated | 10 | Yes | 4 | Lprl;WYL | — |
| 2030927.SAMEA5279175.CAAFXO010 | typeXI | 220,4 | low | Bacteroidota | Bacteroidia | Bacteroidales | — | — | Bacteroidales bacterium | SAMEA5279175 | host-associated | 10 | Yes | 4 | HisKin-conflict;iREC | — |
| 159274.SAMEA5278472.CAAEQB0100 | typeXI | 220,3 | low | Bacteroidota | Bacteroidia | Bacteroidales | Porphyromonadaceae | Porphyromonas | uncultured Porphyromonas sp. | SAMEA5278472 | host-associated | 7 | Yes | 4 | PepSY_like;WYL | — |
| 159272.SAMEA104666937.OMZM0100 | typeXI | 220,3 | low | Bacteroidota | Bacteroidia | Bacteroidales | Prevotellaceae | Prevotella | uncultured Prevotella sp. | SAMEA104666937 | host-associated | 10 | Yes | 4 | WG_beta_rep;Pox_E6;AA_kinase | — |
| 348578.SAMEA7847109.CAJLTU0100 | typeXI | 220,3 | low | Bacteroidota | Bacteroidia | Bacteroidales | Porphyromonadaceae | — | uncultured Porphyromonadaceae bact | SAMEA7847109 | host-associated | 10 | Yes | 4 | ResIII;HTH_17;HisKin-conflict;BAR | — |
| 1898104.SAMN16426341.JADKCP0100 | typeXI | 220,2 | low | Bacteroidota | — | — | — | — | Bacteroidota bacterium | SAMN16426341 | anthropogenic | 10 | Yes | 4 | HTH_31;DnaB_C | — |
| 370804.SAMEA6151724.CADASP0100 | typeXI | 220,1 | low | Bacteroidota | Bacteroidia | Bacteroidales | Prevotellaceae | — | uncultured Prevotellaceae bacterium | SAMEA6151724 | host-associated | 5 | Yes | 2 | LRR_5 | — |
| 348578.SAMEA4891951.URTA0100002 | typeXI | 219,6 | low | Bacteroidota | Bacteroidia | Bacteroidales | Porphyromonadaceae | — | uncultured Porphyromonadaceae bact | SAMEA4891951 | host-associated | 10 | Yes | 4 | Hydrolase_4;Seryl_tRNA_N;WG_b | — |
| 2301481.SAMEA8805709.CAJULY0100 | typeXI | 219,4 | low | Bacteroidota | Bacteroidia | Bacteroidales | Muribaculaceae | — | uncultured Muribaculaceae bacterium | SAMEA8805709 | host-associated | 10 | Yes | 4 | BAR_4;HisKin-conflict;HTH_17;DU | — |
| 2030927.SAMN16344869.JAFUDT0100 | typeXI | 219,4 | low | Bacteroidota | Bacteroidia | Bacteroidales | — | — | Bacteroidales bacterium | SAMN16344869 | host-associated | 10 | Yes | 4 | CdsD_C;WYL;T1-F | — |
| 1872444.SAMN16349454.JAGBAX0100 | typeXI | 219,4 | low | Bacteroidota | Bacteroidia | Bacteroidales | Rikenellaceae | Alistipes | Alistipes sp. | SAMN16349454 | host-associated | 10 | Yes | 4 | WYL;T1-F | — |
| 159272.SAMEA7848212.CAJMJX0100 | typeXI | 219,3 | low | Bacteroidota | Bacteroidia | Bacteroidales | Prevotellaceae | Prevotella | uncultured Prevotella sp. | SAMEA7848212 | host-associated | 10 | Yes | 4 | WYL;WG_beta_rep;Aconitase_C | — |
| 194843.SAMEA6150745.CACZHD0100 | typeXI | 219,3 | low | Bacteroidota | Bacteroidia | Bacteroidales | — | — | uncultured Bacteroidales bacterium | SAMEA6150745 | host-associated | 10 | Yes | 4 | WYL;Baculo_p74 | — |

| | | | | | | | | | | | | | | | | |
|---|---|---|---|---|---|---|---|---|---|---|---|---|---|---|---|---|
| 2498093.SAMN16349242.JAGAST0100 | **typeXI** | 219,1 | low | Bacteroidota | Bacteroidia | Bacteroidales | Muribaculaceae | — | Muribaculaceae bacterium | SAMN16349242 | host-associated | 10 | Yes | 4 | WYL | — |
| 2840839.SAMN15816954.JADIMG0100 | **typeXI** | 218,6 | low | Bacteroidota | Bacteroidia | Bacteroidales | — | Candidatus Gallipaludi | Candidatus Gallipaludibacter merdaviu | SAMN15816954 | host-associated | 10 | Yes | 4 | LysM | — |
| 2053307.SAMN16091041.JADQBJ0100 | **typeXI** | 218,5 | low | Thermodesulfobacteriota | Desulfobulbia | Desulfobulbales | Desulfobulbaceae | — | Desulfobulbaceae bacterium | SAMN16091041 | aquatic | 10 | Yes | 4 | SmpB;Phage_int_M;Response_reg | — |
| 2030927.SAMN18120043.JAGOLH0100 | **typeXI** | 218,5 | low | Bacteroidota | Bacteroidia | Bacteroidales | — | — | Bacteroidales bacterium | SAMN18120043 | anthropogenic | 8 | Yes | 4 | Glyco_hydro_25;DnaB_C;Por_Secr | — |
| 194843.SAMEA6150559.CACYZT0100 | **typeXI** | 218,3 | low | Bacteroidota | Bacteroidia | Bacteroidales | — | — | uncultured Bacteroidales bacterium | SAMEA6150559 | host-associated | 10 | Yes | 4 | DUF488;DuOB;T1-F;Feld-I_B | — |
| 1898104.SAMN17573936.JAFEAA0100 | **typeXI** | 218,2 | low | Bacteroidota | — | — | — | — | Bacteroidota bacterium | SAMN17573936 | anthropogenic | 10 | Yes | 4 | WYL;DUF3127 | — |
| 361581.SAMN08779091.RCCS010000 | **typeXI** | 218,2 | low | Bacteroidota | Flavobacteriia | Flavobacteriales | Flavobacteriaceae | Tenacibaculum | Tenacibaculum discolor | SAMN08779091 | aquatic | 10 | Yes | 4 | HTH_18;NHL;Leu_Phe_trans;SCF | — |
| 246.SAMN10316127.CP033935_66 | **typeXI** | 218,1 | low | Bacteroidota | Flavobacteriia | Flavobacteriales | Weeksellaceae | Chryseobacterium | Chryseobacterium balustinum | SAMN10316127 | aquatic | 10 | Yes | 4 | BAR_4;HisKin-conflict;ResIII;VapC | — |
| 159272.SAMEA7846766.CAJLGE0100 | **typeXI** | 217,6 | low | Bacteroidota | Bacteroidia | Bacteroidales | Prevotellaceae | Prevotella | uncultured Prevotella sp. | SAMEA7846766 | host-associated | 6 | Yes | 3 | WYL;WG_beta_rep;Seryl_tRNA_N | — |
| 2678657.SAMN13336490.WOFO01000 | **typeXI** | 216,8 | low | Bacteroidota | Cytophagia | Cytophagales | Cyclobacteriaceae | Cyclobacterium | Cyclobacterium sp. SYSU L10401 | SAMN13336490 | salinity | 10 | Yes | 4 | iREC;HisKin-conflict;WYL;Peptidas | — |
| 152509.SAMEA6953983.CAINDA0100 | **typeXI** | 216,7 | low | Bacteroidota | — | — | — | — | uncultured Bacteroidota bacterium | SAMEA6953983 | aquatic | 10 | Yes | 4 | YjdM;Seryl_tRNA_N;E2-CBASS | — |
| 152509.SAMEA6955963.CAITNS01000 | **typeXI** | 216,6 | low | Bacteroidota | — | — | — | — | uncultured Bacteroidota bacterium | SAMEA6955963 | aquatic | 5 | Yes | 2 | IL15;WYL | — |
| 194843.SAMEA6951536.CAIYES01000 | **typeXI** | 216,5 | low | Bacteroidota | Bacteroidia | Bacteroidales | — | — | uncultured Bacteroidales bacterium | SAMEA6951536 | aquatic | 7 | Yes | 4 | Response_reg;DUF3127;WYL | — |
| 152509.SAMEA6951679.CAIKXD01000 | **typeXI** | 216,2 | low | Bacteroidota | — | — | — | — | uncultured Bacteroidota bacterium | SAMEA6951679 | aquatic | 3 | Yes | 2 | Methyltransf_23;2OG-FeII_Oxy_2 | — |
| 1284775.SAMN02850936.JRNC010001 | **typeXI** | 215,7 | low | Bacteroidota | Bacteroidia | Bacteroidales | Prevotellaceae | Prevotella | Prevotella sp. S7-1-8 | SAMN02850936 | host-associated | 8 | Yes | 4 | UvrD_C_2;SufE;Seryl_tRNA_N;SL | — |
| 1898104.SAMN20441989.JAIFNN0100 | **typeXI** | 215 | low | Bacteroidota | — | — | — | — | Bacteroidota bacterium | SAMN20441989 | aquatic | 5 | Yes | 2 | DnaB_C;Y1_Tnp | — |
| 2030927.SAMN18118926.JAGNBV0100 | **typeXI** | 215 | low | Bacteroidota | Bacteroidia | Bacteroidales | — | — | Bacteroidales bacterium | SAMN18118926 | anthropogenic | 7 | Yes | 4 | Claudin_TMEM179-179B;Peptidas | — |
| 194843.SAMEA6151813.CADAWB0100 | **typeXI** | 214,8 | low | Bacteroidota | Bacteroidia | Bacteroidales | — | — | uncultured Bacteroidales bacterium | SAMEA6151813 | host-associated | 10 | Yes | 4 | UvrD_C_2;SufE | — |
| 2448779.SAMN18120629.JAGPDS0100 | **typeXI** | 214,8 | low | Bacteroidota | Chitinophagia | Chitinophagales | — | — | Chitinophagales bacterium | SAMN18120629 | anthropogenic | 6 | Yes | 3 | DUF2325;WYL | — |
| 2202734.SAMN16426033.JADJRO0100 | **typeXI** | 214,8 | low | Bacteroidota | Saprospiria | Saprospirales | Saprospiraceae | — | Saprospiraceae bacterium | SAMN16426033 | anthropogenic | 10 | Yes | 4 | DUF7874;E2-CBASS;WYL | — |
| 213322.SAMEA6944250.CAJAII010000 | **typeXI** | 214,6 | low | Bacteroidota | Flavobacteriia | Flavobacteriales | — | — | uncultured Flavobacteriales bacterium | SAMEA6944250 | temperature | 10 | Yes | 4 | PHM7_cyt;DnaB_C;Hydrolase_3;Si | — |
| 212695.SAMEA6944788.CAIXMS0100 | **typeXI** | 214,3 | low | Bacteroidota | Flavobacteriia | — | — | — | uncultured Flavobacteriia bacterium | SAMEA6944788 | temperature | 10 | Yes | 4 | HTH_11;Malic_M;DnaB_C;DNA_pr | — |
| 2748031.SAMN16347797.JAFYQJ0100 | **typeXI** | 214 | low | Bacteroidota | Bacteroidia | Bacteroidales | Paludibacteraceae | — | Paludibacteraceae bacterium | SAMN16347797 | host-associated | 10 | Yes | 4 | HAD_2;Aminotran_1_2;cREC_REC | — |
| 2740583.SAMN15052362.JABUMY0100 | **typeXI** | 213,8 | low | Bacteroidota | Chitinophagia | Chitinophagales | Chitinophagaceae | Flavisolibacter | Flavisolibacter longurius | SAMN15052362 | host-associated | 6 | Yes | 3 | Response_reg;WYL | — |
| 152509.SAMEA6948151.CAJAEO0100 | **typeXI** | 213,8 | low | Bacteroidota | — | — | — | — | uncultured Bacteroidota bacterium | SAMEA6948151 | aquatic | 10 | Yes | 4 | iREC;HisKin-conflict;WYL;DUF671 | — |
| 2498093.SAMN16345734.JAFVPD0100 | **typeXI** | 213,7 | low | Bacteroidota | Bacteroidia | Bacteroidales | Muribaculaceae | — | Muribaculaceae bacterium | SAMN16345734 | host-associated | 10 | Yes | 4 | CstA;PHP;HTS;AAA_lid_6 | — |
| 159272.SAMEA8030440.CAJOPE0100 | **typeXI** | 213,6 | low | Bacteroidota | Bacteroidia | Bacteroidales | Prevotellaceae | Prevotella | uncultured Prevotella sp. | SAMEA8030440 | host-associated | 10 | Yes | 4 | DUF7632;Orf78;DUF8000;TPK_B1 | — |
| 667015.SAMN00713589.CP002530_21 | **typeXI** | 213,1 | low | Bacteroidota | Bacteroidia | Bacteroidales | Bacteroidaceae | Phocaeicola | Phocaeicola salanitronis | SAMN00713589 | host-associated | 10 | Yes | 4 | DNA_processg_A | — |
| 1122985.SAMN02441341.KB899220_5 | **typeXI** | 212,2 | low | Bacteroidota | Bacteroidia | Bacteroidales | Prevotellaceae | Hoylesella | Hoylesella loescheii | SAMN02441341 | host-associated | 10 | Yes | 4 | Exonuc_VII_L;Exonuc_VII_S;RuvA | — |
| 2030927.SAMN13893767.JAAYFR0100 | **typeXI** | 212,1 | low | Bacteroidota | Bacteroidia | Bacteroidales | — | — | Bacteroidales bacterium | SAMN13893767 | anthropogenic | 5 | Yes | 3 | WYL;DUF2325;iREC | — |
| 2044603.SAMN07768642.PEAZ010000 | **typeXI** | 212 | low | Bacteroidota | Sphingobacteriia | Sphingobacteriales | Sphingobacteriaceae | Sphingobacterium | Sphingobacterium sp. 1.A.4 | SAMN07768642 | host-associated | 10 | Yes | 4 | HXXEE;WYL;Yop-YscD_ppl_3rd;A | — |
| 1898104.SAMN17573936.JAFEAA0100 | **typeXI** | 211,9 | low | Bacteroidota | — | — | — | — | Bacteroidota bacterium | SAMN17573936 | anthropogenic | 10 | Yes | 4 | Ribosomal_L17;DnaB_C;ArlS_N;D | — |
| 702447.SAMN00007489.ADKP0100012 | **typeXI** | 211,8 | low | Bacteroidota | Bacteroidia | Bacteroidales | Bacteroidaceae | Bacteroides | Bacteroides xylanisolvens | SAMN00007489 | host-associated | 10 | Yes | 4 | iREC;PucR;Phage_integrase;NusG | — |
| 1852808.SAMN05004131.MWCX01000 | **typeXI** | 211,7 | low | Bacteroidota | — | — | — | — | Bacteroidetes bacterium ADurb.Bin17 | SAMN05004131 | — | 5 | Yes | 0 | — | — |
| 1898104.SAMN10966580.VGMK01000 | **typeXI** | 211,6 | low | Bacteroidota | — | — | — | — | Bacteroidota bacterium | SAMN10966580 | aquatic | 6 | Yes | 3 | DUF7874;DUF6717 | — |
| 159272.SAMEA8030157.CAJOEC0100 | **typeXI** | 211,3 | low | Bacteroidota | Bacteroidia | Bacteroidales | Prevotellaceae | Prevotella | uncultured Prevotella sp. | SAMEA8030157 | host-associated | 10 | Yes | 4 | WDR55;Seryl_tRNA_N | — |
| 162156.SAMEA7847592.CAJLYR0100 | **typeXI** | 210,7 | low | Bacteroidota | Bacteroidia | Bacteroidales | Bacteroidaceae | Bacteroides | uncultured Bacteroides sp. | SAMEA7847592 | host-associated | 10 | Yes | 4 | Tricorn_N;DNA_processg_A | — |
| 213322.SAMEA6950112.CAJAVU0100 | **typeXI** | 210,5 | low | Bacteroidota | Flavobacteriia | Flavobacteriales | — | — | uncultured Flavobacteriales bacterium | SAMEA6950112 | aquatic | 9 | Yes | 4 | DAGAT;DnaB_C;Parathyroid;Lamin | — |
| 1797353.SAMN04313814.MEOQ01000 | **typeXI** | 210,4 | low | Bacteroidota | — | — | — | — | Bacteroidetes bacterium GWF2_43_6 | SAMN04313814 | terrestrial | 10 | Yes | 4 | DuOB;WYL;HisKin-conflict;WHD_e | — |
| 2030927.SAMN16056124.JAHDPK0100 | **typeXI** | 209,9 | low | Bacteroidota | Bacteroidia | Bacteroidales | — | — | Bacteroidales bacterium | SAMN16056124 | anthropogenic | 9 | Yes | 4 | HisKin-conflict;WYL;Transposase_r | — |
| 2301481.SAMEA7847604.CAJKVR0100 | **typeXI** | 209,8 | low | Bacteroidota | Bacteroidia | Bacteroidales | Muribaculaceae | — | uncultured Muribaculaceae bacterium | SAMEA7847604 | host-associated | 9 | Yes | 4 | tRNA-synt_2;Seryl_tRNA_N;DNA_ | — |
| 1423816.SAMD00036561.BACQ010000 | **typeXI** | 209,5 | low | Bacillota | Bacilli | Lactobacillales | Lactobacillaceae | Lacticaseibacillus | Lacticaseibacillus zeae | SAMD00036561 | host-associated | 10 | Yes | 4 | MMR_HSR1;Trypsin_2;DUF2568; | — |
| 152509.SAMEA6954493.CAIYNF01000 | **typeXI** | 209,5 | low | Bacteroidota | — | — | — | — | uncultured Bacteroidota bacterium | SAMEA6954493 | aquatic | 6 | Yes | 3 | Nramp;T2SS-T3SS_pil_N;DnaB_C | — |
| 1898104.SAMN16425514.JADIYW0100 | **typeXI** | 209,3 | low | Bacteroidota | — | — | — | — | Bacteroidota bacterium | SAMN16425514 | — | 6 | Yes | 0 | — | — |
| 2202734.SAMN16425904.JADJNH0100 | **typeXI** | 209,2 | low | Bacteroidota | Saprospiria | Saprospirales | Saprospiraceae | — | Saprospiraceae bacterium | SAMN16425904 | anthropogenic | 10 | Yes | 4 | iREC;HisKin-conflict;WYL;TerC | — |
| 2448779.SAMN18119813.JAGNZZ0100 | **typeXI** | 209,2 | low | Bacteroidota | Chitinophagia | Chitinophagales | — | — | Chitinophagales bacterium | SAMN18119813 | anthropogenic | 1 | Yes | 1 | DnaB_C | — |
| 1974047.SAMN06659620.PCUD01000 | **typeXI** | 208,9 | low | Ignavibacteriota | Ignavibacteria | Ignavibacteriales | — | — | Ignavibacteriales bacterium CG18_big | SAMN06659620 | terrestrial | 7 | Yes | 4 | DEAD;DUF2225;GIDA_C_1st | — |
| 2053306.SAMN09639373.DSML010000 | **typeXI** | 208,6 | low | Ignavibacteriota | Ignavibacteria | — | — | — | Ignavibacteria bacterium | SAMN09639373 | temperature | 10 | Yes | 4 | GHL10;WYL;POTRA_3;iREC | — |
| 880074.SAMN02769649.CP007034_23 | **typeXI** | 208,5 | low | Bacteroidota | Bacteroidia | Bacteroidales | Barnesiellaceae | Barnesiella | Barnesiella viscericola | SAMN02769649 | host-associated | 10 | Yes | 4 | DpnD-PcfM;HisKin-conflict;WHD_e | — |
| 194843.SAMEA6945217.CAIOAD0100 | **typeXI** | 208,4 | low | Bacteroidota | Bacteroidia | Bacteroidales | — | — | uncultured Bacteroidales bacterium | SAMEA6945217 | aquatic | 3 | Yes | 3 | DnaB_C | — |
| 575615.SAMN02595329.GG740078_36 | **typeXI** | 208,2 | low | Bacteroidota | Bacteroidia | Bacteroidales | Prevotellaceae | Prevotella | Prevotella sp. oral taxon 317 | SAMN02595329 | host-associated | 10 | Yes | 4 | NERD;DUF5895;RuvA_N;SprA_N | — |
| 1129257.SAMEA6950934.CAINLJ0100 | **typeXI** | 207,7 | low | Bacteroidota | Bacteroidia | — | — | — | uncultured Bacteroidia bacterium | SAMEA6950934 | aquatic | 10 | Yes | 4 | TnpB_IS66;Phage_T4_Y06J;WYL | — |
| 213322.SAMEA6951797.CAIUSK01000 | **typeXI** | 207,6 | low | Bacteroidota | Flavobacteriia | Flavobacteriales | — | — | uncultured Flavobacteriales bacterium | SAMEA6951797 | aquatic | 8 | Yes | 4 | TatA_B_E;Thiolase_N;DnaB_C;VW | — |
| 1972561.SAMEA5279589.CAAEQH010 | **typeXI** | 207,6 | low | Actinomycetota | Coriobacteriia | Eggerthellales | Eggerthellaceae | — | Eggerthellaceae bacterium | SAMEA5279589 | host-associated | 4 | Yes | 3 | SMC_N | — |
| 1965650.SAMN06473744.NFHT010000 | **typeXI** | 206,8 | low | Bacteroidota | Bacteroidia | Bacteroidales | Rikenellaceae | Alistipes | Alistipes sp. An66 | SAMN06473744 | host-associated | 10 | Yes | 4 | DUF6140;TPR_12;PLDc_2;Cons_ | — |
| 212695.SAMEA6951713.CAIYXD01000 | **typeXI** | 206,7 | low | Bacteroidota | Flavobacteriia | — | — | — | uncultured Flavobacteriia bacterium | SAMEA6951713 | aquatic | 8 | Yes | 4 | DnaB_C;Pol_alpha_B_N | — |
| 1129257.SAMEA6151141.CACZWF010 | **typeXI** | 205,8 | low | Bacteroidota | Bacteroidia | — | — | — | uncultured Bacteroidia bacterium | SAMEA6151141 | host-associated | 10 | Yes | 4 | DUF6242_C;PLDc_2;DUF4172;DU | — |
| 2838443.SAMN15816893.DWXH01000 | **typeXI** | 205,6 | low | Bacteroidota | Bacteroidia | Bacteroidales | Rikenellaceae | Alistipes | Candidatus Alistipes merdigallinarum | SAMN15816893 | host-associated | 10 | Yes | 4 | Mgm101p;TPR_12;DUF6140 | — |
| 1855373.SAMN05216518.FNBQ010000 | **typeXI** | 205,4 | low | Bacteroidota | Bacteroidia | Bacteroidales | — | — | Bacteroidales bacterium KHT7 | SAMN05216518 | host-associated | 10 | Yes | 4 | DAHP_synth_1;SIS;2-Hacid_dh_C | — |
| 2212467.SAMN16348663.JAFZWM010 | **typeXI** | 205,4 | low | Bacteroidota | Bacteroidia | Bacteroidales | Bacteroidaceae | — | Bacteroidaceae bacterium | SAMN16348663 | — | 10 | Yes | 0 | — | — |
| 2487074.SAMN10343198.RJUG010000 | **typeXI** | 205,3 | low | Bacteroidota | Flavobacteriia | Flavobacteriales | Weeksellaceae | Kaistella | Kaistella daneshvariae | SAMN10343198 | host-associated | 10 | Yes | 4 | Peptidase_G2;T1-F;Beta-prop_IFT1 | — |
| 1129257.SAMEA6153040.CADCRA010 | **typeXI** | 205,1 | low | Bacteroidota | Bacteroidia | — | — | — | uncultured Bacteroidia bacterium | SAMEA6153040 | host-associated | 5 | Yes | 4 | DUF87;DUF3920;RyR;LRR_5 | — |
| 475299.SAMN07621195.QKTX0100000 | **typeXI** | 205,1 | low | Bacteroidota | Cytophagia | Cytophagales | Cyclobacteriaceae | Algoriphagus | Algoriphagus aquaeductus | SAMN07621195 | aquatic | 10 | Yes | 4 | YhcG_C;T1-F;WYL;DUF4349 | — |
| 2044944.SAMN16426336.JADKCK0100 | **typeXI** | 204,6 | low | Bacteroidota | Sphingobacteriia | Sphingobacteriales | — | — | Sphingobacteriales bacterium | SAMN16426336 | anthropogenic | 10 | Yes | 4 | DUF4835;4HBT;HisKin-conflict | — |
| 2498093.SAMN12406360.VSPM01000 | **typeXI** | 204,1 | low | Bacteroidota | Bacteroidia | Bacteroidales | Muribaculaceae | — | Muribaculaceae bacterium | SAMN12406360 | host-associated | 7 | Yes | 4 | TPX2;WYL;UPF0227;bpMoxR | — |
| 2026799.SAMN17496206.JAFMKE0100 | **typeXI** | 204 | low | Verrucomicrobiota | — | — | — | — | Verrucomicrobiota bacterium | SAMN17496206 | aquatic | 6 | Yes | 3 | MORN_2;DnaB_C;ABC_tran | — |
| 2301481.SAMEA8805003.CAJTLZ0100 | **typeXI** | 203,5 | low | Bacteroidota | Bacteroidia | Bacteroidales | Muribaculaceae | — | uncultured Muribaculaceae bacterium | SAMEA8805003 | host-associated | 10 | Yes | 4 | Hexapep;WYL;UPF0227;bpMoxR | — |
| 1898104.SAMN16426506.JADKIU0100 | **typeXI** | 203,1 | low | Bacteroidota | — | — | — | — | Bacteroidota bacterium | SAMN16426506 | anthropogenic | 10 | Yes | 4 | TEBP_beta;DnaB_C;Transglut_cor | — |
| 1218108.SAMN02440982.KB908302_8 | **typeXI** | 202,5 | low | Bacteroidota | Flavobacteriia | Flavobacteriales | Weeksellaceae | Empedobacter | Empedobacter brevis | SAMN02440982 | anthropogenic | 10 | Yes | 4 | Phage_integrase;WYL;HisKin-confli | — |
| 1122991.SAMN02745169.QJJX010000 | **typeXI** | 200,3 | low | Bacteroidota | Bacteroidia | Bacteroidales | Prevotellaceae | Hoylesella | Hoylesella shahii | SAMN02745169 | host-associated | 10 | Yes | 4 | SprA_N;RuvA_N;Exonuc_VII_S;Ex | — |
| 28132.SAMD00078755.AP018049_14 | **typeXI** | 200,3 | low | Bacteroidota | Bacteroidia | Bacteroidales | Prevotellaceae | Prevotella | Prevotella melaninogenica | SAMD00078755 | host-associated | 10 | Yes | 4 | Lycopene_cyc;Sigma70_r2;Aminotr | — |
| 213322.SAMEA6956517.CAIQQJ01000 | **typeXI** | 199,7 | low | Bacteroidota | Flavobacteriia | Flavobacteriales | — | — | uncultured Flavobacteriales bacterium | SAMEA6956517 | temperature | 4 | Yes | 2 | ACBP4_C;DnaB_C | — |
| 2099680.SAMN19791121.JAIBBO0100 | **typeXI** | 199,7 | low | Methanobacteriota | Methanobacteria | Methanobacteriales | Methanobacteriaceae | — | Methanobacteriaceae archaeon | SAMN19791121 | anthropogenic | 5 | Yes | 2 | YxeD;BUD31 | — |
| 2030927.SAMEA5278599.CAAFAP0100 | **typeXI** | 199,2 | low | Bacteroidota | Bacteroidia | Bacteroidales | — | — | Bacteroidales bacterium | SAMEA5278599 | host-associated | 10 | Yes | 4 | Methyltransf_23;WYL;UPF0227;PD | — |
| 1869212.SAMN17497506.JAFEIJ01000 | **typeXI** | 199,1 | low | Bacteroidota | Chitinophagia | Chitinophagales | Chitinophagaceae | — | Chitinophagaceae bacterium | SAMN17497506 | aquatic | 10 | Yes | 4 | 2CompART;Methyltransf_23;DnaB_ | — |
| 213322.SAMEA8498464.CAJQJX0100 | **typeXI** | 198,6 | low | Bacteroidota | Flavobacteriia | Flavobacteriales | — | — | uncultured Flavobacteriales bacterium | SAMEA8498464 | aquatic | 10 | Yes | 4 | DUF2805;HpcH_HpaI;SpoU_methy | — |
| 370804.SAMEA6152226.CADBLU0100 | **typeXI** | 198,5 | low | Bacteroidota | Bacteroidia | Bacteroidales | Prevotellaceae | — | uncultured Prevotellaceae bacterium | SAMEA6152226 | host-associated | 5 | Yes | 2 | DUF2201 | — |
| 2486470.SAMN10365287.RIBI0100000 | **typeXI** | 197,8 | low | Bacteroidota | Bacteroidia | Bacteroidales | Muribaculaceae | — | Muribaculaceae bacterium Isolate-102 | SAMN10365287 | host-associated | 10 | Yes | 4 | DEAD;WYL;UPF0227 | — |
| 1236518.SAMN03897722.CP016204_3 | **typeXI** | 197,7 | low | Bacteroidota | Bacteroidia | Bacteroidales | Prevotellaceae | Prevotella | Prevotella scopos | SAMN03897722 | host-associated | 10 | Yes | 4 | Chorismate_bind;Aminotran_4;Sigm | — |

| | | | | | | | | | | | | | | | | |
|---|---|---|---|---|---|---|---|---|---|---|---|---|---|---|---|---|
| 563031.SAMN02463828.JH114149_60 | **typeXI** | 197 | low | Bacteroidota | Bacteroidia | Bacteroidales | Prevotellaceae | Prevotella | Prevotella sp. C561 | SAMN02463828 | host-associated | 10 | Yes | 4 | Lycopene_cyc;Sigma70_r2;Aminotr | — |
| 1898104.SAMN15870297.JACTMM010 | **typeXI** | 196,5 | low | Bacteroidota | — | — | — | — | Bacteroidota bacterium | SAMN15870297 | anthropogenic | 5 | Yes | 2 | WYL | — |
| 1945886.SAMN06298210.FZRE010000 | **typeXI** | 195,9 | low | Bacteroidota | Bacteroidia | Bacteroidales | Prevotellaceae | — | Prevotellaceae bacterium KH2P17 | SAMN06298210 | host-associated | 10 | Yes | 4 | Bac_DnaA;TonB_dep_Rec_b-barre | — |
| 1236517.SAMN03704035.CP012074_1 | **typeXI** | 195,6 | low | Bacteroidota | Bacteroidia | Bacteroidales | Prevotellaceae | Prevotella | Prevotella fusca | SAMN03704035 | — | 10 | Yes | 0 | — | — |
| 152509.SAMEA6951128.CAISCU01000 | **typeXI** | 195,6 | low | Bacteroidota | — | — | — | — | uncultured Bacteroidota bacterium | SAMEA6951128 | aquatic | 10 | Yes | 4 | Esterase;3-dmu-9_3-mt;DnaB_C;UI | — |
| 888743.SAMN00253307.GL872282_81 | **typeXI** | 195,3 | low | Bacteroidota | Bacteroidia | Bacteroidales | Prevotellaceae | Prevotella | Prevotella multiformis | SAMN00253307 | host-associated | 10 | Yes | 4 | Lipase_GDSL_2;LRR_8 | — |
| 348578.SAMEA4891951.URTA0100000 | **typeXI** | 194,9 | low | Bacteroidota | Bacteroidia | Bacteroidales | Porphyromonadaceae | — | uncultured Porphyromonadaceae bact | SAMEA4891951 | host-associated | 8 | Yes | 4 | dCMP_cyt_deam_1;WYL;Ribosoma | — |
| 28132.SAMN18352174.CP072361_100 | **typeXI** | 194,8 | low | Bacteroidota | Bacteroidia | Bacteroidales | Prevotellaceae | Prevotella | Prevotella melaninogenica | SAMN18352174 | host-associated | 10 | Yes | 4 | Chorismate_bind;Aminotran_4;Sigm | — |
| 2840691.SAMN15816958.DVIH010000 | **typeXI** | 194,8 | low | Bacteroidota | Bacteroidia | Bacteroidales | Bacteroidaceae | Candidatus Avibacteroi | Candidatus Avibacteroides excrementi | SAMN15816958 | — | 5 | Yes | 0 | — | — |
| 553174.SAMN00001916.CP002122_18 | **typeXI** | 194,1 | low | Bacteroidota | Bacteroidia | Bacteroidales | Prevotellaceae | Prevotella | Prevotella melaninogenica | SAMN00001916 | — | 10 | Yes | 0 | — | — |
| 1953166.SAMN06454853.DITN0100000 | **typeXI** | 194,1 | low | Bacteroidota | — | — | — | — | Bacteroidetes bacterium UBA6183 | SAMN06454853 | aquatic | 10 | Yes | 4 | PALP;Glucosaminidase;DnaB_C;CI | — |
| 767031.SAMN00031760.CP002589_44 | **typeXI** | 193,4 | low | Bacteroidota | Bacteroidia | Bacteroidales | Prevotellaceae | Prevotella | Prevotella denticola | SAMN00031760 | host-associated | 10 | Yes | 4 | Chorismate_bind;Aminotran_4;GT8 | — |
| 1913989.SAMN09215160.QNFE010000 | **typeXI** | 193 | low | Pseudomonadota | Gammaproteobacteria | — | — | — | Gammaproteobacteria bacterium | SAMN09215160 | temperature | 10 | Yes | 4 | DUF488;dnstrm_HI1420;Response | — |
| 2301481.SAMEA8805108.CAJTOZ0100 | **typeXI** | 191,4 | low | Bacteroidota | Bacteroidia | Bacteroidales | Muribaculaceae | — | uncultured Muribaculaceae bacterium | SAMEA8805108 | host-associated | 10 | Yes | 4 | DUF1453;PLDc_2;PM0109;IDEAL | — |
| 2768039.SAMEA8805111.CAJTQR0100 | **typeXI** | 191,3 | low | Bacteroidota | Bacteroidia | Bacteroidales | Muribaculaceae | Duncaniella | uncultured Duncaniella sp. | SAMEA8805111 | host-associated | 10 | Yes | 4 | GmrSD_N;DpnD-PcfM;PLDc_2 | — |
| 1872444.SAMN16342050.JAFQCA0100 | **typeXI** | 191,1 | low | Bacteroidota | Bacteroidia | Bacteroidales | Rikenellaceae | Alistipes | Alistipes sp. | SAMN16342050 | host-associated | 9 | Yes | 4 | DUF169;HTH_17 | — |
| 2045217.SAMN16425918.CP064974_5 | **typeXI** | 185,5 | low | Candidatus Moraniibacter | — | — | — | — | Candidatus Moraniibacteriota bacteriu | SAMN16425918 | anthropogenic | 10 | Yes | 4 | CCB3_YggT;G0-G1_switch_2;MCM | — |
| 1898104.SAMN21435363.JAIUPL01000 | **typeXI** | 179,2 | low | Bacteroidota | — | — | — | — | Bacteroidota bacterium | SAMN21435363 | anthropogenic | 6 | Yes | 3 | CRISPR_Cas2;Cas_Cas1 | — |
| 1262921.PRJEB697.HF992632_9 | **typeXI** | 176,1 | low | Bacteroidota | Bacteroidia | Bacteroidales | Prevotellaceae | Prevotella | Prevotella sp. CAG:1185 | PRJEB697 | host-associated | 7 | Yes | 4 | WG_beta_rep;Fer4_7 | — |
| 2202734.SAMN14239044.JABWBR010 | **typeXI** | 170,7 | low | Bacteroidota | Saprospiria | Saprospirales | Saprospiraceae | — | Saprospiraceae bacterium | SAMN14239044 | anthropogenic | 8 | Yes | 4 | PIN_3;DUF1801;CRISPR_Cas2;Ca | — |
| 2202734.SAMN16425902.JADJNF0100 | **typeXI** | 164,1 | low | Bacteroidota | Saprospiria | Saprospirales | Saprospiraceae | — | Saprospiraceae bacterium | SAMN16425902 | anthropogenic | 10 | Yes | 4 | TackOD1;CotH;Por_Secre_tail | — |
| 194843.SAMEA6951536.CAIYES01000 | **typeXI** | 155,6 | low | Bacteroidota | Bacteroidia | Bacteroidales | — | — | uncultured Bacteroidales bacterium | SAMEA6951536 | aquatic | 8 | Yes | 4 | OKR_DC_1_N;Cas6b_C;Cas_Cas | — |
| 194843.SAMEA6951804.CAIQUS01000 | **typeXI** | 148,3 | low | Bacteroidota | Bacteroidia | Bacteroidales | — | — | uncultured Bacteroidales bacterium | SAMEA6951804 | aquatic | 8 | Yes | 4 | CRISPR_Cas2;Cas_Cas1;Cas6b_( | — |
| 2013697.SAMN06767598.PHDA010000 | **typeXI** | 139,5 | low | Bacteroidota | — | — | — | — | Bacteroidetes bacterium HGW-Bacter | SAMN06767598 | terrestrial | 7 | Yes | 4 | Phosphoprotein;Cas6b_C;Cas_Cas | — |
| 42895.SAMN17801374.JAHABZ010000 | **typeXII** | 299 | high | Pseudomonadota | Gammaproteobacteria | Enterobacterales | Enterobacteriaceae | Enterobacter | Enterobacter sp. | SAMN17801374 | host-associated | 11 | Yes | 4 | DUF5375;Zn_Ribbon_Prim;RVT_1 | — |
| 685445.SAMN02440965.KB911089_44 | **typeXII** | 295,7 | high | Pseudomonadota | Gammaproteobacteria | Enterobacterales | Enterobacteriaceae | Klebsiella | Klebsiella aerogenes | SAMN02440965 | host-associated | 11 | Yes | 4 | HTH_3;TMEM170A_B;Resolvase;F | — |
| 910996.SAMN10245858.RCWP010000 | **typeXII** | 294,5 | high | Pseudomonadota | Gammaproteobacteria | Enterobacterales | Hafniaceae | Hafnia | Hafnia alvei | SAMN10245858 | — | 11 | Yes | 0 | — | — |
| 1608994.SAMN03328806.JYLF0100000 | **typeXII** | 292 | high | Pseudomonadota | Gammaproteobacteria | Pseudomonadales | Pseudomonadaceae | Pseudomonas | Pseudomonas weihenstephanensis | SAMN03328806 | anthropogenic | 11 | Yes | 4 | Arfaptin;ThiD2 | — |
| 1211707.PRJNA170904.HF570389_10 | **typeXII** | 284,5 | high | Pseudomonadota | Gammaproteobacteria | Lysobacterales | Lysobacteraceae | Xanthomonas | Xanthomonas campestris | PRJNA170904 | anthropogenic | 11 | Yes | 4 | Glyco_hydro_15;Trehalose_PPase; | — |
| 190485.SAMN02603845.AE008922_36 | **typeXII** | 284,5 | high | Pseudomonadota | Gammaproteobacteria | Lysobacterales | Lysobacteraceae | Xanthomonas | Xanthomonas campestris | SAMN02603845 | — | 11 | Yes | 0 | — | — |
| 406819.SAMN02141584.AJHK0200001 | **typeXII** | 284,5 | high | Pseudomonadota | Betaproteobacteria | Burkholderiales | Burkholderiaceae | Burkholderia | Burkholderia sp. SJ98 | SAMN02141584 | terrestrial | 11 | Yes | 4 | ER_lumen_recept;MCPsignal;Dabb | — |
| 1802253.SAMN04313803.MIBZ010000 | **typeXII** | 284,3 | high | Campylobacterota | Epsilonproteobacteria | Campylobacterales | Sulfurimonadaceae | Sulfurimonas | Sulfurimonas sp. RIFCSPLOWO2_12 | SAMN04313803 | terrestrial | 8 | Yes | 4 | PhdYeFM_antitox;ParE_toxin;DUF6 | — |
| 1417228.SAMN04498477.CP014579_1 | **typeXII** | 281,9 | high | Pseudomonadota | Betaproteobacteria | Burkholderiales | Burkholderiaceae | Paraburkholderia | Paraburkholderia phytofirmans | SAMN04498477 | temperature | 11 | Yes | 4 | MacP_activator;SIR2_2;Nuc_deoxy | — |
| 2026748.SAMN16635730.JADMJM0100 | **typeXII** | 279 | high | Pseudomonadota | Alphaproteobacteria | Hyphomonadales | Hyphomonadaceae | — | Hyphomonadaceae bacterium | SAMN16635730 | terrestrial | 11 | Yes | 4 | AAA_31;TPR_12 | — |
| 2282150.SAMN18059877.JAFLCU0100 | **typeXII** | 278,5 | high | Pseudomonadota | Alphaproteobacteria | Caulobacterales | — | — | Caulobacterales bacterium | SAMN18059877 | anthropogenic | 6 | Yes | 2 | Acyl-CoA_dh_M;AKAP2_C | — |
| 224209.SAMEA8805150.CAJTUR0100 | **typeXII** | 273,5 | high | Bacillota | Bacilli | — | — | — | uncultured Bacilli bacterium | SAMEA8805150 | host-associated | 11 | Yes | 4 | HTH_3;FtsX;DnaB_C;Dockerin_1 | — |
| 1913988.SAMN16745442.JAGWZH010 | **typeXII** | 269,8 | high | Pseudomonadota | Alphaproteobacteria | — | — | — | Alphaproteobacteria bacterium | SAMN16745442 | aquatic | 7 | Yes | 3 | Acetyltransf_6;TPR_19 | — |
| 1913988.SAMN19298566.JAHJSN0100 | **typeXII** | 268,6 | high | Pseudomonadota | Alphaproteobacteria | — | — | — | Alphaproteobacteria bacterium | SAMN19298566 | terrestrial | 11 | Yes | 4 | DUF2147;Gly_transporter;DUF128 | — |
| 1479235.SAMN03785500.LJZS0100000 | **typeXII** | 268,6 | high | Pseudomonadota | Gammaproteobacteria | Oceanospirillales | Halomonadaceae | Halomonas | Halomonas sp. HL-48 | SAMN03785500 | salinity | 11 | Yes | 4 | Sbt_1;ZinT;SmpB;Polyketide_cyc | — |
| 1116369.SAMN02441719.KB890024_1 | **typeXII** | 260,8 | low | Pseudomonadota | Alphaproteobacteria | Hyphomicrobiales | Rhizobiaceae | Hoeflea | Hoeflea sp. 108 | SAMN02441719 | anthropogenic | 10 | Yes | 4 | FA_desaturase;DUF5631;Phage_in | — |
| 2862676.SAMN20394465.JAHXYK0100 | **typeXII** | 256,5 | low | Pseudomonadota | Gammaproteobacteria | Lysobacterales | Lysobacteraceae | Lysobacter | Lysobacter sp. ESA13C | SAMN20394465 | host-associated | 10 | Yes | 4 | Zeta_toxin;MraZ;XAC0095_dom;NA | — |
| 194843.SAMEA104666335.ONBO0100 | **typeXII** | 229,4 | low | Bacteroidota | Bacteroidia | Bacteroidales | — | — | uncultured Bacteroidales bacterium | SAMEA104666335 | host-associated | 10 | Yes | 4 | AAA-ATPase_like;HAD_SAK_2 | — |
| 152509.SAMEA6946966.CAITFF01000 | **typeXII** | 226,1 | low | Bacteroidota | — | — | — | — | uncultured Bacteroidota bacterium | SAMEA6946966 | aquatic | 8 | Yes | 4 | MHV_Nsp3_DPUP;DUF4784;DnaE | — |
| 1913988.SAMN12581861.WRAU01000 | **typeXII** | 213,6 | low | Pseudomonadota | Alphaproteobacteria | — | — | — | Alphaproteobacteria bacterium | SAMN12581861 | host-associated | 10 | Yes | 4 | NRDD;Fer4_12;NMO;RNase_PH | — |
| 1300343.SAMN03135132.JSAQ010000 | **typeXII** | 211,4 | low | Bacteroidota | Flavobacteriia | Flavobacteriales | Flavobacteriaceae | Dokdonia | Dokdonia donghaensis | SAMN03135132 | host-associated | 10 | Yes | 4 | Ribosomal_S15;RNase_PH;GH85_ | — |
| 2055788.SAMN08019674.DOVO01000 | **typeXIII** | 364,3 | high | Cyanobacteriota | — | — | — | — | Cyanobacteria bacterium UBA9226 | SAMN08019674 | anthropogenic | 10 | Yes | 4 | Response_reg;LCIB_C_CA;CBM_1 | — |
| 2055779.SAMN08018872.DNXU010000 | **typeXIII** | 363,4 | high | Cyanobacteriota | — | — | — | — | Cyanobacteria bacterium UBA8803 | SAMN08018872 | anthropogenic | 4 | Yes | 3 | Response_reg | — |
| 2692870.SAMN13702772.JACJPF0100 | **typeXIII** | 362,5 | high | Cyanobacteriota | Cyanophyceae | Leptolyngbyales | Trichocoleusaceae | Trichocoleus | Trichocoleus sp. FACHB-40 | SAMN13702772 | terrestrial | 9 | Yes | 4 | SWIM;VWA | — |
| 1211.SAMEA6944671.CAIRPA0100003 | **typeXIII** | 362,4 | high | Cyanobacteriota | — | — | — | — | uncultured cyanobacterium | SAMEA6944671 | aquatic | 11 | Yes | 4 | WGR;SWIM;PaO;Metallophos | — |
| 1454205.SAMN05826284.CP017708_2 | **typeXIII** | 359,2 | high | Cyanobacteriota | Cyanophyceae | Coleofasciculales | Coleofasciculaceae | Moorena | Moorena producens | SAMN05826284 | aquatic | 11 | Yes | 4 | DUF2157;Pneumo_M2;XisI;XisH | — |
| 393003.SAMN09074762.QLNL0100000 | **typeXIII** | 355,3 | high | Bacteroidota | Chitinophagia | Chitinophagales | Chitinophagaceae | Chitinophaga | Chitinophaga ginsengisegetis | SAMN09074762 | host-associated | 11 | Yes | 4 | WGR;SWIM;TonB_dep_Rec_b-bar | — |
| 1157708.SAMN02440495.KB907450_1 | **typeXIII** | 353,1 | high | Pseudomonadota | Betaproteobacteria | Burkholderiales | Comamonadaceae | Variovorax | Variovorax atrisoli | SAMN02440495 | host-associated | 11 | Yes | 4 | HEAT_2;SWIM;Acetyltransf_1;RIO | — |
| 1869181.SAMN17140208.JAFAIX0100 | **typeXIII** | 352,6 | high | Bacteroidota | Chitinophagia | Chitinophagales | Chitinophagaceae | Chitinophaga | Chitinophaga sp. | SAMN17140208 | terrestrial | 11 | Yes | 4 | WGR;SWIM;TonB_dep_Rec_b-bar | — |
| 1095768.PRJEA70551.HE578945_39 | **typeXIII** | 350,3 | high | Pseudomonadota | Gammaproteobacteria | Enterobacterales | Enterobacteriaceae | Phytobacter | Phytobacter massiliensis | PRJEA70551 | host-associated | 11 | Yes | 4 | WGR;SWIM;HATPase_c;Lipoprotei | — |
| 1849603.SAMN17801224.JAGZWF010 | **typeXIII** | 349,3 | high | Pseudomonadota | Gammaproteobacteria | Enterobacterales | Enterobacteriaceae | — | Enterobacteriaceae bacterium | SAMN17801224 | host-associated | 11 | Yes | 4 | WGR;SWIM;HATPase_c;Lipoprotei | — |
| 1884385.SAMN05518669.FNOL010000 | **typeXIII** | 348,9 | high | Pseudomonadota | Betaproteobacteria | Burkholderiales | Comamonadaceae | Variovorax | Variovorax sp. YR634 | SAMN05518669 | host-associated | 11 | Yes | 4 | HEAT_2;SWIM;Acetyltransf_1;RIO | — |
| 1871043.SAMN18061513.JAFKMS0100 | **typeXIII** | 347,2 | high | Pseudomonadota | Betaproteobacteria | Burkholderiales | Comamonadaceae | Variovorax | Variovorax sp. | SAMN18061513 | anthropogenic | 11 | Yes | 4 | HEAT_2;SWIM;Acetyltransf_1;RIO | — |
| 1882827.SAMN05443579.FOWG01000 | **typeXIII** | 347,2 | high | Pseudomonadota | Betaproteobacteria | Burkholderiales | Comamonadaceae | Variovorax | Variovorax sp. PDC80 | SAMN05443579 | anthropogenic | 11 | Yes | 4 | HEAT_2;SWIM;4HBT;RIO1 | — |
| 1297865.SAMN02952940.APJD010000 | **typeXIII** | 346,9 | high | Pseudomonadota | Alphaproteobacteria | Hyphomicrobiales | Nitrobacteraceae | Bradyrhizobium | Bradyrhizobium sp. OHSU_III | SAMN02952940 | anthropogenic | 11 | Yes | 4 | LplA-B_cat;FliMN_C;SWIM;WGR | — |
| 1761785.SAMN04487979.FZNL010000 | **typeXIII** | 346,3 | high | Bacteroidota | Flavobacteriia | Flavobacteriales | Flavobacteriaceae | Flavobacterium | Flavobacterium sp. ov086 | SAMN04487979 | host-associated | 11 | Yes | 4 | SWIM;WGR;AMP-binding;DUF771 | — |
| 1128427.SAMN02256433.KB904821_2 | **typeXIII** | 346,1 | high | Cyanobacteriota | Cyanophyceae | Oscillatoriales | — | — | filamentous cyanobacterium ESFC-1 | SAMN02256433 | salinity | 11 | Yes | 4 | Endo_dU;Uma2;Psb34 | — |
| 2789216.SAMN12641115.CP043489_3 | **typeXIII** | 344,8 | high | Pseudomonadota | Alphaproteobacteria | Hyphomicrobiales | Xanthobacteraceae | Labrys | Labrys sp. KNU-23 | SAMN12641115 | host-associated | 11 | Yes | 4 | WGR;SWIM;Rubrerythrin;Yip1 | — |
| 2056868.SAMN08111093.QEHP010001 | **typeXIII** | 344,8 | high | Bacteroidota | Flavobacteriia | Flavobacteriales | Weeksellaceae | Chryseobacterium | Chryseobacterium sp. HMWF035 | SAMN08111093 | anthropogenic | 11 | Yes | 4 | Arm-DNA-bind_5;WGR;SWIM | — |
| 2578106.SAMN11620940.CP042171_9 | **typeXIII** | 344,2 | high | Bacteroidota | Sphingobacteriia | Sphingobacteriales | Sphingobacteriaceae | Pedobacter | Pedobacter sp. KBS0701 | SAMN11620940 | host-associated | 11 | Yes | 4 | DUF4953;LRR_ComC;WGR;SWIM | — |
| 2587064.SAMN12024188.JACHWM010 | **typeXIII** | 343 | high | Pseudomonadota | Alphaproteobacteria | Hyphomicrobiales | Methylobacteriaceae | Methylobacterium | Methylobacterium sp. R2-1 | SAMN12024188 | terrestrial | 11 | Yes | 4 | WGR;SWIM;Acetyltransf_6 | — |
| 651561.SAMN06453202.DCFI0100002 | **typeXIII** | 343 | high | Bacteroidota | Flavobacteriia | Flavobacteriales | Weeksellaceae | Chryseobacterium | Chryseobacterium arthrosphaerae | SAMN06453202 | host-associated | 11 | Yes | 4 | Amidohydro_1;Urocanase_C;WGR; | — |
| 946333.SAMN04621830.CP015118_35 | **typeXIII** | 342,7 | high | Pseudomonadota | Betaproteobacteria | Burkholderiales | Sphaerotilaceae | Piscinibacter | Piscinibacter gummiphilus | SAMN04621830 | host-associated | 11 | Yes | 4 | XPC-binding;SWIM;LysR_substrate | — |
| 2026763.SAMN10607053.SKYM010003 | **typeXIII** | 341,6 | high | Myxococcota | Myxococcia | Myxococcales | — | — | Myxococcales bacterium | SAMN10607053 | salinity | 8 | Yes | 4 | SWIM;Beta-prop_IFT122_1st;DUF2 | — |
| 1230476.SAMN01893863.KE747879_2 | **typeXIII** | 338,7 | high | Pseudomonadota | Alphaproteobacteria | Hyphomicrobiales | Nitrobacteraceae | Bradyrhizobium | Bradyrhizobium sp. DFCI-1 | SAMN01893863 | anthropogenic | 11 | Yes | 4 | Polysacc_deac_1;DUF4148;FliMN_ | — |
| 2015576.SAMN07280669.NKIY0100000 | **typeXIII** | 338,7 | high | Pseudomonadota | Alphaproteobacteria | Hyphomicrobiales | Nitrobacteraceae | — | Bradyrhizobiaceae bacterium PARB1 | SAMN07280669 | — | 11 | Yes | 0 | — | — |
| 1045867.SAMN18262562.JAGGDU010 | **typeXIII** | 338,1 | high | Cyanobacteriota | Cyanophyceae | Oscillatoriales | Microcoleaceae | Trichodesmium | Trichodesmium erythraeum | SAMN18262562 | aquatic | 11 | Yes | 4 | GATase;Ribonucleas_3_3;Glyco_tr | — |
| 2040292.SAMEA104414344.OEPV010 | **typeXIII** | 338 | high | Bacteroidota | Bacteroidia | Bacteroidales | Dysgonomonadaceae | Pseudodysgonomonas | Pseudodysgonomonas massiliensis | SAMEA104414344 | host-associated | 11 | Yes | 4 | Bgal_small_N;Alpha_L_fucos;WGR | — |
| 693986.SAMN03075686.CP003811_16 | **typeXIII** | 337,9 | high | Pseudomonadota | Alphaproteobacteria | Hyphomicrobiales | Methylobacteriaceae | Methylobacterium | Methylobacterium oryzae | SAMN03075686 | — | 11 | Yes | 0 | — | — |
| 426355.SAMN00000277.CP001001_13 | **typeXIII** | 337,6 | high | Pseudomonadota | Alphaproteobacteria | Hyphomicrobiales | Methylobacteriaceae | Methylobacterium | Methylobacterium radiotolerans | SAMN00000277 | anthropogenic | 11 | Yes | 4 | Enterotoxin_HS1;SWIM;WGR | — |
| 179408.SAMN02261335.CP003614_51 | **typeXIII** | 335,8 | high | Cyanobacteriota | Cyanophyceae | Oscillatoriales | Oscillatoriaceae | Phormidium | Phormidium nigroviride | SAMN02261335 | anthropogenic | 11 | Yes | 4 | HicA_toxin;HicB_lk_antitox;DUF567 | — |
| 2587122.SAMN12024204.JACHYS0100 | **typeXIII** | 335,6 | high | Pseudomonadota | Betaproteobacteria | Burkholderiales | Sphaerotilaceae | Roseateles | Mitsuaria sp. BK037 | SAMN12024204 | host-associated | 11 | Yes | 4 | XPC-binding;SWIM;YitT_membrane | — |
| 2777973.SAMN16351716.JADEWD010 | **typeXIII** | 334,5 | high | Cyanobacteriota | Cyanophyceae | Nostocales | Aphanizomenonaceae | Cuspidothrix | Cuspidothrix issatschenkoi | SAMN16351716 | aquatic | 6 | Yes | 2 | HSP70;NAGPA | — |
| 398578.SAMN00623041.CP000884_16 | **typeXIII** | 332,6 | high | Pseudomonadota | Betaproteobacteria | Burkholderiales | Comamonadaceae | Delftia | Delftia acidovorans | SAMN00623041 | host-associated | 11 | Yes | 4 | TPR_NOC4;HTH_18;SWIM;XPC-b | — |
| 2448779.SAMN18059842.JAFLBL0100 | **typeXIII** | 331,4 | high | Bacteroidota | Chitinophagia | Chitinophagales | — | — | Chitinophagales bacterium | SAMN18059842 | anthropogenic | 11 | Yes | 4 | WGR;SWIM;Amidohydro_1;LacI | — |

| | | | | | | | | | | | | | | | | | |
|---|---|---|---|---|---|---|---|---|---|---|---|---|---|---|---|---|---|
| 1032.SAMN18076803.JAGVHW010000 | **typeXIII** | | 330,9 | **high** | Pseudomonadota | Gammaproteobacteria | Thiotrichales | Thiotrichaceae | Thiothrix | Thiothrix sp. | SAMN18076803 | anthropogenic | 11 | Yes | 4 | Aminotran_5;WGR;SWIM | — |
| 2052484.SAMN07200918.NIOE0100000 | **typeXIII** | | 330,8 | **high** | Pseudomonadota | Betaproteobacteria | Burkholderiales | Sphaerotilaceae | Roseateles | Roseateles noduli | SAMN07200918 | host-associated | 11 | Yes | 4 | Amidohydro_2;YitT_membrane;SWI | — |
| 513160.SAMN12129767.VIDT01000849 | **typeXIII** | | 330,6 | **high** | Pseudomonadota | Betaproteobacteria | Burkholderiales | Sphaerotilaceae | Ideonella | Ideonella azotifigens | SAMN12129767 | host-associated | 2 | Yes | 1 | SWIM | — |
| 2692887.SAMN13702731.JACJQT0100 | **typeXIII** | | 330,3 | **high** | Cyanobacteriota | Cyanophyceae | Nostocales | Aphanizomenonaceae | Aphanizomenon | Aphanizomenon flos-aquae | SAMN13702731 | aquatic | 11 | Yes | 4 | HicA_toxin;Peptidase_C13;AdoHcy | — |
| 2745558.SAMN15428588.CP058988_2 | **typeXIII** | | 328,1 | **high** | Bacteroidota | Flavobacteriia | Flavobacteriales | Flavobacteriaceae | Cellulophaga | Cellulophaga sp. HaHa_2_95 | SAMN15428588 | host-associated | 11 | Yes | 4 | WGR;SWIM;LRR_14 | — |
| 100233.SAMEA6952382.CAIXQV01000 | **typeXIII** | | 327,7 | **high** | Planctomycetota | Planctomycetia | Planctomycetales | Planctomycetaceae | — | uncultured Planctomycetaceae bacteri | SAMEA6952382 | aquatic | 4 | Yes | 3 | Sulfatase;RlaP | — |
| 179408.SAMN02261335.CP003614_38 | **typeXIII** | | 326,5 | **high** | Cyanobacteriota | Cyanophyceae | Oscillatoriales | Oscillatoriaceae | Phormidium | Phormidium nigroviride | SAMN02261335 | anthropogenic | 11 | Yes | 4 | DEAD;Band_7;FrhB_FdhB_C | — |
| 2026763.SAMN16426551.JADKKK0100 | **typeXIII** | | 326,4 | **high** | Myxococcota | Myxococcia | Myxococcales | — | — | Myxococcales bacterium | SAMN16426551 | anthropogenic | 11 | Yes | 4 | WGR;SWIM;Sel1;ABC_tran | — |
| 2651161.SAMN12878373.WBUJ010000 | **typeXIII** | | 325,9 | **high** | Pseudomonadota | Gammaproteobacteria | — | Candidatus Competibacte | Candidatus Contendiba | Candidatus Contendobacter sp. | SAMN12878373 | — | 9 | Yes | 0 | — | — |
| 2052164.SAMN08179277.PLDW01000 | **typeXIII** | | 325,5 | **high** | Planctomycetota | Planctomycetia | Gemmatales | Gemmataceae | — | Gemmataceae bacterium | SAMN08179277 | temperature | 11 | Yes | 4 | Trypsin_2;PDZ_2;SWIM;HEAT_2 | — |
| 2053538.SAMN10587497.RXIW010000 | **typeXIII** | | 324,9 | **high** | Pseudomonadota | Gammaproteobacteria | — | Candidatus Competibacte | — | Candidatus Competibacteraceae bacte | SAMN10587497 | anthropogenic | 9 | Yes | 4 | WGR;SWIM;Lactate_perm | — |
| 2026763.SAMN16426013.JADJQX0100 | **typeXIII** | | 324,1 | **high** | Myxococcota | Myxococcia | Myxococcales | — | — | Myxococcales bacterium | SAMN16426013 | anthropogenic | 11 | Yes | 4 | WGR;SWIM;DUF937 | — |
| 2562705.SAMEA6956407.CAIXXB0100 | **typeXIII** | | 324,1 | **high** | Verrucomicrobiota | Verrucomicrobiia | Verrucomicrobiales | Akkermansiaceae | — | Akkermansiaceae bacterium | SAMEA6956407 | aquatic | 6 | Yes | 2 | NHL;PIN | — |
| 2024858.SAMN07620073.PABA010001 | **typeXIII** | | 323,8 | **high** | Myxococcota | — | Polyangiales | Sandaracinaceae | Sandaracinus | Sandaracinus sp. | SAMN07620073 | aquatic | 8 | Yes | 4 | HEAT_2;SWIM;PEGA | — |
| 92487.SAMN02745130.FUYB01000016 | **typeXIII** | | 321,8 | **high** | Pseudomonadota | Gammaproteobacteria | Thiotrichales | Thiotrichaceae | Thiothrix | Thiothrix eikelboomii | SAMN02745130 | anthropogenic | 11 | Yes | 4 | DCC1-like;dCache_2;SWIM;HEAT_ | — |
| 2053538.SAMN16426005.JADJQQ0100 | **typeXIII** | | 321,7 | **high** | Pseudomonadota | Gammaproteobacteria | — | Candidatus Competibacte | — | Candidatus Competibacteraceae bacte | SAMN16426005 | anthropogenic | 11 | Yes | 4 | WGR;Chalcone_3;SWIM | — |
| 502025.SAMN00002596.CP001804_15 | **typeXIII** | | 321,5 | **high** | Myxococcota | — | Haliangiales | Kofleriaceae | Haliangium | Haliangium ochraceum | SAMN00002596 | salinity | 11 | Yes | 4 | HEAT_2;SWIM;WGR | — |
| 1131567.SAMD00245618.BMYU010000 | **typeXIII** | | 321,3 | **high** | Pseudomonadota | Betaproteobacteria | Burkholderiales | Oxalobacteraceae | Undibacterium | Undibacterium squillarum | SAMD00245618 | anthropogenic | 11 | Yes | 4 | Phosphoesterase;CC_CEP250;SW | — |
| 2052181.SAMN11369556.SYPF010001 | **typeXIII** | | 321,3 | **high** | Planctomycetota | Planctomycetia | — | — | — | Planctomycetia bacterium | SAMN11369556 | aquatic | 8 | Yes | 4 | Pan_kinase;MMR_HSR1;SWIM | — |
| 2053538.SAMN16426005.JADJQQ0100 | **typeXIII** | | 320,5 | **high** | Pseudomonadota | Gammaproteobacteria | — | Candidatus Competibacte | — | Candidatus Competibacteraceae bacte | SAMN16426005 | anthropogenic | 11 | Yes | 0 | — | — |
| 1951615.SAMN06451712.DCNP010000 | **typeXIII** | | 320 | **high** | Pseudomonadota | Gammaproteobacteria | — | Candidatus Competibacte | — | Competibacteraceae bacterium UBA15 | SAMN06451712 | anthropogenic | 8 | Yes | 4 | AAA_22;SWIM;WGR | — |
| 1965545.SAMN06473586.NFLT010000 | **typeXIII** | | 318,9 | **high** | Bacillota | Clostridia | Lachnospirales | Lachnospiraceae | Tyzzerella | Tyzzerella sp. An114 | SAMN06473586 | host-associated | 11 | Yes | 4 | DUF6849;SWIM;PTS_EIIA_2;PTS_ | — |
| 2769423.SAMD00245656.BNCN010000 | **typeXIII** | | 318,3 | **high** | Pseudomonadota | Betaproteobacteria | Burkholderiales | Comamonadaceae | Comamonas | Comamonas sp. KCTC 72670 | SAMD00245656 | host-associated | 11 | Yes | 4 | GMC_oxred_C;Hydrolase_4;SWIM | — |
| 156588.SAMEA6945809.CAJCBL01000 | **typeXIII** | | 318,2 | **high** | Verrucomicrobiota | — | — | — | — | uncultured Verrucomicrobiota bacteriu | SAMEA6945809 | aquatic | 10 | Yes | 4 | OTCace;DUF3239;SWIM;HEAT_2 | — |
| 261164.SAMN04549283.CP014945_13 | **typeXIII** | | 315,6 | **high** | Pseudomonadota | Gammaproteobacteria | Moraxellales | Moraxellaceae | Psychrobacter | Psychrobacter alimentarius | SAMN04549283 | anthropogenic | 11 | Yes | 4 | HEAT_2;SWIM;Lipase_GDSL_2;C | — |
| 2024858.SAMN07620073.PABA010001 | **typeXIII** | | 315,5 | **high** | Myxococcota | — | Polyangiales | Sandaracinaceae | Sandaracinus | Sandaracinus sp. | SAMN07620073 | aquatic | 7 | Yes | 3 | 23S_rRNA_IVP;NTP_transf_9;DUF | — |
| 244328.SAMEA8805375.CAJTZO01000 | **typeXIII** | | 315,5 | **high** | Bacillota | Clostridia | — | — | — | uncultured Clostridia bacterium | SAMEA8805375 | host-associated | 11 | Yes | 4 | CoaE;DNA_pol_A;SWIM;DUF6849 | — |
| 2053632.SAMN19225299.DXZA010000 | **typeXIII** | | 315,4 | **high** | Bacillota | Clostridia | Lachnospirales | Lachnospiraceae | Tyzzerella | Tyzzerella sp. | SAMN19225299 | host-associated | 11 | Yes | 4 | DUF6849;SWIM;ProRS-C_1;Lar_r | — |
| 244328.SAMEA7202299.CAJFIR01000 | **typeXIII** | | 314,8 | **high** | Bacillota | Clostridia | — | — | — | uncultured Clostridia bacterium | SAMEA7202299 | host-associated | 10 | Yes | 4 | DUF6849;SWIM;Pantoate_transf;A | — |
| 2053538.SAMN16425960.JADJPA0100 | **typeXIII** | | 314,1 | **high** | Pseudomonadota | Gammaproteobacteria | — | Candidatus Competibacte | — | Candidatus Competibacteraceae bacte | SAMN16425960 | — | 11 | Yes | 0 | — | — |
| 244328.SAMEA8805706.CAJUKL01000 | **typeXIII** | | 313,9 | **high** | Bacillota | Clostridia | — | — | — | uncultured Clostridia bacterium | SAMEA8805706 | — | 11 | Yes | 0 | — | — |
| 297314.SAMEA8805501.CAJUDQ0100 | **typeXIII** | | 312,8 | **high** | Bacillota | Clostridia | Lachnospirales | Lachnospiraceae | — | uncultured Lachnospiraceae bacterium | SAMEA8805501 | host-associated | 11 | Yes | 4 | ABC_membrane;Response_reg;SW | — |
| 297314.SAMEA8805536.CAJUIB01000 | **typeXIII** | | 312,8 | **high** | Bacillota | Clostridia | Lachnospirales | Lachnospiraceae | — | uncultured Lachnospiraceae bacterium | SAMEA8805536 | — | 11 | Yes | 0 | — | — |
| 997898.SAMN02596761.KB851130_27 | **typeXIII** | | 312,7 | **high** | Bacillota | Clostridia | Eubacteriales | Clostridiaceae | Clostridium | Clostridium butyricum | SAMN02596761 | host-associated | 11 | Yes | 4 | MarR;A_deaminase;SWIM;DUF684 | — |
| 1946346.SAMN06454910.DCDL010000 | **typeXIII** | | 312,2 | **high** | Bacillota | Clostridia | Eubacteriales | Clostridiaceae | Clostridium | Clostridium sp. UBA1056 | SAMN06454910 | anthropogenic | 11 | Yes | 4 | DUF6849;SWIM;Resolvase | — |
| 244328.SAMEA8805806.CAJUOZ01000 | **typeXIII** | | 311,5 | **high** | Bacillota | Clostridia | — | — | — | uncultured Clostridia bacterium | SAMEA8805806 | host-associated | 11 | Yes | 4 | DUF6849;SWIM;AA_permease_2; | — |
| 2052164.SAMN08179277.PLDW01000 | **typeXIII** | | 311,4 | **high** | Planctomycetota | Planctomycetia | Gemmatales | Gemmataceae | — | Gemmataceae bacterium | SAMN08179277 | temperature | 11 | Yes | 4 | HEAT_2;SWIM;Pkinase;Sigma70_ | — |
| 297314.SAMEA8805527.CAJUDE01000 | **typeXIII** | | 311,2 | **high** | Bacillota | Clostridia | Lachnospirales | Lachnospiraceae | — | uncultured Lachnospiraceae bacterium | SAMEA8805527 | host-associated | 11 | Yes | 4 | DUF6849;SWIM;DUF6056;DUF60 | — |
| 2024858.SAMN07620073.PABA010001 | **typeXIII** | | 311 | **high** | Myxococcota | — | Polyangiales | Sandaracinaceae | Sandaracinus | Sandaracinus sp. | SAMN07620073 | aquatic | 11 | Yes | 4 | DUF779;Abhydrolase_1;SWIM;HE | — |
| 446043.SAMEA8805476.CAJUBU01000 | **typeXIII** | | 310,9 | **high** | Bacillota | Clostridia | Lachnospirales | Lachnospiraceae | Lachnospira | uncultured Lachnospira sp. | SAMEA8805476 | host-associated | 11 | Yes | 4 | DUF6849;SWIM;SBP_bac_1;HAT | — |
| 100233.SAMEA6949581.CAIYBH01000 | **typeXIII** | | 310,7 | **high** | Planctomycetota | Planctomycetia | Planctomycetales | Planctomycetaceae | — | uncultured Planctomycetaceae bacteri | SAMEA6949581 | aquatic | 11 | Yes | 4 | LRR_RI_capping;Methyltransf_11; | — |
| 1898207.SAMN13894064.JAAYMT0100 | **typeXIII** | | 310,6 | **high** | Bacillota | Clostridia | Eubacteriales | — | — | Clostridiales bacterium | SAMN13894064 | temperature | 7 | Yes | 3 | SWIM;DUF4179 | — |
| 297314.SAMEA8805036.CAJTOU01000 | **typeXIII** | | 309,9 | **high** | Bacillota | Clostridia | Lachnospirales | Lachnospiraceae | — | uncultured Lachnospiraceae bacterium | SAMEA8805036 | — | 6 | Yes | 0 | — | — |
| 707003.SAMEA8805243.CAJTTB01000 | **typeXIII** | | 309,5 | **high** | Bacillota | Clostridia | Eubacteriales | Oscillospiraceae | — | uncultured Oscillospiraceae bacterium | SAMEA8805243 | host-associated | 11 | Yes | 4 | MIF;NUDIX;SWIM;DUF6849 | — |
| 935198.SAMN02603538.CP001056_18 | **typeXIII** | | 309,3 | **high** | Bacillota | Clostridia | Eubacteriales | Clostridiaceae | Clostridium | Clostridium botulinum | SAMN02603538 | host-associated | 11 | Yes | 4 | Exo_endo_phos;DUF4004;SWIM;D | — |
| 1946690.SAMN06453118.DESG01000 | **typeXIII** | | 309,3 | **high** | Bacillota | Clostridia | Lachnospirales | Lachnospiraceae | Lachnoclostridium | Lachnoclostridium sp. UBA3320 | SAMN06453118 | host-associated | 10 | Yes | 4 | DUF6849;SWIM;DUF848 | — |
| 244328.SAMEA8805759.CAJUNT01000 | **typeXIII** | | 309,1 | **high** | Bacillota | Clostridia | — | — | — | uncultured Clostridia bacterium | SAMEA8805759 | host-associated | 8 | Yes | 3 | DUF6548;SWIM;DUF6849 | — |
| 100233.SAMEA6949573.CAIYBK01000 | **typeXIII** | | 308,6 | **high** | Planctomycetota | Planctomycetia | Planctomycetales | Planctomycetaceae | — | uncultured Planctomycetaceae bacteri | SAMEA6949573 | aquatic | 10 | Yes | 4 | GFO_IDH_MocA_C2;ACP_syn_III_ | — |
| 297314.SAMEA8805577.CAJUGZ01000 | **typeXIII** | | 308,3 | **high** | Bacillota | Clostridia | Lachnospirales | Lachnospiraceae | — | uncultured Lachnospiraceae bacterium | SAMEA8805577 | host-associated | 9 | Yes | 4 | Response_reg;HATPase_c_5;SWI | — |
| 251229.SAMN02261359.CP003597_35 | **typeXIII** | | 307,8 | **high** | Cyanobacteriota | Cyanophyceae | Chroococcidiopsidales | Chroococcidiopsidaceae | Chroococcidiopsis | Chroococcidiopsis thermalis | SAMN02261359 | anthropogenic | 11 | Yes | 4 | DUF7755;ParE_toxin;Band_7 | — |
| 2026763.SAMN17180964.JAEUJY0100 | **typeXIII** | | 307,7 | **high** | Myxococcota | Myxococcia | Myxococcales | — | — | Myxococcales bacterium | SAMN17180964 | anthropogenic | 7 | Yes | 4 | HEAT_2;SWIM;WGR;DNA_ligase_ | — |
| 297314.SAMEA8805519.CAJUCS01000 | **typeXIII** | | 307,7 | **high** | Bacillota | Clostridia | Lachnospirales | Lachnospiraceae | — | uncultured Lachnospiraceae bacterium | SAMEA8805519 | host-associated | 11 | Yes | 4 | DUF6849;SWIM;ROK;Response_r | — |
| 297314.SAMEA8804990.CAJTJY01000 | **typeXIII** | | 307,6 | **high** | Bacillota | Clostridia | Lachnospirales | Lachnospiraceae | — | uncultured Lachnospiraceae bacterium | SAMEA8804990 | host-associated | 11 | Yes | 4 | DUF6849;SWIM;NMO;ACCA | — |
| 297314.SAMEA8804971.CAJTJZ01000 | **typeXIII** | | 307,2 | **high** | Bacillota | Clostridia | Lachnospirales | Lachnospiraceae | — | uncultured Lachnospiraceae bacterium | SAMEA8804971 | host-associated | 11 | Yes | 4 | GAIN_ADGRA3;SWIM;MCPsignal; | — |
| 297314.SAMEA8805211.CAJTRU01000 | **typeXIII** | | 306,8 | **high** | Bacillota | Clostridia | Lachnospirales | Lachnospiraceae | — | uncultured Lachnospiraceae bacterium | SAMEA8805211 | — | 9 | Yes | 0 | — | — |
| 297314.SAMEA8805591.CAJUGJ01000 | **typeXIII** | | 306,8 | **high** | Bacillota | Clostridia | Lachnospirales | Lachnospiraceae | — | uncultured Lachnospiraceae bacterium | SAMEA8805591 | host-associated | 11 | Yes | 4 | HTH_28;Fic;SWIM;OrfB_IS605 | — |
| 297314.SAMEA8805451.CAJUEI01000 | **typeXIII** | | 306,4 | **high** | Bacillota | Clostridia | Lachnospirales | Lachnospiraceae | — | uncultured Lachnospiraceae bacterium | SAMEA8805451 | — | 6 | Yes | 0 | — | — |
| 297314.SAMEA8805268.CAJTXA01000 | **typeXIII** | | 306,3 | **high** | Bacillota | Clostridia | Lachnospirales | Lachnospiraceae | — | uncultured Lachnospiraceae bacterium | SAMEA8805268 | — | 11 | Yes | 0 | — | — |
| 297314.SAMEA8805166.CAJTUU01000 | **typeXIII** | | 305,8 | **high** | Bacillota | Clostridia | Lachnospirales | Lachnospiraceae | — | uncultured Lachnospiraceae bacterium | SAMEA8805166 | host-associated | 11 | Yes | 4 | WYL;SWIM;DUF6849 | — |
| 297314.SAMEA8805034.CAJTNT01000 | **typeXIII** | | 305,3 | **high** | Bacillota | Clostridia | Lachnospirales | Lachnospiraceae | — | uncultured Lachnospiraceae bacterium | SAMEA8805034 | host-associated | 11 | Yes | 4 | ATP_bind_3;LapA_dom;SWIM;DUF | — |
| 446043.SAMEA8805821.CAJUTV01000 | **typeXIII** | | 305,3 | **high** | Bacillota | Clostridia | Lachnospirales | Lachnospiraceae | Lachnospira | uncultured Lachnospira sp. | SAMEA8805821 | — | 11 | Yes | 0 | — | — |
| 297314.SAMEA8805562.CAJUHN01000 | **typeXIII** | | 305,2 | **high** | Bacillota | Clostridia | Lachnospirales | Lachnospiraceae | — | uncultured Lachnospiraceae bacterium | SAMEA8805562 | host-associated | 6 | Yes | 2 | SWIM;DUF6849 | — |
| 297314.SAMEA8805329.CAJTXH01000 | **typeXIII** | | 304,7 | **high** | Bacillota | Clostridia | Lachnospirales | Lachnospiraceae | — | uncultured Lachnospiraceae bacterium | SAMEA8805329 | host-associated | 8 | Yes | 4 | UvrD-helicase;DUF2157;SWIM;DU | — |
| 297314.SAMEA8805824.CAJUSO01000 | **typeXIII** | | 304,4 | **high** | Bacillota | Clostridia | Lachnospirales | Lachnospiraceae | — | uncultured Lachnospiraceae bacterium | SAMEA8805824 | host-associated | 11 | Yes | 4 | DUF4179;ATP_bind_3;SWIM;DUF | — |
| 34034.SAMEA5364575.CAADGB01000 | **typeXIII** | | 304,1 | **high** | — | Deltaproteobacteria | — | — | — | uncultured delta proteobacterium | SAMEA5364575 | temperature | 6 | Yes | 4 | DNA_ligase_OB_2;WGR;SWIM;HE | — |
| 297314.SAMEA8805835.CAJUPJ01000 | **typeXIII** | | 303,2 | **high** | Bacillota | Clostridia | Lachnospirales | Lachnospiraceae | — | uncultured Lachnospiraceae bacterium | SAMEA8805835 | host-associated | 11 | Yes | 4 | UvrD-helicase;DUF2157;SWIM;DU | — |
| 100371.SAMEA6947806.CAIJGL01000 | **typeXIII** | | 299,5 | **high** | Planctomycetota | Planctomycetia | Planctomycetales | — | — | uncultured Planctomycetales bacteriun | SAMEA6947806 | aquatic | 11 | Yes | 4 | SpoIIE;MR_MLE_C;HEAT_2;PTP-I | — |
| 2800791.SAMN17140641.JAFAZH0100 | **typeXIII** | | 298,8 | **high** | Abditibacteriota | Abditibacteriia | Abditibacteriales | Abditibacteriaceae | — | Abditibacteriaceae bacterium | SAMN17140641 | host-associated | 11 | Yes | 4 | SWIM;PSD4;HXXSHH | — |
| 642492.SAMN00013959.CP002582_83 | **typeXIII** | | 298,5 | **high** | Bacillota | Clostridia | Lachnospirales | Cellulosilyticaceae | Cellulosilyticum | Cellulosilyticum lentocellum | SAMN00013959 | host-associated | 11 | Yes | 4 | Flagellin_N;tRNA-synt_2b;SWIM;D | — |
| 360056.SAMN09639612.DSVQ010000 | **typeXIII** | | 298,3 | **high** | Planctomycetota | Planctomycetia | Planctomycetales | Planctomycetaceae | Schlesneria | Schlesneria paludicola | SAMN09639612 | temperature | 11 | Yes | 4 | GHMP_kinases_N;SGSH_C;DUF1 | — |
| 2052181.SAMD00166091.BJHJ010000 | **typeXIII** | | 298,2 | **high** | Planctomycetota | Planctomycetia | — | — | — | Planctomycetia bacterium | SAMD00166091 | aquatic | 9 | Yes | 4 | CbiN;BD-FAE | — |
| 2485925.SAMN12581690.WQUF01000 | **typeXIII** | | 297,3 | **high** | Bacillota | Clostridia | Eubacteriales | Oscillospiraceae | — | Oscillospiraceae bacterium | SAMN12581690 | host-associated | 6 | Yes | 2 | PIN | — |
| 48.SAMN07426514.QFQP01000022_11 | **typeXIII** | | 291,9 | **high** | Myxococcota | Myxococcia | Myxococcales | Archangiaceae | Archangium | Archangium gephyra | SAMN07426514 | anthropogenic | 11 | Yes | 4 | Peptidase_M54;YrhK;SWIM | — |
| 100233.SAMEA6952302.CAITYU01000 | **typeXIII** | | 291,4 | **high** | Planctomycetota | Planctomycetia | Planctomycetales | Planctomycetaceae | — | uncultured Planctomycetaceae bacteri | SAMEA6952302 | aquatic | 6 | Yes | 3 | DUF7474 | — |
| 2049045.SAMN12582053.WRIE010000 | **typeXIII** | | 290,6 | **high** | Bacillota | Clostridia | Eubacteriales | Eubacteriaceae | — | Eubacteriaceae bacterium | SAMN12582053 | host-associated | 11 | Yes | 4 | HATPase_c;SAICAR_synt;SWIM;D | — |
| 2073117.SAMN15013936.JABTTZ0100 | **typeXIII** | | 290,3 | **high** | Chloroflexota | Anaerolineae | Anaerolineales | — | — | Anaerolineales bacterium | SAMN15013936 | temperature | 11 | Yes | 4 | Peptidase_S51;Peripla_BP_2;DUF | — |
| 2026780.SAMN17496266.JAFMMM010 | **typeXIII** | | 289,1 | **high** | Planctomycetota | — | — | — | — | Planctomycetota bacterium | SAMN17496266 | aquatic | 6 | Yes | 2 | ACP_syn_III_C;GFO_IDH_MocA_ | — |
| 1952931.SAMN06452880.DIXK010000 | **typeXIII** | | 287,8 | **high** | Verrucomicrobiota | Verrucomicrobiia | — | — | — | Verrucomicrobia subdivision 3 bacteriu | SAMN06452880 | anthropogenic | 8 | Yes | 4 | SWIM;GFO_IDH_MocA | — |
| 257465.SAMEA6953639.CAINLR01000 | **typeXIII** | | 287,5 | **high** | Verrucomicrobiota | Verrucomicrobiia | Limisphaerales | — | — | uncultured Verrucomicrobia subdivisio | SAMEA6953639 | aquatic | 11 | Yes | 4 | SWIM;NAPRTase_N;NAD_synthas | — |

| | | | | | | | | | | | | | | | | |
|---|---|---|---|---|---|---|---|---|---|---|---|---|---|---|---|---|
| 2052179.SAMN12582057.WRII0100002 | **typeXIII** | 285,2 | **high** | Bacillota | Clostridia | Eubacteriales | Peptococcaceae | — | Peptococcaceae bacterium | SAMN12582057 | host-associated | 11 | Yes | 4 | DUF6849;SWIM;DA_C | — |
| 2026780.SAMN09639629.DSWH01000 | **typeXIII** | 285 | **high** | Planctomycetota | — | — | — | — | Planctomycetota bacterium | SAMN09639629 | temperature | 11 | Yes | 4 | N6_N4_Mtase;GFO_IDH_MocA;PI | — |
| 2026780.SAMN09639583.DSUN01000 | **typeXIII** | 285 | **high** | Planctomycetota | — | — | — | — | Planctomycetota bacterium | SAMN09639583 | temperature | 11 | Yes | 4 | Glyco_hydro_10;NPCBM;GFO_IDH | — |
| 156588.SAMEA6944372.CAIMXY01000 | **typeXIII** | 281,9 | **high** | Verrucomicrobiota | — | — | — | — | uncultured Verrucomicrobiota bacteriu | SAMEA6944372 | aquatic | 10 | Yes | 4 | SWIM;TIR_2 | — |
| 2052186.SAMN11532986.VKHA010000 | **typeXIII** | 280,5 | **high** | Verrucomicrobiota | Verrucomicrobiia | Limisphaerales | — | — | Limisphaerales bacterium | SAMN11532986 | terrestrial | 7 | Yes | 3 | SWIM | — |
| 2026763.SAMN10607053.SKYM010002 | **typeXIII** | 279,1 | **high** | Myxococcota | Myxococcia | Myxococcales | — | — | Myxococcales bacterium | SAMN10607053 | salinity | 8 | Yes | 4 | Radical_SAM;IPPT | — |
| 2026735.SAMN10966762.VGTK010002 | **typeXIII** | 274,5 | **high** | Myxococcota | Myxococcia | — | — | — | Deltaproteobacteria bacterium | SAMN10966762 | aquatic | 5 | Yes | 4 | SWIM;DUF2779;FGE-sulfatase | — |
| 100233.SAMEA6956128.CAIXVO01000 | **typeXIII** | 272,2 | **high** | Planctomycetota | Planctomycetia | Planctomycetales | Planctomycetaceae | — | uncultured Planctomycetaceae bacteri | SAMEA6956128 | aquatic | 3 | Yes | 2 | N6_N4_Mtase | — |
| 1913988.SAMN17181029.JAEUML0100 | **typeXIII** | 269,7 | **high** | Pseudomonadota | Alphaproteobacteria | — | — | — | Alphaproteobacteria bacterium | SAMN17181029 | anthropogenic | 5 | Yes | 3 | SWIM;tRNA-synt_1g | — |
| 1951185.SAMN06451746.DDIR010000 | **typeXIII** | 257,9 | **high** | Planctomycetota | Phycisphaerae | Phycisphaerales | — | — | Phycisphaerales bacterium UBA1845 | SAMN06451746 | anthropogenic | 11 | Yes | 4 | NUDIX;Aminotran_5;CoA_trans;ME | — |
| 297314.SAMEA8804971.CAJTJZ01000 | **typeXIII** | 255,2 | **high** | Bacillota | Clostridia | Lachnospirales | Lachnospiraceae | — | uncultured Lachnospiraceae bacterium | SAMEA8804971 | host-associated | 8 | Yes | 0 | — | — |
| 2026763.SAMN17180967.JAEUKB0100 | **typeXIII** | 254,1 | **high** | Myxococcota | Myxococcia | Myxococcales | — | — | Myxococcales bacterium | SAMN17180967 | anthropogenic | 11 | Yes | 4 | DUF4139;Polysacc_synt_3;Aldo_k | — |
| 1632864.SAMN03417854.CP011270_1 | **typeXIII** | 250,7 | **high** | Planctomycetota | Planctomycetia | Planctomycetales | Planctomycetaceae | Planctomyces | Planctomyces sp. SH-PL14 | SAMN03417854 | anthropogenic | 11 | Yes | 4 | DUF899;DUF7569;PSD1;HEAT_2 | — |
| 2052164.SAMN08179277.PLDW010000 | **typeXIII** | 248,1 | **high** | Planctomycetota | Planctomycetia | Gemmatales | Gemmataceae | — | Gemmataceae bacterium | SAMN08179277 | temperature | 11 | Yes | 4 | Pkinase;DUF3024;FGE-sulfatase | — |
| 100233.SAMEA6946666.CAIVBC01000 | **typeXIII** | 240,6 | **high** | Planctomycetota | Planctomycetia | Planctomycetales | Planctomycetaceae | — | uncultured Planctomycetaceae bacteri | SAMEA6946666 | aquatic | 3 | Yes | 2 | SWIM | — |
| 2026780.SAMN11533026.WSZJ010001 | **typeXIII** | 227,5 | **high** | Planctomycetota | — | — | — | — | Planctomycetota bacterium | SAMN11533026 | terrestrial | 9 | Yes | 4 | HEAT_2;DNA_pol3_alpha;Beta-pro | — |
| 2094028.SAMN13893733.JAAYEJ0100 | **typeXIII** | 219,1 | **low** | Planctomycetota | Candidatus Brocadiia | Candidatus Brocadiales | Candidatus Brocadiaceae | — | Candidatus Brocadiaceae bacterium | SAMN13893733 | temperature | 7 | Yes | 4 | Lipase_GDSL_2;DUF4011;Creatini | — |
| 1129257.SAMEA6153040.CADCRA010 | **typeXIII** | 218,6 | **low** | Bacteroidota | Bacteroidia | — | — | — | uncultured Bacteroidia bacterium | SAMEA6153040 | host-associated | 10 | Yes | 4 | FtsK_SpoIIIE;TIR_2 | — |
| 2026780.SAMN09639622.DSWA01000 | **typeXIII** | 217,7 | **low** | Planctomycetota | — | — | — | — | Planctomycetota bacterium | SAMN09639622 | temperature | 10 | Yes | 4 | GH123_N;SGL;Alpha_L_fucos;GH | — |
| 2026779.SAMN13893644.JAAYAY0100 | **typeXIII** | 215,8 | **low** | Planctomycetota | Planctomycetia | Planctomycetales | Planctomycetaceae | — | Planctomycetaceae bacterium | SAMN13893644 | temperature | 6 | Yes | 3 | TGT;RNA_pol_A_CTD | — |
| 2026779.SAMN14914873.JABIHZ0100 | **typeXIII** | 213,7 | **low** | Planctomycetota | Planctomycetia | Planctomycetales | Planctomycetaceae | — | Planctomycetaceae bacterium | SAMN14914873 | aquatic | 9 | Yes | 4 | Ldh_2;SGL;Lum_binding;DUF6009 | — |
| 156588.SAMEA9695016.CAJXYV01000 | **typeXIII** | 211,3 | **low** | Verrucomicrobiota | — | — | — | — | uncultured Verrucomicrobiota bacteriu | SAMEA9695016 | anthropogenic | 10 | Yes | 4 | Metallophos;AAA_23;DUF3293;Ser | — |
| 2840714.SAMN15817152.DVNY010001 | **typeXIII** | 209,5 | **low** | Bacteroidota | Bacteroidia | Bacteroidales | Prevotellaceae | Candidatus Caccomon | Candidatus Caccomonas pullistercoris | SAMN15817152 | host-associated | 6 | Yes | 3 | DUF7017;AAA_30 | — |
| 1262935.PRJEB997.HF989231_22 | **typeXIII** | 207,6 | **low** | Bacteroidota | Bacteroidia | Bacteroidales | Prevotellaceae | Prevotella | Prevotella sp. CAG:755 | PRJEB997 | — | 10 | Yes | 0 | — | — |
| 2026778.SAMN13893681.JAAYCJ0100 | **typeXIII** | 201,5 | **low** | Planctomycetota | Phycisphaerae | — | — | — | Phycisphaerae bacterium | SAMN13893681 | temperature | 1 | Yes | 1 | URO-D | — |
| 2723666.SAMN14513815.CP050848_3 | **typeXIII** | 193,7 | **low** | Planctomycetota | Planctomycetia | Planctomycetales | — | — | Planctomycetales bacterium ZRK34 | SAMN14513815 | aquatic | 10 | Yes | 4 | Secretin;CPSase_sm_chain;MgtE; | — |
| 2026780.SAMN09724347.QWPT010000 | **typeXIII** | 183,5 | **low** | Planctomycetota | — | — | — | — | Planctomycetota bacterium | SAMN09724347 | temperature | 10 | Yes | 4 | AAA_5;DUF2201;LTI65_LTI78_NY | — |
| 286133.SAMEA104667185.OMXR0100 | **typeXIII** | 160,8 | **low** | Pseudomonadota | Betaproteobacteria | Burkholderiales | Sutterellaceae | Sutterella | uncultured Sutterella sp. | SAMEA104667185 | host-associated | 10 | Yes | 4 | AAA_5;DUF2201_N;N6_N4_Mtase | — |
| 2021391.SAMN07620311.PBEC010000 | **typeXIII** | 158 | **low** | Bacteroidota | Flavobacteriia | Flavobacteriales | — | — | Flavobacteriales bacterium | SAMN07620311 | aquatic | 5 | Yes | 2 | iREC;HisKin-conflict | — |
| 40545.SAMN17800730.JAGZDF010000 | **typeXIII** | 149,6 | **low** | Pseudomonadota | Betaproteobacteria | Burkholderiales | Sutterellaceae | Sutterella | Sutterella wadsworthensis | SAMN17800730 | host-associated | 10 | Yes | 4 | PSF2_N;SNF2-rel_dom;Radical_S | — |
| **TOTAL: 1,286 candidates \| 934 high \| 25 borderline \| 327 low** | | | | | | | | | | | | | | | | |
