## Supplementary Table S3 for "Landscape of retron diversity across the SPIRE prokaryotic metagenome resource reveals candidate novel type XI-like lineages"

**Supplementary Table S3. Systematic evidence review of all 28 SPIRE retron classification groups. For each group, five evaluation criteria are scored: absent (−), weak (+), moderate (++), strong (+++). Scores were assigned using uniform quantitative thresholds applied across all groups. The composite novelty priority score (none/low/medium/high) integrates all five criteria. Groups marked as 'known-type expansion' were excluded from the curated evidence matrix (Figure 3A) as they displayed fully canonical characteristics providing no discriminatory information.**

| Retron group | N total | N high (%) | N border (%) | N low (%) | Top phylum | Top biome | RT phylogenetic coherence | Accessory architecture | Confidence support | Taxonomic coherence | Ecological coherence | Novelty priority | Review classification |
|---|---|---|---|---|---|---|---|---|---|---|---|---|---|
| **typeXI** | 232 | 65 (28%) | 0 (0%) | 167 (72%) | Bacteroidota | host-associated | +++ | +++ | ++ | +++ | ++ | high | **candidate Type XI-like lineage** |
| **clade2_Ec107like** | 192 | 188 (98%) | 0 (0%) | 4 (2%) | Bacillota | host-associated | ++ | − | +++ | ++ | ++ | low | known-like/minimal expansion |
| **typeXIII** | 143 | 129 (90%) | 0 (0%) | 14 (10%) | Bacillota | host-associated | ++ | ++ | +++ | +++ | ++ | medium | known-type expansion |
| **typeVI** | 93 | 90 (97%) | 0 (0%) | 3 (3%) | Bacteroidota | host-associated | — | — | — | — | — | — | known-type expansion |
| **clade11-A** | 71 | 40 (56%) | 0 (0%) | 31 (44%) | Pseudomonadota | host-associated | − | − | + | − | − | none | heterogeneous/low priority |
| **typeIII-A5** | 70 | 32 (46%) | 0 (0%) | 38 (54%) | Bacillota | host-associated | + | ++ | + | + | + | low | known-type expansion |
| **typeIII-A1** | 50 | 43 (86%) | 0 (0%) | 7 (14%) | Bacillota | host-associated | — | — | — | — | — | — | known-type expansion |
| **typeII-A1** | 47 | 46 (98%) | 0 (0%) | 1 (2%) | Pseudomonadota | host-associated | — | — | — | — | — | — | known-type expansion |
| **typeIII-A2** | 42 | 30 (71%) | 0 (0%) | 12 (29%) | Bacteroidota | host-associated | — | — | — | — | — | — | known-type expansion |
| **typeI-C1** | 44 | 44 (100%) | 0 (0%) | 0 (0%) | Pseudomonadota | host-associated | — | — | — | — | — | — | known-type expansion |
| **typeIII-A3** | 40 | 40 (100%) | 0 (0%) | 0 (0%) | Pseudomonadota | aquatic | — | — | — | — | — | — | known-type expansion |
| **clade11-B** | 39 | 18 (46%) | 0 (0%) | 21 (54%) | Actinomycetota | aquatic | − | − | + | + | + | none | heterogeneous/low priority |
| **typeI-C2** | 31 | 31 (100%) | 0 (0%) | 0 (0%) | Bacillota | host-associated | — | — | — | — | — | — | known-type expansion |
| **typeII-A3** | 28 | 27 (96%) | 0 (0%) | 1 (4%) | Bacillota | host-associated | — | — | — | — | — | — | known-type expansion |
| **typeI-B2** | 21 | 9 (43%) | 12 (57%) | 0 (0%) | Bacillota | host-associated | ++ | +++ | + | ++ | + | medium | known-type expansion |
| **typeIII-A4** | 21 | 19 (90%) | 2 (10%) | 0 (0%) | Bacteroidota | host-associated | — | — | — | — | — | — | known-type expansion |
| **typeI-A** | 21 | 18 (86%) | 0 (0%) | 3 (14%) | Pseudomonadota | host-associated | — | — | — | — | — | — | known-type expansion |
| **typeXII** | 21 | 15 (71%) | 0 (0%) | 6 (29%) | Pseudomonadota | host-associated | — | — | — | — | — | — | known-type expansion |
| **typeX** | 16 | 6 (38%) | 3 (19%) | 7 (44%) | Planctomycetota | aquatic | + | + | + | ++ | +++ | none | low-count known group |
| **typeI-C3** | 14 | 3 (21%) | 2 (14%) | 9 (64%) | Bacteroidota | aquatic | + | ++ | + | ++ | + | none | low-count known group |
| **typeV** | 11 | 10 (91%) | 1 (9%) | 0 (0%) | Pseudomonadota | anthropogenic | + | +++ | +++ | +++ | + | none | low-count known group |
| **clade11** | 9 | 9 (100%) | 0 (0%) | 0 (0%) | Actinomycetota | aquatic | — | — | — | — | — | — | known-type expansion |
| **typeVII-A1** | 7 | 6 (86%) | 1 (14%) | 0 (0%) | Pseudomonadota | anthropogenic | — | — | — | — | — | — | known-type expansion |
| **typeII-A2** | 7 | 3 (43%) | 3 (43%) | 1 (14%) | Pseudomonadota | aquatic | — | — | — | — | — | — | known-type expansion |
| **typeIX** | 5 | 5 (100%) | 0 (0%) | 0 (0%) | Pseudomonadota | aquatic | — | — | — | — | — | — | known-type expansion |
| **typeIV** | 5 | 5 (100%) | 0 (0%) | 0 (0%) | Pseudomonadota | aquatic | — | — | — | — | — | — | known-type expansion |
| **typeVII-A2** | 4 | 1 (25%) | 1 (25%) | 2 (50%) | Pseudomonadota | anthropogenic | — | — | — | — | — | — | known-type expansion |
| **typeI-B1** | 2 | 2 (100%) | 0 (0%) | 0 (0%) | Pseudomonadota | aquatic | — | — | — | — | — | — | known-type expansion |

*Score legend: − (absent) = no evidence; + (weak) = marginal evidence; ++ (moderate) = consistent evidence; +++ (strong) = robust evidence across multiple independent indicators*

*Note: Groups classified as 'known-type expansion' were not included in the curated evidence matrix (Figure 3A) because they displayed fully canonical characteristics across all criteria, providing no discriminatory information for novelty assessment. Scores marked '—' indicate groups for which detailed criterion-level scoring was not performed as part of the curated review.*
